# Prediction of the reaction mechanisms of sesquiterpene coumarin synthases supports a direct evolutionary link with triterpene biosynthesis

**DOI:** 10.1101/2024.10.24.619981

**Authors:** Michael J. Stephenson, Peter E. Brodelius

## Abstract

Sesquiterpene coumarins are primarily produced by plants of the Apiaceae and Asteraceae families. Farnesylation of 7-hydroxycoumarins such as umbelliferone, scopoletin or isofraxidin yield linear 7-farnesyloxycoumarins that are converted to various cyclic sesquiterpene coumarins by sesquiterpene coumarin synthases (cyclases). The terminal double bond of the linear 7-farnesyloxycoumarins is epoxidized by a sesquiterpene coumarin epoxidase. The diverse 7-(10′,11′-oxidofarnesyloxy)-coumarins produced are protonated by various sesquiterpene coumarin synthases to generate a carbocation that initiates cyclization of the farnesyl moiety (A process analogous to the carbocation cascades observed with sesquiterpene synthases and other cyclases involved in the biosynthesis of additional terpene classes, such as the triterpenes). These reaction mechanisms typically include Wagner-Meerwein rearrangements, such as hydride, methyl, and other alkyl shifts, but can also involve more complex processes including Grob fragmentations. Around 260 sesquiterpene coumarins based on 7-farnesyloxycoumarins have been described, but essentially nothing is known about the biosynthetic enzymes involved, *i.e*., farnesyltransferase, sesquiterpene coumarin epoxidase and synthase. In this review, putative reaction pathways for formation of the carbon skeletons of all known 7-farnesyloxycoumarins-derived sesquiterpene coumarins are presented.

## 1. Introduction

Sesquiterpene coumarins are a family of plant secondary metabolites characterized by the inclusion of both a sesquiterpene and a coumarin moiety in their structures (as illustrated in Figure 1). These natural products have been shown to exhibit a range of biological activities, many of which have potential pharmaceutical applications [Gliszczynska & Brodelius, 2012; Li et al, 2018]. The common coumarin building blocks observed in these natural products are shown in Figure 2. In total, around 370 sesquiterpene coumarins have been isolated and at least partly characterized from various plant species. Most have been isolated from plants belonging to just a few genera of the Apiaceae (Umbelliferae) and Asteraceae (Compositae) families, with the richest source being the genus *Ferula* (Apiaceae). The structures of the sesquiterpene coumarins can be divided into two main classes distinguished by the chemical nature of the linkage between the coumarin and sesquiterpene fragments. Both ether and carbon-carbon linkages are observed. Here, we define these as *O*- and *C*-prenylated sesquiterpene coumarins, respectively. This review, which is part one of two reviews, will cover *O*-prenylated sesquiterpene coumarins. In part two, *C*-prenylated sesquiterpene coumarins will be discussed.

**Figure 1.**
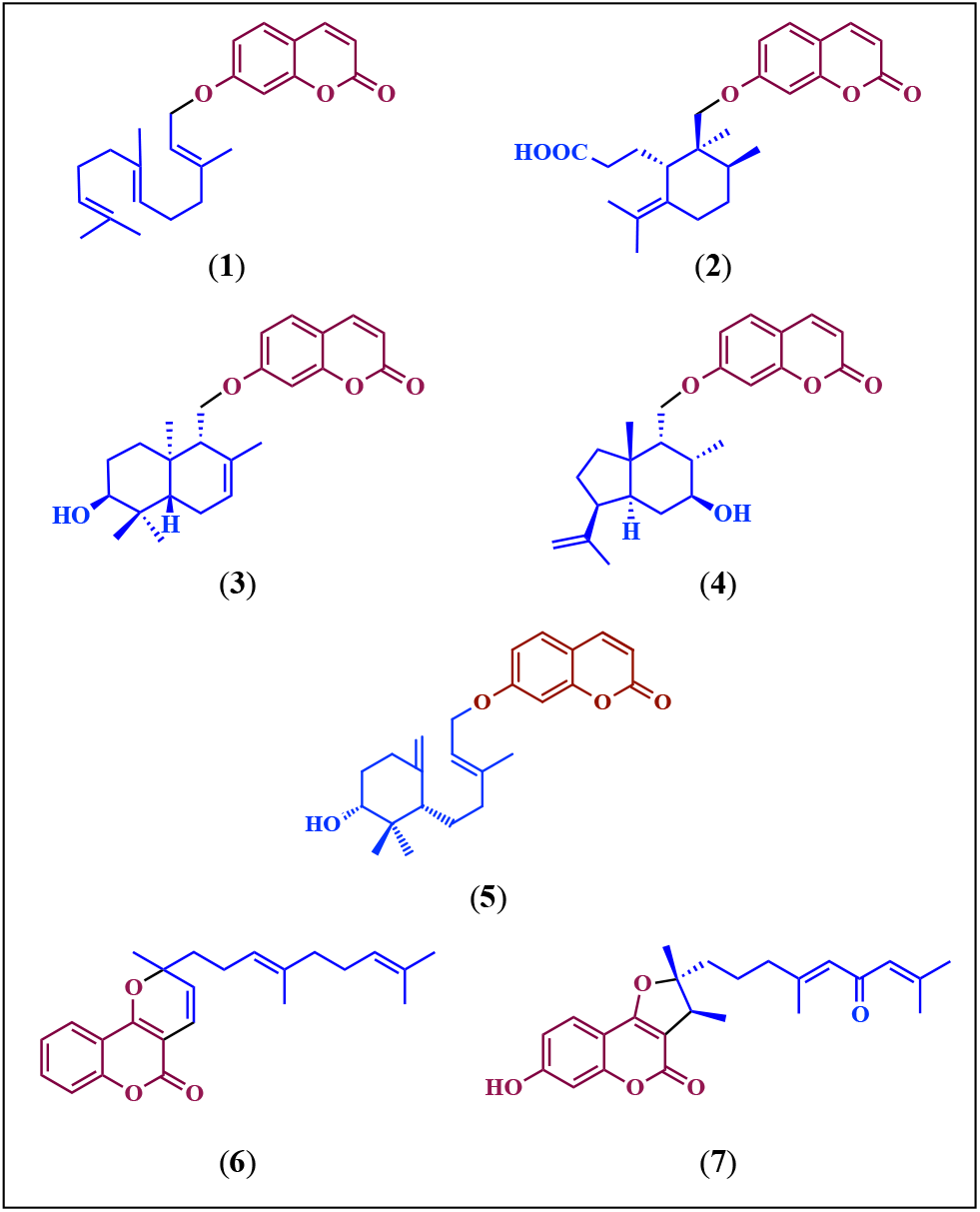
Examples of the structure of some sesquiterpene coumarins. (**1)** to (**5)** are examples of *O*-farnesylated umbelliferone and (**6)** to (**7)** are examples of *C*-farnesylated 4-hydroxycoumarin. (**1**) umbelliprenin; (**2**) galbanic acid; (**3**) conferol; (**4**) ferusinol; (**5**) farnesiferol B; (**6**) ferprenin; (**7**) fukanefuromarin A. Coumarin moieties are in brown and sesquiterpene moieties in blue.

**Figure 2.**
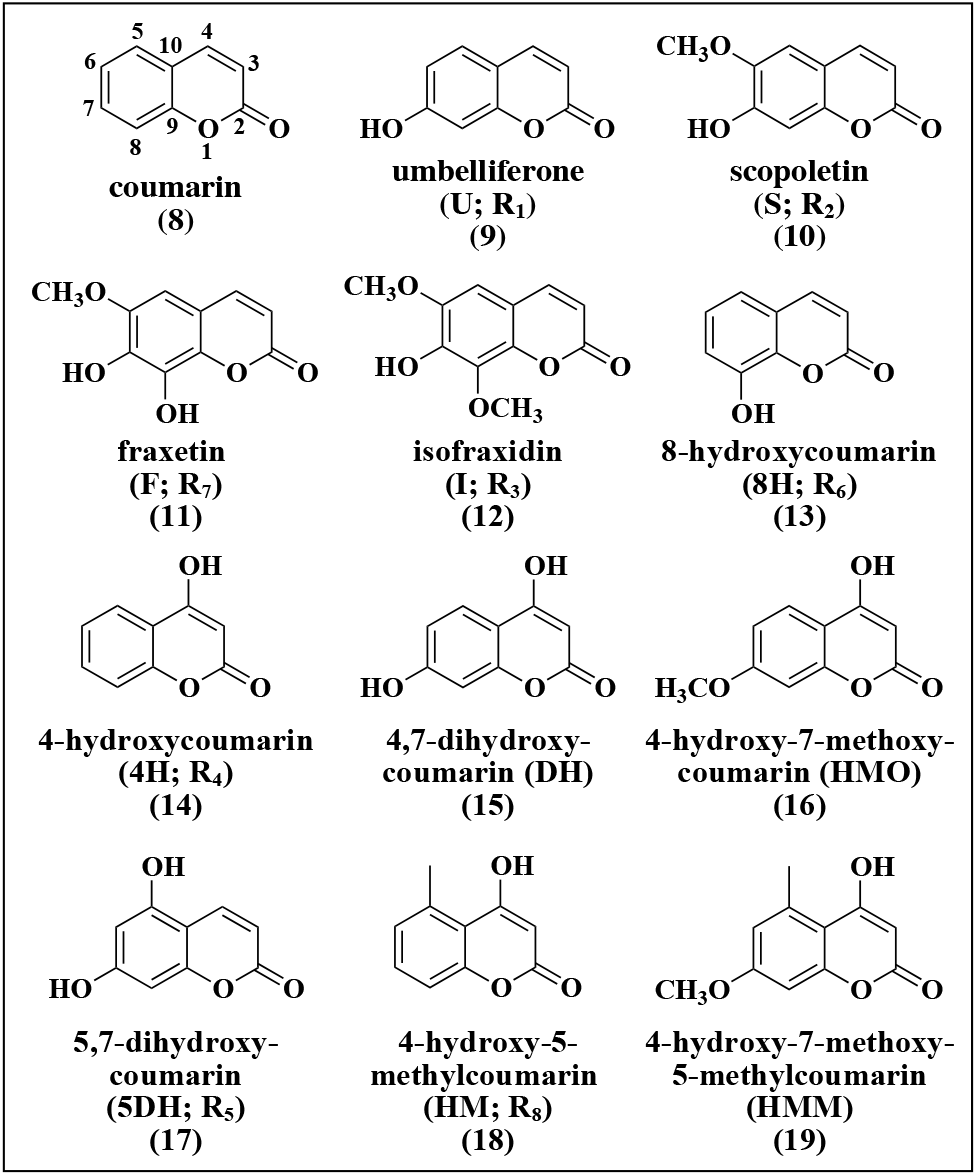
Structure of coumarins found as building blocks of sesquiterpene coumarins. Abbreviations used are shown.

The individual structures of 277 *O*-prenylated sesquiterpene coumarins are shown in Figures S2 to S9. All *O*-prenylated sesquiterpene coumarins (except for only a handful of examples) have been isolated from plants belonging to the Apiaceae and Asteraceae families (as summarized in supplementary Table S1). In total, 79,2 % of the *O*-prenylated sesquiterpene coumarins have been isolated from Apiaceae plants, with the majority coming from the genus *Ferula* while 17,3% of the *O*-prenylated sesquiterpene coumarins have been isolated from plants belonging to the Asteraceae family (Table 1). The most common coumarin moiety of the Apiaceae sesquiterpene coumarins is umbelliferone (**9**), but 4-hydroxycoumarin (**14**) and 5,7-dihydroxycoumarin (**17**) have also been found, as summarized in Table 1. The coumarin moiety of the Asteraceae sesquiterpene coumarins is usually scopoletin (**10**) or isofraxidin (**12**). However, sesquiterpene coumarins containing umbelliferone (**9**) have also been isolated from 13 of 42 studied *Artemisia* species (Asteraceae) [Al-Hazimi and Basha, 1991].

**Table 1.** Number of isolated sesquiterpene coumarins with different coumarin moieties from different plant families.

| coumarin | Apiaceae | Asteraceae | Rutaceae | Euphorbiaceae | Amaranthaceae | Lythraceae | Myrtaceae | Solanaceae | Total |  |
| --- | --- | --- | --- | --- | --- | --- | --- | --- | --- | --- |
| number of species | 72<sup>§</sup> | 21 | 2 | 2 | 3 | 1 | 1 | 1 | 103 | 100% |
| umbelliferone (9) | 242 | 14 | 1<sup>#</sup> |  | 3<sup>#</sup> | 1<sup>#</sup> | 1<sup>#</sup> | 1<sup>#</sup> | 263 | 84% |
| scopoletin (10) |  | 4 |  | 1 |  |  |  |  | 5 | 1.6% |
| fraxetin (11) |  |  |  | 1* |  |  |  |  | 1 | 0.3% |
| isofraxidin (12) |  | 31 |  |  |  |  |  |  | 31 | 9.9% |
| 8-hydroxycoumarin (13) |  |  | 2* |  |  |  |  |  | 2 | 0.6% |
| 4-hydroxycoumarin (14) | 3 |  |  |  |  |  |  |  | 3 | 1.0% |
| 5,7-dihydroxycoumarin (17) | 3 |  |  |  |  |  |  |  | 3 | 1.0% |
| 4-hydroxy-5-methylcoumarin (18) |  | 5 |  |  |  |  |  |  | 5 | 1.6% |
| **Total** | **248** | **54** | **3** | **2** | **3** | **1** | **1** | **1** | **313** |  |
|  | 79.2% | 17.3% | 1.0% | 0.7% | 1.0 % | 0.3% | 0.3% | 0.3% |  |  |

<sup>§</sup>Including 51 *Ferula* species. There are 204 accepted *Ferula* species [http://www.theplantlist.org/browse/A/Apiaceae/Ferula/]. Most of these, if not all, produce sesquiterpene coumarins.
\*These sesquiterpene coumarins are formed by coupling a preformed sesquiterpene to the coumarin moiety.
<sup>#</sup>These sesquiterpene coumarins are all the linear umbelliprenin

The different carbon skeletons of the sesquiterpene moiety are summarized in Figure 3. Sesquiterpene coumarins with the same carbon skeleton have been grouped together. Their individual structures are shown in supplementary Figures S2 to S9. The structures of all sesquiterpene coumarins are presented and labeled in a standardized style as shown in supplementary Figure S1. The code in the upper left corner gives the following: prenylation type (in this case an *O* for *O*-prenylation), coumarin moiety (abbreviations are given in Figure 2), designated carbon skeleton of the sesquiterpene moiety and finally the identification number. For instance, the code for umbelliprenin (**1**) is ***O*-U-A1** (*O*-prenylation, umbelliferone (U) (**9**), carbon skeleton A and identification number 1). In the upper right corner a reference is given. Finally, the trivial name is provided at the bottom. The major carbon skeletons are ***O*-A, *O*-Ha, *O*-Ia**, and ***O*-Ja** with 35, 38, 62 and 42 compounds containing these scaffolds, respectively.

**Figure 3.**
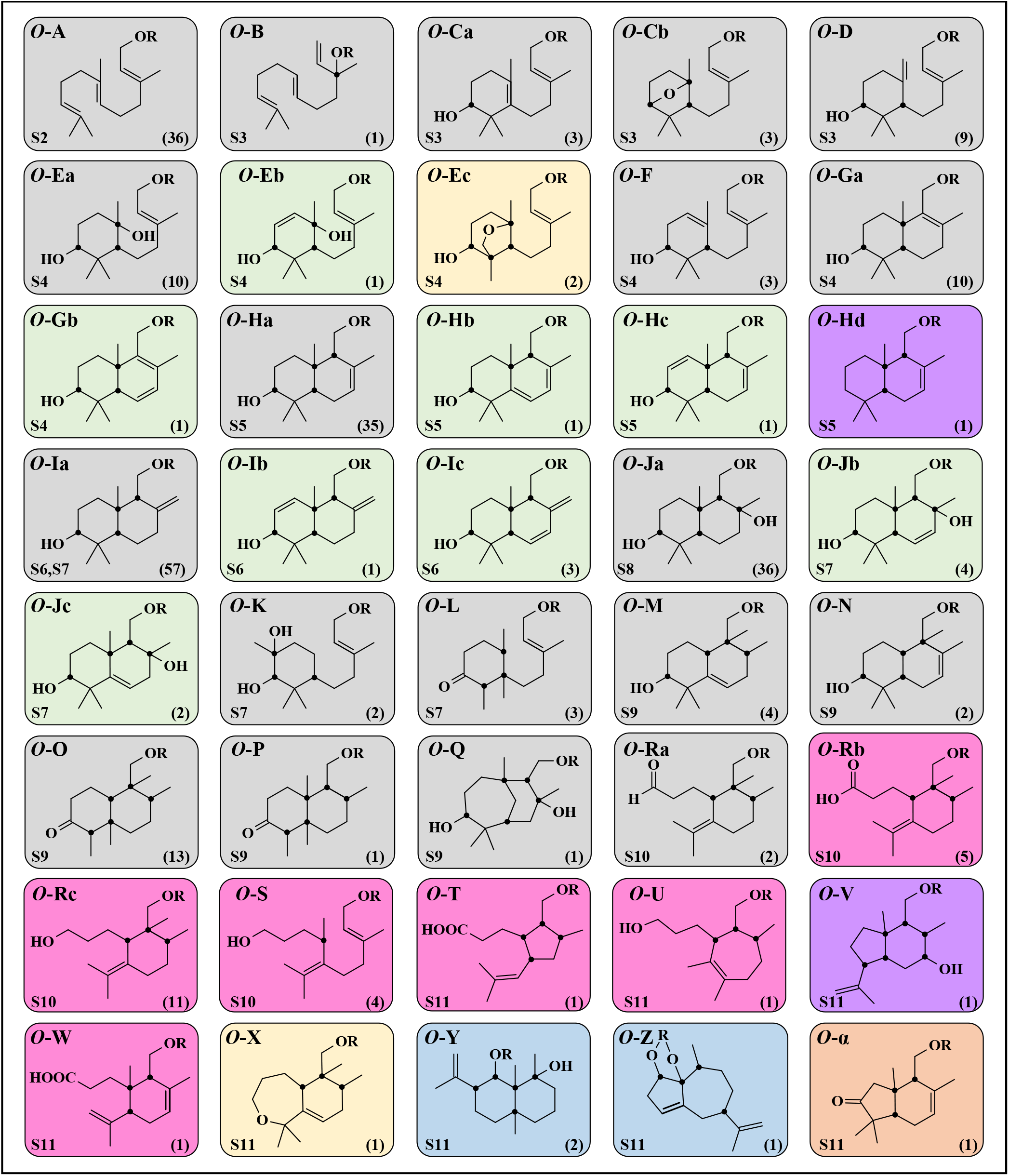
Carbon skeletons of isolated *O*-prenylated sesquiterpene coumarins. The colour coding of the boxes is based on the putative pathways presented in this review. Carbon skeletons obtained after cyclization without alterations are shown in grey boxes. Carbon skeletons obtained after protonation of a double bond is shown in violet boxes. Carbon skeletons obtained after desaturation are shown in green boxes. Carbon skeletons obtained after oxidation/reduction by oxidoreductases are shown in magenta boxes. Carbon skeletons obtained by biotransformation of sesquiterpene coumarins are shown in yellow boxes. Carbon skeletons preformed before coupling to coumarins are shown in blue boxes. The carbon skeleton obtained after decarboxylation is shown in orange box. The numbers in the lower right corner show the number of sesquiterpene coumarins with that carbon skeletons and the number shown in the lower left corner refer to the supplementary figure in which the structure of all the sesquiterpene coumarins with that carbon skeleton are shown. Black dots indicate chimeric carbons.

Although, the individual biogenic origins of coumarins and sesquiterpenes are well-understood, very little is known about the enzymatic genesis of sesquiterpene coumarins. It is well-established that cyclic sesquiterpenes are produced by sesquiterpene synthases (STSs) from linear substrates using carbocationic reaction mechanisms [Christianson, 2017]. However, no enzymes involved in the biosynthesis of sesquiterpene coumarins have so far been identified. The formation of these products could be predicted to arise from either, the conjugation of preformed cyclic sesquiterpene and coumarin building blocks, or through the cyclisation and rearrangement of linear sesquiterpene precursors after conjugation to the coumarin moiety. The presence of simple linear sesquiterpene coumarins, such as umbelliprenin (**1**), in species also producing more complex cyclic variants, could be seen to support the latter hypothesis. In this review, we present the structures of all known *O*-prenylated sesquiterpene coumarins in the context of the proposal that simple linear variants serve as common precursors acting as substrates for hypothesized sesquiterpene coumarin synthases (STCS). To illustrate, we provide putative carbocation reaction pathways that could be invoked to access the diversity of known *O*-prenylated sesquiterpene coumarins observed. We also highlight similarities and draw parallels with triterpene biosynthesis and suggest that these STCSs could share an evolutionary origin more closely related to triterpene synthases (oxidosqualene cyclases) than sesquiterpene synthases; perhaps even arising from duplication and neofunctionalization of triterpene synthases in the Apiaceae family, and that prospecting for STCSs in divergent Apiaceae triterpene synthases genes could yield their discovery.

## 2. Terpene synthases

A brief review of terpene synthases (TPSs) will give the basis for our putative mechanisms of the formation of the sesquiterpene moiety of sesquiterpene coumarins. Carbocationic reaction mechanisms are used by TPSs. The reaction is initiated by ionization of the substrate with the formation of a carbocationic intermediate. This intermediate undergoes a series of cyclizations, hydride shifts and/or other rearrangements (such as methyl and other alkyl shifts) until the reaction is terminated by proton abstraction or the addition of a nucleophile such as water. Generally, 1,2- and 1,3-hydride shifts are allowed. In addition to the 1,2- and 1,3-hydride shifts, some terpene synthases catalyze long-range hydride transfers (1,4- and 1,5-hydride shifts). However, these long-range migrations are rarely observed [Williams et al, 2000; Meguro et al, 2015; Driller et al, 2018; Chiba et al, 2013]. Only 1,2-alkyl shifts are generally allowed. Such an alkyl shift can result in a change of ring size (both ring contractions and ring expansions are observed) or the translocation of a methyl group depending on the migrating bond. Recently it has been shown that 1,3-methyl shifts can occur [Meguro et al, 2015; Driller et al, 2018]. Despite utilizing the same substrate and exhibiting significant sequence and structural homology, terpene synthases form a large group of products with different carbon skeletons.

Many TPSs have been isolated and characterized [Christianson, 2017; Karunanithi & Zerbe, 2019]. These enzymes are the key to the enormous structural diversity of terpenoids. TPSs are traditionally divided into two categories depending on how the initial carbocation is generated. In type I TPSs, the diphosphate group of the prenyl diphosphate is abstracted resulting in an allylic carbocation on the prenyl substrate, which initiates a carbocation cascade reaction. In type II TPSs, the initial carbocation is formed through protonation of a double bond or an epoxide group. In the latter case, a hydroxyl group is left on the product.

The key catalytic amino acid residues in most type I TPSs are a DDXXD-motif and a N/DxxS/TxxxE-motif. The aspartate residues (D) of these motifs bind divalent metal ions, typically Mg^2+^, which coordinate the diphosphate of the substrate and provide the electrophilic driving force for ionization. In most type II TPSs, a DXDD-motif is involved in protonation of the substrate. However, this motif is unrelated to the aspartate-rich DDXXD-motif of class I TPSs [Christianson, 2017].

Canonical TPSs are also classified according to structural features of the protein. Phylogenetic analyses of TPS protein sequences from gymnosperms and angiosperms recognized seven major clades (or subfamilies), designated TPSs *a* through TPSs *g* [Bohlmann et al, 1998]. An updated analysis including TPS sequences from the sequenced genomes of several plant species also recognized seven TPS subfamilies - the original *a, b, c, d*, and *g*, a merged clade of the original *e* and *f* subfamilies (designated as *e/f*), and a new subfamily *h* [Chen et al, 2011].

Other TPSs are classified as non-canonical. TPSs involved in the biosynthesis of meroterpenoids belong to this class of TPSs. Meroterpenoids are characterized by a terpenoid structure linked to another type of natural compound, *e.g*., a phenolic structure or an alkaloid. These TPSs can be of type I or type II but may lack the characteristic aspartate-rich motif. TPSs involved in the biosynthesis of sesquiterpene coumarins would belong to type II of this group.

In Figure 4A, two well-characterized reaction mechanism involving type II terpene synthases are shown. In both examples the starting substrate is squalene (**20**). For lanosterol (**24**) biosynthesis, squalene epoxidase (SQE) produces 2,3-oxidosqualene (**22**), which is protonated and converted to lanosterol (**24**) and lupeol (**26**) by lanosterol synthase (LAS) and lupeol synthase (LUP1) respectively. In the second example, the terminal double bond of squalene (**20**) is pronated and converted to hopene (**21**) by squalene hopene cyclase (SHC). We suggest that similar reactions catalyzed by type II terpene synthases are involved in the biosynthesis of sesquiterpene coumarins as exemplified by the putative biosynthetic pathway for the sesquiterpene coumarin conferol (**3**) from umbelliferone (**9**) and farnesyl diphosphate (**27**) as shown in Figure 4B. First, umbelliferone (**9**) is farnesylated by *O*-farnesyltransferase (*O*-FT) to yield umbelliprenin (**1**), which is converted to 10′,11′-oxidoumbelliprenin (**28**) by umbelliprenin monooxidase (UMO). Next protonation and cyclization of 10′,11′-oxidoumbelliprenin (**28**) by a STCS gives the sesquiterpene coumarin conferol (**3**). During cyclization a monocyclic carbocation, intermediate carbocation I (**29**) and a bicyclic carbocation, intermediate carbocation II (**30**) are formed. Observe the different numberings of linear, monocyclic and the bicyclic sesquiterpene coumarins. We have chosen to number the sesquiterpene coumarins in analogy to the numbering of monocyclic and bicyclic triterpenes. Despite utilizing the same substrate and exhibiting significant sequence and structural homology, terpene synthases form a large group of products with different carbon skeletons (Figure 3). These two cyclic carbocations (**29, 30**) are common for the biosynthesis of a number of sesquiterpene coumarins. In the following, various putative biosynthetic pathways catalyzed by different STCSs will start from one of these intermediate carbocations. In order to emphasize that different STCSs are involved in the biosynthesis of different carbon skeletons of the sesquiterpene moiety, the enzyme abbreviation STCS-Xx (Xx = carbon skeleton of the product) is used.

**Figure 4.**
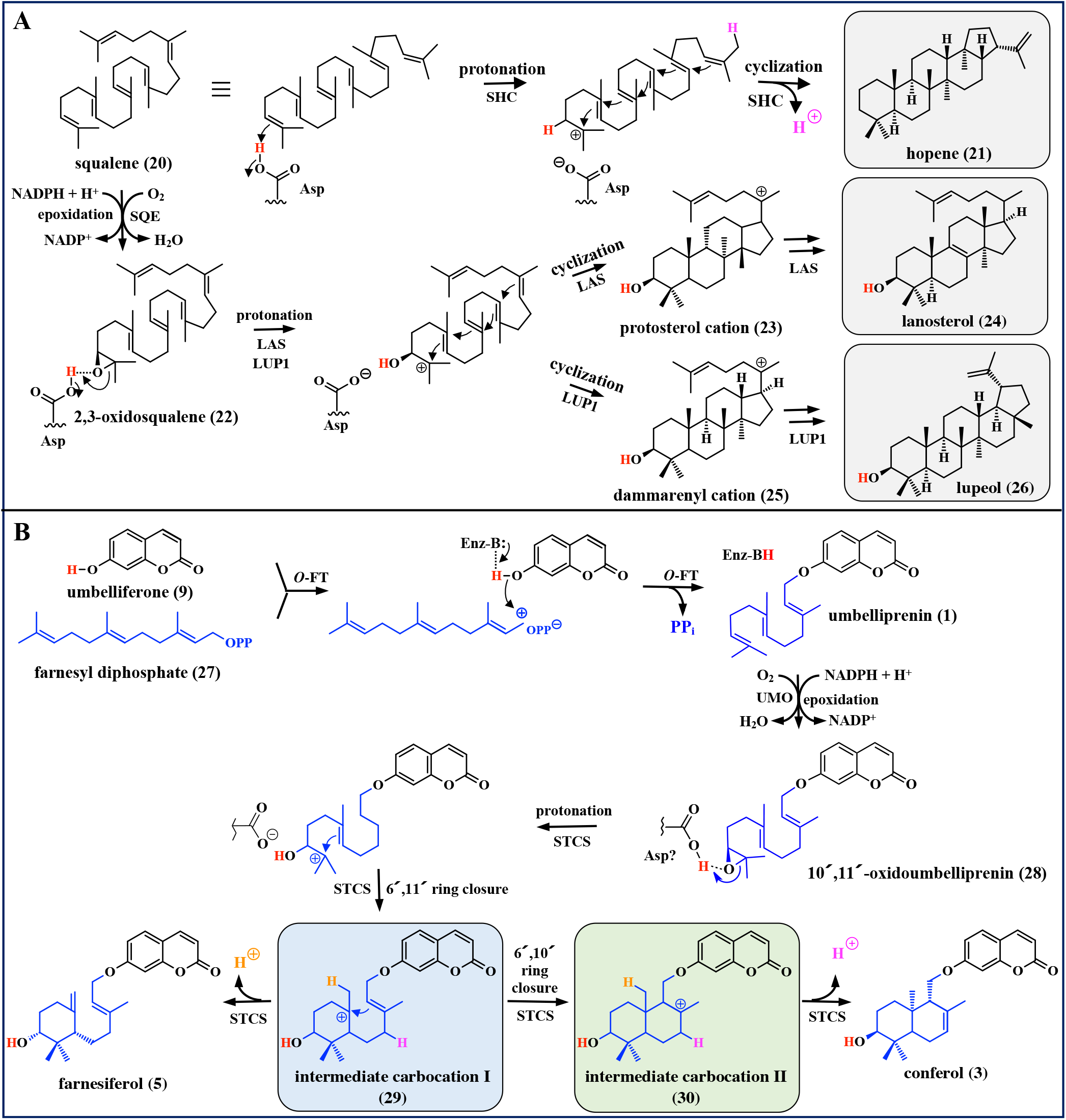
(**A**) Biosynthesis of hopene (**21**) from squalene catalyzed by squalene hopene cyclase (SHC) and biosynthesis of lanosterol (**24**) and lupeol (**26**) from squalene (**20**) catalyzed by squalene epoxidase (SQE) and lanosterol/lupeol synthase (LAS/LUP1). (**B**) Putative biosynthetic pathways for the sesquiterpene coumarins conferol (**3**) and farnesiferol B (**5**) from umbelliferone (**9**) and farnesyl diphosphate (**27**) catalyzed by *O*-farnesyltransferase (O-FT), umbelliprenin monooxygenase (UMO) and sesquiterpene coumarin synthases (STCS). The boxed intermediate carbocations (**29** and **30**) are intermediates in several putative biosynthetic pathways of sesquiterpene coumarins.

## 3. Sesquiterpene Coumarin Biosynthesis

Sesquiterpene coumarins are formed by farnesylation/nerolidylation of different coumarins followed by modifications of the prenylated product obtained. Biochemical studies on the biosynthesis of sesquiterpene coumarins are very limited. However, our knowledge of terpene biosynthesis involving canonical TPSs and the prenylation of natural products will be used to suggest putative biosynthetic routes for these lesser studied meroterpenoids. Please note that in proceeding figures, for conciseness, some sequences of 1,2-shifts are depicted as concerted when in reality individual transition states and intermediary cations would be predicted to be required. For example, the sequential suprafacial migration of synperiplanar bonds.

### 3.1 Biosynthesis of farnesyl diphosphate

The universal five-carbon terpene building blocks isopentenyl diphosphate (IDP) and its isomer dimethylallyl diphosphate (DMADP) are produced in plants by two different pathways. The classical mevalonic acid (MVA) pathway, which produces IDP, is localized partly in the cytosol and partly in peroxisomes, which harbor the last enzymes of the pathway. IDP is enzymatically isomerized to DMADP. These C_5_ terpenoid diphosphates serve as substrates for peroxisomal farnesyl diphosphate synthase to form FDP (**27**), which is used to produce C_15_ sesquiterpenes directly and other terpenoids, such as C_30_ triterpenes following further condensation to a squalene precursor. The plastidic methylerythritol phosphate (MEP) pathway produces both IDP and DMADP for the formation of geranyl diphosphate (GDP) and geranylgeranyl diphosphate (GGDP) by the plastidic enzymes GDP and GGDP synthase, respectively. GDP and GGDP are used for mono- and diterpene biosynthesis, respectively.

Canonical sesquiterpene synthases (STSs) convert FDP (**27**) to C_15_ terpenoids. Different STSs produce sesquiterpenes of differing carbon skeletons from this common linear substrate. Around 300 different carbon skeletons have been described for sesquiterpenes [Fidan & Zhan, 2018].

### 3.2 Biosynthesis of coumarins

More than 1300 natural coumarins from plants, fungi and bacteria have been described [Matos et al, 2015]. Most of these have been isolated from about 150 plant species distributed over approximately 30 different families [Sharifi-Rad et al, 2021]. The function of coumarins is not fully known. Many coumarins are induced by various abiotic and biotic stresses indicting that they are plant defense substances. A few coumarins are building blocks of sesquiterpene coumarins (Figure 2). The biosynthesis of some of these coumarins has been established and will be described next.

Intermediates of the phenylpropanoid pathway are used in the biosynthesis of 7-hydroxycoumarins such as umbelliferone (**9**), scopoletin (**10**), fraxetin (**11**), and isofraxidin (**12**), which are the coumarin moiety of 95.3 % of *O*-prenylated sesquiterpene coumarins (Table 1). The CoA-thioesters of coumaric, ferulic and sinapic acids are hydroxylated at the *ortho* position by coumaroyl-2′-hydroxylase (C2′H) or feruloyl-6′-hydroxylase [Shimizu, 2014]. *Trans*–*cis* isomerization of the hydroxylated CoA esters is induced by UV-light, which leads to spontaneous lactonization and formation of the corresponding coumarins [Edwards et al, 1967].

Recently, an enzyme (coumarin synthase; COSY) catalyzing the conversion of *ortho*-hydroxycoumaroyl CoAs to the corresponding coumarins has been described [Vanholme et al, 2019]. COSY is able to produce 7-hydroxycoumarins by a two-step reaction. Initially, a *trans*– *cis* isomerization takes place, which is followed by lactonization. COSY is important for the biosynthesis of coumarins in organs that are shielded from light, such as roots [Vanholme et al, 2019].

4-Hydroxycoumarin (**14**) is the building block of a few sesquiterpene coumarins, such as linear and angular furanocoumarins, as well as pyranocoumarins. A biosynthetic pathway for the synthesis of 4-hydroxycoumarin (**14**) has been presented [Liu et al, 2010]. Salicylic acid, which is a precursor, can be synthesized from phenylalanine via cinnamic and benzoic acid or from isochorismate. The salicylic acid is converted to its CoA-ester by salicyloyl-CoA synthase. The final step in the biosynthesis of 4-hydroxycoumarin (**14**) is catalyzed by bisphenyl synthase (BIS) or 4-hydroxycoumarin synthase, which is a type III polyketide synthase [Liu et al, 2010]. A BIS cDNA clone was isolated from an elicited cell culture of *Sorbus aucuparia* (Rosaceae). Recombinant enzyme was produced by heterologous expression of the BIS cDNA in *E. coli*. Assays of the purified BIS showed that the enzyme preferred salicyloyl-CoA as a starter substrate and catalyzed a single decarboxylative condensation with malonyl-CoA to give 4-hydroxycoumarin (**14**). Type III polyketide synthases are involved in the biosynthesis of many plant secondary metabolites of medicinal interest [Abe, 2020].

4-Hydroxy-5-methylcoumarin (**18**) is found in many sesquiterpene coumarins. In contrast, sesquiterpene coumarins utilizing 4-hydroxy-7-methoxy-5-methylcoumarin (**19**) as a building block are rare [Li et al, 2020b]. Type III polyketide synthases are involved in the biosynthesis of these two coumarins. The biosynthesis of 4-hydroxy-5-methylcoumarin (**18**) was suggested based on results obtained in ^14^C-feeding experiments on *Gerbera jamesonii* (Asteraceae) [Inoue et al, 1989]. Similar results were obtained with the ascomycetes *Aspergillus* and *Emericella* producing the coumarin siderin (4,7-dimethoxy-5-methylcoumarin) [Pietiäinen et al, 2016]. *O*-Methylation of the intermediate metabolite 4,7-dihydroxy-5-methylcoumarin by *O*-methyltransferase yields 4-hydroxy-7-methoxy-5-methylcoumarin (**19**), while *O*-methylation of the 4- and 7-hydroxyl groups yields siderin. A type III polyketide synthase involved in the biosynthesis of 4-hydroxy-5-methylcoumarin (**18**) has been cloned from *Gerbera hybrid* (Asteraceae) [Girol et al, 2012]. When this enzyme was heterologously expressed in *Nicotiana benthamiana*, the isolated product was 4,7-dihydroxy-5-methylcoumarin. However, in *G. hybrid* this intermediate is converted to 4-hydroxy-5-methylcoumarin (**18**) by a reductase present in the plant [Girol et al, 2012]. Finally, a type III polyketide synthase from *Hypericum perforatum* (Hypericaceae) involved in hypericine biosynthesis can produce 4,7-dihydroxy-5-methylcoumarin *in vitro* [Karppinen et al, 2008].

## 4. Enzymes of sesquiterpene coumarin biosynthesis

The putative biosynthetic pathway of sesquiterpene coumarins, as proposed here, exhibits striking similarities to the biosynthesis of triterpenes (Figure 4). The first step is prenylation. In both cases, the donor molecule is FDP (**27**) while the acceptor molecule is 7-hydroxycoumarins (*e.g*., umbelliferone (**9**)) and FDP (**27**), respectively. The products formed are 7-farnesyloxycoumarins (*e.g*., umbelliprenin (**1**)) and squalene (**20**). The prenylation is followed by epoxidation to prepare for the cyclization. Both epoxides undergo cyclization reactions after protonation of the epoxy group by either STCSs or 2,3-oxidosqualene cyclases (OSCs).

Other groups of natural meroterpenoids are indoloditerpenes, which are found in filamentous fungi [Saikia et al, 2008; Hou et al, 2022], indolosesquiterpenes, which have been isolated from both plants and fungi [Li et al, 2015b], and sesquiterpene quinone/quinols found in marine sponges, algae, ascidians, coral, fungi, and plants [Tian et al, 2023]. The same enzymatic steps, *i.e*., prenylation, epoxidation, protonation, cyclization, and termination, are involved in the formation of sesquiterpene coumarins, indolosesquiterpenes, indolodi-terpenes, sesquiterpene quinone/quinols, and triterpenes. These steps are catalyzed by prenyltransferases, epoxidases and terpene cyclases (synthases), which will be discussed below.

### 4.1 Step 1: Prenylation

Prenyltransferases (PTs) catalyze the transfer reactions of prenyl moieties from different prenyl donors, such as DMADP (C_5_), GDP (C_10_), FDP (C_15_) (**27**), or GGDP (C_20_), to various acceptors of both low and high molecular weight, including proteins and nucleic acids [Winkelblech et al, 2015]. A large PT group, aromatic PTs, use secondary metabolites including indole alkaloids, flavonoids, coumarins, xanthones, quinones, and naphthalenes as acceptor molecules.

Prenylcoumarins are produced by plant aromatic PTs. These can be grouped into two classes based on the type of prenylation of coumarins. 7-Hydroxycoumarins such as umbelliferone (**9**) (Figure 2) are the most common substrates for *O*-farnesyl/nerolidyl-PTs, and 4-hydroxycoumarin (**14**) (Figure 2) is the most common substrate for *C*-farnesyl/nerolidyl-PTs. Most prenylated coumarins are further modified by cyclizations, hydroxylations, oxidations, esterification, etc.

Plant aromatic PTs belong to the UbiA superfamily of PTs, and they are localized to the outer membrane of the plastid envelope [de Bruijn et al, 2020]. In contrast, aromatic prenyltransferases from fungi and bacteria are soluble. Based on protein amino acid sequences, these enzymes can be divided into two families: p-hydroxybenzoic acid (PHB) PTs and homogentisate (HG) PTs. In the HG PT family, the aspartate-rich motifs are NQxxDxxxD and KDxxDxxGD, whereas for members of the PHB PT family the corresponding motifs are NDxxDxxxD and DKxDDxxxG [Li, 2016; de Bruijn et al, 2020]. Both these aspartate-rich motifs coordinate Mg^2+^ ions, which are involved in the binding, stabilization, and orientation of the diphosphate group of the donor substrate. An electrophilic prenyl carbocation is formed in the same way as seen with either type I TPSs or type II TPSs. In the case of type II TPSs by either protonation of a double bond or an epoxide. The carbocation generated reacts with an aromatic acceptor resulting in a prenylated substrate.

The canonical aromatic PT, belonging to the HG family, is a protein containing a N-terminal transit peptide, 9 transmembrane helices, and two conserved aspartate-rich motifs. These aspartate-rich motifs are found in protein loops 2 and 6, and result in the localization of the active site of the transferase to the cytoplasmic or intramembrane side of the outer envelope membrane of plastids. Examples of canonical aromatic PTs are naringenin-8-dimethylallyltransferase from *Sophora flavescens* [Yazaki et al, 2009], pterocarpen-4-dimethylallyltransferase from *Glycine max* (soybean) [Akashi et al, 2009] and resveratrol-4-dimethylallyltransferase from *Arachis hypogaea* (peanut) [Yang et al, 2018].

Plant aromatic PTs exhibit specificity for donor and acceptor substrates. Reports on the identification, gene cloning and characterization of aromatic *C*-PTs from various plant species, such as soybean (*Glycine max*) [Sukumaran et al, 2018], peanut (*Arachis hypogaea*) [Yang et al, 2018], hop (*Humulus lupulus*) [Ban et al, 2018] and *Artemisia capillaris* [Munakata et al, 2019], have been published.

So far, most characterized plant aromatic PTs perform *C*-prenylations. However, recently one study on aromatic *O*-PT genes from plants was published [Munakata et al, 2021]. Aromatic *O*-PTs from *Angelica keiskei* (Apiacaea) and grapefruit (*Citrus paradisi*) (Rutaceae) were characterized. cDNAs of two *O*-PT genes, *CpPT1* and *AkPT1*, belonging to the UbiA superfamily, were isolated and analyzed. CpPT1 was found to exhibit bergaptol-5-geranyltransferase activity and AkPT1 bergaptol-5-dimethylallyltransferase activity. *In silico* analysis predicted that the proteins CpPT1 and AkPT1 each include a N-terminal transit peptide, two aspartate-rich motifs (NQxxDxxxD and DxxDxxxD) and nine transmembrane regions, all of which are characteristics of plant UbiA PT proteins [Munakata et al, 2021].

The position of the aspartate-rich motifs on the cytosolic side of the outer membrane of the plastid envelope is favorable since both the donor (FDP (**27**)) and the acceptor (coumarin) for sesquiterpene coumarin biosynthesis are produced in the cytosol. It has been suggested that the supply of substrates may be problematic if the aspartate-rich motifs of an aromatic *O*-PT are localized to the intermembrane space of the plastid envelope [Saeki et al, 2018]. The inner membrane of the chloroplast envelope has been recognized as the permeability barrier of the plastid envelope. By contrast, the chloroplast outer envelope membrane has been shown to be permeable to compounds of low molecular weight [Inoue, 2007]. Consequently, it is likely that farnesylation of coumarins can also take place in the intermembrane space utilizing FDP (**27**) from the cytosol. For prenyltransferases using GDP and GGDP as donors, which are produced in the stroma, transport over the inner envelope membrane is required. Such transport systems for the translocation of IDP. DMADP, GDP and GGDP have been predicted [Dudareva et al, 2013; Gutensohn et al, 2013].

The biosynthesis of daurichromenic acid involves a *C*-farnesyltransferase, which converts orsellinic acid to grifolic acid. This *C*-farnesyltransferase has been cloned from *Rhododendron dauricum* (Ericaceae) [Saeki et al, 2018], and is predicted to be a typical aromatic PT with 9 transmembrane domains and two conserved aspartate-rich motifs. In this case, the authors suggest that loops 2 and 6, with the aspartate-rich motifs, are localized to the intermembrane space. The authors also suggest that the FDP (**27**) required for the farnesylation of orsellinic acid is synthesized in the plastid. However, it is generally accepted that in plants FDP (**27**) is synthesized in the cytosol. The acceptor orsellinic acid is also produced in the cytosol by a type III polyketide synthase from acetyl-CoA and three molecules of malonyl-CoA [Taura et al, 2016]. Both the donor FDP (**27**) and acceptor orsellinic acid may be passively transported into the intermembrane space of the plastid envelope for the prenylation reaction as outlined above.

So far, no information can be found on aromatic PTs involved in the biosynthesis of sesquiterpene coumarins. However, recently it was shown that umbelliprenin (**1**) biosynthesis was induced in callus cultures of *Ferulago campestris* (Apiaceae) after treatment with ferulic acid [Fiorito et al, 2022]. Elicitation of the culture with 10 μM ferulic acid resulted in more than a 200-fold increase in the level of umbelliprenin (**1**) from 12 μg/g to 2.6 mg/g dry weight. The level of umbelliferone (**9**) in the elicited cells was increased 40-fold from 6.4 to 270 μg/g dry weight. Obviously, the elicitation of the callus culture resulted in the induction of enzymes involved in the biosynthesis of umbelliprenin (**1**). Isolation of an *O*-farnesyltransferase cDNA from an elicited cell culture should be possible using a suitable prenyltransferase probe. This would be a step forward in our understanding of sesquiterpene coumarin biosynthesis.

A total of 277 different sesquiterpene coumarins are formed through *O*-prenylation of the 7-hydroxycoumarins umbelliferone (**9**), scopoletin (**10**), fraxidin (**11**) or isofraxidin (**12**) by *O*-farnesyltransferase as summarized in Table 1. As shown in Figure 5, ionization of FDP (**27**) takes place in the active site of the *O*-PT by abstraction of the diphosphate group by Mg^2+^-ions coordinated to the aspartate-rich motif. A C-O bond is formed by abstraction of the hydroxyl proton of the coumarin by a basic amino acid with simultaneous attack by the oxygen on the carbocation. In this way, the ether linkage of the sesquiterpene coumarin is established. The 7-farnesyloxycoumarins produced are linear sesquiterpenes, which are designated to the carbon skeleton group ***O*-A** (Figure 3). Only one 7-nerolidyloxycoumarin isolated from *Ferula cocanica* has been reported and is assigned carbon skeleton ***O*-B** (Figure 3) [Kiryalov, 1961]. The products umbelliprenin (**1**), scopofarnol (**31**) and farnochcrol (**32**) belong to the carbon skeleton groups ***O*-U-A, *O*-S-A** and ***O*-I-A**, respectively. Additional coumarins found as building blocks of sesquiterpene coumarins are fraxetin (**11**), 8-hydroxycoumarin (**13**), 4-hydroxycoumarin (**14**), 5,7-dihydroxycoumarin (**17)** and 4-hydroxy-5-methylcoumarin (**18**).

**Figure 5.**
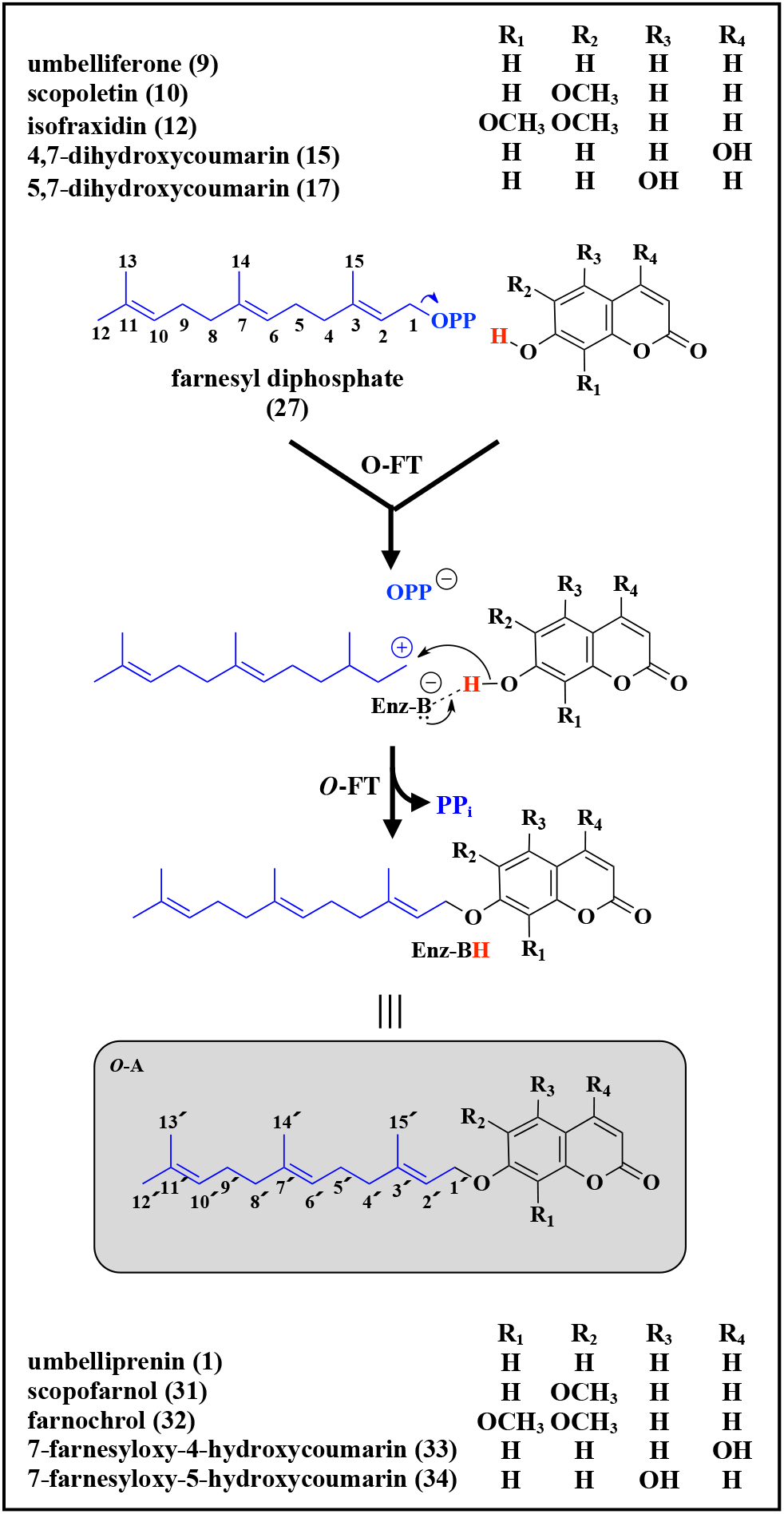
*O*-Farnesylation of 7-hydroxycoumarins by *O*-farnesyltransferase (*O*-FT). Sesquiterpene coumarin carbon skeletons ***O*-A** is formed. Nerolydation in the same way results in formation of carbon skeleton ***O*-B**

The linear sesquiterpene coumarins are modified by hydroxylation, esterfication, glycosylation and/or epoxidation reactions. The structures of linear sesquiterpene coumarins belonging to carbon skeleton groups ***O*-A** and ***O*-B** are shown in supplementary Figure S2. One chiral atom is found in structure ***O*-B**. It is important to determine the stereochemistry of sesquiterpene coumarins. The biological activity is dependent on the stereochemistry of the molecule. In the various carbon skeleton structures obtained in biosynthetic schemes, chiral carbons of the end products are marked with black dots. When more than one diastereomer of a specific carbon skeleton have been isolated, the suggested biosynthetic pathways do not show any stereochemistry. The stereochemistry of such sesquiterpene coumarins is shown in Figures S2 to S9.

### 4.2. Step 2: Epoxidation

Essentially all cyclic sesquiterpene coumarins derived from 7-farnesyloxycoumarins carry a hydroxyl group on C3′. This is a strong indication that linear sesquiterpene coumarins, such as umbelliprenin (**1**), scopofarnol (**31**) and farnochrol (**32**), are first epoxidized to prepare for the generation of a carbocation by protonation of the epoxide (Figure 4). This carbocation will initiate cyclization of the linear substrate to cyclic products using a carbocation mechanism. This assumption is supported by the fact that 10′,11′-oxidoumbelliprenin (**28**) (***O*-U-A2**) has been isolated from *Ferula turcica* [Erucar et al, 2023], *Heptaptera cilicica* [Güvenalp et al, 2017] and 10′,11′-oxidofarnochrol (***O*-I-A2)** from *Achillea ochroleuca* [Greger et al, 1983b; Jandl *et al*., 1997], *Anthemis cretica* [Hofer & Greger, 1985] *Artemisia tripartita* [Greger et al, 1983a], and *Ferula jaeschkeana* [Razdan et al, 1989].

An additional indirect proof for 10′,11′-oxidoumbelliprenin (**28**) as an intermediate of the pathway is the isolation of linear sesquiterpene coumarins with 10′- and 11′-hydroxyl groups, such as 10′R-karatavicinol (**35**) (***O*-U-A10**), which has been isolated from several *Ferula* species [Abd El-Razek et al, 2007; Ahmed, 1999; Amin et al, 2016; Hofer et al, 1984; Lee et al, 2009; Shomirzoeva et al, 2021; Teng et al, 2013; Wang et al, 2020; Xing et al, 2017] and *Heptaptera* species [Cicek Kaya et al, 2022; Miski et al, 2015; Tosun et al, 2019, 2021]. Karatavicinol (**35**) is obtained from 10′,11′-oxidoumbelliprenin (**28**) by an epoxyhydrolase as shown in Figure 6.

**Figure 6.**
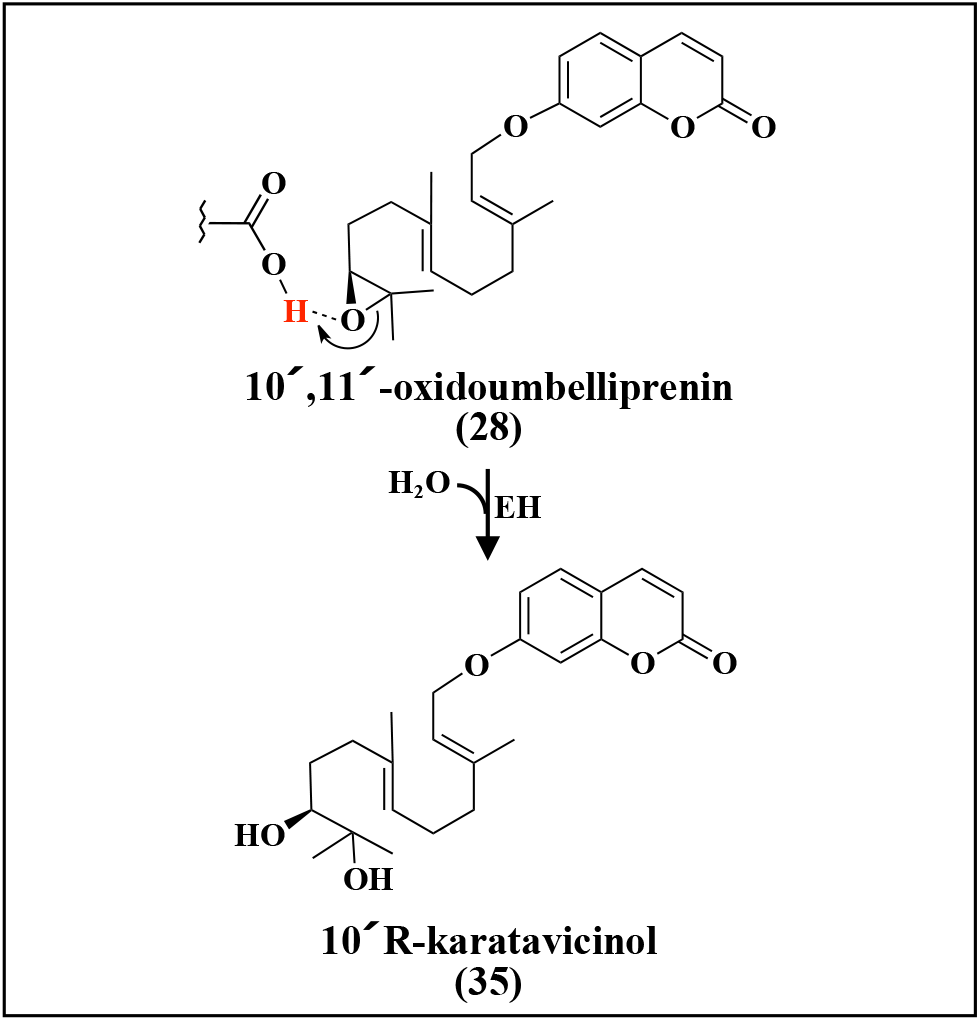
Formation of 10′-R-karatavicinol (**35**) from 10′,11′-oxidoumbelliprenin (**28**) by epoxyhydrolase (EH).

Epoxidation of 7-farnesyloxy-5-hydroxycoumarin (**34**) with subsequent hydrolysis by epoxyhydrolase will yield 10′,11′,5-trihydroxyumbelliprenin (***O*-DH-A1** (Figure S2), which has been isolated from *Heptaptera anatolica* and *Heptaptera anisoptera* [Appendino et al, 1992]. The sesquiterpenes ***O*-DH-A2** and ***O*-DH-A3** (Figure S2) are obtained from 10′,11′,5-trihydroxyumbelliprenin. Other examples are the linear sesquiterpene coumarins ***O*-U-A11** to ***O*-U-A17** and ***O*-U-A19** to ***O*-U-A21** (Figure S2), all of which are most likely produced from karatavicinol (**35**). All these findings support the assumption that 10′,11′-epoxidation of 7-farnesyloxycoumarins with subsequent protonation is the mechanism used for the generation of initial carbocations for biosynthesis of cyclic sesquiterpene coumarins (as discussed below).

There are at least three different types of monooxygenases that can catalyze the epoxidation of metabolites in plants. These belong to the families of cytochrome P450 monooxygenases (P450s), FAD monooxygenases (FMOs) and nonheme iron(II)- and 2- oxoglutarate-dependent oxygenases (2OGXs). No information is available on which type of monooxygenase is involved in the biosynthesis of sesquiterpene coumarins. The first two types of monooxygenases catalyze the same overall reaction:

7-farnesyloxycoumarin + O_2_ + NADPH + H^+^ →7-(10′,11′-oxidofarnesyloxy)coumarin + H_2_O + NAD

In the third case, the epoxidation is catalyzed by 2OGX, which utilizes a Fe(IV)-oxo intermediate to initiate diverse oxidative transformations including epoxidation. No information is available on sesquiterpene coumarin epoxidase (STCE).

#### 4.2.a. Cytochrome P450 monooxygenase

Cytochrome P450 monooxygenases (P450s) are a diverse superfamily of heme-dependent enzymes that catalyze the introduction of one atom of molecular oxygen into nonactivated C-H bonds, often in a regio- and stereoselective manner. Epoxidation of plant metabolites is a common reaction that is catalyzed by P450s [Mitchell & Weng, 2019]. P450s require electrons for activation of O_2_ by the heme b prosthetic group for the epoxidation reaction [Jin et al, 2003; Coleman et al, 2021]. These electrons are obtained from NADPH by the help of NADPH-cytochrome P450 reductase (CPR). In the first step, the substrate binds to the Fe^III^-enzyme displacing a water molecule in the active site. This is followed by the transfer of one electron to the heme-iron thereby reducing it from Fe^III^ to Fe^II^. Subsequently, the binding of molecular oxygen results in an oxy-P450 complex. This complex is reduced by a second electron transfer, and after a double protonation at the distal oxygen and water release, the O-O bond is cleaved, resulting in the reactive enzyme intermediate (compound I (**129**)), which is an Fe^IV^-oxo porphyrin radical cation. This intermediate inserts the oxygen atom into a double bond. The epoxide is formed before release of the product and the enzyme returns to the ferric resting state with a water molecule bound in the active site.

#### 4.2.b. FAD monooxygenase

The flavin monooxygenases (FMOs) utilize FAD to catalyze various oxidative reactions, including epoxidations, by forming a reactive intermediate between singlet oxygen and the C4a of the flavin moiety [Deng et al., 2022]. To activate the enzyme, FAD is reduced to FADH_2_ by obtaining electrons from NADPH via CPR with subsequent reaction of the flavin with singlet oxygen. During binding of oxygen, a proton is abstracted from an amino acid residue of the active site leading to the formation of a reactive flavin hydroperoxide intermediate. After monooxygenation of the substrate, and formation of the epoxy group by abstraction of a proton, the formed hydroxyflavin decays and yields the oxidized flavin (FAD) with H_2_O as side-product.

Squalene epoxidase (squalene monooxygenase) (SQE) is a rate-limiting FMO of triterpene biosynthesis [Chua et al, 2020]. In the formation of triterpenes from 2,3-oxidosqualene (**22**), protonation of the epoxy group generates the carbocation needed for the cyclization [Thimmappa et al, 2014; da Silva Magedans et al, 2021]. All plant triterpenes carry a hydroxyl or keto group on carbon 3, which is the product of the protonation of the 2,3-epoxide of oxidosqualene (**22**). SQEs from some plants have been cloned and characterized [He et al, 2008; Gao et al, 2016].

Analysis of SQE amino acid sequences predicted a transmembrane domain at the N-terminus, suggesting that the SQE protein is anchored to the endoplasmic reticulum membrane as exemplified in *Dioscorea zingiberensis* [Song et al, 2019], *Bupleurum chinense* [Gao et al, 2016], *Panax vietnamensis* [Ma et al, 2016] and *Betula platyphylla* [Zhang et al, 2016]. SQE is co-localized with CPR on the cytosolic side of the endoplasmatic reticulum membrane as seen with other monooxygenases [Barnaba et al, 2017].

#### 4.2.c. 2-Oxoglutarate-dependent oxygenase

The family of 2-oxoglutarate-dependent oxygenases (2OGXs) includes enzymes catalyzing a diverse range of biologically important reactions in plants [Islam et al, 2018]. These enzymes function in biosynthesis and catabolism of cellular metabolites, including secondary metabolites as exemplified by the biosynthesis of scopolamine by hyoscyamine 6β-hydroxylase [Hashimoto et al, 1991]. 2OGXs are involved in hydroxylations, desaturations, cyclizations, halogenation and other reactions [Herr & Hausinger, 2018]. Indeed, recently 2OGXs have been shown to catalyze furan ring formation in plant limonoid biosynthesis [De La Peñe et al, 2023]. Here, the epoxidation activity of 2OGXs is of relevance. The reaction mechanism of epoxidation via oxygen atom transfer has been studied [Li et al, 2020c].

The mechanism of epoxidation by 2OGXs is initiated with Fe(II) coordinated to a His-Asp/Glu-His triad, with three additional coordination sites occupied by water molecules [Martinez et al, 2015]. Two of the metal-bound water molecules are displaced by binding of 2-oxoglutarate to the Fe(II) center. Upon binding of the substrate to the enzyme active site (not to the metal ion) the third metal-bound water is displaced. This substrate-triggered process creates a site for binding of an O_2_ molecule, generating a Fe(III)-superoxo intermediate. The distal oxygen atom of the Fe(III)-superoxo species attacks C2 of 2-oxoglutarate to yield a peroxohemiketal bicyclic intermediate. This species initiates the oxidative decarboxylation of 2-oxoglutarate releasing CO_2_ and yielding a Fe(IV)-oxospecies (called the ferryl intermediate), which is bound to succinate. The ferryl intermediate attacks the double bond of the substrate to generate a Fe(III)-O-substrate radical. The subsequent C−O bond formation completes the epoxide ring formation. The product and succinate are released, and three molecules of water are added to complete the reaction cycle and regenerate the starting Fe(II) complex.

### 4.3. Ionization of 10′,11′-oxidoumbelliprenin

The first step in the cyclization of 10′,11′-oxidoumbelliprenin (**28**) is protonation of the epoxide by the STCS to generate the initial carbocation, as shown in Figure 4B. This step is the same for various STCSs using 7-(10′,11′-oxidofarnesyloxy)-coumarins as a substrate. The same type of carbocation generation takes place in the biosynthesis of triterpenes from 2,3-oxidosqualene (**22**) (Figure 4A). In triterpene synthases (oxidosqualene cyclases), the proton donor is an aspartic acid residue, which is found within a highly conserved amino acid sequence. This conserved sequence is Asp-Cys-Thr-Ala-Glu, and it has been defined as the protonation and initiation site of triterpene synthases. Examples are cycloartenol synthase [Corey et al, 1993], lanosterol synthase [Forestier et al, 2019], α-amyrin synthase [Yu et al, 2018], lupeol synthase [Guhling et al, 2006], cucurbitadienol synthase [Qiao et al, 2018], marneral synthase [Xiong et al, 2006], butyrospermol synthase [Forestier et al, 2019], and dammarenediol-II synthase [Tansakul et al, 2006]. Future cloning of STCS genes will show if the same conserved sequence is used for protonation and initiation of the carbocation cascade reactions.

### 4.4. Cyclization of initial carbocation

In Figure 4B, putative pathway for the biosynthesis of farniseferol B (**5**) conferol (**3**) from umbelliferone (**9**) and FDP (**27**) are shown. After prenylation and epoxidation, 10′,11′-oxidoumbelliprenin (**28**) is protonated by STCS and 10′,11′-oxidoumbelliprenin (**28**) is protonated by STCS and a C11′ carbocation is obtained. This carbocation is converted to a monocyclic sesquiterpene carbocation (**29**) by C6′, C11′-ring closure. By proton abstraction the monocyclic product farnesiferol B (**5**) is obtained. The carbocation can go through a second ring closure (C6′, C10′-closure) to generate the bicyclic drimane-type sesquiterpene carbocation (**30**). By proton abstraction the bicyclic product conferol (**3**) is generated. The two carbocations (**29**) and (**30**) are intermediates in many of the putative pathways leading to various carbon skeletons (Figure 3). In the following, putative pathways for the biosynthesis of various carbon skeleton will start from the intermediate carbocations (**29**) or (**30**).

It is well known that terpene synthases exhibit different degrees of product specificity. High-fidelity terpene syntases produce close to 100% of a specific product, while low-fidelity terpene synthases produce a number of products [Christianson, 2017]. Cotton (+)-δ-cadinene synthase producing more than 98% (+)-δ-cadinene is an example of a high-fidelity sesquiterpene synthase [Yoshikoni et al, 2006], while γ-humulene synthase from *Abies grandis*, a highly promiscuous sesquiterpene synthase, produce 28.6% of the main product γ-humulene and 51 other sesquiterpenes in amounts between 0.1 to 15.1% [Steele et al, 1998]. The correct sequence of carbon−carbon bond forming reactions in the cyclization cascade is directed by the precatalytic binding conformation of the substrate and the accessible conformation(s) of subsequently formed carbocation intermediates, so the fidelity of the cyclization pathway is encoded in the three-dimensional contour of the cyclase active site is directing the correct sequence of carbon−carbon bond forming reactions in the cyclization cascade, which encodes the fidelity of the terpene synthase.

Nothing is known about the fidelity of STCSs. This cannot be investigated until the cloning of STCSs has been achieved and recombinant enzyme has been produced and characterized. In this situation, we assume that all carbon skeletons of sesquiterpene coumarins (Figure 3) are produced by specific STCSs, which is indicated in all putative biosynthetic pathways by addition of carbon skeleton type to the enzyme abbreviation. For example, the enzyme STCS-Ca (Figure 7A) is catalyzing the formation of carbon skeleton Ca. However, it is highly likely that STCSs are at least to a certain degree promiscuous and produce more than one product. If we look at Figure 7 where the putative biosynthesis of 8 different carbon skeletons (O-C to O-J) from the two intermediate carbocation (**29**) and (**30**) is shown. Six of these skeletons are obtained by quenching the carbocation by abstraction of different protons resulting in a double bond at different locations and two are obtained by quenching the carbocation by addition of water. It is possible that some or even all these products are produced by one STCS exhibiting a broad product specificity. *Ferula sinkiangensis* is probably the richest source of sesesquiterpene coumarins. Khayat et al [2023] list 60 sesquiterpene coumarins isolated from *F. sinkiangensis*. Five of these are farnesiferol B (**5**) (***O*-U-D1**), sinkianol B (**38**) (***O*-U-Ea6**), feselol (**37**) (***O*-U-Ha7**), colladonin (**36**) (***O*-U-Ia6**) and isosamarcandin (**39**) (***O*-U-Ja13**), which have the same stereochemistry of the sesquiterpene moiety except for location of the double bond or the hydroxyl group. Thus, it is possible that only one STCS is involved in the formation of these carbon skeleton from 10′,11′-oxidoumbelliprenin (**28**) as outlined in Figure 8.

**Figure 7.**
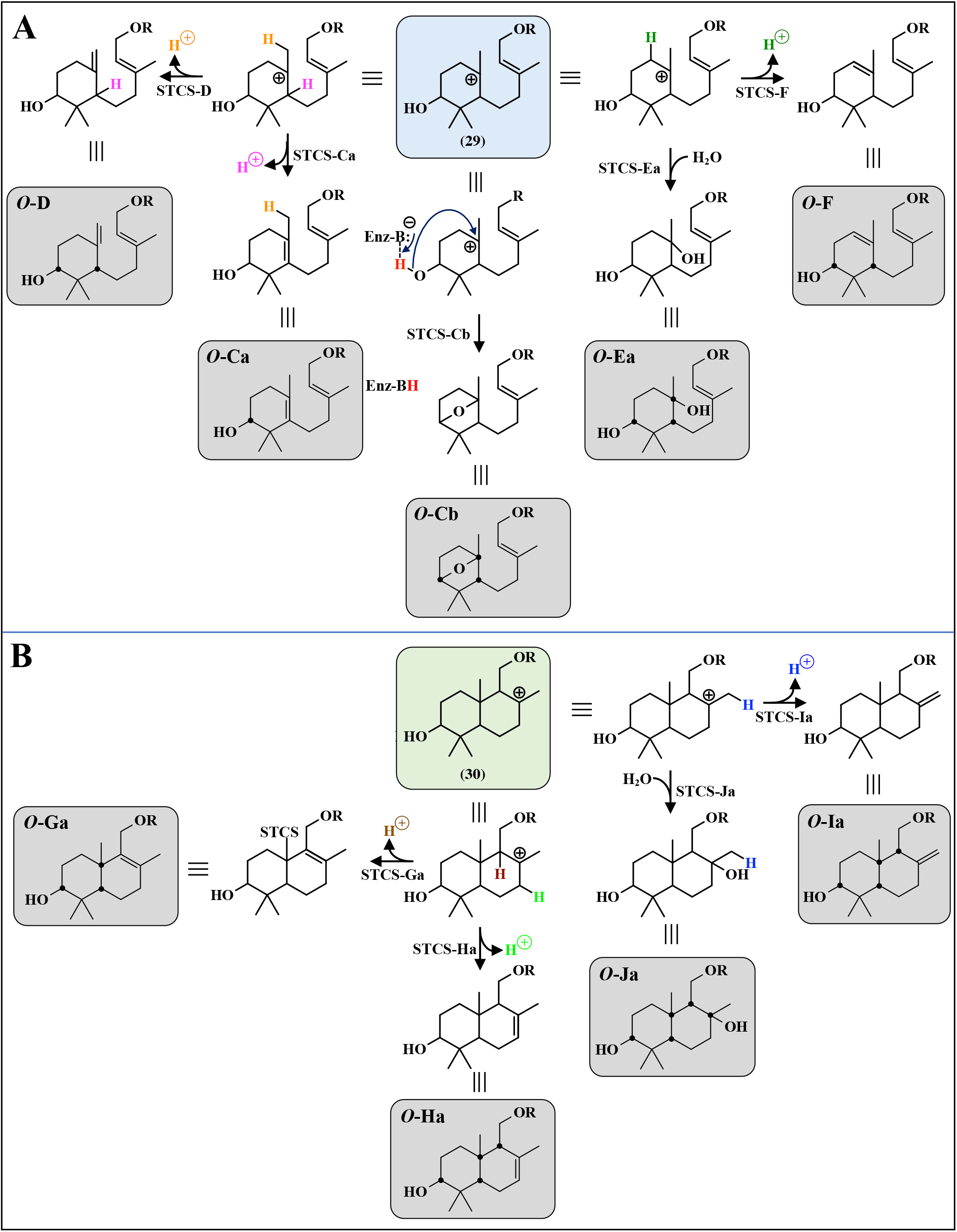
**A:** Putative pathways from the intermediate carbocation I (**29**) generated by sesquiterpene synthases (STCSs) to the sesquiterpene coumarins with carbon skeletons ***O*-C** to ***O*-E** after cyclizations and quenching of the final carbocation. **B:** Putative pathways from the intermediate carbocation II (**30**) generated by sesquiterpene synthases (STCSs) to the sesquiterpene coumarins with carbon skeletons ***O*-C** to ***O*-E** after cyclizations and quenching of the final carbocation The carbon skeletons ***O*-C** to ***O*-J** are the products of different STCSs, which generates the common carbocations I and II (see Figure 4). Chiral carbon atoms are indicated by black dots in the final structures.

**Figure 8.**
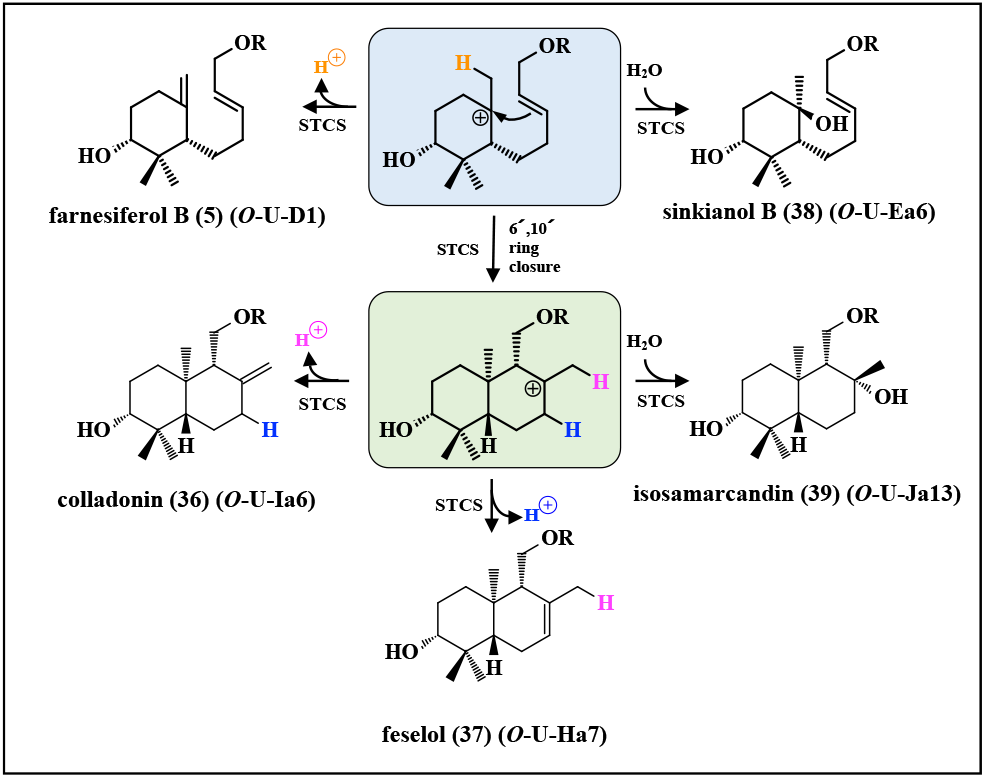
Example of a putative promiscuous STCS producing five different sesquiterpene carbon skeletons found in *Ferula sinkiangensis*.

The monocyclic carbocation (**29**) is quenched by proton eliminations to generate the monocyclic carbon skeletons ***O*-Ca, *O*-D**, and ***O*-F** or by the addition of water to generate the monocyclic carbon skeleton ***O*-E** as shown in Figure 7A. In addition, a C4′, C8′-ether bridge can be formed to obtain skeleton ***O*-Cb** by quenching of the monocyclic C6′-carbocation (**29**) with the hydroxyl group at C4′ (Figure 7A). This is analogous to the formation of the monoterpene 1,8-cineole from α-terpineol [Piechulla et al, 2016] and to the formation of the triterpene baccharis oxide by an oxidosqualene cyclase (OSC), the cDNA of which was cloned from the roots of *Stevia rebaudiana* and expressed in yeast [Shibuya et al., 2008].

Quenching of the bicyclic C′8-carbocation (**30**) through different proton eliminations yields the end products with carbon skeletons ***O*-G, *O*-H, *O*-I** as shown in Figure 7B. The alternative addition of water yields carbon skeleton ***O*-J** (Figure 7B).

The diversity of sesquiterpene coumarins is increased considerably by modification of the carbon skeletons by hydroxylations, epoxidations, esterfications, desaturations, glycosylations, *etc*. A particularly important group of enzymes responsible for many of these modifications in sesquiterpene coumarins is the P450s [Villa-Ruano et al, 2015; Banerjee & Hamberger, 2018; Li et al, 2020a]. An extensive review on hydroxylases involved in terpenoid biosynthesis is of particular relevance [Zhang et al, 2023b].

### 4.5. The evolution of sesquiterpene epoxidase and sesquiterpene coumarin synthase

As pointed out above the proposed putative biosynthetic pathway of sesquiterpene coumarins exhibits striking similarities to the biosynthesis of triterpenes (Figure 4). The substrates umbelliprenin (**1**) (C_24_O_3_H_30_; MW 366) and squalene (**20**) (C_30_H_50_; MW 410) are relatively large linear molecules. First, an epoxy group is introduced at the terminal double bond of a farnesyl moiety by an epoxidase

In the next step both epoxides undergo cyclization reactions after protonation of the epoxy group by cyclases (synthases).

Although no plant enzymes have been studied, the structures of both the catalytic domain of human SQE (aa 118 to 574) and the human OSC lanosterol synthase have been solved [Padyana et al, 2019, Thoma et al. 2004]. *In silico* docking of umbelliprenin (**1**) and squalene (**20**) to the active site of human SQE, and the docking of 10′,11′-oxidoumbelliprenin (**28**) and 2,3-oxidosqualene (**22**) to the active site of lanosterol synthase are shown in Figure 9A and B. In both cases it was found that the unnatural substrate (umbelliprenin (**1**) or 10′,11′-oxidoumbelliprenin (**28**)) could adopt a binding mode that was in close agreement to the lowest energy binding model of the respective enzyme’s natural substrate (squalene (**20**) or 2,3-oxidosqualene (**22**)). These binding models show that umbelliprenin (**1**) and 10′,11′-oxidoumbelliprenin (**28**) are well-accommodated and display the correct orientation with respective to the catalytic domain of each enzyme (the FAD cofactor in SQE, and the ASP455 residue in lanosterol synthase) for the anticipated equivalent epoxidation or protonation reactions observed with the natural substates to appear feasible. This is best illustrated by the overlays of the docked confirmations of umbelliprenin (**1**) with squalene (**20**) and 10′,11′-oxidoumbelliprenin (**28**) with 2,3-oxidosqualene (**22**) shown in the bottom panels of Figure 9A and B.

**Figure 9.**
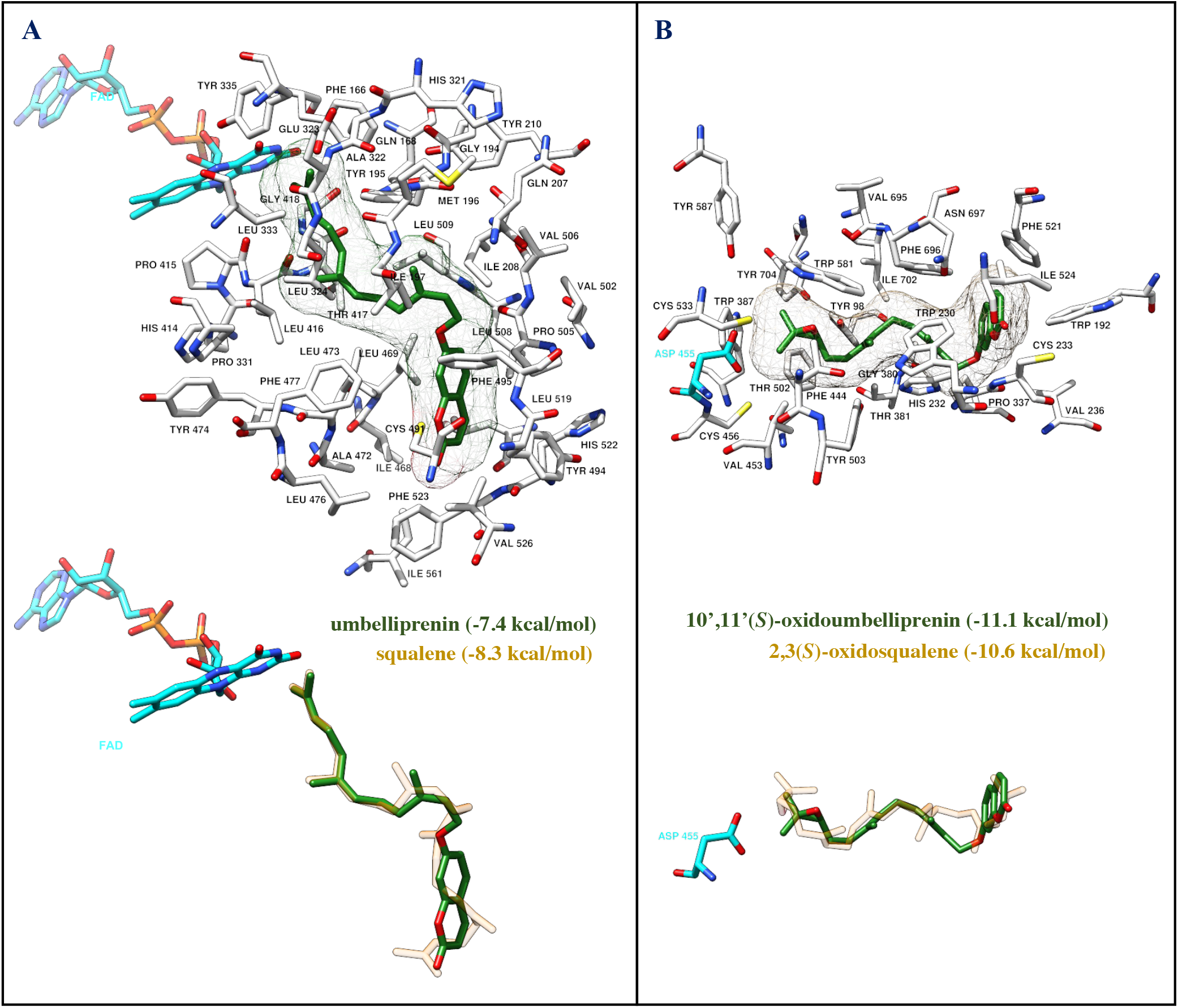
**(A)** Top: umbelliprenin (**1**) (green) docked to *Human Squalene Epoxidase* (PDB: 6C6R) with residues within 5 ångströms shown (surface of umbelliprenin (**1**) shown). FAD highlighted in cyan. [Binding affinity, umbelliprenin (**1**): -7.4 kcal/mol] Bottom: Docked umbelliprenin (**1**) (green) overlayed with docked squalene (**20**) (transparent orange) [Binding affinity, squalene (**20**): -8.4 kcal/mol]. **(B)** Top: 10’,11’(*S*)-oxidoumbelliprenin (**28**) (green) docked to *Human lanosterol synthase* (PDB: 1W6K) with residues within 5 ångströms shown (surface of 10’,11’(S)-oxidoumbelliprenin (**28**) shown). The catalytic ASP 455 residue is highlighted in cyan. [Binding affinity, 10’,11’(S)-oxidoumbelliprenin (**28**): -11.1 kcal/mol]. Bottom: Docked 10’,11’(S)-oxidoumbelliprenin (**28**) (green) overlayed with docked 2,3(S)-oxidosqualene (**22**) (transparent orange) [Binding affinity, 2,3(S)-oxidosqualene (**22**): -10.6 kcal/mol]. Docking experimental details. *Autodock vina*, rigid receptor (default receptor prep with *Mgltools*), flexible ligands (default ligand prep with *Mgltools*), configuration: exhaustiveness = 24, energy range = 10, number of modes = 10, search space; [6C6R: seed = 166819220, center x = -8.689, center y = -54.255, center z = -1.066, size x = 16.8979, size y = 27.45, size z = 17.3766], [1W6K, : seed = 166819220, center x = 31.09, center y = 70.677, center z = 6.968, size x = 30, size y = 30, size z = 30]. Ligand starting geometries were optimised by Molecular Mechanics before preparation [*Avogadro*, force field = MMFF94s, Number of steps = 500, Algorithm = Steepest Descent, Convergence = 10e-7]. Program names in italics.

Squalene hopene cyclase (SHC), often used as a model system for OSCs, protonates the terminal double bond of squalene (**20**) to initiate cyclization and formation of hopene (**21**) (Figure 4A). Studies on substrate specificity of SHC have shown that different substrates with a terminal double bond are converted to cyclic products (Table 2). Depending on the substrate, one or more products are formed. It is interesting to note that 2-(10′,11′-oxidofarnesyl)-phenol (**53**), 3-(farnesyldimethylallyl)-pyrrole (**54**) and 3-(farnesyl-dimethylallyl)indole (**55**), which are similar in structure and size to 10′,11-oxidoumbelliprenin (**28**), are substrates.

**Table 2.**
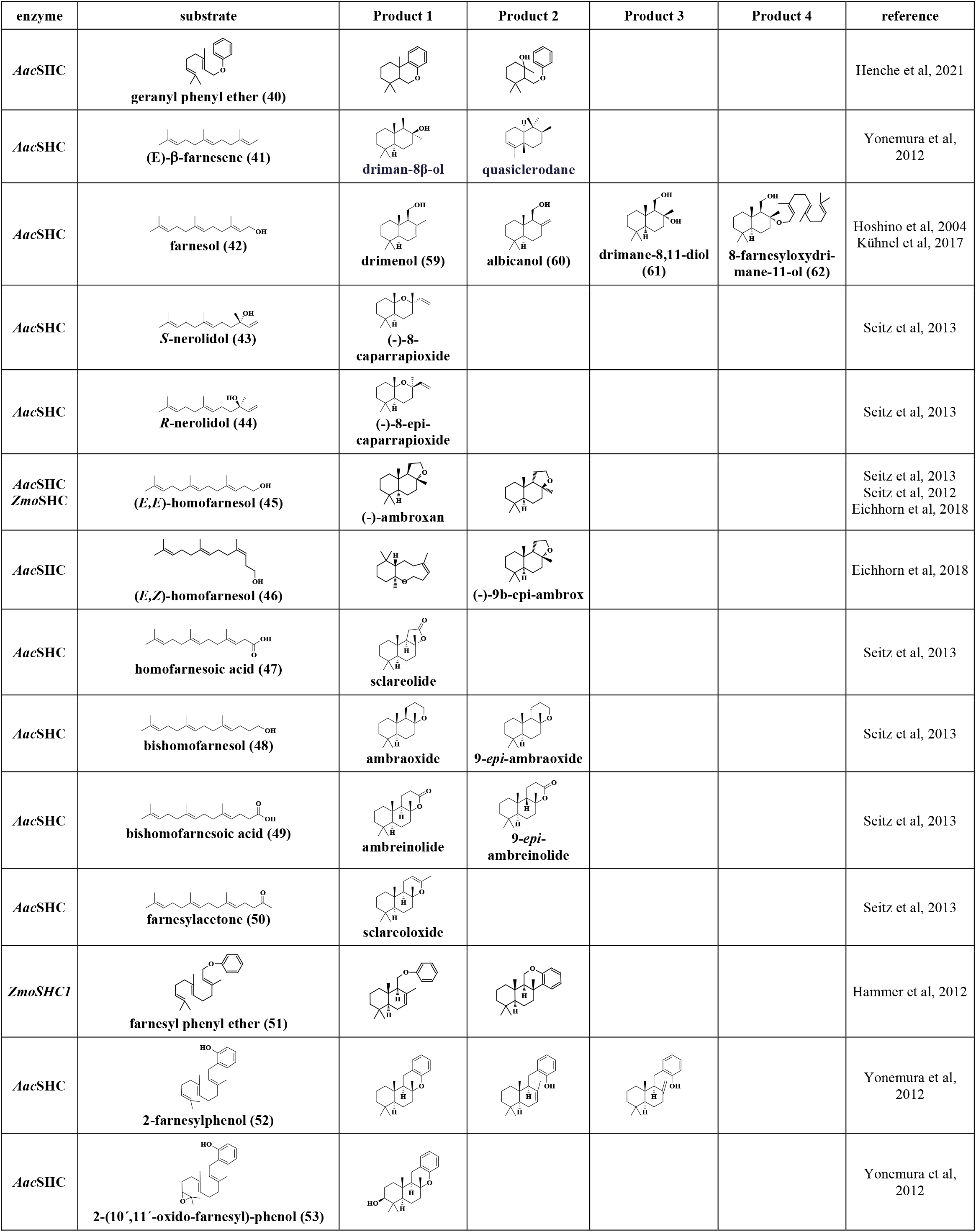
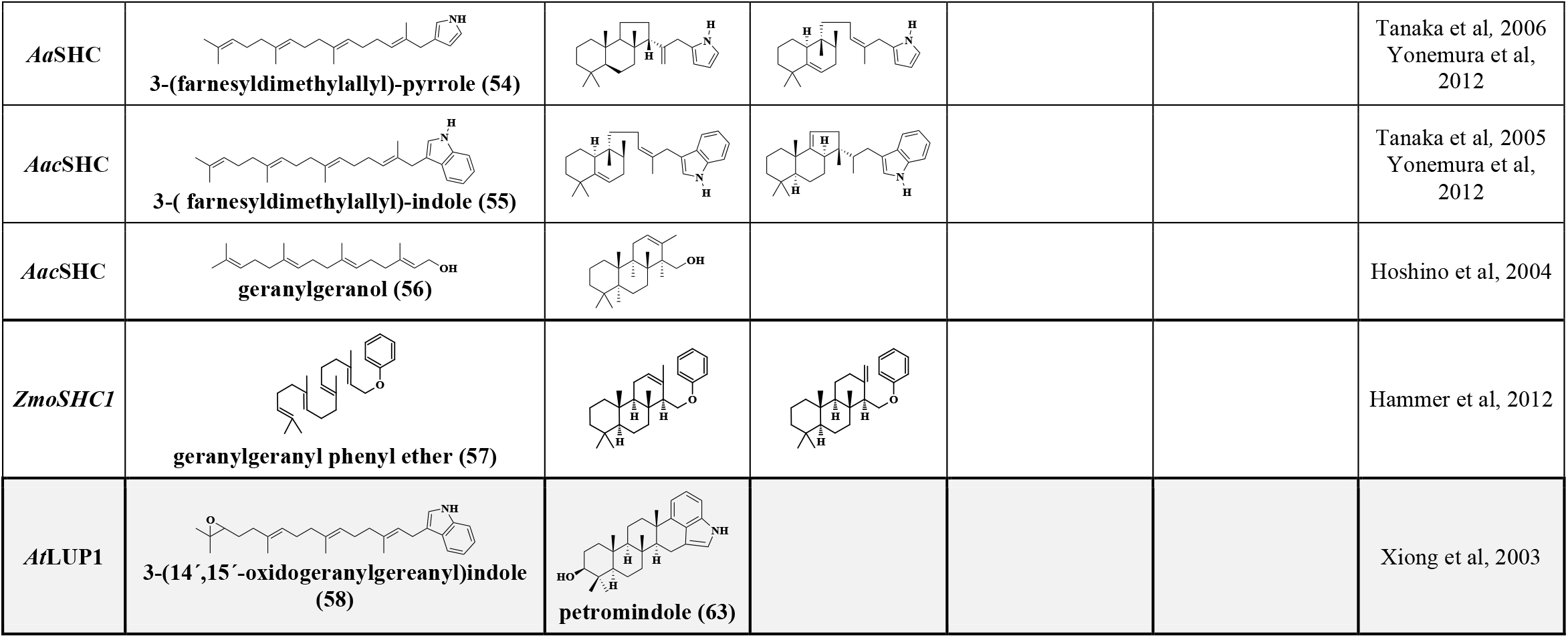
Activity of squalene hopene cyclase from *Alicyclobacillus acidocaldarius* (*Aac*SHC), *Zymomonas mobilis* (*Zmo*SHC) and lupeol synthase from *Arabidopsis thaliana* (*At*LUP1) with different substrates.

Squalene hopene cyclase (SHC) converts farnesol (**42**) to three bicyclic sesquiterpenes as shown in Table 2. The product mixture contained 4% drimenol (**59**), 7% albicanol (**60**) and 85% drimane-8,11-diol (**61**) [Hoshino et al, 2004; Kühnel et al, 2017]. The carbon skeletons of these three sesquiterpenes are the same as those of sesquiterpene coumarins ***O*-Ha, *O*-Ia** and ***O*-Ja**, respectively (Figure 7B).

This means that around 46% of all sesquiterpene coumarins discussed in this review have the same carbon skeletons as those sesquiterpenes produced by SHC from farnesol (**42**). Obviously, the active site configurations of SHC and STCS are guiding the cyclization of the farnesyl moiety in a common way. Furthermore, it is interesting to note that the remaining 4% of the product mixture was shown to be 8-farnesyloxydrimane-11-ol (**62**) (Table 2) [Kühnel et al, 2017]. In this case the intermediate bicyclic carbocation is quenched by addition of a second farnesol molecule. This means that SHC functions as a farnesyltransferase but with a different reaction mechanism compared to normal *O*-farnesyltransferases using FDP (**27**) (*cf* Figure 5).

The OSC lupeol synthase (LUP1) converts 3-(14′,15′-oxidogeranylgeranyl)-indole (**58**) to the indole diterpene petromindole (**63**) (Table 2 shaded row) [Xiong et al, 2003].

It is interesting to note that a triterpene synthase (lupeol synthase) and an indole ditepene synthase (petromindole synthase) produce identical products from 3-(14′,15′-oxidogeranylgeranyl)-indole (**58**). Pentacyclic triterpenes, such as lupeol (**26**) and hopene (**21**), share the same relative stereochemistry in rings A-D and the same substituents in rings A and B as petromindole (**63**). Based on these similarities it was suggested that petromindole synthase may be closely related to plant triterpene synthases [Xiong et al, 2003]. It would be worthwhile to test if 10′,11-oxidoumbelliprenin (**28**) is protonated and cyclized by SHC and/or an OSC such as lanosterol synthase.

The OSC lupeol synthase (LUP1) converts 3-(14′,15′-oxidogeranylgeranyl)-indole (**58**) to the indole diterpene petromindole (**63**) (Table 2 shaded row) [Xiong et al, 2003]. It is interesting to note that a triterpene synthase (lupeol synthase) and an indole ditepene synthase (petromindole synthase) produce identical products from 3-(14′,15′-oxidogeranylgeranyl)-indole (**58**). Pentacyclic triterpenes, such as lupeol (**26**) and hopene (**21**), share the same relative stereochemistry in rings A-D and the same substituents in rings A and B as petromindole (**63**). Based on these similarities it was suggested that petromindole synthase may be closely related to plant triterpene synthases [Xiong et al, 2003]. It would be worthwhile to test if 10′,11-oxidoumbelliprenin (**28**) is protonated and cyclized by SHC and/or an OSC such as lanosterol synthase.

In addition, some inhibition experiments on SHCs and OSCs are summarized in Table 3. 6′.7′:10′,11′-Bisoxidoumbelliprenin (**64**) (IC_50_ = 1.5 μM) is a somewhat stronger inhibitor of squalene hopene cyclase than 10′,11′-oxidoumbelliprenin (**28**) (IC_50_ = 2.5 μM). C6 substituents on the coumarin moiety of 10′,11′-oxidoumbelliprenin (**28**) lowers the potency of the inhibitors (**68, 69**). It is interesting to note that the sesquiterpene coumarin farnesiferol C (**67**) is a relative potent inhibitor of squalene hopene cyclase (IC_50_ = 7 μM). It is possible that 10′,11′-oxidoumbelliprenin (**28**) is a substrate for SHC. No attempts to show this in the inhibition experiments was reported by Cravotto et al (2004). Lanosterol synthase and cycloartenol synthase from various plants are strongly inhibited by inhibitors 1 (**68**) and 2 (**69**) (IC_50_ = 0.01 to 2.7 μM). These inhibitors carry a umbelliferone (**9**) moiety and a side chain with the same length as 10′,11′-oxidoumbelliprenin (**28**). It may be concluded that 10′,11′-oxidoumbelliprenin (**28**) can bind into the active site of SHCs and OSCs.

**Table 3.**
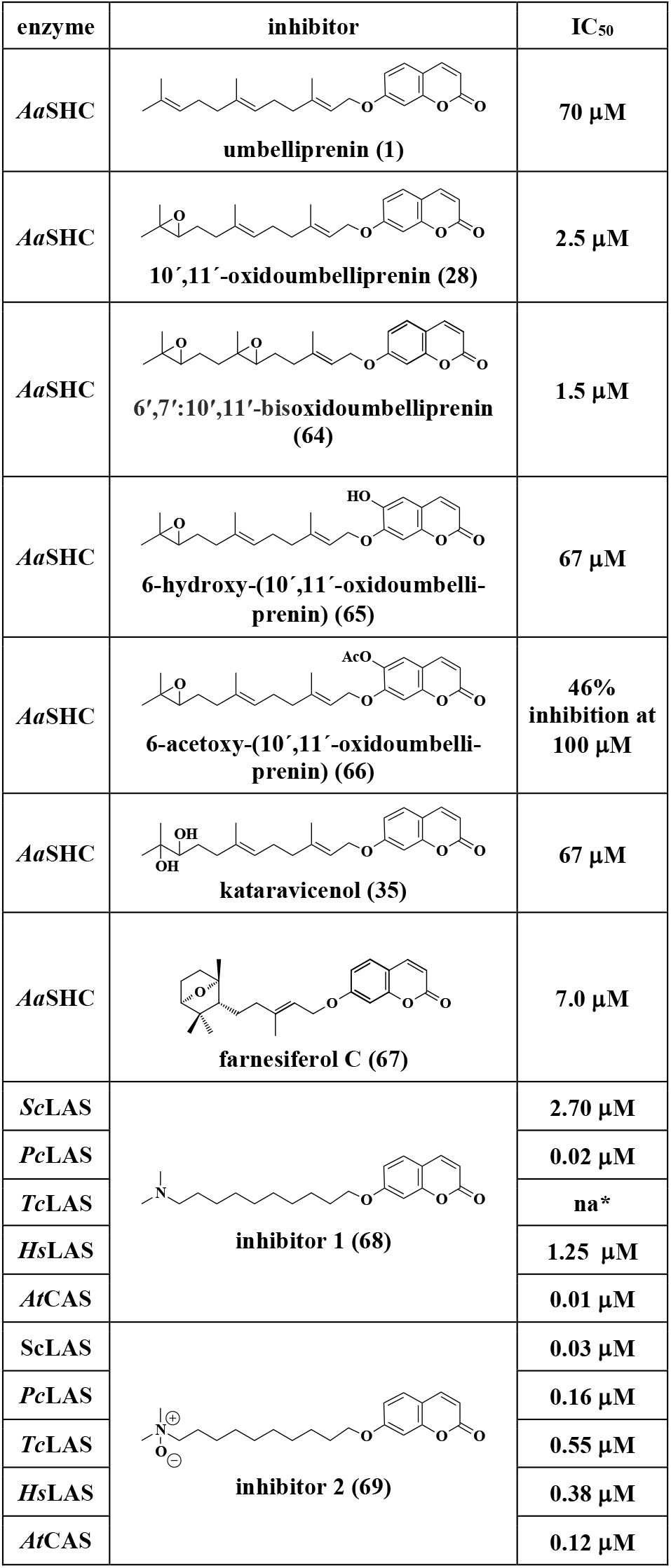
Inhibition of squalene hopene cyclase from *Alicyclobacillus acidocaldarius* (*Aa*SHC) (Cravotto et al, 2004) and lanosterol cyclase from *Saccharomyces cerevisiae* (*Sc*LAS), *Pheumocycsis carinii* (*Pc*LAS), *Trypanosoma cruzi* (*Tc*LAS), *Homo sapiens* (*Hs*LAS) and cycloartenol cyclase from *Arabidpsis thaliana* (*At*CAS) by umbelliferone derivatives [Oliaro-Bosso et al, 2007].

| enzyme | inhibitor | IC<sub>50</sub> |
| --- | --- | --- |
| *AaSHC* | umbelliprenin (1) | 70 µM |
| *AaSHC* | 10',11'-oxidoumbelliprenin (28) | 2.5 µM |
| *AaSHC* | 6',7':10',11'-bisoxidoumbelliprenin (64) | 1.5 µM |
| *AaSHC* | 6-hydroxy-(10',11'-oxidoumbelliprenin) (65) | 67 µM |
| *AaSHC* | 6-acetoxy-(10',11'-oxidoumbelliprenin) (66) | 46% inhibition at 100 µM |
| *AaSHC* | kataravicenol (35) | 67 µM |
| *AaSHC* | farnesiferol C (67) | 7.0 µM |
| *ScLAS* | inhibitor 1 (68) | 2.70 µM |
| *PcLAS* |  | 0.02 µM |
| *TcLAS* |  | na* |
| *HsLAS* |  | 1.25 µM |
| *AtCAS* |  | 0.01 µM |
| *ScLAS* | inhibitor 2 (69) | 0.03 µM |
| *PcLAS* |  | 0.16 µM |
| *TcLAS* |  | 0.55 µM |
| *HsLAS* |  | 0.38 µM |
| *AtCAS* |  | 0.12 µM |

Nearly all *O*-prenylated sesquiterpene coumarins have been isolated from plants belonging to the closely related Apiaceae and Asteraceae families, as summarized in Table S1. A phylogenetic tree of Angiosperms is shown in 10A (adapted from Cole *et al*., 2019) and a detailed phylogenetic tree of the clade Campanulids is shown in Figure 10B (adapted from Magallón *et al*., 2015). The numbers show the estimated divergence time in million years ago (mya). The families Apiaceae and Asteraceae belong to the orders Apiales and Asterales, respectively. The branching of the two linages Apiales and Asterales occurred 93.7 million years ago (mya) while the Apiaceae and Asteraceae diverted 58.0 and 49.3 mya, respectively [Magallón et al, 2015].

**Figure 10.**
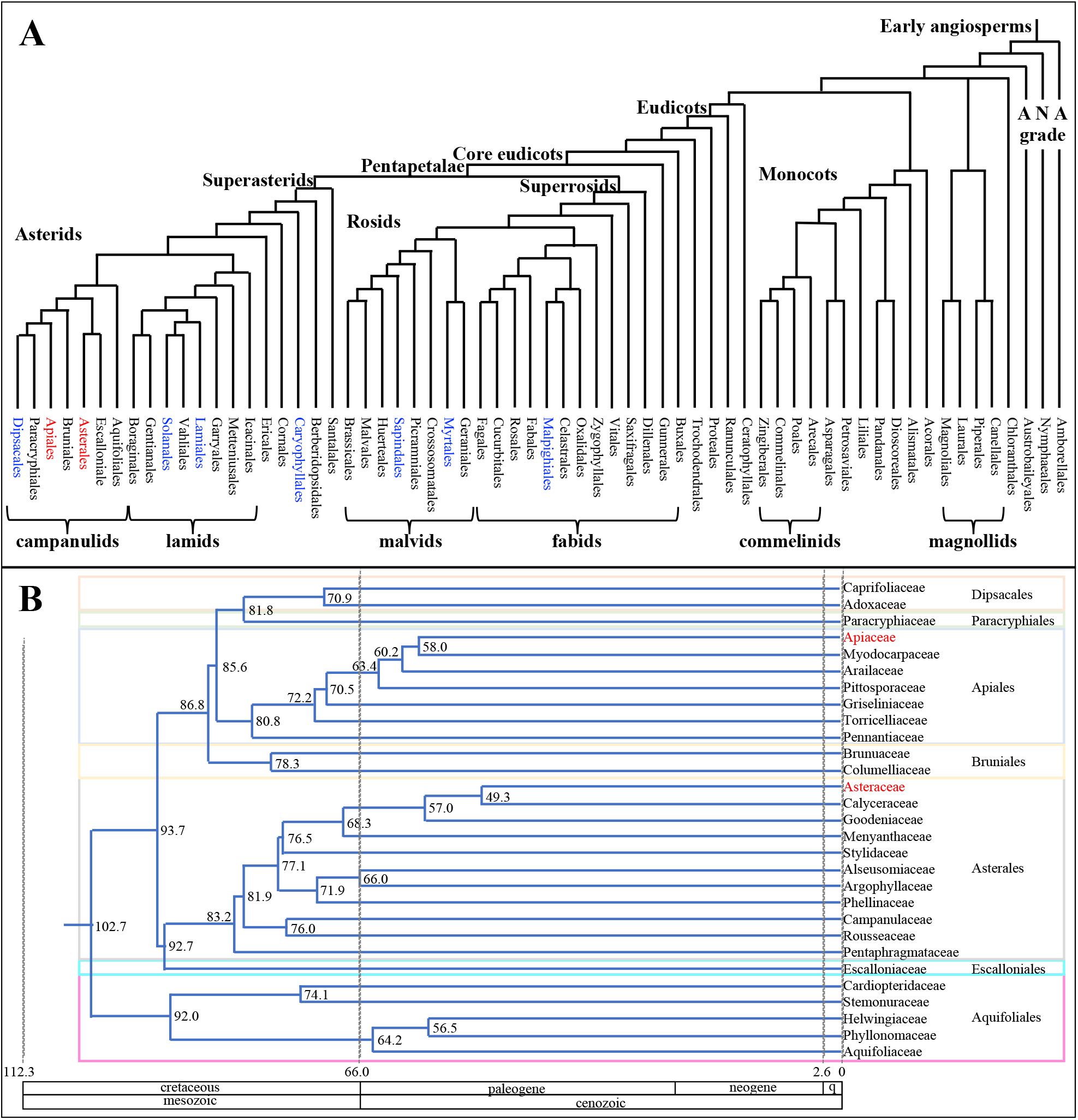
(**A)** Phylogenetic tree of Angiosperms (adopted from Cole *et al*., 2019). Orders with species producing linear sesquiterpene coumarins are in blue text; (**B)** Detailed phylogenetic tree of the clade Campanulids [adopted from Magallón *et al*., 2015]. The numbers show the estimated divergence time in million years ago (mya). Sesquiterpene coumarins are found in plants belonging to Apiacea and Asteraceae.

The sesquiterpene coumarins isolated from Apiaceae are all from plants belonging to the subfamily Apioideae. However, the phylogenic characterization of Apioideae is complicated since the major classifications do not agree between molecular phylogenetic and morphological studies, or even among different molecular phylogenetic analyses [Wen et al, 2021]. Regardless, to further evaluate the evolution of STCS a chronogram of the subfamily Apioideae based on 90 complete plastid genomes will be used [Wen et al, 2021]. This chronogram, presenting estimated divergence times, is shown in Figure 11A (Adapted from [Wen et al, 2021]). Tribes with plants producing sesquiterpene coumarins are shown in red. The genus *Magydaris* has not been ascribed to any tribe. However, *Magydaris* has been included in the *Opopanax* group, which is found in the Apioid superclade [Ajani et al, 2008]. The sesquiterpene coumarins producing tribes belong to different, clearly separated clades, indicating parallel evolution of STCSs. Only the linear sesquiterpene coumarin umbelliprenin (**1**) has been isolated from the taxa *Angelica, Herachum* and *Magydaris* indicating that these plants belonging to the Apiaceae family do not express any STCS. To produce umbelliprenin (**1**) from umbelliferone (**9**) and FDP (**27**) only a *O*-farnesyltransferase is required.

**Figure 11.**
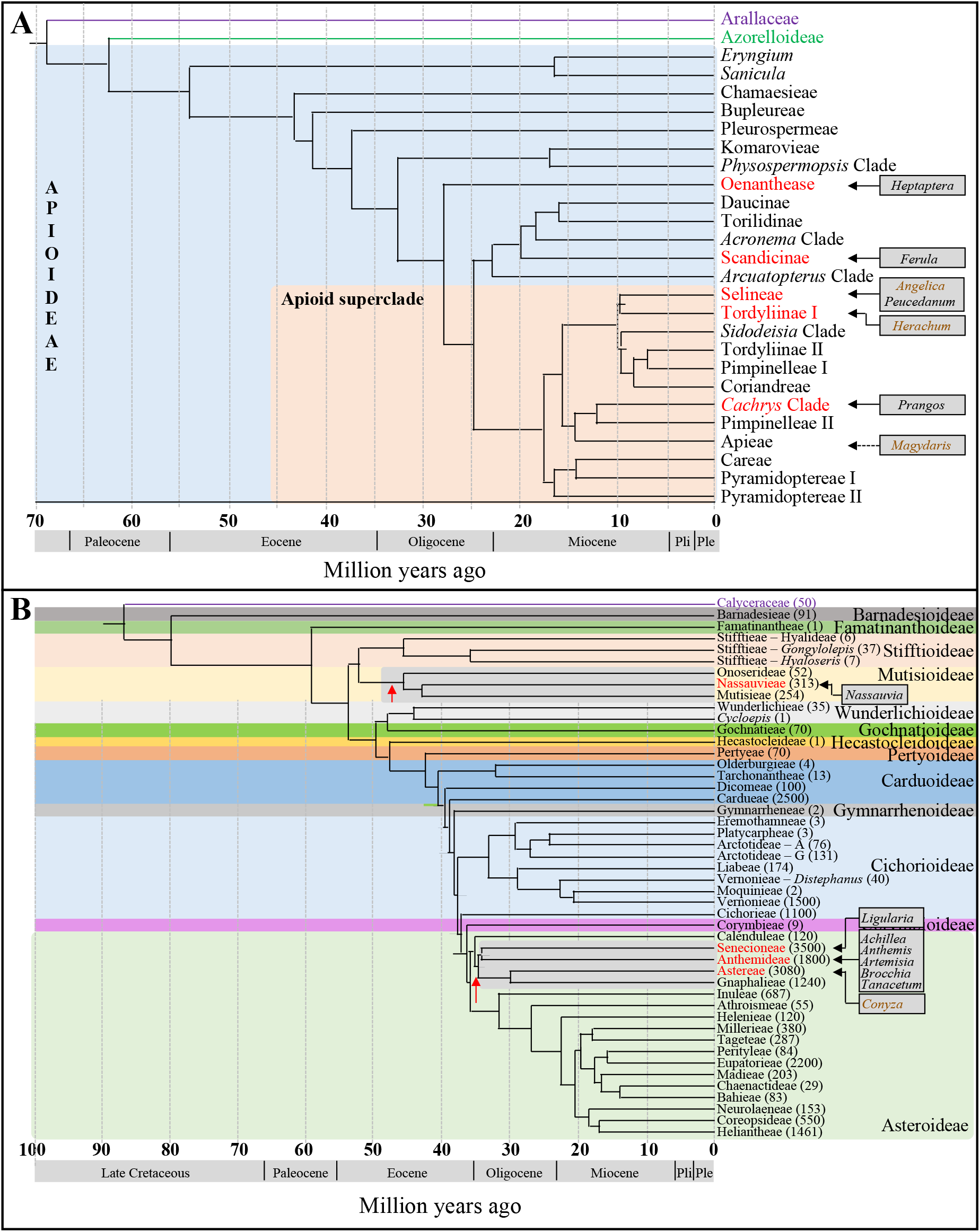
(**A**) Chronogram of the subfamily Apioideae (Apiaceae) presenting estimated divergence times (Adopted from Wen *et al*, 2021). This chronogram is based on 90 complete plastid genomes. Tribes with plants producing sesquiterpene coumarins are shown in red. The genus *Magydaris* has not been ascribed to any tribe. However, *Magydaris* has been included in the *Opopanax* group, which is found in the Apioid superclade [Ajani *et al*., 2008]. (**B**) Tribe-level chronogram of the Asteraceae family. The 12 subfamilies are colour coded. The estimated number of species in each tribe is shown in brackets. Tribes with plants producing sesquiterpene coumarins are shown in red. Genera are shown in boxes. The red arrows show possible time of gene duplication for sesquiterpene coumarin synthase development. [Adopted from Mandel *et al*., 2019]

With at least 25,000 named species and more than 1,700 genera, the backbone phylogeny of the Asteraceae family has been difficult to resolve. However, with increasing access to next generation sequencing technologies,resolving the phylogenies of megafamilies has become a reality. Highly resolved and well-supported nuclear phylogeny is now available. A tribe-level chronogram of the Asteraceae family is shown in Figure 11B. The 12 subfamilies are color coded. Tribes with plants producing sesquiterpene coumarins are shown in red. Taxa producing sesquiterpene coumarins are shown in boxes.

There are three tribes in the subfamily Mutisioideae. Sesquiterpene coumarins based on *O*-farnesylation have been isolated from the Nassauvieae tribe, while sesquiterpene coumarins based on *C*-farnesylation have been isolated from all three tribes of the subfamily Mutisioideae. It is interesting to note that the coumarin moiety of both *O*- and *C*-prenylated sesquiterpene coumarins isolated from subfamily Mutisioideae is 4-hydroxy-5-methylcoumarin (**18**). In the subfamily Asteroidae, some plants belonging to the tribes Senecioneae, Anthemideae and Astereae produce sesquiterpene coumarins. The taxon *Conyza* (Asteraceae) produces the linear sesquiterpene coumarin scopofarnol (**31**) from scopoletin (**10**) and FDP (**27**) indicating that no STCS is expressed. The most recent common ancestor of the two remaining tribes (Senecioneae and Anthemideae) indicates that parallel evolution of STCSs has occurred after a gene duplication around 34-35 mya.

Gene duplications have played a very important role in the evolution of plant metabolism. These duplication events can range from single genes to gene clusters to whole genomes. They have contributed to the evolution of novel functions in plants, such as the production of floral structures, induction of disease resistance, and adaptation to stress [Panchy et al, 2016]. Gene duplication has been an important mechanism for plants to generate the enormous diversity of terpenes [Hofberger et al, 2015]. One of the gene copies keeps its original function, while the other gene copy can go through a neofunctionalization resulting in a gene product with altered specificity for substrate and/or product. The expression pattern of the new gene will adapt to the needs of the new pathway in which it is involved.

For the evolution of sesquiterpene coumarin biosynthesis, it may be relevant to look at the evolution of SQE and OSCs. As shown in Figure 9A, umbelliprenin (**1**) fits very well in the active site of human SQE, which is an indication that umbelliprenin epoxidase may be closely related to SQE. The number of SQE and SQE-like proteins vary in different species. There is only one human *SQE* gene while there are six *SQEs* genes (*SQE1*-*SQE6*) in *Arabidopsis thaliana* [Rasbery et al, 2007]. However, to the best of our knowledge there are not any SQE-like enzymes characterized which use a substrate other than squalene (**20**). Functional analysis of the six Arabidopsis *SQEs* showed that *SQE1*-*SQE3* are squalene epoxidases while substrates and products for *SQE4*-*SQE6* could not be established [Rasbery et al, 2007]. Obviously, there are squalene epoxidase like enzymes expressed in *A. thaliana* with other functions than the epoxidation of squalene (**20**). It remains to be determined if any have adopted new catalytic functions [Philips et al, 2006].

The ancestral lanosterol synthase-like and cycloartenol synthase-like cyclases appeared about 140 million years ago before the differentiation of mono- and dicotyledonous plants. After differentiation, the lanosterol synthase-like gene was replicated multiple times, resulting in the expansion of the OSC genes in dicotyledons [Wang et al, 2022a]. The amplification of the OSC gene in the genomes of dicotyledons was mainly due to tandem duplications [Xue et al, 2012]. Tandem duplication may contribute to plant defense against biological and abiotic stresses. It is interesting to note that the function of some OSCs is unknown (2,3-oxidosqualene (**22**) is not a substrate), which illustrates that duplication of ancestral OSC genes led to the evolution of enzymes with altered substrate specificities as discussed above for SQE. This mechanism may have led to the evolution of cyclases involved in the biosynthesis of sesquiterpene coumarins (Figure 4B) and various meroterpenoids. An interesting observation in this respect is that Arabidopsis lupeol synthase efficiently converts 3-(14′,15′-oxidogeranylgeranyl)-indole (**58**) to the diterpeneindole petromindole (**63**) (Table 2) [Xiong et al, 2003]. An obvious experiment is to test if 10′,11′-oxidoumbelliprenin (**28**) is a substrate for the recombinant Arabidopsis lupeol synthase. We expect that a bicyclic sesquiterpene coumarin is obtained in such an experiment.

Another triterpene synthase that makes an intermediate bicyclic triterpene is marneral synthase from *Arabidopsis thaliana* [Xiong et al, 2006]. This enzyme, which will discussed below, has evolved through multiple gene duplications and is part of a biosynthetic gene cluster (BGC) [Xue et al, 2012]. It is evolutionarily closely related to thalianol and baruol synthases. During recent years it has been shown that BGCs have also evolved in other higher plants [Bharadvaj et al, 2021; Smit & Lichman, 2022]. Plant BGCs, consisting of between three to fifteen genes, have been identified and implicated in the biosynthesis of a diverse range of secondary metabolites. Notable examples include: the benzylisoquinoline and opiate alkaloids noscapine and thebaine in Papaver somniferum [Yang et al, 2021]; triterpenoids such as thalianin and cucurbitacin in *Arabidopsis thaliana* [Liu et al, 2020] and *Cucumis sativus* [Zhou et al, 2016b], respectively; and the triterpenoid-saponin avenacin in *Avena strigose* [Li et al, 2021].

Based on docking experiments (Figure 9), activity studies (Table 2) and inhibition studies (Table 3), we propose that STCE and STCS in Apiaceae and Asteraceae have their origin in the duplication of ancestral SQE and OSC genes, respectively. The enzyme functions are conserved while the active sites have evolved by amino acid substitutions to accommodate the new substrates, *i.e*., 7-farnesyloxycoumarins and 7-(10′,11′-oxidofarnesyloxy)-coumarins, respectively. In conclusion, we suggest that STCE is an FMO as other meroterpenoid synthases (Matsuda et al, 2016; Yuan et al, 2022). It appears that a parallel evolution of sesquiterpene coumarin biosynthesis has occurred in some tribes of the Apiaceae and Asteraceae families.

The coumarin moiety of sesquiterpene coumarins may reflect the availability of coumarins and/or the substrate specificity of the *O*-farnesylprenyltransferase. An example is the 5 sesquiterpene coumarins based on 4-hydroxy-5-methylcoumarin (**18**) (Table 1). These 5 compounds were isolated from two species belonging to the genus *Nassauvia*, as discussed above. 5-Methylcoumarins have been isolated from a few species of plants, but all from within the Asteraceae family. Furthermore, they are also restricted to representatives of the subfamily Mutisiodeae [Vestena et al, 2022]. In a similar way, umbelliferone (**9**) is the coumarin used in Apiaceae, while scopoletin (**10**) and isofraxidin (**12**) are used in Asteraceae (Table 1).

### 4.6. Desaturation

Some of the sesquiterpene coumarin carbon skeletons are modified by desaturation, most likely by P450 enzymes similar to desaturases involved in sterol biosynthesis, such as Δ^7^-sterol-C5(6)-desaturase [Taton & Rahier, 1996]. Δ^7^-Sterol-C5(6)-desaturase is membrane bound and catalyzes introduction of a C5 double bond into the B ring of Δ^7^-sterols to yield the corresponding Δ^5,7^-sterols (Figure 12A). Enzymatic activity requires molecular oxygen, NADH or NADPH, and membrane-bound cytochrome b5, which is an electron carrier from NAD(P)H to the desaturase through NAD(P)H cyt b5 reductase. Three examples of desaturation of sesquiterpene coumarins are shown in Figures 12B to 12D. Feselol (**37**) (***O*-U-Ha6**), colladonin (**36**) (***O*-U-Ia6**) and ferukrin (**74**) (***O*-U-Ja18**) are transformed to seravschanin B (**72**) (***O*-U-Hc1**), cauferidine (**73**) (***O*-U-Ic1**) and 5,6-ene-ferukrin (**75**) (***O*-U-Jc1**), respectively. A summary of desaturations of sesquiterpene coumarins is shown in Figure 13.

**Figure 12.**
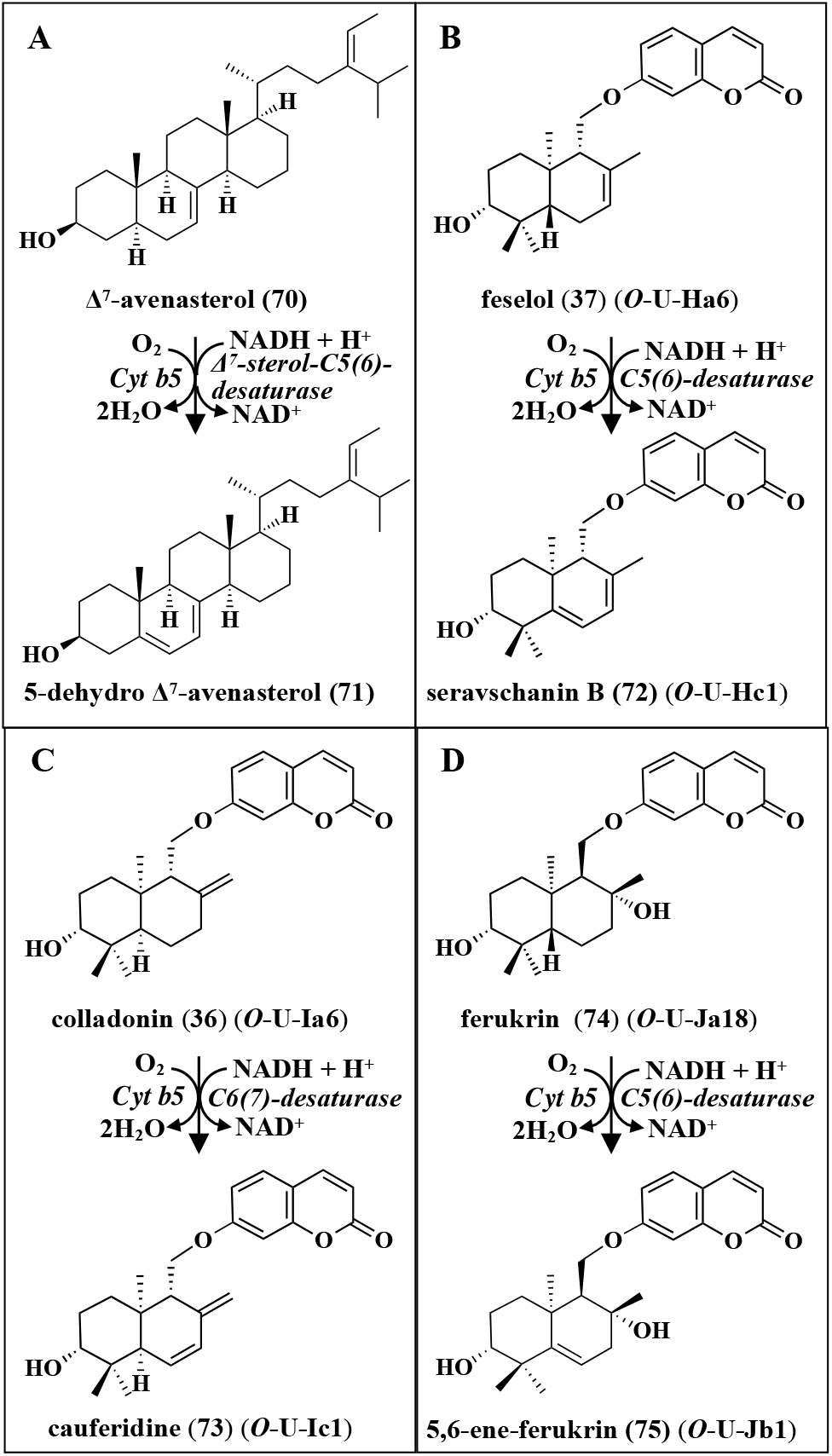
Examples of desaturation. (**A)** desaturation of the sterol Δ7-avenasterol (**70**) by Δ^7^-sterol-C5(6)-desaturase; (**B)** to (**D**) desaturation of three sesquiterpene coumarins by putative desaturases involving membrane bound cyt b5 as electron carrier.

**Figure 13.**
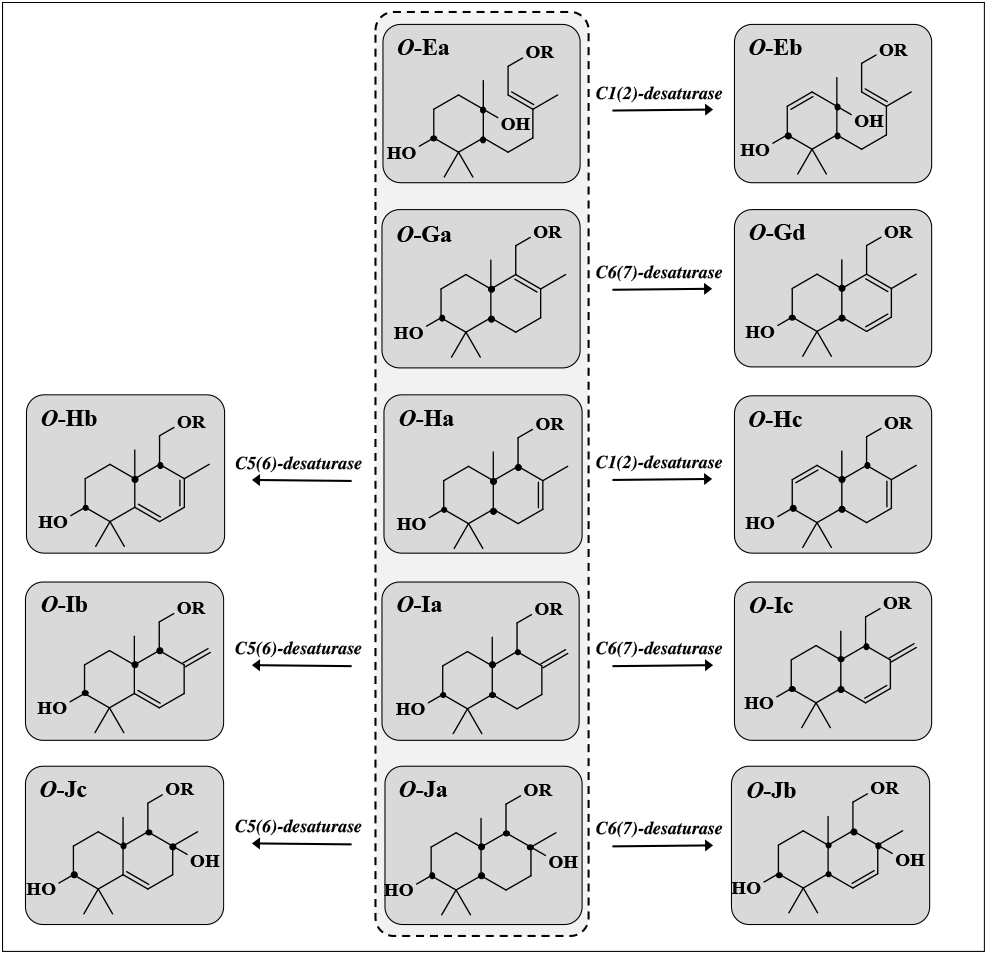
Summary of desaturations of 7-*O*-sesquiterpene coumarins by putative desaturases to generate the carbon skeletons ***O*-Eb, *O*-Gd, *O*-Hb, *O*-Hc, *O*-Ib, *O*-Ic, *O*-Jb**, and ***O*-Jc**

### 4.7 Peroxidation

Recently, the structural diversity of sesquiterpene coumarins was extended by a study on *Ferula bungeana*, which is found in Mongolia and Northern China [Guo et al, 2022]. Eight new sesquiterpene coumarins were isolated. Seven of these sesquiterpene coumarins are derived from conferol (**3**) with carbon skeleton ***O*-U-Ha1** (Figure S4) while one is derived from the sesquiterpene coumarin ***O*-U-Ha26** (**76**). These novel structures were the first examples of sesquiterpene coumarins possessing hydroperoxyl groups isolated as natural products. Hydroperoxide groups can be formed through either an autoxidation process or through the action of a dioxygenase. In both cases, oxygen is introduced into the substrate by a mechanism involving radicals. The mechanism of peroxidation has been studied on different plant metabolites such as fatty acids [Hajeyah et al, 2020], sterols [Porter, 2013] and other terpenoids [Nikolaiczyk et al, 2022]. Introduction of oxygen by autoxidation results in an epimeric mixture of the product, while introduction of oxygen by a dioxygenase gives a single epimer. The four sesquiterpene coumarins, ferubungeanol B (**77)**, ferubungeanol C (**78)**, ferubungeanol F (**79)**, and ferubungeanol H (**80**) with hydroperoxyl groups isolated from *F. bungeana* are epimers indicating that they have been produced by a dioxygenase as shown in Figure 14. In step 1, a hydrogen radical is regioselectivity abstracted from the sesqui-terpene coumarin by an active site tyrosyl radical. Next, rearrangement of the radical (step 2) occurs before oxygen is stereospecifically introduced by the enzyme (step 3). The hydroperoxide is formed through addition of hydrogen radical from the active site tyrosine residue (step 4), which restores the active site tyrosyl radical.

**Figure 14.**
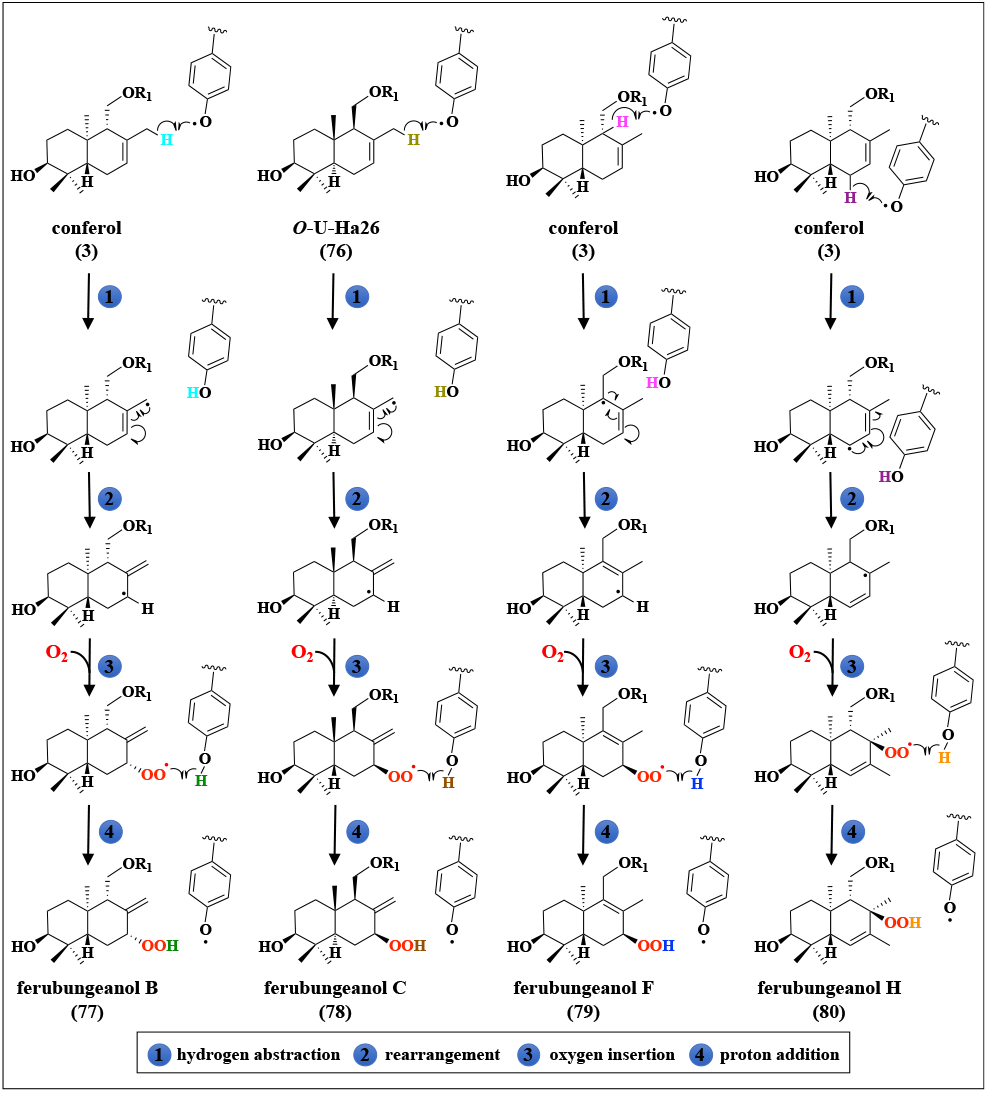
Putative reactions for the hydroperoxidation of conferol (**3**) and ***O*-U-Ha26** (**76**) in different positions by a dioxygenase involving an active site tyrosyl radical. The ***O*-Ha** carbon structure of the substrates is converted to ***O*-Ga** (**79**), ***O*-Ia** (**77**,**78**) and ***O*-Jb** (**80**) carbon skeletons.

However, nonenzymatic formation of the four sesquiterpene coumarins (**77-80**) as outlined in Figure 14 cannot be excluded. There are three steps involved in nonenzymatic peroxidation. The first step is generation of a terpene radical by abstraction of a hydrogen radical. In the second step, oxygen is introduced and in the final step a hydrogen radical is added to terminate the reaction and to give the epimeric hydroperoxides. As outlined in Figure 15, the two epimeric mixtures **75** and **76** are obtained by peroxidation of badrakemin (**81**) and turcicanol A (**82)**. One of the epimers of each epimeric mixtures (**83** and **84)** is the substrate of an enzyme converting the 7′-hydroperoxides to (*S*)-7′,8′-oxidoconferol (**83**) and (*R*)-7′,8′-oxidoconferol (**84**), respectivel. The conversions start with homolytic cleavage of the hydroperoxide groups and release of hydroxyl radicals. The sesquiterpene radicals are converted to epoxides and the reactions are terminated by addition of hydrogen radicals, possibly from tyrosine residues in the active sites, according to a postulated mechanism for the formation of 1,10-oxidovalencene from valencene-1-hydroperoxide [Nikolaiczyk et al, 2022]. Alternatively, formation of the intermediate 7′,8′-oxidoconferol epimers (**83** and **84**) can be obtained by epoxidation of conferol (**3**) by two monooxygenases w ith different product specificities. The 7′,8′-oxidoconferol epimers (**83** and **84**) are protonated to yield C7′-hydroxy groups that retain the stereochemical configuration of C7’ seen in their respective epoxide precursor. The resulting C8′-carbocations are quenched by proton abstraction from either C9′ or C12′ to give carbon skeleton ***O*-Ga** and ***O*-Ia**, respectively. The compounds produced are ferubungeanol A (**85**) (***O*-U-Ia32**) (Figure S5) and ferubungeanol E (**86**) (***O*-U-Ga2**) (Figure S4). Ferubungeanol G (**87**) may be formed by oxidation of the 7′hydroxyl group of ferubungeanol E (**86**) by a P450 (Figure 15). Alternatively, turcicanol A (**82**) (Figure 15) can be converted to ferubungeanol G (**87**) by an allylic oxidation with ferubungeanol E (**86**) as an intermediate product (see section 4.9).

**Figure 15.**
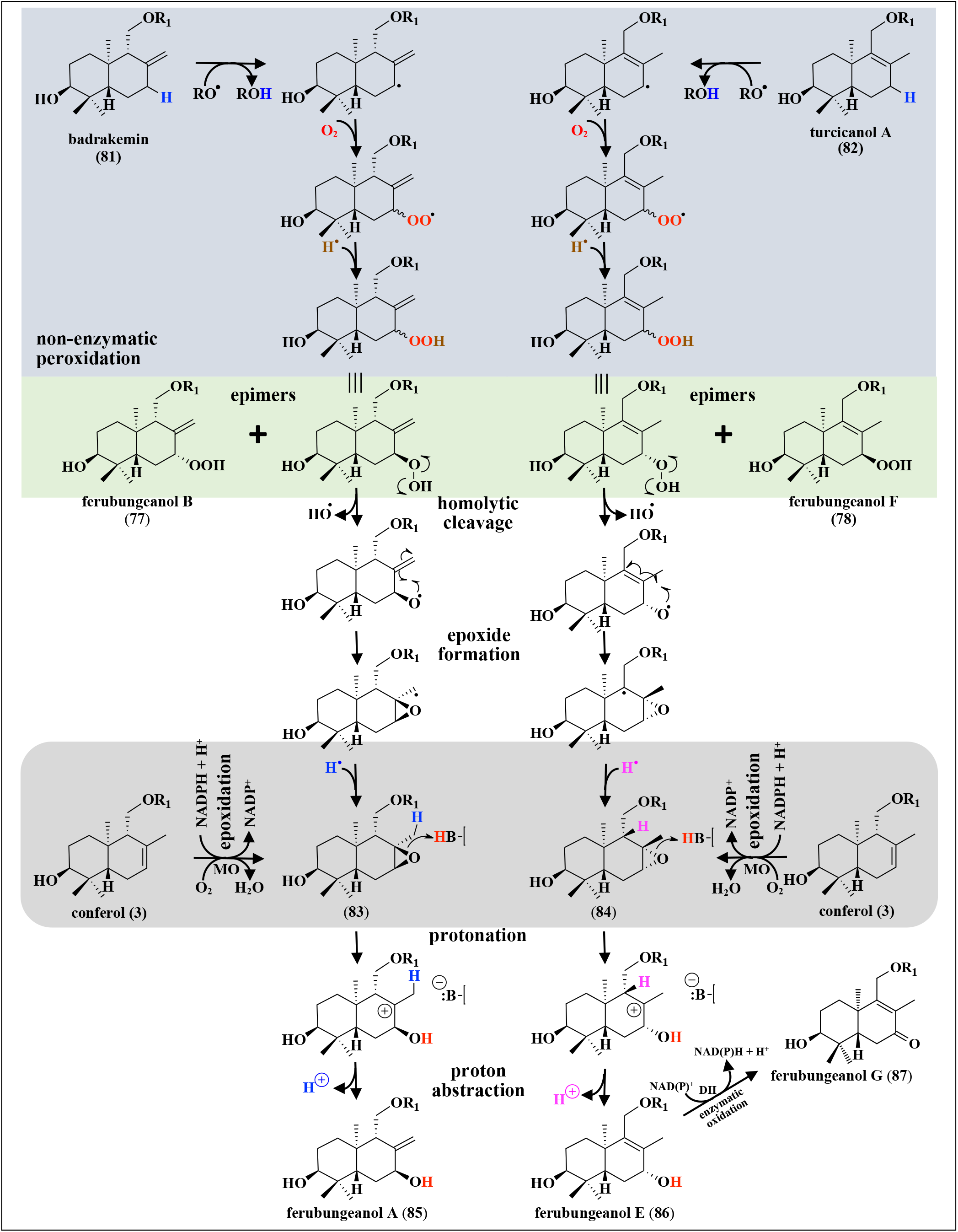
Nonenzymatic peroxidation of badrakemin (**81**) and tutcicanol A (**82**) yields epimeric mixtures of ferubungeanol B (**77**) and ferubungeanol F (**79**), respectively. One epimer of each mixture is enzymatically converted to 7′,8′(*S*)-oxidoconferol (**83**) and 7′,8′(*R*)-oxidoconferol (**84**), respectively. Alternatively 7′,8′(*S*)-oxidoconferol (**83**) and 7′,8′(*R*)-oxidoconferol (**84**) are obtained by epoxidation of conferol (**3**) by epoxidases with different product specificity. Protonation of 7′,8′(*S*)-oxidoconferol (**83**) and 7′,8′(*R*)-oxidoconferol (**84**) give carbocations, which can be converted to ferubungeanol A (**85**) (carbon skeleton *O*-Ia) and ferubungeanol E (**86**) (carbon skeleton *O*-Ia) by abstraction of proton H-12′ and H-9′, respectively.

### 4.8 Rearrangements

The diversity of sesquiterpene coumarins is increased by different rearrangements of the carbon skeletons of the two primary carbocations (**29, 30**) obtained in Figure 4. These rearrangements include hydride shifts and alkyl or aryl group migrations, which move the carbocation from one carbon to another.

As an example, a highly speculative reaction pathway starting from the monocyclic carbocation (**29**) leading to the formation of sinkiangenorin F (**88**) with carbon skeleton ***O*K** is shown in Figure 16. This pathway involves two methyl shifts and three 1,2-hydride shifts. An alternative route involving one 1,3-methyl shift is also shown (shaded area Figure 16). A third pathway involving one 1,3-alkyl shift is also included (framed). Two sesquiterpene coumarins with this carbon skeleton have been isolated from seeds of *Ferula sinkiangensis* (Figure S7) [Li et al, 2015a]

**Figure 16.**
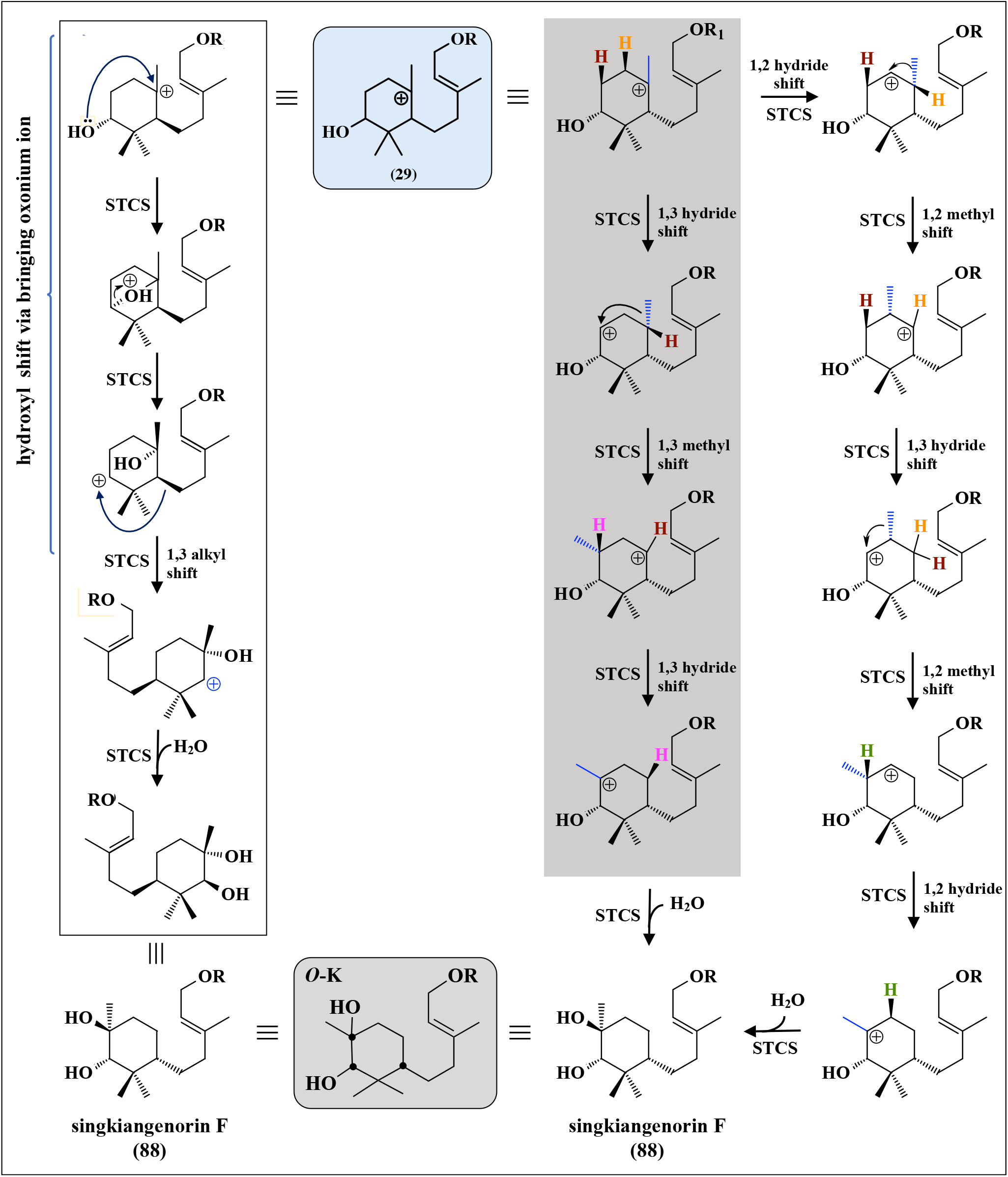
Alternative putative pathways for the biosynthesis of sinkiangenorin F (**88**) (carbon skeleton ***O*-K**) from the intermediate carbocation I (**29**) (see Figure 4). Chiral carbon atoms are indicated by black dots in the final structures. R = umbelliferone.

#### 4.8.a. Wagner Meerwein rearrangements

Wagner–Meerwein rearrangements are characterized by one or more 1,2-rearrangement reactions in which a hydrogen, alkyl or aryl group migrates from one carbon to a neighboring cationic carbon. These migrations result in rearrangement of the carbon skeleton of the sesquiterpene moiety and the formation of additional sesquiterpene coumarins [Lin et al, 2017; Quílez del Moral et al, 2020].

Classical examples of Wagner–Meerwein rearrangements are the conversion of 2,3-oxidosqualene (**22**) to lanosterol (**24**) and cycloartenol by lanosterol (LAS) and cycloartenol (CAS) synthase, respectively. Lanosterol (**24**) is a key intermediate in cholesterol biosynthesis in animals and cycloartenol in sitosterol biosynthesis in plants. The reactions catalyzed by these enzymes have been extensively studied [Nes, 2011; Chen et al, 2015; Diao et al, 2020]. First, the enzymes protonate the 2,3-oxidosqualene (**22**) to generate a C2 carbocation, as discussed above (Figure 4). This initiates cyclization reactions, which result in the formation of the intermediate tetracyclic protosterol cation (**23**). Next, a Wagner– Meerwein rearrangement involving two 1,2-hydride shifts and two 1,2-methyl shifts change the carbon skeleton to that of lanosterol (**24**). In the LAS reaction, the C8 carbocation is quenched by elimination of the proton from C9 to release lanosterol (**24**). In the CAS reaction, an additional 1,2-hydride shift followed by proton elimination from C19 leads to the formation of the cyclopropane ring of cycloartenol.

A putative reaction pathway leading to carbon skeletons ***O*-L** from the monocyclic carbocation (**29**) is shown in Figure 17A and pathways leading to carbon skeletons ***O*-M** to ***O*-Q** from the bicyclic carbocation (**30**) are shown in Figure 17B. All these pathways involve Wagner–Meerwein rearrangements of the decalin ring system. Cascades of 1,2-rearrangements move the carbocation in a stepwise fashion long distance. Carbon skeleton ***O*-L** (sinkianone (**89**)) is obtained from the monocyclic carbocation (**29**) by a Wagner–Meerwein rearrangement, which includes three 1,2-shifts (1,2-hydride, 1,2-methyl- and 1,2-hydride shifts). Abstraction of the proton from the 3′-hydroxyl group yields a keto-group on carbon 3′in the final product. The formation of carbon skeleton ***O*-N** (fnarthexol (**91**)) involves a Wagner–Meerwein rearrangement over five bonds of the decalin ring system (shown in the red frames). This cascade reaction involves five 1,2-shifts (both hydride and methyl) and is finally terminated by abstraction of the proton from C8′to obtain the 7′,8′-double bond of carbon skeleton ***O*-N**. The formation of carbon skeleton ***O*-O** involves a Wagner– Meerwein rearrangement over six bonds (shown in the blue frame). Abstraction of the proton from the 3′-hydroxyl group yields a keto-group on carbon 3′in the final product. The Wagner–Meerwein rearrangement of the bicyclic carbocation (**30**) leading to carbon skeleton ***O*-P** includes a 1,4-hydride shift, which is rare. The carbon skeleton ***O*-Q** is obtained from the bicyclic intermediate carbocation (**30**) through Wagner–Meerwein rearrangement involving two 1,2-hydride shifts followed by a 1,2-alkyl shift. The carbon skeleton is changed from a 6,6 bicyclic structure to a 6,7 bicyclic structure with a carbocation on C11′. This C11′ carbocation is subsequently moved to C3′ by one 1,4 hydride shift or by a combination of 1,2- and 1,3-hydride shifts. The carbocation is quenched by addition of water to yield the final product, sinkiangenorin E (**94**), which is a unique sesquiterpene structure [Li et al, 2016].

**Figure 17.**
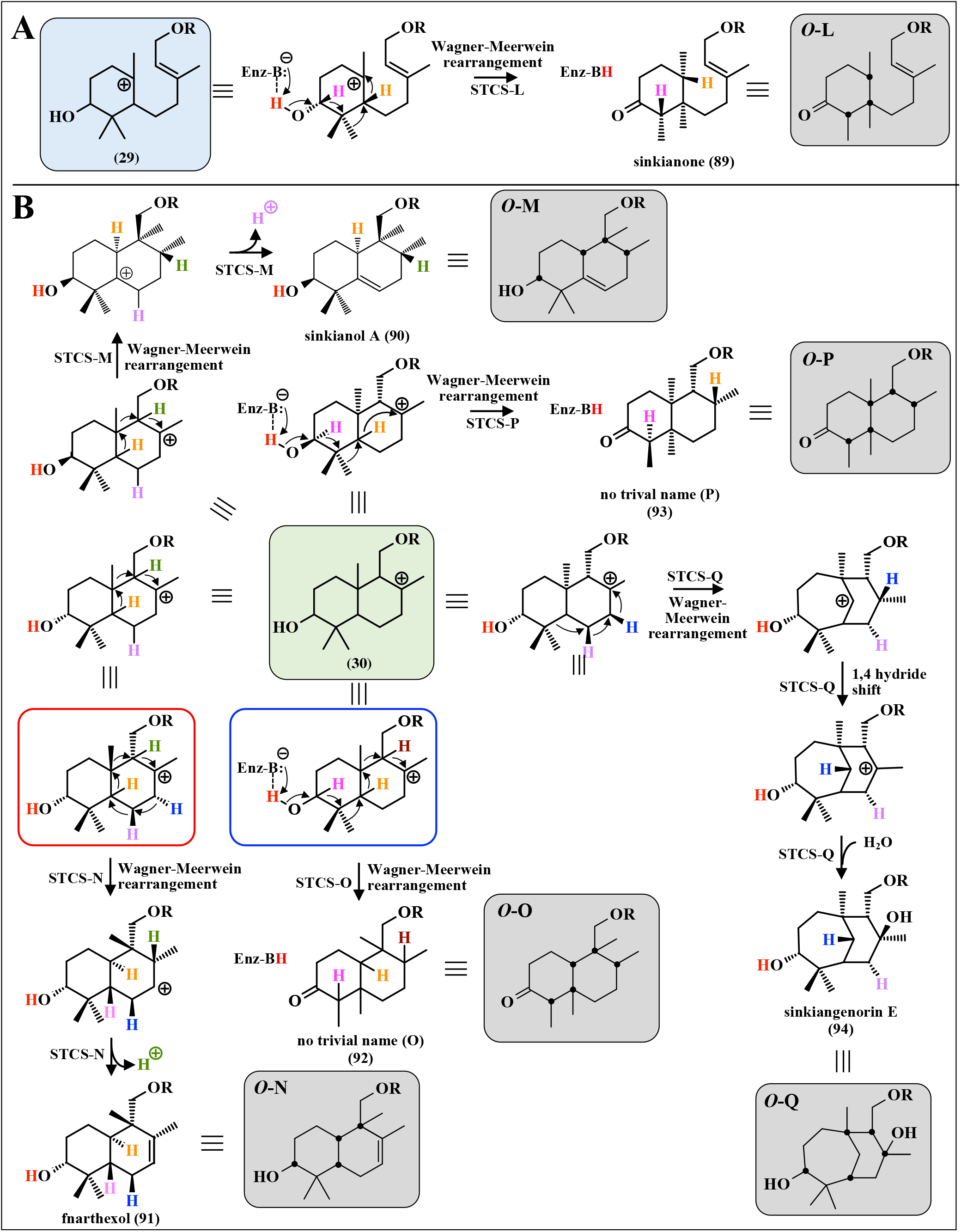
A: Putative pathway for the cyclization of intermediate carbocation I (29) (see Figure 4) to sesquiterpene coumarins with carbon skeletons *O*-L involving Wagner-Meerwein rearrangement. B: Putative pathways for the cyclization of intermediate carbocation II (30) (see Figure 4) to sesquiterpene coumarins with carbon skeletons *O*-L (sinkianone (89), *O*-M (sinkianol A (90), *O*-N (fnarthexol (91), *O*-O (no trivial name (O) (92)), *O*-P (no trivial name (P) (93)), and *O*-Q (sinkiangenorin E (94)) involving Wagner-Meerwein rearrangements. Chiral carbon atoms are indicated by black dots in the final structures.rmediate car

It is interesting to note that five different Wagner– Meerwein rearrangements of the bicyclic carbocation (**30**) are proposed in Figure 17B. These rearrangements ranging from two to six bonds are involved in the biosynthesis of carbon skeletons ***O*-M, *O*-N, *O*-O, *O*-P**, and ***O*-Q**. The structures of the sesquiterpene coumarins with these carbon skeletons are shown in Figure S7.

#### 4.8.b. Grob fragmentations

Grob fragmentations [Grob & Baumann, 1955] are important reactions for breaking C-C-bonds of the carbon skeletons of natural products. The reaction, named after Cyril A. Grob, is a concerted C-C bond cleavage involving a five-atom system [Lin et al, 2017]. An OSC will be used to illustrate the important Grob fragmentation reaction. The triterpene marneral (**95**) has only recently been isolated from a natural source. It was for a long time assumed to be an intermediate in the biosynthesis of iridals in plants of the Iridaceae (Lily) family [Marner et al, 1989; Hasegawe et al, 2011]. A gene from Arabidopsis (*MRN1*) encoding an enzyme that produces marneral (**95**) has been cloned and the recombinant protein characterized [Xiong et al, 2006; Go et al, 2012]. The enzymatic reaction of MRN1 proceeds as outlined in Figure 18A through the cyclization of 2,3-oxidosqualene (**22**) to a bicyclic intermediate, which undergoes a Wagner-Meerwein rearrangement to generate a C5 cation. Subsequently, a Grob fragmentation of the A-ring cleaves the carbon-carbon bond between C3 and C4 to generate a monocyclic aldehyde, marneral (**95**). The production of α- and β-seco-amyrin by a recombinant OSC from Arabidopsis involves Grob fragmentation of the C-ring [Shibuya et al, 2007]. A seco-triterpene suggested to be produced by Grob fragmentation is sasanquol, which is a by-product in the formation of baccharide oxide by baccharide oxide synthase from *Stevia rebaudiana* [Shibuya et al, 2008]. *MRN1* is part of a three gene cluster found on chromosome 5 of *Arabidopsis thaliana* [Field et al, 2011]. In addition to *MRN1*, a marneral oxidase (*Cyp71A16/MRO*) [Kranz-Finger et al, 2018] and an uncharacterized gene (*At5g42591*) constitute this gene cluster. These genes are coexpressed in *A. thaliana* and marneral (**95**) is not accumulated as it is hydroxylated by marneral oxidase to 23-hydroxymarneral.

**Figure 18.**
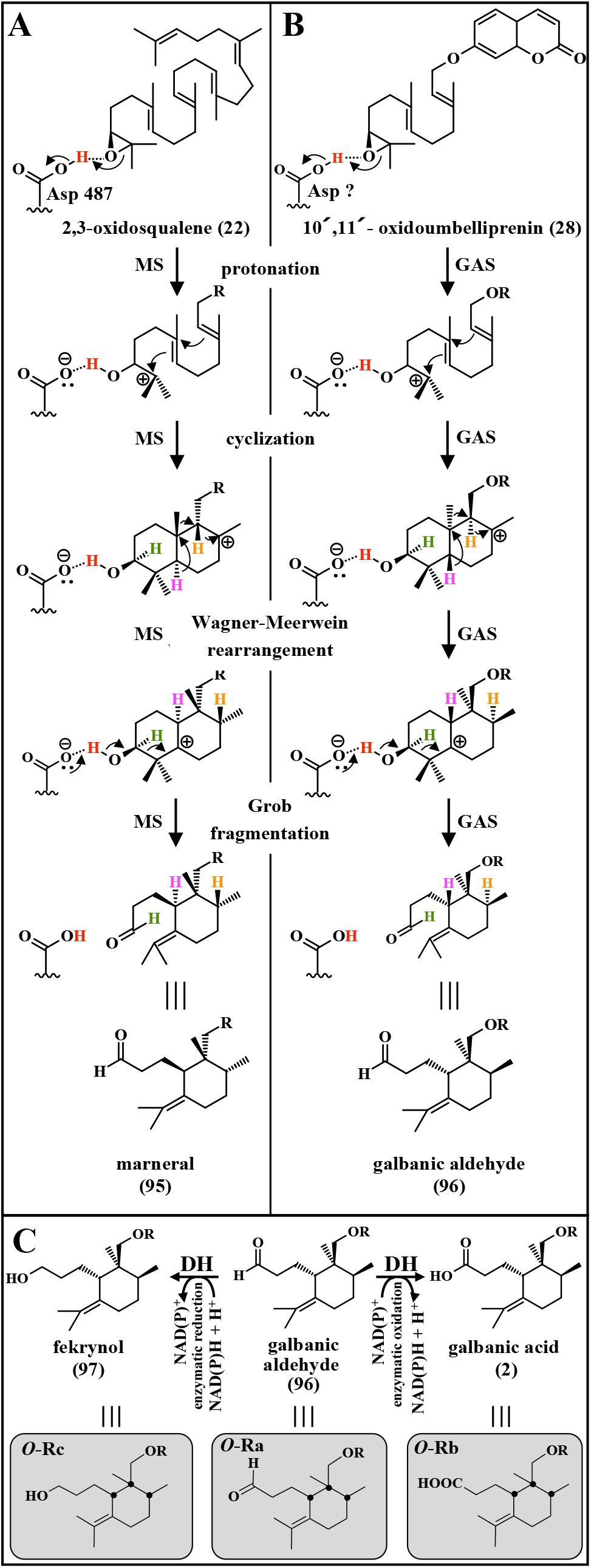
Comparison of the biosynthesis of the triterpene marneral (**95**) (**A**) from 2,3-oxidosqualene (**22**) and galbanic aldehyde (**96**) (**B**) from 10′,11′-oxidoumbelliprenin (**28**) involving Grob fragmentation reactions. MS: marneral synthase; GAS: galbanic aldehyde synthase. (**C**) Putative enzymatic reduction and oxidation of galbanic aldehyde (**96**) to fekrynol (**97**) and galbanic acid (**2**), repectively.

In analogy with the marneral pathway, a hypothetical galbanic aldehyde synthase (GAS) converts 10′,11′-oxidoumbelliprenin (**28**) to the bicyclic intermediate (**30**) (Figure 18B). A Wagner-Meerwein rearrangement generates a C5′ cation, which undergoes Grob fragmentation to cleave the C3′-C4′-bond forming the 3,4 secodrimane skeleton in the same way as has been shown for marneral (**95**). The galbanic aldehyde (**96; *O*-U-Ra1**) formed can be enzymatically oxidized to the carboxylic acid (galbanic acid (**2**); ***O*-U-Rb1**; Figure S8) or enzymatically reduced to the hydroxyl derivative (fekrynol (**97**); ***O*-U-Rc1**; Figure S8) by NAD(P)^+^-dependent oxidoreductases as outlined in Figure 18C. Galbanic acid (**2**) has been isolated from various species of the *Ferula* genus. It exhibits a variety of interesting biological activities. The cloning of a *GAS* gene from a *Ferula* species may be possible using the information obtained from *MRN1*.

Two different proposals for the biosynthesis of galbanic acid (**2**) have been presented. Marner and Kasel [1995] suggested a mechanism involving a Grob fragmentation which is in accordance with the mechanism suggested here. Appendino et al [1993] suggested a biosynthesis of galbanic acid (**2**) starting from mogoltadone (***O*-U-Ia29**). It is highly likely that galbanic aldehyde (**96**) is biosynthesized in analogy with marneral (**95**) as suggested here.

In addition to carbon skeleton ***O*-R**, we suggest that the formation of carbon skeletons ***O*-S, *O*-T**, and ***O*-U** involve Grob fragmentations as shown in Figure 19. Ferusingensine A (**98**) with carbon skeleton ***O*-S** is obtained from the monocyclic carbocation (**29**) by a Grob fragmentation followed by reduction of the aldehyde by an oxidoreductase (Figure 19A). Formation of the sinkiangenorin D (**100**) with carbon skeleton ***O*-U** and ferulsinaic acid (**99**) with carbon skeleton ***O*-T** include 1,2-alkyl shifts for expansion (sinkiangenorin D (**100**)) or reduction (ferulsinaic acid (**99**)) of the ring size (Figure 19B). Both these carbon skeletons are obtained by Grob fragmentation of the bicyclic C8′ intermediate carbocation (**30**). In the case of ***O*-S**, the C8′ carbocation is moved to C4′ through a Wagner-Meerwein rearrangement. Next, an alkyl shift leading to a 5,7-ring structure prepares the molecule for the Grob fragmentation reaction, and the resulting aldehyde is reduced by an oxidoreductase to give the final product. In the paper reporting the isolation of sinkiangenrin D (**100**) (***O*-U-U1**; Figure S8), a pathway for its formation is proposed [Li et al, 2015a]. This pathway as outlined in Figure 19C most likely requires a second sesquiterpene synthase. It starts by protonation of fekrynol (**97**) (***O*-U-Rc1**; Figure S8) followed by an alkyl shift before termination of the reaction by proton abstraction.

**Figure 19.**
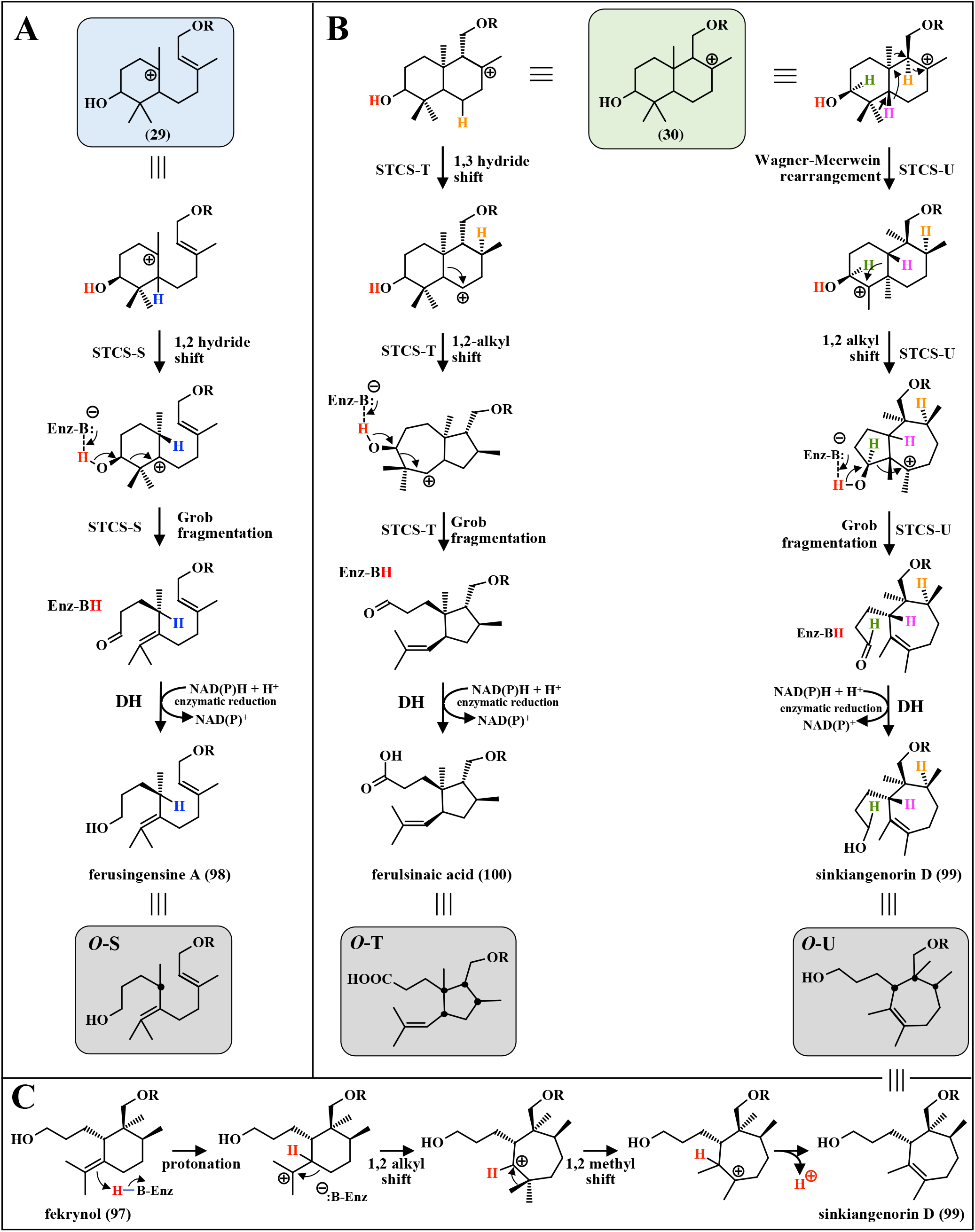
Putative pathways involving Grob fragmentation reactions for the formation of sesquiterpene coumarins. **A:** Formation of carbon skeleton ***O*-S** (ferisingensine A (**98**)) from intermediate carbocation I (**29**). **B:** Formation of carbon skeletons ***O*-T** (ferulsinaic acid (**98**)) and ***O*-U** (sinkiangenorin D (**99**)) from intermediate carbocation II (**30**). **C**: Alternative pathway for the formation of sinkiangenorin D (**99**) from fekrynol (**97**) as suggested by [Li et al, 2015a]. The formation of intermediate carbocation I (**29**) and intermediate carbocation II (**30**) is shown in Figure 4. Chiral carbon atoms are indicated by black dots in the final product.

Biosynthesis of ***O*-T** from the bicyclic carbocation (**30**) is initiated by an 1,3 hydride shift followed by an alkyl shift, which prepares the molecule for Grob fragmentation. In this case, the aldehyde formed is oxidized to the carboxylic acid by an oxidoreductase. Ferulsinic acid (**99**) (***O*-U-T1**) is the only sesquiterpene coumarin with carbon skeleton ***O*-T** [Ahmed et al, 2007]. A biosynthetic route starting from feselol (**37**) (***O*-U-Ha7**) was suggested. The suggested reaction pathway included oxidation of the C3′-hydroxyl group, hydroxylation at C6′ followed by acetylation before the Grob fragmentation without participation of any preceding carbocation rearrangements. This is a highly unlikely biosynthesis of the ***O*-T** carbon skeleton. The biosynthesis of karatavic acid (**136**) is one more example of a Grob fragmentation, which will be discussed later in this review.

Until recently all isolated sesquiterpene coumarins with 3′,4′ secodrimane skeletons (***O*-R**) exhibited the same stereochemistry (Figure S8). However, recently five new sesquiterpene coumarins with 3′,4′ secodrimane moieties were isolated from *F. sinkiangensis* [Wang et al, 2023]. The carbon skeletons of these five sesquiterpene coumarins named fesinkin A to D are diastereomers of fekrynol (**97**); ***O*-U-Rc1**) and are formed in a similar way as shown for galbanic aldehyde (**96**) in Figure 18B. Putative pathways for the biosynthesis of fesinkin A (**101**), fesinkin C (**102**), 4′Z fesinkin D (**103**) and 4′E fesinkin D (**104**) from the bicyclic carbocation (**30**) are shown in Figure 20. 4′Z fesinkin D (**103**) and 4′E fesinkin D (**104**) are derivatives of fekrynol diastereomer (**107**). The carbon skeleton of fesinkin E (**105**) and fesinkin F (**106**) is ***O*-U-Ga** (Figure 7B and Figure S4). Recently, a sesquiterpene coumarin named turicanol B was isolated from *Ferula turcica* [Erucar et al, 2023]. The structure of turicanol B is the same as that of fesinkin E (**105**).

**Figure 20.**
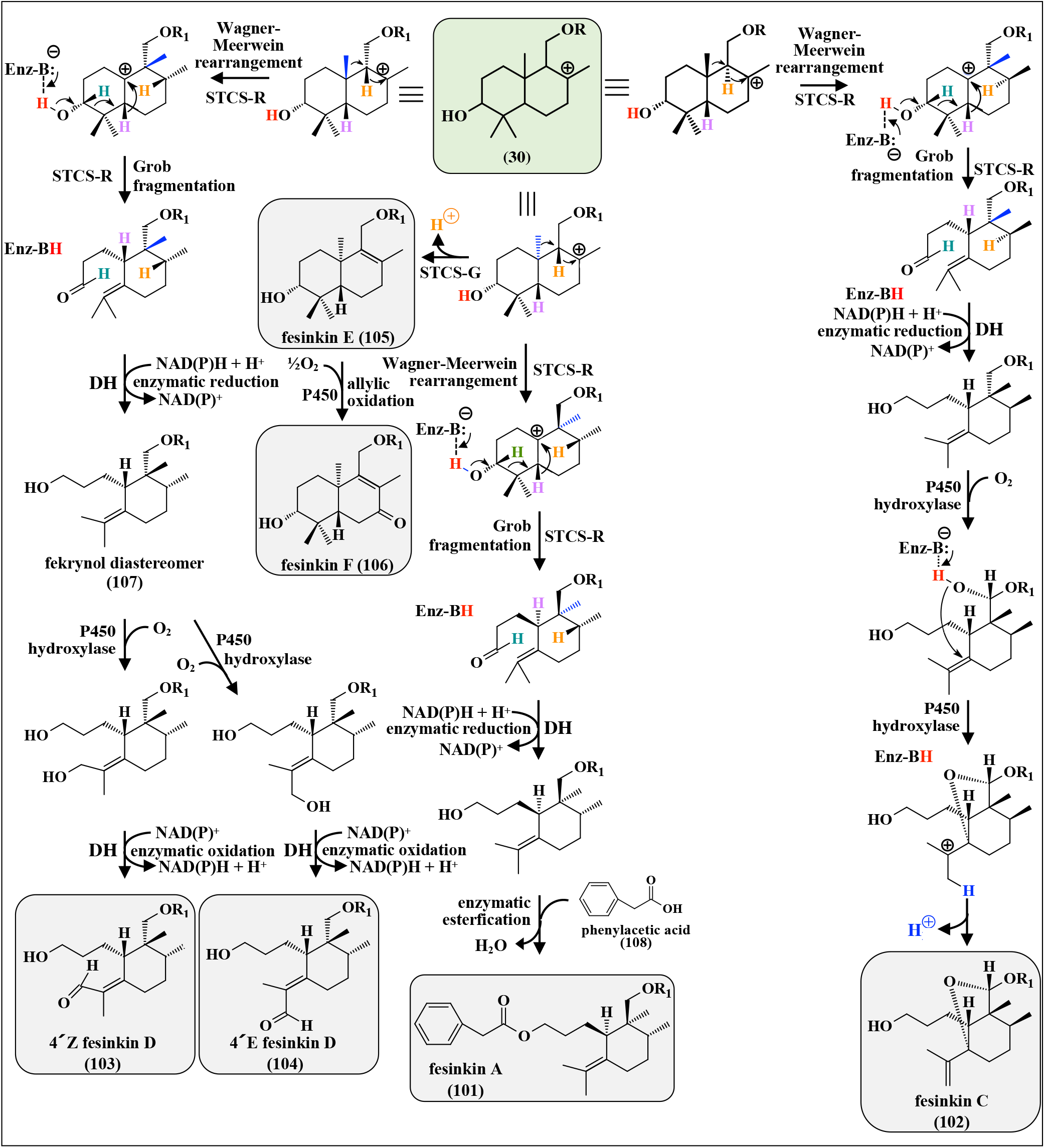
Putative pathways for the biosynthesis of fekrynol (97), fesinkins A (99), C (101) and D (102 and 103), E (104) and F (105), fekrynol (95) and fekrynol diastereomer (107) from intermediate carbocation II (30) (see Figure 4).

Fesinkins A to D are the first examples of 3′,4′ secodrimane skeletons with different stereochemistry from that of galbanic acid (**2**), which extend the structural diversity of sesquiterpene coumarins. Fesinkin B (**109**) is a hydroperoxide sesquiterpene coumarin. Some hydroperoxyl sesquiterpene coumarins isolated from *Ferula bungeana* have been discussed above (Figures 14 and 15) [Guo et al, 2022]. A putative pathway for the peroxidation of fekrynol diastereomer (**107**) is shown in Figure 21A. During the radical reactions, the 4′,5′-double bond of the fekrynol diastereomer (**107**) is converted to the 4′,14′-double bond of fesinkin B (**109**). Finally, fesinkin C (**102**) has a unique 5′,11′-oxido bridge and fesinkin G (**110**) a very unusual endoperoxide bridge.

**Figure 21.**
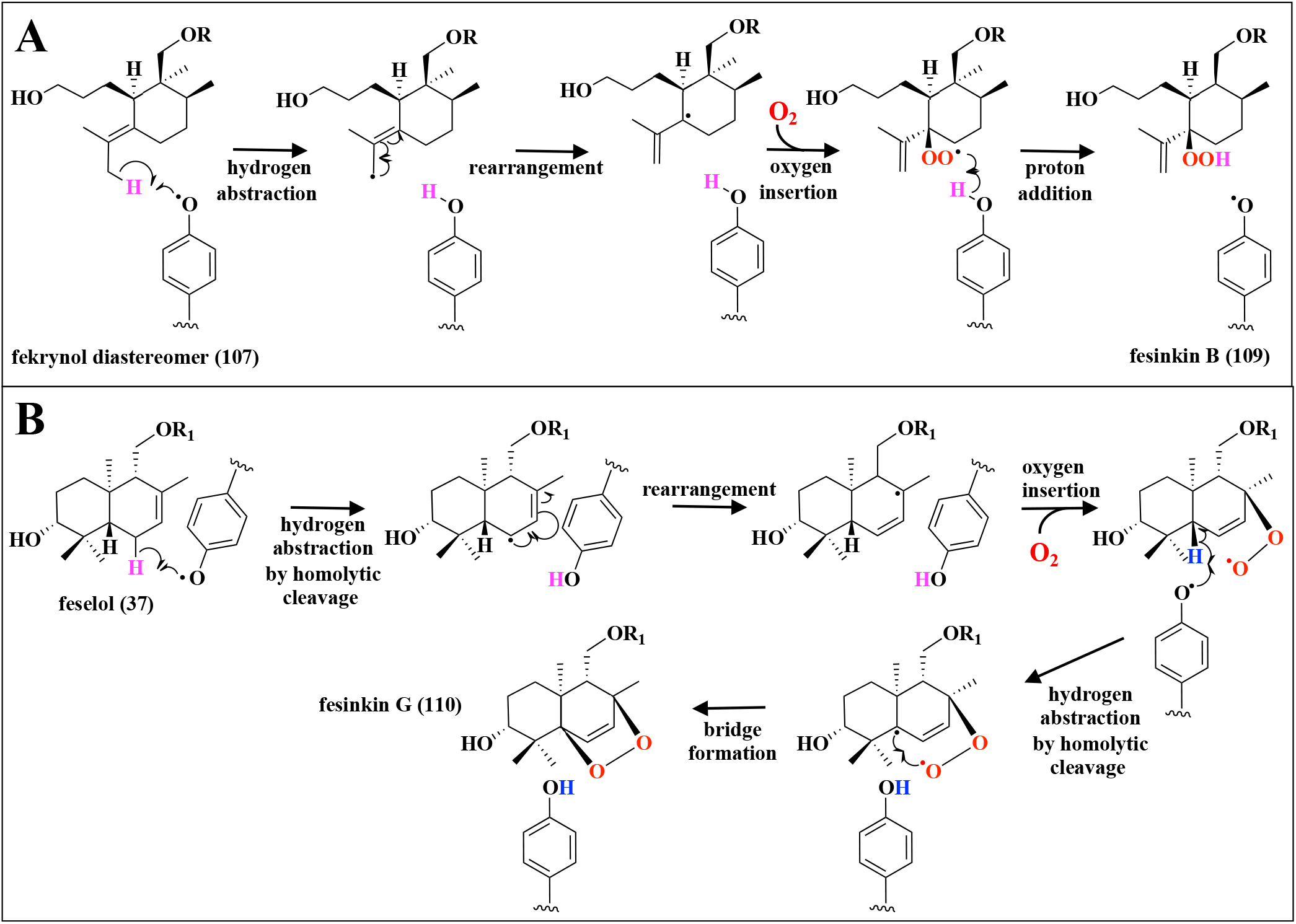
**A**. Putative biosynthesis of fesinkin B (**109**) from the fekrynol diastereomer (**107**). **B**. Putative pathway for the biosynthesis of the 5′,8′-endoperoxide sesquiterpene coumarin fesinkin G (**100**) from feselol (**37**).

Fesinkin G (**110**) (***O*-U-Jb4**) is the first sesquiterpene coumarin endoperoxide that has been isolated [Wang et al, 2023]. It may be obtained by peroxidation of feselol (**37**) (***O*-U-Ha7**), which was co-isolated with fesinkin G (**110**) from *Ferula sinkiangensis*. A putative pathway, based on the peroxidation of conferol (**3**) as outlined in Figure 14, is shown in Figure 21B. This pathway starts by homolytic cleavage of the C6′-hydrogen bond of feselol (**37**) by an active site tyrosyl radical followed by rearrangement to generate the 6′,7′-double bond and a radical on C8′. Oxygen is inserted to obtain the 8′-peroxide. Instead of proton addition, a second homolytic cleavage of the C5′-hydrogen bond prepares for the formation of the endoperoxide by bond formation via fusion of the two radicals.

### 4.9. Biosynthesis of a sesquiterpene coumarin not involving epoxidation

The biosynthesis of sesquiterpene coumarins lacking a 3′-hydroxyl group does not involve the epoxidation of umbelliprenin. 10′,11′-Oxidoumbelliprenin (**28**) is not a substrate. However, this is extremely rare. An example is the biosynthesis of a sesquiterpene coumarin in *Ferula galbaniflua* as shown in Figure 22A [Graf & Alexa, 1985]. The terminal double bond of umbelliprenin (**1**) is protonated to generate a carbocation that can go through cyclizations to a bicyclic sesquiterpene moiety. The reaction is terminated by abstraction of a proton. However, this sesquiterpene coumarin has not been isolated. The carbon skeleton of this sesquiterpene coumarin has been assigned ***O*-U-Hd**. We suggest that the last step of biosynthesis of sesquiterpene coumarin ***O*-U-Hd1 (111)** involves an allylic oxidation as shown in Figure 22A.

**Figure 22.**
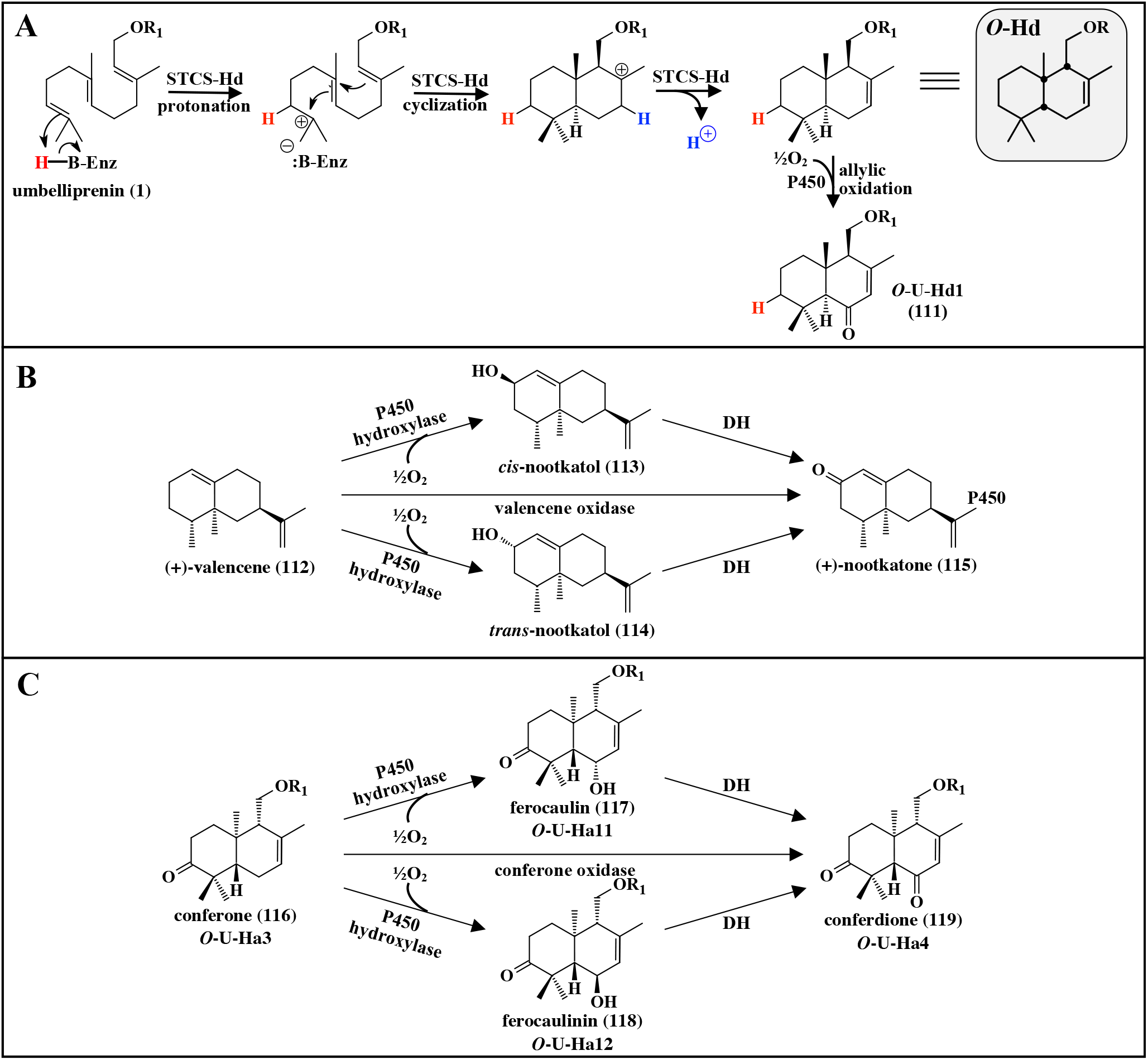
**A:** Putative biosynthesis of a sesquiterpene coumarin (**111**) (**O-U-Hd1**) not involving epoxidation of umbelliprenin. The final step, allylic oxidation, is catalyzed by a putative P450 oxidase. **B:** One step oxidation of (+)-valencene (**112**) to (+)-nootkatone (**115**) by valencene oxidase (VO) or two step oxidation by P450 hydroxylase and NAD(P)H-dependent oxidoreductase (dehydrogenase DH). **C:** Putative allylic oxidation of conferone (**116**) to conferdione (**119**) in *Ferula conocaula*.

Allylic oxidation of terpenoids has been studied in various organisms including plants [Wang et al, 2022b].

Allylic oxidation of (+)-valencene (**112**) to (+)-nootkatone (**115**) as outlined in Figure 22B has been extensively investigated. (+)-Nootkatone (**115**), with a grapefruit-like flavor, is a sesquiterpene found in *Citrus paradisi, Citrus maxima* (Rutaceae), and *Callitropsis nootkatensis* (Cupressaceae) widely used in the food, flavor and fragrance industries [Zhang et al, 2023a]. A two-step enzymatic conversion of (+)-valencene (**112**) to (+)-nootkatone (**115**) has been proposed. The first step is a regioselective allylic hydroxylation of the 2-position of (+)-valencene (**112**) to *cis*-nootkatol (**113**) or *trans*-nootkanol (**114**), followed by the oxidation to (+)-nootkatone (**115**) (Figure 22B) [Fraatz et al, 2009]. A single multifunctional cytochrome P450 enzyme may catalyze both reaction steps. Alternatively, first (+)-valencene (**112**) is hydroxylated to nootkatol (**113** or **114**) by a cytochrome P450 enzyme with subsequent oxidation to (+)-nootkatone (**115**) by a NAD(P)H-dependent oxidoreductase. Some cytochrome P450 enzymes from the CYP71 family have been shown to exhibit valencene oxidase activity. Premnaspirodiene oxygenase CYP71D55 from *Hyoscyamus muticus* (Solanaceaae) was shown to convert (+)-valencene (**112**) to *trans*-nootkatol (**114**) [Takahashi et al, 2007]. Co-expression of valencene oxidase CYP71AV8 from *Cichorium intybus* (Asteraceae) with valencene synthase in yeast, resulted in the formation of *trans*-nootkatol (**114**) and small quantities of (+)-nootkatone (**115**) [Cankar et al, 2011]. Finally, valencene oxidase CYP706M1, which belongs to the CYP706 family, was cloned from *Callitropsis nootkatensis* (Alaska cedar) (Cupressaceae) [Cankar et al, 2014]. It was shown that this cytochrome P450 enzyme converts (+)-valencene (**112**) to (+)-nootkatone (**115**).

We postulate that some sesquiterpene coumarins are modified by allylic oxidation of the sesquiterpene moiety by cytochrome P450 enzymes similar to the enzymes converting (+)-valencene (**112**) to (+)-nootkatone (**115**). An example of putative allylic oxidation of a sesqui-terpene coumarin is shown in Figure 22C. The four sesquiterpene coumarins shown (**116** to **119**) were all isolated from *Ferula conocaula* [Vandyshev et al, 1972a, 1972b; Kuliev & Khasanov, 1978]. In this case, conferone (**116**) is hydroxylated by a P450 hydroxylase belonging to the CYP71 family. The intermediate product ferocaulin (**117**) and/or ferocaulinin (**118**) are/is subsequently oxidized to conferdione (**119**) by a NAD(P)H-dependent oxidoreductase. Alternatively, the reaction can be catalyzed by an oxidase belonging to the CYP706 family.

Examples of other sesquiterpene coumarins, which may have been obtained by allylic oxidations are shown in Figure 23. These allylic oxidations occur in different *Ferula* species, *i.e. F. gummosa, F. galbaniflua, F. conocaula*, and *F. sinkiangensis*, as shown in Figure 23A to 23D. In these cases, no intermediate hydroxylated sesquiterpene coumarins have been isolated, which indicates that the allylic oxidation is catalyzed by oxidases belonging to the CYP706 family. The two reactions in *F. gummosa* (Figure 23A) may be catalyzed by an oxidase with broad substrate specificity.

**Figure 23.**
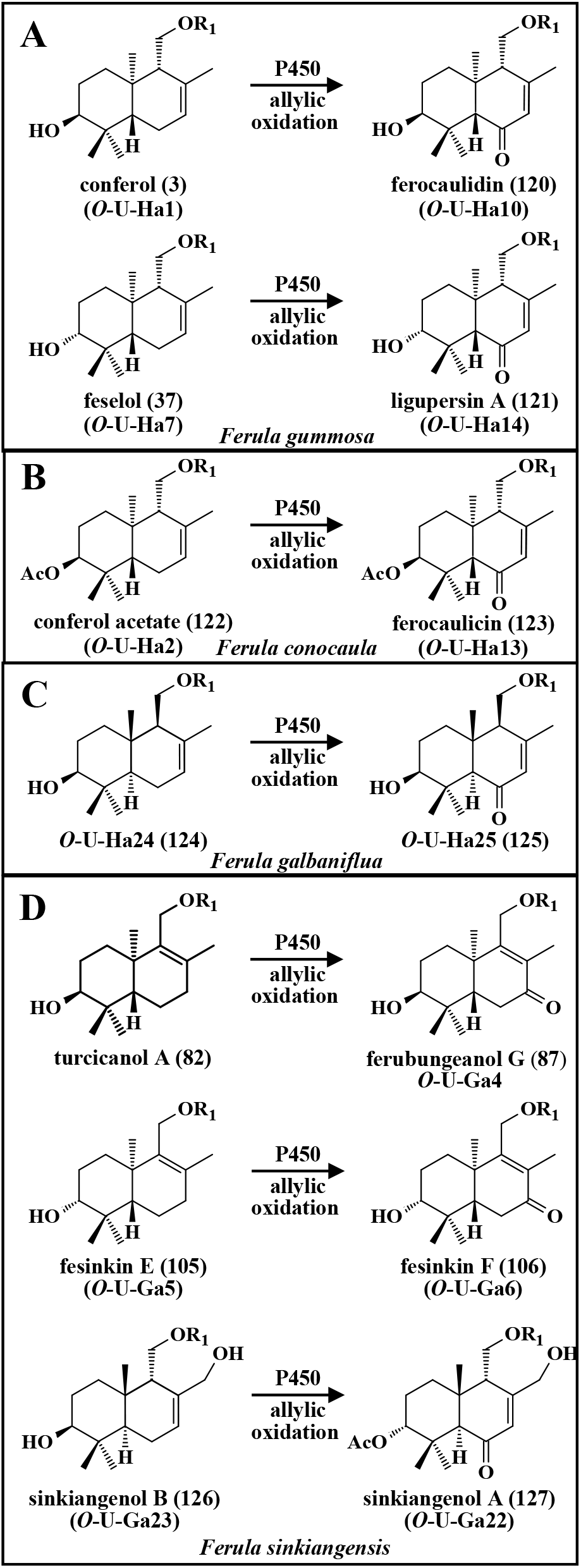
Examples of putative allylic oxidation of sesquiterpene coumarins in various *Ferula* species.

Another example of a sesquiterpene coumarin without the 3′-hydroxyl group on the sesquiterpene moiety is structure ***O*-V** (Figure 3). The putative pathway leading to structure ***O*-V**, shown in Figure 24A, starts with 4′-hydroxylation of umbelliprenin (**1**) by a P450 hydroxylase. 4-Hydroxyumbelliprenin (**126**) is a unique structure. It has not been isolated. Protonation of the 2′,3′-double bond of 4′-hydroxyumbelliprenin (**126**) leads to ionization and a carbocation on C-2′. Two ring closures (2′,7′- and 6′,10′) generate the 5,6-ring structure of ferusinol (**4**) [Bandyopadhyay et al, 2006]. Next, a proton is abstracted to yield the final product. The STCS catalyzing this reaction is most likely of a different type to the STCSs using 10′,11′-oxidoumbelliprenin (**28**) as a substrate; a distinction analogous to the comparison between SHCs and OSCs in triterpene biosynthesis.

**Figure 24.**
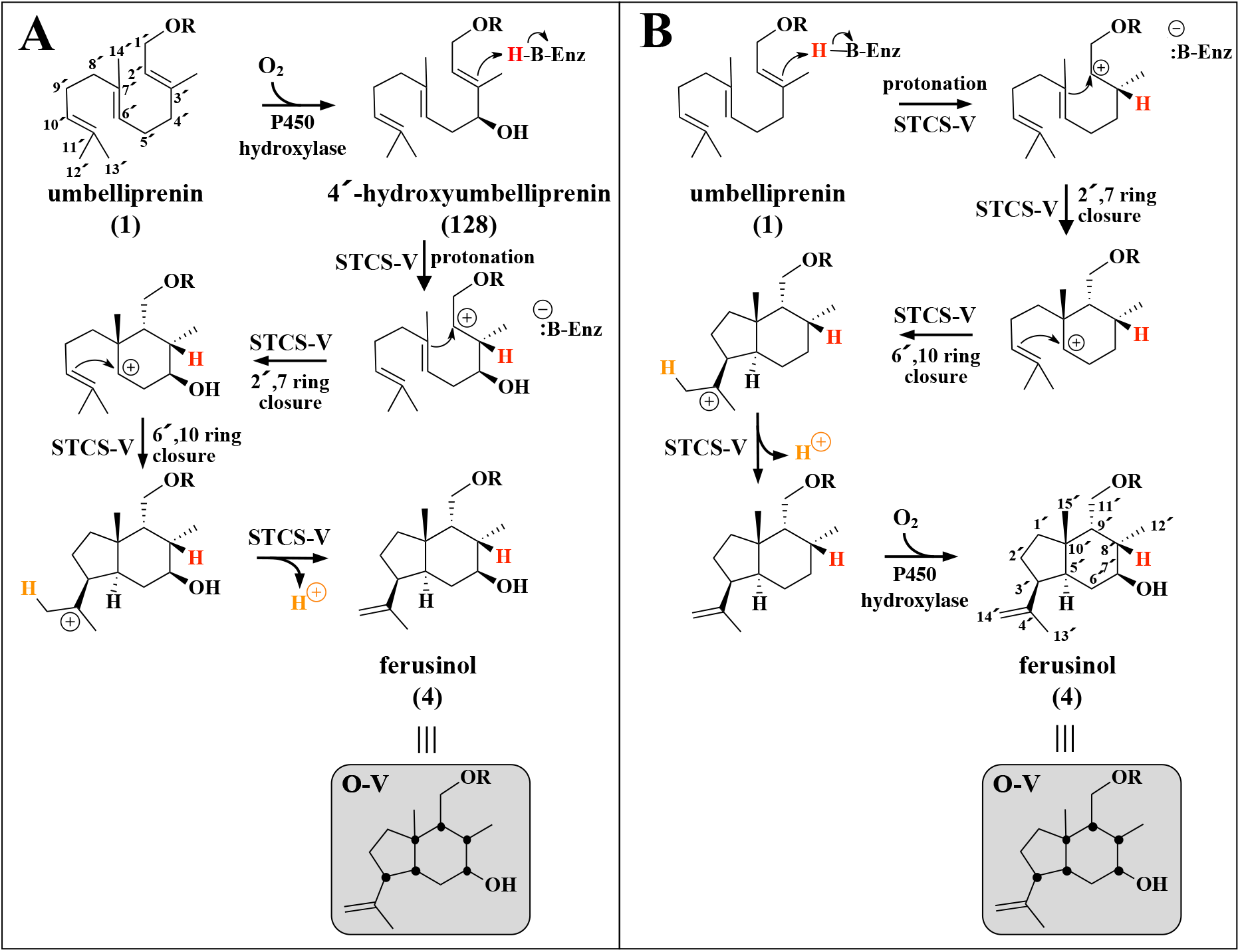
Putative biosynthesis of ferusinol (**4**) with carbon skeleton ***O*-V** from umbelliprenin (**1**). Chiral carbon atoms are indicated by black dots in the final products.

An alternative putative pathway leading to ferusinol (**4**) is shown in Figure 24B. In this case the starting substrate is umbelliprenin (**1**) (the general precursor of 7-farnesyloxycoumarins). The 2′,3′-double bond of umbelliprenin (**1**) is protonated, which is followed by the two ring closures to generate the 5,6-ring structure of ferusinol (**4**). After proton abstraction a P450 hydroxylase introduces the 7′-hydroxyl group to yield the final product.

The STCS catalyzing the formation of carbon skeleton ***O*-V** has most likely evolved from a SHC-like ancestral enzyme after gene duplication. As summarized in Table 2, it has been shown that SHCs can use many substrates (probably including umbelliprenin (**1**)) in addition to squalene (**20**). We suggest that ferusinol (**4**) is produced by a SHC-like STCS according to the pathway shown in Figure 24B. Umbelliprenin (**1**) is probably better accommodated in the active site of such an SHC-like enzyme than 4′-hydroxyumbelliprenin (**126**).

Bungeanin A (**129**), recently isolated from *Ferula bungeana*, is a sesquiterpene coumarin with a unique structure (Figure 25A) [Kang et al. 2024]. It is most likely produced from umbelliprenin (**1**) by formation of a C5′-C9′ bond. The three double bonds of umbelliprenin are conserved in the product indicating that no carbocations are involved in the formation of bungeanin A (**129**). We propose it is produced by a P450 enzyme catalyzing formation of the C5′-C9′ bond of the product as schematically shown in Figure 25A. Carbon-carbon bond formation in plant secondary metabolism has been shown to involve P450 enzymes [Tang et al, 2017; Guengerich & Yoshimoto, 2018]. The P450 enzyme generates two radicals by abstraction of two hydrogens. These two radicals form the new C5′-C9′ carbon-carbon bond.

**Figure 25.**
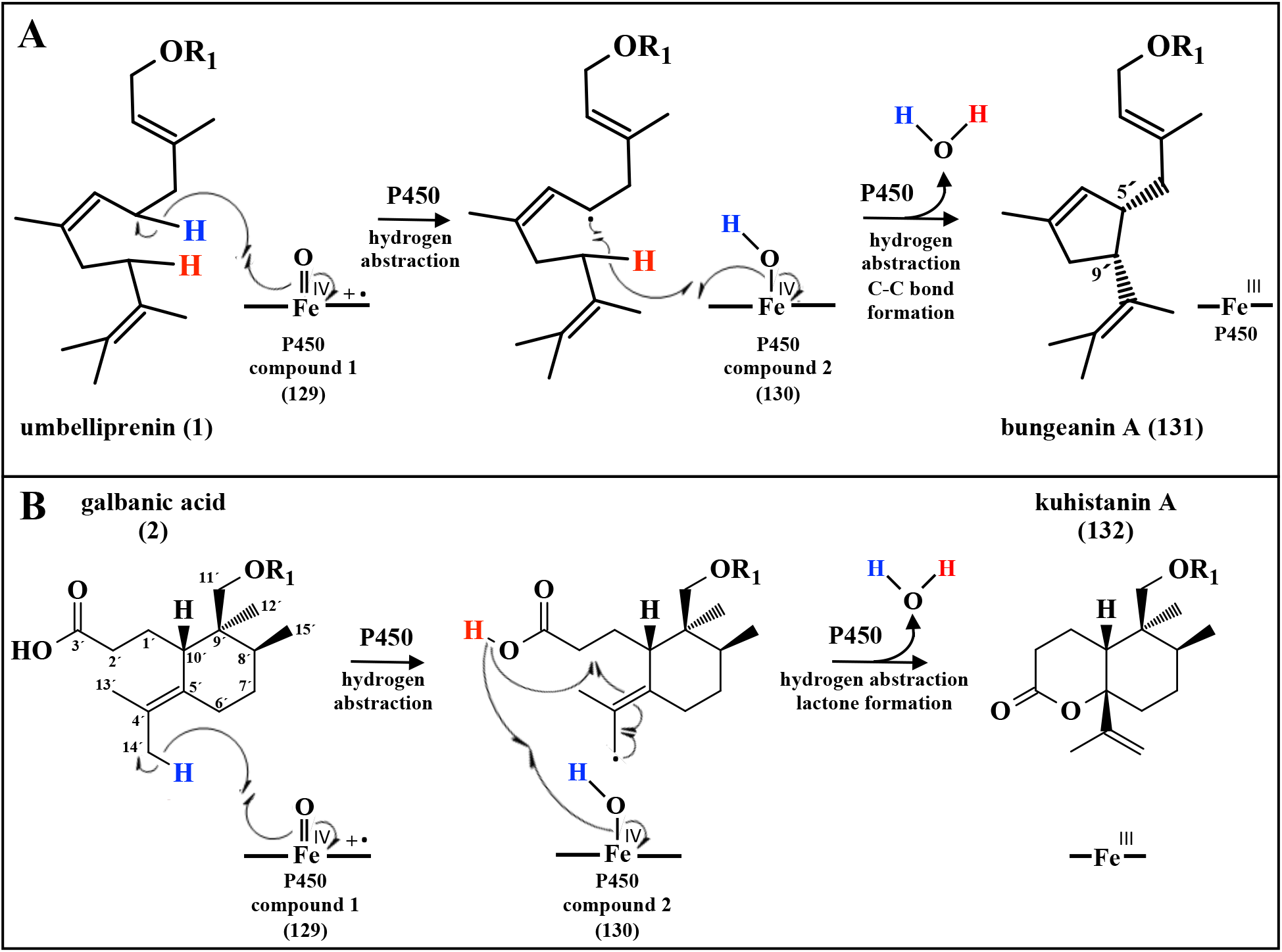
**A**. Putative biosynthesis of bungeanin A (**131**) from umbelliprenin (**1**) by a P450 enzyme forming a 5′,9′ C-C bond. **B**. Putative biosynthesis of kuhistanin A (**132**) from galbanic acid (**2**) by lactonization by a P450 enzyme.

### 4.10. Biotransformation of sesquiterpene coumarins

In a few cases, sesquiterpene coumarins are the starting substance for the biosynthesis of other sesquiterpene coumarins as shown in Figures 25 to 27. The lactone kuhistanin A (**130**), isolated from *Ferula kuhistanica*, exhibits the same stereochemistry as galbanic acid (**2**) (Figure 25B) indicating that it is derived from galbanic acid (**2**) [Ashurov et al. 2024]. Figure 25B shows the putative pathways for the conversion of galbanic acid (**2**) to kuhistanin A (**132**). The radical mechanism shown in Figure 25B starts by homolytic cleavage of the C14′-hydrogen bond of galbanic acid (**2**) by P450 compound 1 (**127**). A rearrangement results in the abstraction of the hydroxyl proton by P450 compound 2 (**128**) and formation of the lactone structure of kuhistanin A (**130**). An alternative pathway for the biosynthesis of kuhistanin A (**130**), based on classical terpene carbocation chemistry, is also possible. First the 4′,5′-double bond of galbanic acid (**2**) is epoxidized. The formed 4′,5′-oxidogalbanic acid is protonated and the carbocation formed is quenched by elimination of the C14′-hydrogen. The formed 5′-hydroxygalbanic acid, a hydroxycarboxylic acid, is spontaneously converted to the lactone kuhistanin A (**130**). 5′-Hydroxygalbanic acid has not been isolated from *Ferula kuhistanica* [Ashurov et al. 2024].

The product ligupersin B (**132**) was isolated from roots *Ligularia persica* (Asteraceae) along with twelve other sesquiterpene coumarins [Marco et al., 1991]. A potential precursor of ligupersin B (**132**) biosynthesis is kopeolin (***O*-U-Ea3**) (**131**), which has the same stereochemistry as the product. 12′-Hydroxylation of kopeolin (**131**) by a P450 followed by elimination of water will yield ligupersin B (**132**) with an C7′,C12′-ether bridge (Figure 26). This type of hydroxylation catalyzed by a cytochrome P450 enzyme with subsequent elimination of water to generate the ether bridge has been shown for other natural products [Rudolf et al, 2018].

**Figure 26.**
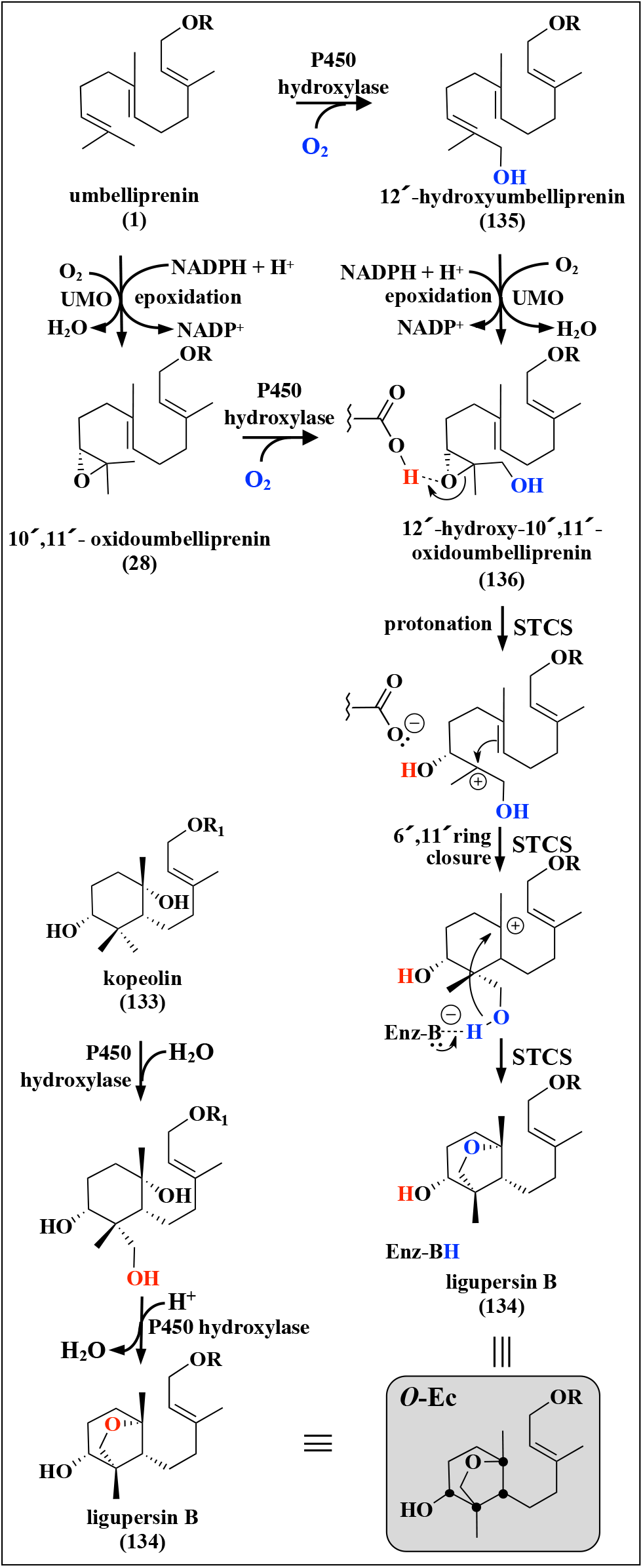
Putative biosynthesis of ligupersin B (**134**) with carbon skeleton ***O*-Ec** from kopeolin (**133**) or alternatively from umbelliprenin (**1**). Chiral carbon atoms are indicated by black dots in the final products.

An alternative biosynthesis of ligupersin B (**132**) starts from umbelliprenin (**1**). In this case, the initial step is a 12′-hydroxylation by a cytochrome P450 hydroxylase to yield 12′-hydroxyumbelliprenin (**133**). Epoxidation of 12′-hydroxyumbelliprenin (**133**) gives 12′-hydroxy-10′,11′-oxidoumbelliprenin (**134**). However, this could also be obtained by 12′-hydroxylation of 10′,11′-oxidoumbelli-prenin (**28**). Regardless, the epoxide is subsequently protonated and undergoes a 6′, 11′-cyclization, which results in a carbocation on C7′. The C7′, C12′-ether bridge of ligupersin B (**132**) is formed by quenching of the C7′-carbocation with the C12′ hydroxyl group in analogy to the formation of the C4′, C7′-ether bridge seen in structure ***O*-Cb** (shown in Figure 7A). This second pathway for the formation of ligupersin B (**132**) may be less plausible because binding of the substrate to the STCS may be prevented due to steric hindrance imposed by the 12′-hydroxy group.

Putative pathways leading to carbon skeletons ***O*-W** (karatavic acid (**135**)) and ***O*-X** (ferusingensine H (**136**)) are shown in Figure 27A and B, respectively. The pathway leading to karatavic acid (**135**) starts from structure ***O*-U-Ha24** (**124**) (Figure S4). The structure of karatavic acid (**135**) is similar to that of galbanic acid (**2**) being almost a positional isomer. We suggest that a cytochrome P450 enzyme in state I (compound I) (**127**) abstracts a hydrogen atom from the 12′-methyl group. The resulting carbon radical serves as an electron donor giving rise to a carbocation and the P450 enzyme in state II (compound II) for hydroxylation of the substrate, is abstracted from the (**128**). Subsequently, the hydroxyl, which normally is used enzyme. This mechanism has been suggested for a few reactions involving carbocations on various natural products [Zhang & Li, 2017]. In the next step, the same cytochrome P450 enzyme converts the carbocation intermediate into the 3,4 seco-drimane structure by a Grob fragmentation. In the final step, the aldehyde is enzymatically oxidized by a NAD(P)^+^-dependent reductase to karatavic acid (**135**) with carbon skeleton ***O*-W**.

**Figure 27.**
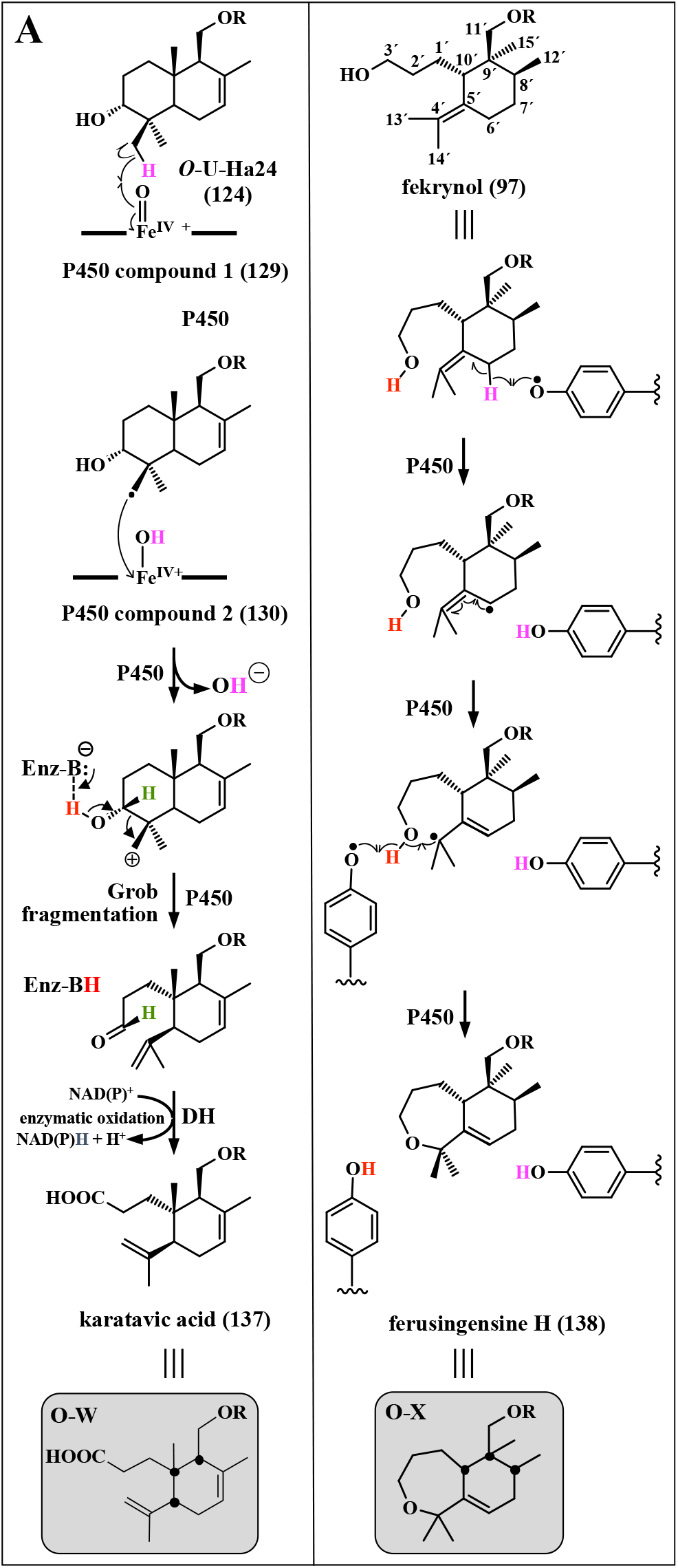
Putative biosynthesis of karatavic acid (**137**) and ferusingensine H (**138**) with carbon skeletons ***O*-W** and ***O*-X** from ***O*-U-Ha24** (**124**) and fekrynol (**97**), respectively. Chiral carbon atoms are indicated by black dots in the final products.

We suggest that a radical-based reaction mechanism is involved in the biosynthesis of structure ***O*-X** (ferusingensine H (**136**)) from the 3,4 seco-drimane structure ***O*-U-Rc** (fekrynol (**97**)) (Figure 18). The reaction shown in Figure 27B is initiated by homolytic cleavage of the terminal hydroxyl group, which results in a radical on C6′. A rearrangement generates a C5′,C6′double bond and a radical on C4′. After homolytic cleavage of the C3′-hydroxyl group a radical on the oxygen is formed, which combines with the radical on C4′to form the ether bridge and ring closure.

#### 4.1.1. Sesquiterpene coumarins from 4-*O*-farnesyloxy-5-methylcoumarin

Five sesquiterpene coumarins obtained from 4-*O*-farnesyloxy-5-methylcoumarin (**137**) have been isolated from *Nassauvia revoluta* [Bittner et al, 1988a] and *Nassauvia argentea* (Asteracea) [Bittner et al, 1988b], plants native to Argentina, Bolivia, Chile, and the Falkland Islands. The biosynthesis of the sesquiterpene moieties is the same as for those of 7-O-farnesylated sesquiterpene coumarins (Figure 7). First, 4-(10′,11′-oxidofarnesyloxy)-5-methylcoumarin (**138**) is formed, which is subsequently protonated to give a 10′-hydroxyl group and a carbocation on the 11′-C. From there, different routes lead to the biosynthesis of the three sesquiterpene coumarins nassuvirevoltin A (**139**), nassuvirevoltin C (**140**) and 7′,14′-dehydro-7′,10′-oxidonassauvirevolutin A (**141**) based on 4-hydroxy-5-methylcoumarin (**18**), as shown in Figure 28. The carbon skeleton of these sesquiterpene coumarins belong to ***O-*Cb** (**141**), ***O-*D** (**139**) and ***O-*Ia** (**140**).

**Figure 28.**
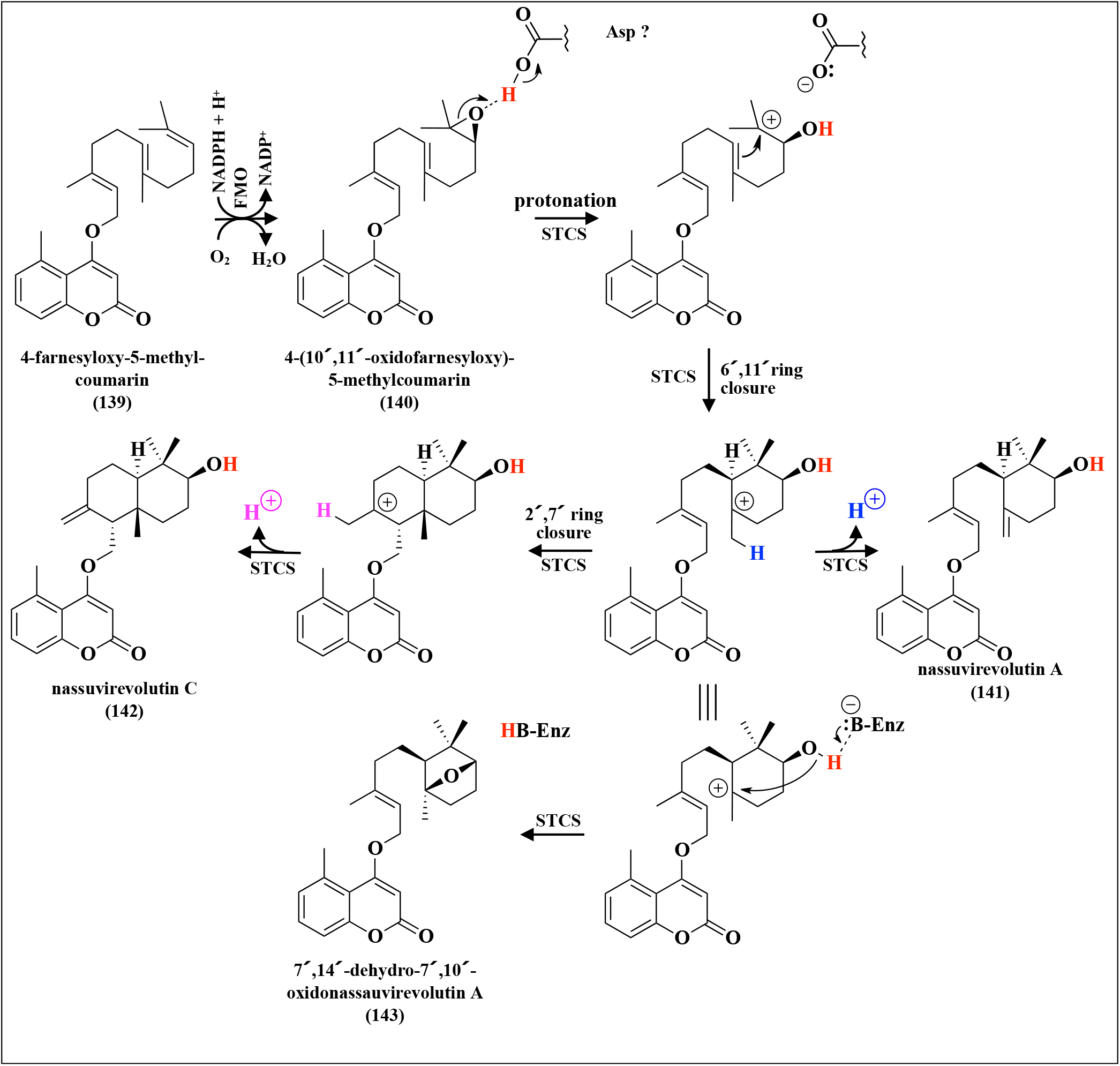
Putative biosynthesis of various sesquiterpene coumarins from 4-farnesyloxy-5-methylcoumarin (**139**).

#### 4.1.2. Other *O*-farnesylated sesquiterpene coumarins

Fraxetin (**11**) (Figure 2) is very rarely found as a building block of sesquiterpene coumarins. An example is 8-farnesyloxyfraxetin (**142**) (***O*-F-A1**) (Figure 29A), which was isolated from *Brochia cinerea* (Asteraceae) [Greger & Hofer, 1985]. This linear sesquiterpene coumarin is obtained by farnesylation of fraxetin (**11**) by a *O*-farnesyltransferase. It is interesting to note that nearly all the sesquiterpene coumarins described in this text have been isolated from plants belonging to the Apiaceae and Asteraceae families, except for a few examples (shown in Figure 29B to E). The linear sesquiterpene coumarin umbelliprenin (**1**) (***O*-U-A1**) (Figure 29B) is obtained by farnesylation of umbelliferone (**9**) (Figure 4) by a *O*-farnesyltransferase. Umbelliferone (**9**) is widely distributed within the Rutaceae and Apiaceae families [Dawidowicz et al, 2018] but it occurs in the flowers, fruits, and roots of all higher plants [Bourgaud et al, 2006]. Umbelliprenin (**1**) has been isolated from plants belonging to Amaranthaceae (*Amaranthus retroflexus, Chenopodium quinoa*, and *Spinacia oleraceae*) [Fiorito et al, 2017; Fiorito et al, 2019b], Caprifoliaceae (*Scabiosa comosa*) [Dargaeva & Brutko, 1976], Lamiaceae (*Scutellaria baicalensis)* [Murch et al, 2004], Lythraceae (*Punica granatum*) [Fiorito et al, 2019a], Myrtaceae (*Melaleuca alternifolia*) [Scotti et al, 2018], Rutaceae (*Citrus limon* and *Haplophyllum patavinum*) [Sidana et al, 2013; Filippini *et al*., 1998] and Solanaceae (*Lycium barbarum*) [Fiorito et al, 2019b] families (Figure 29B; Table 1; Table S1). 10′R-Karatavicinol (**35**) was isolated from *Scutellaria baicalensis* in addition to umbelliprenin (**1**) indicating that epoxidation of umbelliprenin (**1**) takes place in this species [Murch et al, 2004].

**Figure 29.**
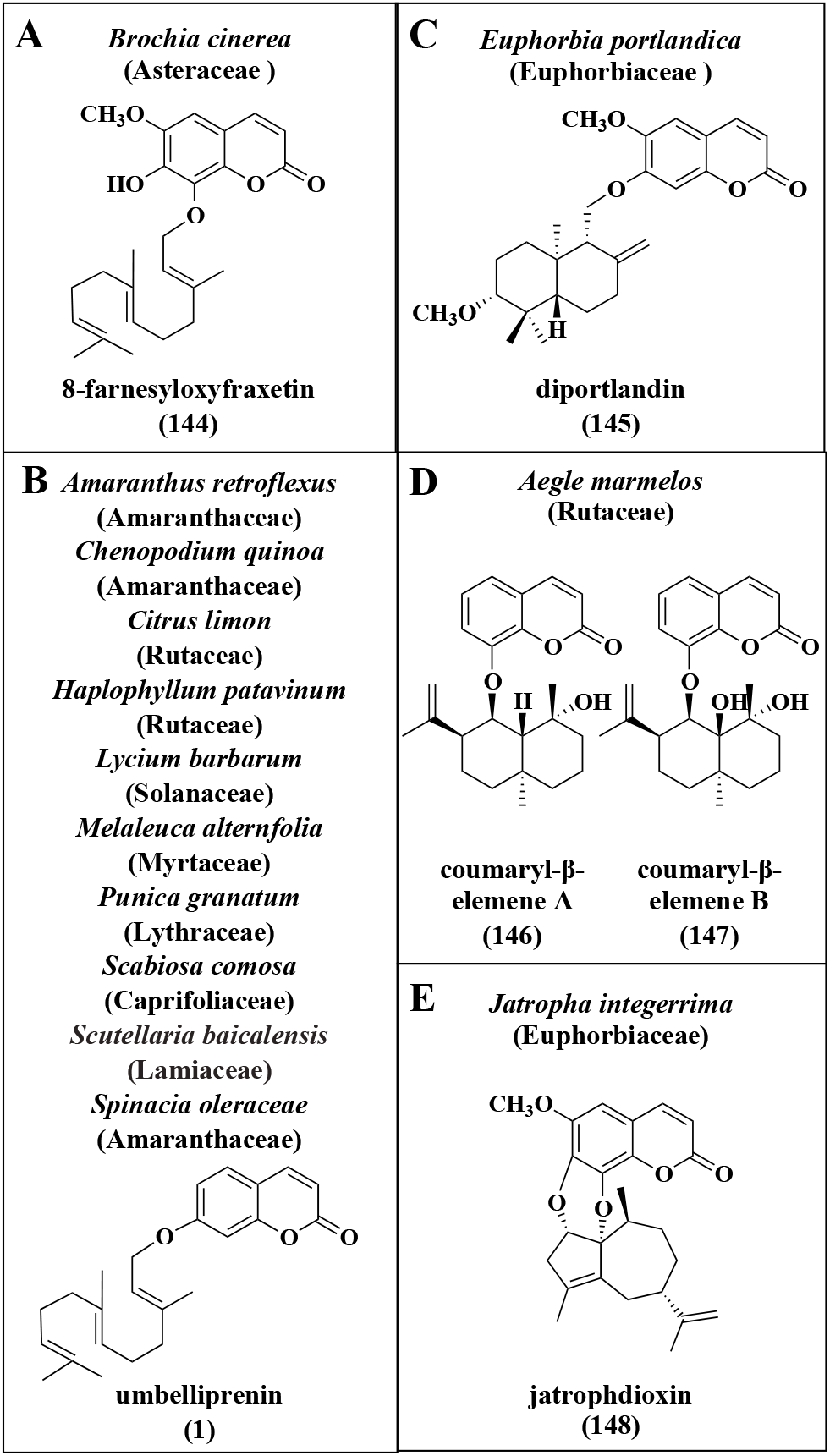
A: 8-Farnesyloxyfraxetin (**148**) from *Brochia cinerea* (Asteraceae); B to D: Sesquiterpene coumarins isolated from plants not belonging to the Apiaceae or Asteraceae families (**145-148**).

The other three compounds have been isolated from the Euphorbiaceae and Rutaceae families (Figure 29C to 30E). They are not derived from 10′,11′-oxidofarnesyloxy-coumarins except for diportlandin (**143**) (***O*-S-Ia1**) from *Euphorbia portlandica*, which carries a methoxy group on C10′. This indicates that it may have been formed from 10′,11′-oxidoscopofarnol according to the biosynthesis shown in Figure 7 even though it has been isolated from a plant belonging to Euphorbiaceae.

8-Hydroxycoumarin (**13**) is the coumarin moiety of two sesquiterpene coumarins, coumaryl-Δ-elemene A (**144**) and coumaryl-β-elemene B (**145**), with carbon skeleton ***O-*Y** (Figure 29D) isolated from roots of *Aegle marmelos* [Ahmad et al, 2012]. The suggested biosynthesis, shown in Figure 30, involves a sesquiterpene synthase (STS) that first produces an intermediate (+)-germacrene A (**148**), which remains bound to the active site of the enzyme. The reaction is reinitiated by protonation of the 1 ’,10 ’-double bond to generate a germacryl carbocation (**147**). This suggested reaction is based on the reported reaction mechanism of aristolochene synthase from *Penicillium roqueforti* [Calvert et al, 2002] and 5-epi-aristolochene synthase from tobacco [Rising et al, 2000]. The generated germacryl carbocation (**147**) is converted to an eudesmyl carbocation (**149**) through a 5 ’,10 ’-ring closure. In aristolochene synthase and 5-*epi*-aristolochene synthase, the final products are obtained after 1,2-hydride and 1,2-alkyl shifts followed by abstraction of a proton, as indicated in Figure 30. In the reaction of the *A. marmelos* enzyme, abstraction of a proton from the eudesmyl carbocation (**149**) generates the sesquiterpene eudesma-4,11-diene (**150**), which in the next step is epoxidized to 4′,5′-oxidoeudesma-11-ene (**151**) by a monooxygenase. Protonation of the epoxide generates a carbocation on C5 ’ and a hydroxyl group on C4′. After a 1,2-hydride shift a carbocation on C6′ is obtained. This intermediate cation reacts with the 8-hydroxy group of the coumarin to form the ether bridge of the final product with carbon skeleton ***O*-Y** (Figure S9).

**Figure 30.**
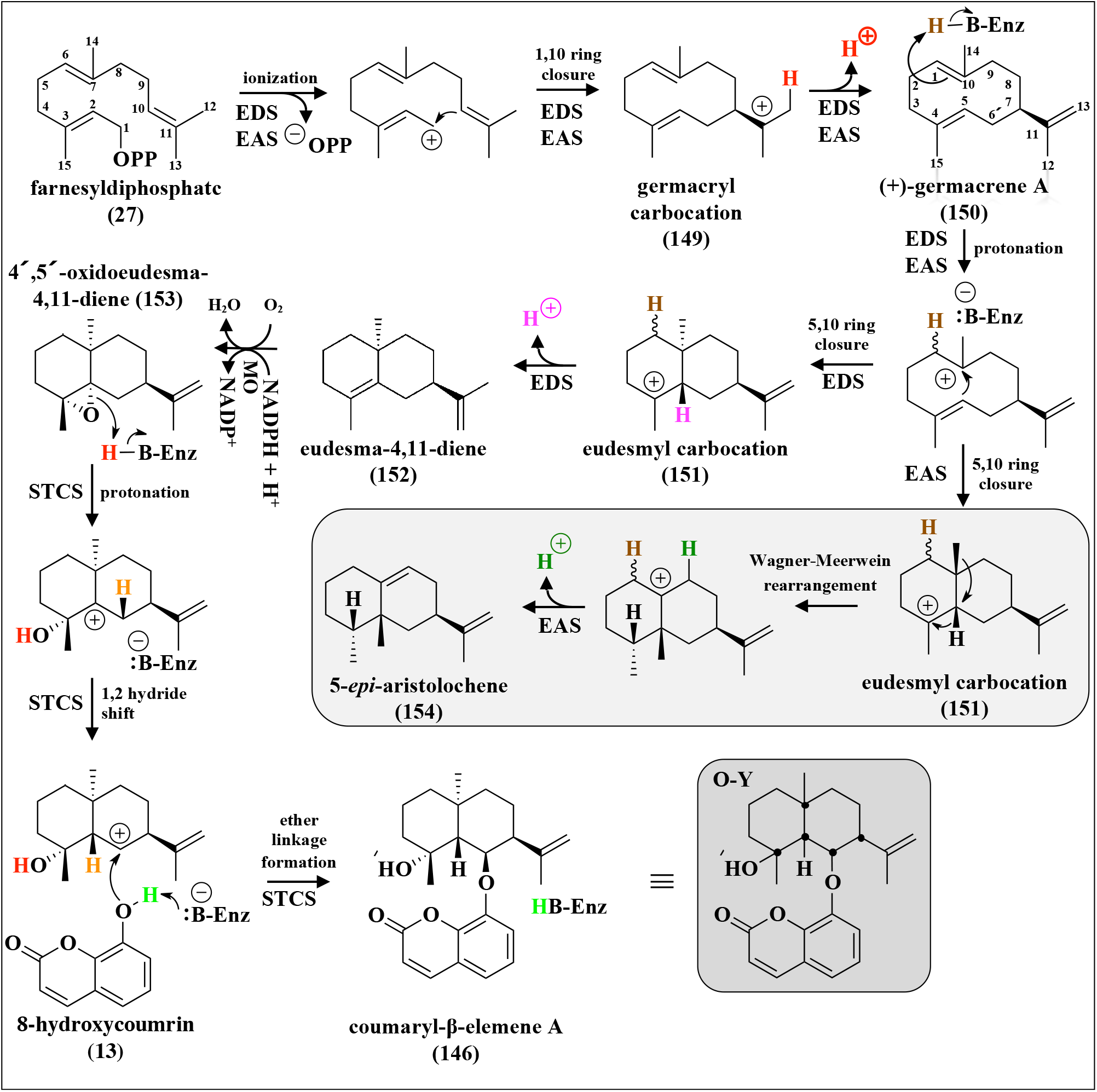
Putative biosynthesis of coumaryl-β-elemene A (**146**) with carbon skeleton ***O*-Y**. Chiral carbon atoms are indicated by black dots in the final products

Jatrophadioxin (**146**) (***O*-U-Z1** Figure S9) (Figure 29E), isolated from *Jatropha intergerrima* (Euphorbiaceae) [Sutthivaiyakit et al, 2009], is a second example of a sesquiterpene coumarin with fraxetin (**11**) as coumarin moiety. In this case, two ether bridges link the sesquiterpene moiety to the coumarin. It is suggested that thesesquiterpene moiety 7-*epi*-aciphyllene (**154**), which is of guaiene type, is formed before it is coupled to fraxetin (**11**) as shown in Figure 31. (-)-Germacrene A (**153**) is also an enzyme boundintermediate in this biosynthesis but here the C4,C5-double bond of (-)-germacrene A (**153**) is reprotonated followed by a 1,5-ring closure. Abstraction of a proton gives the sesquiterpene 7-*epi*-aciphyllene (**154**) (guaia-4,11-diene). We suggest that the coupling of fraxetin (**11**) and 7-*epi*-aciphyllene (**154**) is catalyzed by a cytochrome P450 monooxygenase in a manner similar to that reported for the formation of ether bridges in secondary metabolites [Rudolf et al, 2018]. The P450 enzyme hydroxylates the sesquiterpene and forms an ether linkage by elimination of water. This reaction is repeated, possibly by the same P450 enzyme, to form the second ether bridge. The order of the last reactions can be reverse.

**Figure 31.**
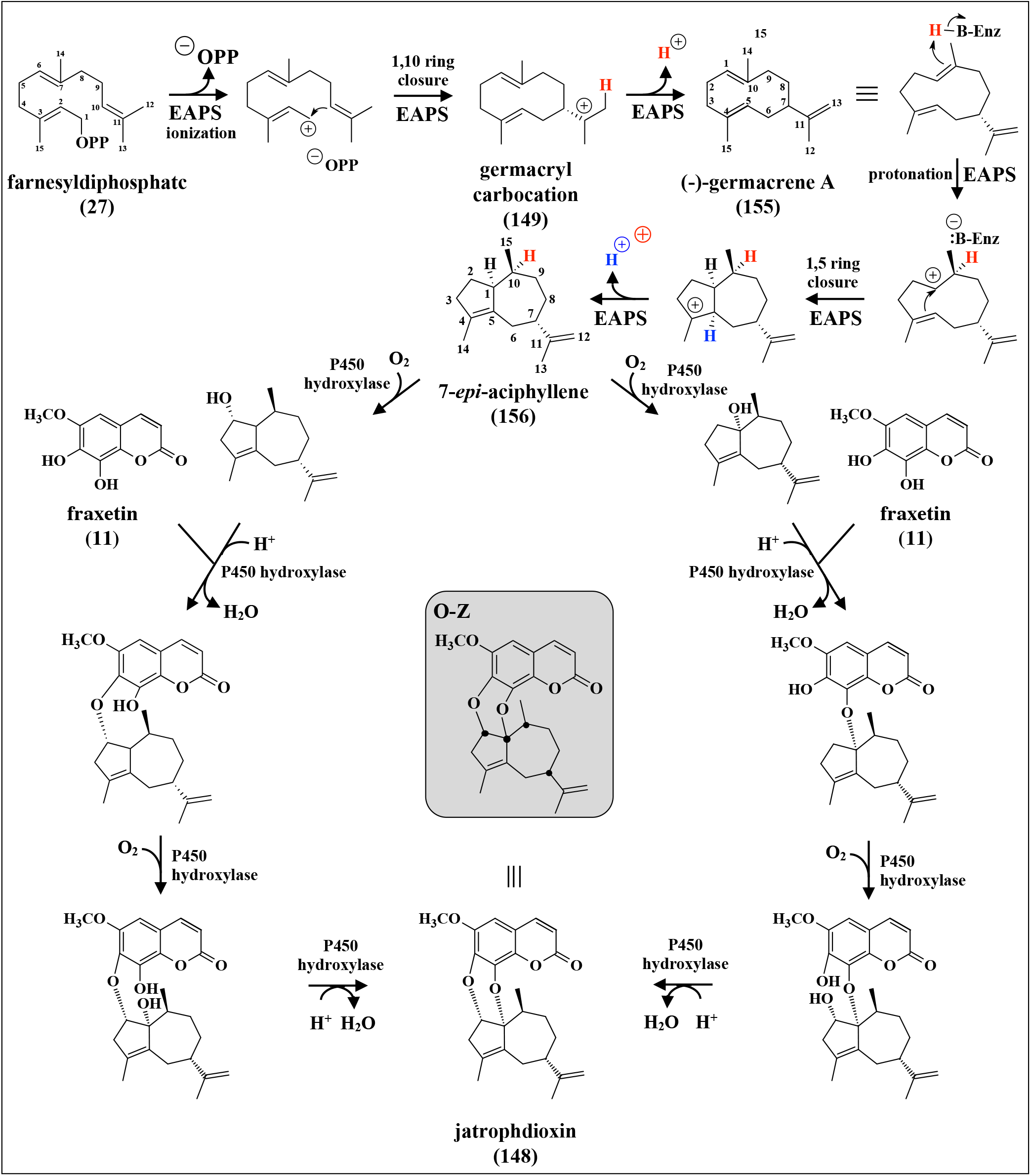
Putative biosynthesis of jatrophdioxin (**148**) with carbon skeleton ***O*-Z**. Chiral carbon atoms are indicated by black dots in the final product.

This is in line with the above suggestion (discussed in section 4.2) that the sesquiterpene coumarin synthase genes coding for synthases using 7-(10′,11′-oxidofarnesyloxy)-coumarin as a substrate, have evolved from ancestor triterpene synthase genes in Apiaceae and Asteraceae but not in other plant families

#### 4.1.3. A norsesquiterpene coumarin

The norsesquiterpene coumarin tavicone (**156**) (**O-U-α1;** Figure S9) has been isolated from *Ferula karatavica* [Bagirov et al, 1969]. The structure of tavicone (**156**) was later corrected [Bagirov & Sheichenko, 1976]. The terpene moiety of tavicone (**156**) is a norsesquiterpene with 14 carbon atoms (as opposed to the normal 15 carbons of sesquiterpenes), which has similarities to drimenol (**59**). This is the only norsesquiterpene coumarin that has been reported. During the biosynthesis of tavicone (**156**) an intermediate sesquiterpene coumarin (**155**) is formed, which is demethylated in later steps of the pathway. A putative biosynthetic route from the monocyclic carbocation (**29**) to tavicone (**156**) with carbon skeleton ***O*-α** is shown in Figure 32. The putative pathway can be divided into three sets of reactions. The first set (box A) is the formation of an intermediate sesquiterpene coumarin (**155**) by a STCS. The next set of reactions (box B) is catalyzed by a methyl oxidase (MO) that introduces a hydroxyl group and further oxidizes it to a carboxyl group via the corresponding aldehyde. Introduction of this carboxyl group, vicinal to the A-ring alcohol, provides a handle for the subsequent decarboxylation that ultimately affords the norsesquiterpene skeleton. This suggestion is based on the demethylation of cycloartenol to norcycloartenol in sterol biosynthesis in plants [Rahier, 2011; Pascal et al, 1993]. MO is a membrane-bound non-heme iron oxygenase, which uses O_2_ and ferrocytochrome b5 (Fe^2+^). The ferrocytochrome b5 (Fe^2+^) is oxidized to ferricytochrome b5 (Fe^3+^) during theformation of the carboxy-sesquiterpene coumarin metabolite. The third set of reactions (box C) is catalyzed by a NADP^+^-dependent decarboxylating dehydrogenase, which produces the final product by deprotonation and release of CO_2_. This reaction sequence is based on a model proposed for a membrane bound (ER) multienzyme complex in plants that is involved in sterol biosynthesis. It consists of sterol methyl oxidase (SMO), a non-heme iron monooxygenase; 3β-hydroxysteroid dehydrogenases/C4 decarboxylase (HSD/D), a member of the short-chain dehydrogenases/ reductases family; 3-oxosteroid reductase (SR), a small transmembrane protein (supposed to tether SMO, HSD/D and SR to the ER); and cytochrome b5 and NADH cytochrome b5 reductase [Rahier, 2011]. The reductase, which reduces the keto group is not required for tavicone (**156**) biosynthesis. However, it is likely that a similar membrane bound multienzyme complex is involved in the demethylation of the intermediate sesquiterpene coumarin in tavicone (**156**) biosynthesis.

**Figure 32.**
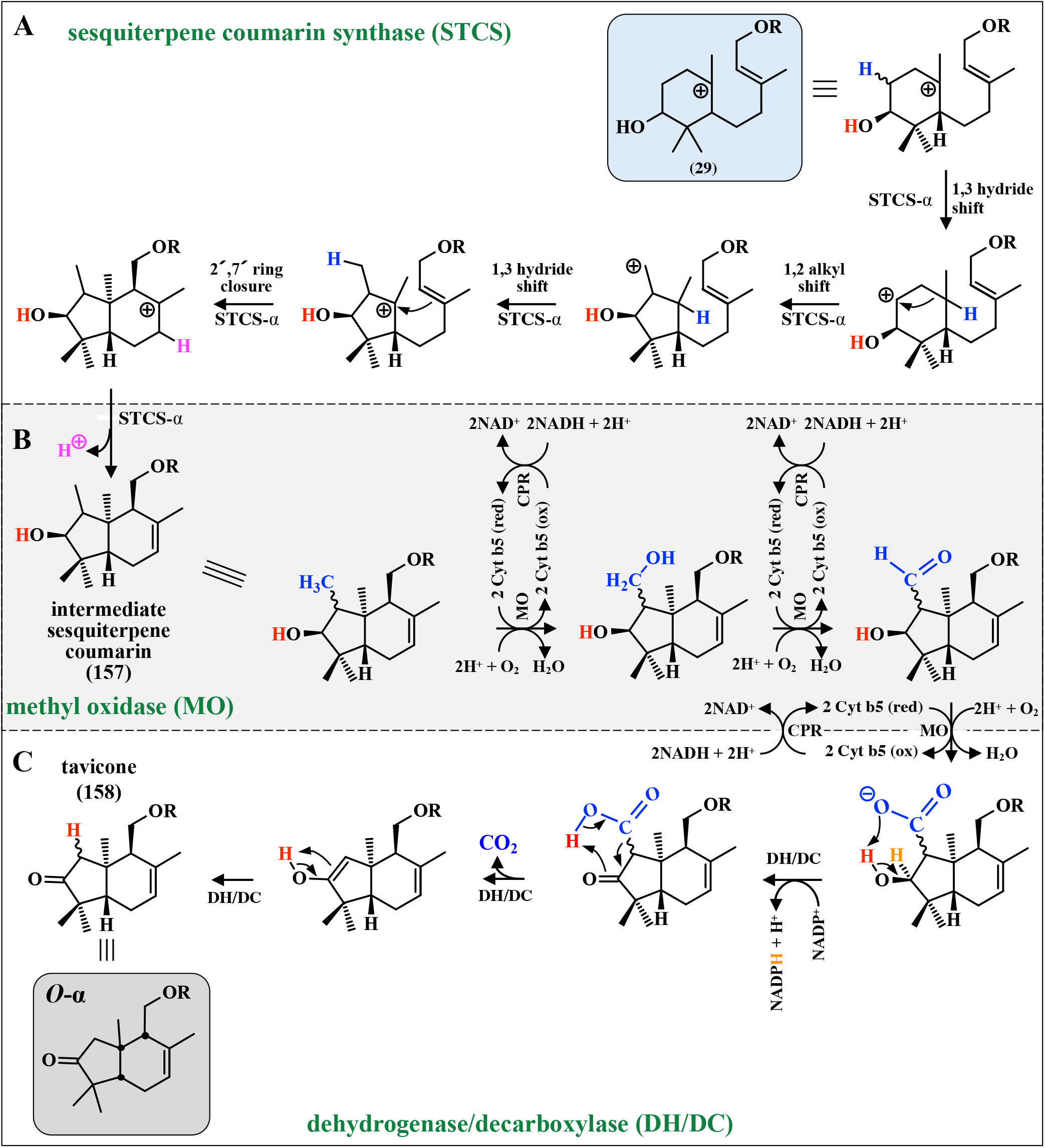
Putative biosynthesis of the norsesquiterpene coumarin tavicone (**158**) from intermediate carbocation I (**29**). STCS: sesquiterpene coumarin synthase; MO: methyl oxidase; DH: dehydrogenase; DC: decarboxylase. Chiral carbon atoms are indicated by black dots in the final structure.

#### 4.1.4. Diversity of sesquiterpene coumarins

The enormous diversity of sesquiterpenes is reflected in the sesquiterpene coumarins. The cyclization of farnesyloxycoumarins gives rise to a broad spectrum of carbon skeletons as discussed above. In this diversification, Wagner–Meerwein rearrangements and Grob fragmentations play important roles. The chemical space of each constitutionally identical carbon skeleton is further expanded due to the range of possible diastereomers. As an example, the bicyclic sesquiterpene moiety of the sesquiterpene coumarins shown in Figure 33 contains five asymmetric centers. The three marked with black dots in Figure 33 can adopt either *R*- or *S*-configuration. The other

**Figure 33.**
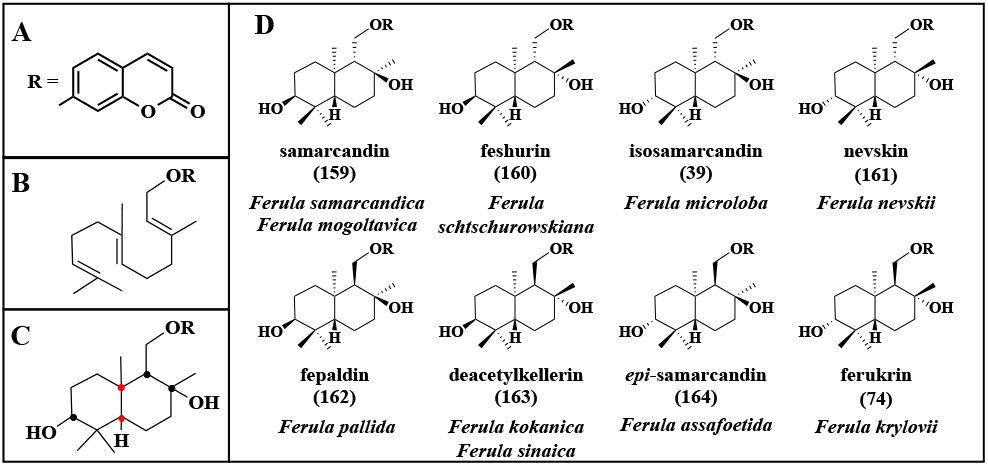
Eight diastereomers of a bicyclic sesquiterpene coumarin (***O*-U-Ja**; Figure S8). Three chiral centers (black dots) can adopt *R*- or *S*-configurations and two centers (red dots) are unchanged.

#### 4.14. Diversity of sesquiterpene coumarins

The enormous diversity of sesquiterpenes is reflected in the sesquiterpene coumarins. The cyclization of farnesyloxycoumarins gives rise to a broad spectrum of carbon skeletons as discussed above. In this diversification, Wagner–Meerwein rearrangements and Grob fragmentations play important roles. The chemical space of each constitutionally identical carbon skeleton is further expanded due to the range of possible diastereomers. As an example, the bicyclic sesquiterpene moiety of the sesquiterpene coumarins shown in Figure 33 contains five asymmetric centers. The three marked with black dots in Figure 33 can adopt either *R*- or *S*-configuration. The other two marked with red dots remain unchanged in this example. Consequently eight (*i.e*., 2^3^) diastereomers can be formed. The structures of the eight possible natural diastereomers are shown in Figure 33. All eight diastereomers have been isolated and their absolute configurations have been confirmed [Tashkhodzhaev et al, 2015]. Obviously, sesquiterpene coumarin synthases from different closely related plant species have evolved subtle differences in their active site directing the stereochemical arrangement of the sesquiterpene moiety.

## 5. Concluding remarks

In this review we have suggested putative biosynthetic routes for sesquiterpene coumarins according to a general scheme starting with the substrate farnesyl diphosphate (**27**) or nerolidyl diphosphate and a coumarin. The first step is the farnesylation/nerolydation of a hydroxycoumarin by a *O*-prenyltransferase to yield farnesyloxy/nerolidyloxy coumarins. The second step is epoxidation of the farnesyloxy/nerolidyloxy coumarins to introduce a 10′,11′-epoxy group. In the next step, sesquiterpene coumarin synthases, which are type II terpene synthases, initiate the cyclization reactions by protonation of the epoxy group resulting in the formation of an intermediate carbocation with a C10′-hydroxyl group. This carbocation initiates cyclization reactions similar to those catalyzed by terpene synthases [Christianson 2017]. Based on our knowledge of terpene synthases we have suggested reaction mechanisms for the various sesquiterpene coumarins. Some of these reactions also involve rearrangements of the carbon skeleton by Wagner-Meerwein-rearrangements and Grob fragmentation reactions.

To our knowledge, there is no report on the isolation or cloning of a STCS. The reaction mechanism of such an enzyme is most likely similar to other terpene synthases, which initiate cyclization by protonation of an epoxy-group, such as lanosterol and cycloartenol cyclase [Thimmappa et al, 2014, Wang et al, 2022a]. These OSCs are localized to lipid particles in which the highly hydrophobic substrate and products can be stored and transported. The aspartic residue of the conserved sequence DCTAE has been shown to be involved in the initial protonation of the epoxide [Qiao et al, 2018]. The motif QxxxxxW, which is important for stabilization of the carbocation during cyclization, is repeated three times. Finally, a histidine residue of the conserved motif MWCHCR is assumed to stabilize the protosteryl cation (**23**). It is likely that these conserved sequences can also be found in STCSs.

Cell cultures of sesquiterpene coumarin-producing plants offer potential to serve as powerful tools for studying the biosynthesis of sesquiterpene coumarins. Biotic elicitors have been widely used to induce enzymes of biosynthesis and increase the production of secondary metabolites in hairy root cultures [Alcalde et al, 2022]. *In vitro* induction of the enzymes involved in the biosynthesis of the sesquiterpene coumarins (such as *O*-farnesyltransferase, farnesyloxy-coumarin epoxidase and/or sesquiterpene coumarin synthase) through elicitation could simplify their identification and subsequent cloning. Some *in vitro* cell culture systems of sesquiterpene coumarin-producing species have been reported (Table 4). One example has already been discussed above, i.e., induction of umbelliprenin (**1**) formation in callus cultures of *Ferulago campestris* (Apiaceae) after elicitation with ferulic acid [Fiorito et al, 2022]. This platform could be excellent for studying the formation of umbelliprenin (**1**) from FDP (**27**) and umbelliferone (**9**) catalyzed by *O*-farnesyltransferase. The induction of hairy root cultures of *Ferula pseudalliacea* (Apiaceae) has been reported [Khazaei et al, 2019]. Farnesiferol B (**5**) (***O*-U-Da1**; Figure S3) was extracted at a yield of 12 and 7.5 mg/g dry weight for the wildtype and hairy roots, respectively. This culture will be useful for studies on the biosynthesis of farnesiferol B (**5**) from ubelliferone (**9**) and farnesyldiphosphate (**27**) as outlined in Figure 4B. The three enzymes *O*-farnesyltransferase (*O-* FT), umbelliprenin epoxidase (UMO) and sesquiterpene coumarin synthase (STCS), which are involved in the biosynthesis of most sesquiterpene coumarins, may be cloned and characterized using the *F. pseudalliacea* hairy root culture. Hairy root cultures of both *Tanacetum parthenium* (Asteraceae) and *Artemisia annua* (Asteraceae) have also been reported to produce sesquiterpene coumarins. The *T. parthenium* cultures produced isofraxidine (**12**) and the sesquiterpene coumarin 9-*epi*-pectachol B (**O-I-Ia9**; Figure S5) [Kisiel & Stojakowska, 1997]. Isofraxidine (**12**) is the coumarin moiety of 9-*epi*-pectachol B. The co-production of isofraxidine (**12**) and 9-*epi*-pectachol B indicates that the formation of the two products is coordinated. The sesquiterpene coumarin drimartol A (***O*-I-Ha1**; Figure S4) was isolated from the *A. annua* culture at a yield of 20 mg/g dry weight [Zhai & Zhong, 2010]. In addition, callus cultures of the sesquiterpene coumarin-producing species *Ferula assa-foetida, Ferula ferulaeoides, and Ferula gummosa* (Apiaceae) have been described. Although, no investigations of the production of sesquiterpene coumarins were reported [Zare et al, 2010; Suran et al, 2016; Hadi et al, 2011] the conditions for the establishment of these undifferentiated cell cultures may be of interest.

**Table 4.** Examples of sesquiterpene coumarins produced in callus or hairy root cultures.

| Species | Type of culture | Product | Yield | Reference |
| --- | --- | --- | --- | --- |
| *Ferula assa-foetida* | callus | - | - | Zare et al, 2010 |
| *Ferula ferulaeoides* | callus | - | - | Suran et al, 2016 |
| *Ferula gummosa* | callus | - | - | Hadi et al, 2011 |
| *Ferulago campestris* | callus | umbelliprenin (**O-U-A1**) | 2.6 mg/g DW* | Fiorito et al, 2022 |
| *Ferula pseudalliacea* | hairy root | farnesiferol B (**O-U-D1**) | 7.5 mg/g DW | Khazaei et al, 2019 |
| *Tanacetum parthenium* | hairy root | 9-*epi*-pectachol B (**O-I-Ia9**) | 0.14 mg/g DW | Kisiel & Stojakowska, 1979 |
| *Artemisia annua* | hairy root | drimartol A (**O-I-Ha1**) | 20 mg/g DW | Zhai & Zhong, 2010 |

\*after elicitation with 10 μM ferulic acid

An interesting experiment was reported by Zhou et al [2016a]. In this study, the bicyclic sesquiterpene coumarin kellerin (**163**) (***O*-U-Ja34**, Figure S6) was fed to a cell culture of *Angelica sinensis* (Apiaceae). As shown in Table S1, plants belonging to the genus *Angelica* produce the linear sesquiterpene coumarin umbelliprenin (**1**) but no cyclic sesquiterpene coumarins. Biotransformation of kellerin by various enzyme systems in the cultivated *Angelica* cells resulted in the formation of eight sesquiterpene coumarins as shown in Figure 34. Five of these (**164**-**168**) have not been isolated before. It may be concluded that the diversity of sesquiterpene coumarins may be increased by feeding sesquiterpene coumarins to plant cell cultures and possibly cell cultures of fungi and bacteria.

**Figure 34.**
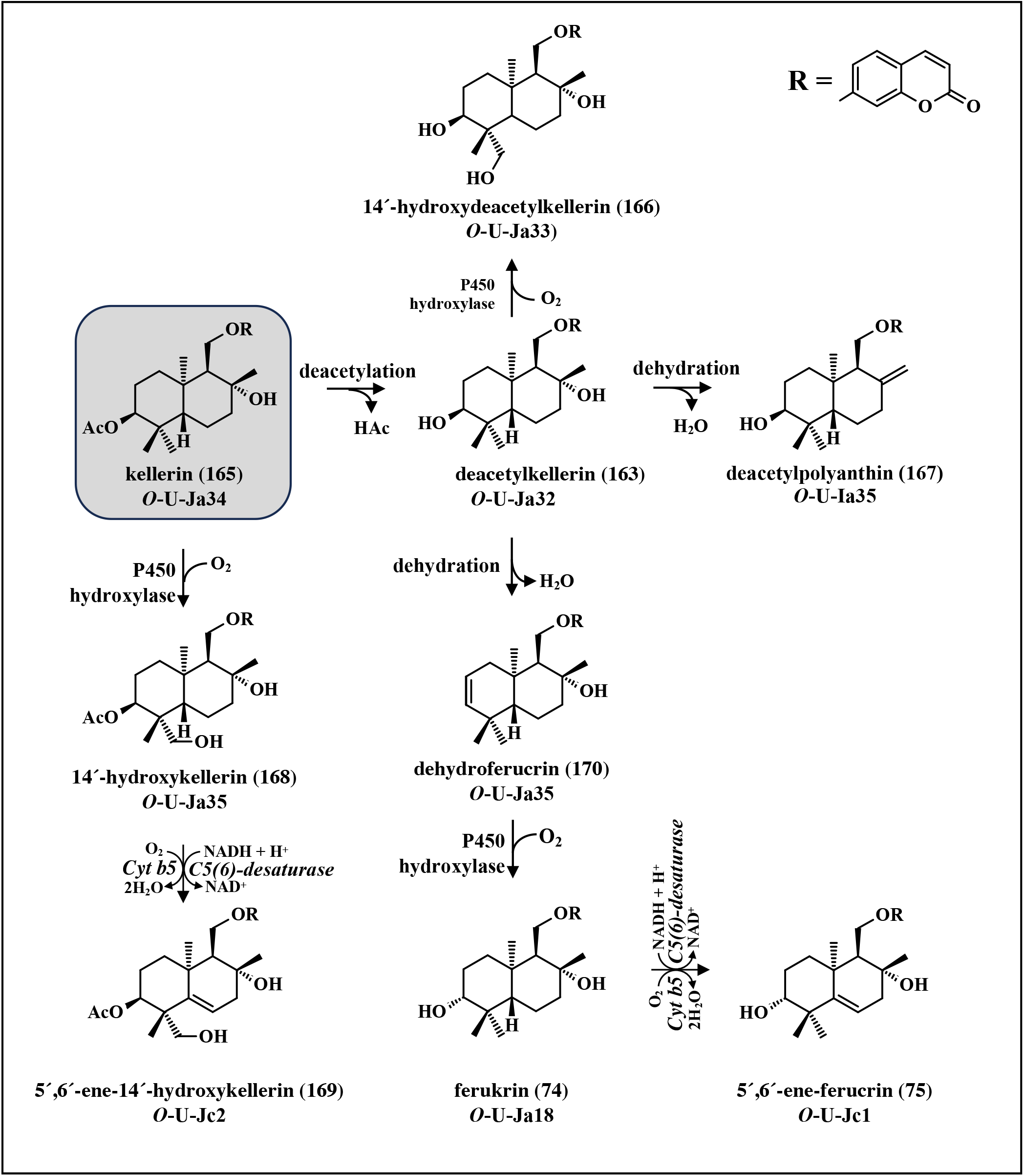
Putative reaction pathways to various metabolism after feeding kellerin to callus culture of *Angelica sinensis*. Adopted from Zhou et al, 2016.

10′,11′-Oxidoumbelliprenin (**28**) (***O*-U-A2**) is the substrate for many STCSs from various plants. The availability of this substrate will make it possible to screen for putative STCSs in extracts from plants tissues producing sesquiterpene coumarins. It is likely that these enzymes only require the appropriate substrate to exhibit activity. Furthermore, the substrate is important for the functional characterization of putative STCS genes, which will be cloned as more plant transcriptomes and genomes become available. A convenient method to synthesize 10′,11′-oxidoumbelliprenin (**28**) in 60-65 % yield has been described by Cravotto et al [2004].

An analysis of the transcriptome of *Ferula gummosa* (Apiaceae) has been reported [Najafabadi et al, 2017]. All the enzymes of the MVA and MEP pathways were expressed, as well as, some prenylsynthases such as geranyl- and farnesyl synthases. The expression of some mono-sesqui- and triterpene synthases were also confirmed. However, no enzyme specifically involved in the biosynthesis of sesquiterpene coumarins was identified.

In a recent study, comparative *de novo* transcriptome analysis of roots, leaves, stems, and flowers of *Ferula assafoetida* (Apiaceae) was combined with computational annotation to identify candidate genes with probable roles in terpenoid and coumarin biosynthesis [Amini et al, 2019]. Genes of the phenylpropanoid pathway identified were phenylalanine lyase, 4-coumaryl CoA-ligase and C2′H. C2′H is important in the biosynthesis of 7-hydroxycoumarins. Genes of terpene meeolism, including a number or mono-, sesqui- and diterpene synthases, were identified. Finally, 245 putative cytochrome P450 enzyme transcripts were identified. As discussed above, many cytochrome P450 enzymes are involved in the biosynthesis of sesquiterpene coumarins.

The genome of *Artemisia annua* is the first sequenced genome of a plant belonging to the Asteraceae family [Shen et al, 2018]. *A. annua* is cultivated as the source of the anti-malarial sesquiterpene artemisinin. Sesquiterpene coumarins have been isolated from a number of Artemisia species including *A. annua* (Table S1). The 1.74-gigabase genome of *A. annua* contains 63226 protein-coding genes, which makes it a rich source for identification and isolation of enzymes involved in the biosynthesis of sesquiterpene coumarins.

*Tanacetum cinerariifolium* (Asteraceae) is known for the biosynthesis of the pyrethrin class of insecticides. A 7.1-Gb draft genome of *T. cinerariifolium*, consisting of 60080 genes, has been reported [Yamashiro et al, 2019]. This genome is another rich source for the identification and isolation of enzymes involved in the biosynthesis of sesquiterpene coumarins. At least two reports on two *Tanacetum* species (*T. heterotumum* and *T. parthenium*) producing sesquiterpene coumarins have been published (Table 1S) [Goren et al, 1988; Kisiel & Stojakowska, 1997].

It has been pointed out that nearly all the sesquiterpene coumarins described in this text have been isolated from plants belonging to the Apiaceae and Asteraceae families. Only a handful of sesquiterpene coumarins, shown in Figure 29, have been isolated from other families (Amaranthaceae, Caprifoliaceae, Euphorbiaceae, Lamiaceae, Lythraceae, Myrtaceae, Rutaceae, Solanaceae). However, the biosynthesis of the sesquiterpene coumarins from these families does not involve STCSs. It is suggested that STCS genes of plants belonging to the Apiaceae and Asteraceae families have evolved from ancestorial OSC genes. An indication of this is that the 10′,11′-oxidoumbelliprenin (**28**) fits well in the active site of lanosterol cyclase as shown by docking experiments (Figure 9). Strong evidence would be to show that OSCs can use 10′,11′-oxidoumbelliprenin (**28**) as a substrate (*cf*. Tables 2 and 3). The final proof will the cloning of genes encoding STCSs.

## Abbreviations

2OGX: 2-oxoglutarate-dependent oxygenase
4-CL: 4-coumaryl CoA-ligase
BGC: biosynthetic gene cluster
BIS: bisphenyl synthase
C2′H: coumaroyl-2′-hydroxylase
CAS: cycloartenol synthase
COSY: coumarin synthase
CPR: NADPH-cytochrome P450 reductase
DMADP: dimethylallyl diphosphate
FDP: farnesyl diphosphate
FMO: flavin monooxygenase
GAS: galbanic aldehyde synthase
GDP: geranyl diphosphate
GGDP: geranylgeranyl diphosphate
HG: homogentisate
HSD/D: 3β-hydroxysteroid dehydrogenases/C4 decarboxylase
IDP: isopentenyl diphosphate
LAS: lanosterol synthase
MEP: methylerythritol phosphate
MVA: mevalonic acid
OSC: oxidosqualene cyclase
P450: cytochromeP450 monooxygenase
PHB: p-hydroxybenzoic acid
PT: prenyltransferases
SHC: squalene hopene cyclase
SMO: sterol methyl oxidase
SR: 3-oxosteroid reductase
STCE: sesquiterpene coumarin epoxidase
STCS: sesquiterpene coumarin synthase
STS: sesquiterpene synthase
SQE: squalene epoxidase
TPS: terpene synthases
UMO: umbelliprenin monooxygena

## References

Abd El-Razek MH, Wu Y-Ch, Chang F-R (2007) Sesquiterpene coumarins from Ferula foetida. J Chin Chem Soc 54:235–238 10.1002/jccs.200700035

Abe I (2020) Biosynthesis of medicinally important plant metabolites by unusual type III polyketide synthases. J Nat Med 74:639–646 10.1007/s11418-020-01414-9

Ahmad MZ, Ali M, Showkat R, Mir SR (2012) New sesqui- and diterpenic coumarin ethers from the roots of Aegle marmelos (L.) Corr. Nat Prod J 2:252–258 10.2174/2210315511202040252

Ahmed AA (1999) Sesquiterpene coumarins and sesquiterpenes from Ferula sinaica. Phytochemistry 50:109–112 10.1016/S0031-9422(98)00489-0

Ahmed AA, Mohamed AE-HH, Abd El-Razek MH, Hegazy M-EF (2007) Ferulol and epi-samarcandin, two new sesquiterpene coumarins from Ferula sinaica. Nat Prod Comm 2:521–524 https://doi-org.proxy.lnu.se/10.1177/1934578X070020050

Ajani Y, Ajani A, Cordes JM, Watson MF, Downie SR (2008). Phylogenetic analysis of nrDNA ITS sequences reveals relationships within five groups of Iranian Apiaceae subfamily Apioideae. Taxon 57:383–401 http://www.jstor.org/stable/25066011

Akashi T, Sasaki K, Aoki T, Ayabe S, Yazaki K (2009) Molecular cloning and characterization of a cDNA for pterocarpan 4-dimethylallyltransferase catalyzing the key prenylation step in the biosynthesis of glyceollin, a soybean phytoalexin. Plant Physiol 149:683–693 http://www.plantphysiol.org/cgi/doi/10.1104/pp.108.123679

Alcalde MA, Perez-Matas E, Escrich A, Cusido RM, Palazon J, Bonfill M (2022) Biotic elicitors in adventitious and hairy root cultures: A review from 2010 to 2022. Molecules 27:5253 10.3390/molecules27165253

Al-Hazimi HMG, Basha RMY (1991) Phenolic compounds from various Artemisia species. J Chem Soc Pak 13:277–289

Amin A, Tuenter E, Cos P, Maes L, Exarchou V, Apers S, Pieters L (2016) Antiprotozoal and antiglycation activities of sesquiterpene coumarins from Ferula narthex exudate. Molecules 21:1287 10.3390/molecules21101287

Amini H, Naghavi MR, Shen T, Wang Y, Nasiri J, Khan IK, Fiehn O, Zerbe P, Maloof JN (2019) Tissue-specific transcriptome analysis reveals candidate genes for terpenoid and phenylpropanoid metabolism in the medicinal plant Ferula assafoetida. G3 9:807–816 10.1534/g3.118.200852

Appendino G, Ozen HC, Nano GM, Cisero M (1992) Sesquiterpene coumarin ethers from the genus Heptaptera. Phytochemistry 31:4223–4226 10.1016/0031-9422(92)80447-m

Appendino G, Tagliapietra S, Nano GM, Jakupovic J (1993) Sesquiterpene coumarin ethers from Asafetida. Phytochemistry 35:183–186 10.1016/S0031-9422(00)90530-2

Ashurov K, Numonov S, Guoruoluo Y, Aisa H-A, Turak A (2024) Unveiling the anti-vitiligo, anti-inflammatory, and antitumor activities of sesquiterpene coumarins isolated from Ferula kuhistanica. Fitoterapia 176:106035 10.1016/j.fitote.2024.106035

Bagirov VYu, Kiryalov NP, Sheichenko VI (1969) The coumarin tavicone from the roots of Ferula karatavica. Chem Nat Compd 5:504–505 10.1007/bf00568601

Bagirov VYu, Sheichenko VI (1976) The structure of tavicone. Chem Nat Compd 12:399–401 10.1007/BF00564795

Ban ZN, Qin H, Mitchell AJ, Liu B, Zhang F, Weng J-K, Dixon RA, Wang G (2018) Noncatalytic chalcone isomerase-fold proteins in Humulus lupulus are auxiliary components in prenylated flavonoid biosynthesis. Proc Natl Acad Sci USA, 115:E5223–E5232 10.1073/pnas.1802223115

Bandyopadhyay D, Basak B, Chatterjee A, Lai TK, Banerji A, Banerji J, Neuman A, Prangé T (2006) Saradaferin, a new sesquiterpenoid coumarin from Ferula assafoetida. Nat Prod Res 20:961–965 10.1080/1478641060082343

Banerjee A, Hamberger B (2018) P450s controlling metabolic bifurcations in plant terpene specialized metabolism. Phytochem Rev 17:81–111 10.1007/s11101-017-9530-4

Bharadwaj R, Kumar SR, Sharma A, Sathishkumar R (2021) Plant metabolic gene clusters: evolution, organization, and their applications in synthetic biology. Frontiers Plant Sci 12:697318 https://www.frontiersin.org/articles/10.3389/fpls.2021.697318

Barnaba C, Gentry K, Sumangala N, Ramamoorthy A (2017) The catalytic function of cytochrome P450 is entwined with its membrane-bound nature. F1000 Research 6:662 10.12688/f1000research.11015.1

Bittner M, Jakupovic J, Bohlmann F, Silva M (1988a) Coumarins and guaianolides from further Chilean representatives of the subtribe Nassauviinae. Phytochemistry 27:2867–2868 10.1016/S0031-9422(00)98112-3

Bittner M, Jakupovic J, Bohlmann F, Silva M (1988b) 5-Methylcoumarins from Nassauvia Species. Phytochemistry 27:3845–3847 10.1016/0031-9422(88)83029-2

Bohlmann J, Meyer-Gauen G, Croteau R (1998) Plant terpenoid synthases: Molecular biology and phylogenetic analysis. Proc Natl Acad Sci USA, 95:4126-4133 10.1073/pnas.95.8.4126

Bourgaud F, Hehn A, Larbat R, Doerper S, Gontier E, Kellner S, Matern U (2006) Biosynthesis of coumarins in plants: a major pathway still to be unravelled for cytochrome P450 enzymes. Phytochem Rev 5:293–308 10.1007/s11101-006-9040-2

Calvert MJ, Ashton PR, Allemann RK (2002) Germacrene A is a product of the aristolochene synthase-mediated conversion of farnesylpyrophosphate to aristolochene. J Am Chem Soc 124:11636–11641 10.1021/ja020762p

Cankar K, van Houwelingen A, Bosch D, Sonke T, Bouwmeester H, Beekwilder J (2011) A chicory cytochrome P450 mono-oxygenase CYP71AV8 for the oxidation of (+)-valencene. FEBS Lett 585:178–182 10.1016/j.febslet.2010.11.040

Cankar K, van Houwelingen A, Goedbloeda M, Reniriec R, de Jong RM, Bouwmeester H, Bosch D, Sonke T, Beekwilder J (2014) Valencene oxidase CYP706M1 from Alaska cedar (Callitropsis nootkatensis). FEBS Lett 588:1001–1007 10.1016/j.febslet.2014.01.061

Chen F, Tholl D, Bohlmann J, Pichersky E (2011) The family of terpene synthases in plants: a mid-size family of genes for specialized metabolism that is highly diversified throughout the kingdom. Plant J 66:212–229 10.1111/j.1365-313X.2011.04520.x

Chen NH, Wang SL, Smentek L, Hess BA, Wu RB (2015) Biosynthetic mechanism of lanosterol: cyclization. Angew Chem Int Ed 54:8693–8696 10.1002/anie.201501986

Chiba R, Minami A, Gomi K, Oikawa H (2013) Identification of ophiobolin F synthase by a genome mining approach: A sesterterpene synthase from Aspergillus clavatus. Org Lett 15:594–597 https://doi.org.proxy.lnu.se/10.1021/o1303498a

Christianson DW (2017) Structural and chemical biology of terpenoid cyclases. Chem Rev 117:11570–11648 10.1021/acs.chemrev.7b00287

Chua NK, Coates HW, Brown AJ (2020) Squalene monooxygenase: a journey to the heart of cholesterol synthesis. Prog Lipid Res 79:101033 10.1016/j.plipres.2020.101033

Çiçek Kaya A, Özbek H, Yuca H, Yilmaz G, Bingöl Z, Kazaz C, Gülçin İ, Güvenalp Z (2022). Phytochemical analysis and screening of acetylcholinesterase and carbonic anhydrase I and II isoenzymes inhibitory effect of Heptaptera triquetra (Vent.) Tutin root. Fabad J Pharm Sci 47:381–392 10.55262/fabadeczacilik.1147174

Cole TCH, Hilger HH, Stevens P (2019) Angiosperm phylogeny poster (APP) – Flowering plant systematics, 2019. PeerJ Preprints 7:e2320v6 10.7287/peerj.preprints.2320v6

Coleman T, Kirk AM, Chao RR, Podgorski MN, Harbort JS, Churchman LR, Bruning JB, Bernhardt PV, Harmer JR, Krenske EH, de Voss JJ, Bell SG (2021) Understanding the mechanistic requirements for efficient and stereoselective alkene epoxidation by a cytochrome P450 enzyme. ACS Catal 11:1995–2010 https://doi-org.proxy.lnu.se/10.1021/acscatal.0c04872

Corey EJ, Matsuda SP, Bartel B (1993) Isolation of an Arabidopsis thaliana gene encoding cycloartenol synthase by functional expression in a yeast mutant lacking lanosterol synthase by the use of a chromatographic screen. Proc Natl Acad Sci U S A 90:11628–11632. 10.1073/pnas.90.24.11628

Cravotto G., Balliano G., Robaldo B., Oliaro-Bosso S., Chimichi S., Boccalini M. (2004) Farnesyloxycoumarins, a new class of squalene-hopene cyclase inhibitors Bioorg Med Chem Lett 14:1931–1934 10.1016/j.bmcl.2004.01.085

Dargaeva TD, Brutko LI (1976) Coumarins of the epigeal part of Scabiosa comosa. Chem Nat Compd 12:337 https://doi-org.proxy.lnu.se/10.1007/BF00567814

da Silva Magedans YV, Phillips MA, Fett-Neto AG (2021) Production of plant bioactive triterpenoid saponins: from metabolites to genes and back. Phytochem Rev 20:461–482 https://doi-org.proxy.lnu.se/10.1007/s11101-020-09722-4

Dawidowicz AL, Bernacik K, Typek R (2018) Umbelliferone instability during an analysis involving its extraction process. Monatsh Chem 149:1327–1340 10.1007/s00706-018-2188-9

de Bruijn WJC, Levisson M, Beekwilder J, van Berkel WJH, Vincken J-P (2020) Plant aromatic prenyltransferases: tools for microbial cell factories. Trends Biotech 38:917–934 10.1016/j.tibtech.2020.02.006

De La Peña R, Hodgson H, Liu JC-T Stephenson MJ, Martin AC, Owen C, Harkess A, Leebens-Mack J, Jimenez LE, Osbourn A, Sattely ES (2023) Complex scaffold remodeling in plant triterpene biosynthesis. Science 379:361–368 10.1126/science.adf1017

Deng Y, Zhou Q, Wu Y, Chen X, Zhong F (2022) Properties and mechanisms of flavin-dependent monooxygenases and their applications in natural product synthesis. Int J Mol Sci 23:2622 10.3390/ijms23052622

Diao H, Chen N, Wang K, Zhang F, Wang Y-H, Wu R (2020) Biosynthetic mechanism of lanosterol: A completed story. ACS Catalysis 10:2157–2168 10.1021/acscatal.9b05221

Driller R, Janke S, Fuchs M, Warner E, Mhashal AR, Major DT, Christmann M,Brück T, Loll B (2018) Towards a comprehensive understanding of the structural dynamics of a bacterial diterpene synthase during catalysis. Nat Commun 9:3971 10.1038/s41467-018-06325-8

Dudareva N, Klempien A, Muhlemann JK, Kaplan I (2013) Biosynthesis, function and metabolic engineering of plant volatile organic compounds. New Phytol 198:16–22 10.1111/nph.12145

Edwards KG, Stoker RJ (1967) Biosynthesis of coumarin: The isomerization stage. Phytochemistry, 6:655–661 10.1016/S0031-9422(00)86004-5

Eichhorn E, Locher E, Guillemer S, Wahler D, Fourage L, Schilling B (2018) Biocatalytic process for (-)-ambrox production using squalene hopene cyclase. Adv Synth Catal 360:2339–2351 10.1002/adsc.201800132

Eruçar FM, Senadeera SPD, Wilson JA, Goncharova E, Beutler JA, Miski M (2023) Novel Cytotoxic Sesquiterpene Coumarin Ethers and Sulfur-Containing Compounds from the Roots of Ferula turcica. Molecules 28:5733 10.3390/molecules28155733

Fidan O, Zhan J (2018) Reconstitution of medicinally important plant natural products in microorganisms. In: Molecular pharming: Applications, challenges, and emerging areas. Kermode AR, ed. pp. 383–415. John Wiley & Sons, Inc https://ebookcentral-proquest-com.proxy.lnu.se/lib/linne-ebooks/detail.action?docID=5322112

Field B, Fiston-Lavier A-S, Kemen A, Geisler K, Quesneville H, Osbourn AE (2011) Formation of plant metabolic gene clusters within dynamic chromosomal regions. Proc Natl Acad Sci USA 108:16116–16121 10.1073/pnas.1109273108

Filippini R, Piovan A, Innocenti G, Caniato R, Cappelletti EM (1998) Production of coumarin compounds by Haplophyllum patavinum in vivo and in vitro. Phytochemistry 49:2337–2340 10.1016/S0031-9422(98)00356-2

Fiorito S, Epifano F, Palmisano R, Genovese S, Taddeo VA (2017) A re-investigation of the phytochemical composition of the edible herb Amaranthus retroflexus L. J Pharm Biomed Anal 143:183–187 https://doi-org.proxy.lnu.se/10.1016/j.jpba.2017.05.051

Fiorito S, Ianni F, Preziuso F, Epifano F, Scotti L, Bucciarelli T, Genovese S (2019a) UHPLC-UV/Vis quantitative analysis of hydroxylated and O-prenylated coumarins in Pomegranate seed extracts. Molecules 24:1963 10.3390/molecules24101963

Fiorito S, Preziuso F, Epifano F, Scotti L, Bucciarelli T, Taddeo VA, Genovese S (2019b) Novel biologically active principles from spinach, goji and quinoa. Food Chem 276:262–265 https://doi-org.proxy.lnu.se/10.1016/j.foodchem.2018.10.018

Fiorito S, Genovese S, Palumbo L, Scotti L, Ciulla M, di Profio P, Epifano F. (2020) Umbelliprenin as a novel component of the phytochemical pool from Artemisia spp. J Pharm Biomed Anal 184:113205 10.1016/j.jpba.2020.113205

Fiorito, S., Palumbo, L., Epifano, F. Fraternale D, Collevecchio C, Genovese S (2022) Modulation of the biosynthesis of oxyprenylated coumarins in calli from Ferulago campestris elicited by ferulic acid. Biomass Conversion and Biorefinery 10.1007/s13399-022-03309-z

Forestier E, Romero-Segura C, Pateraki I, Centeno E, Compagnon V, Preiss M, Berna A, Boronat A, Bach TJ, Darnet S, Schaller H (2019) Distinct triterpene synthases in the laticifers of Euphorbia lathyris. Sci Rep 9:4840 10.1038/s41598-019-40905-y

Fraatz MA, Berger RG, Zorn H (2009) Nootkatone-a biotechnological challenge. Appl Microbiol Biotechnol 83:35–4110.1007/s00253-009-1968-x

Gao K, Xu J, Sun J, Xu Y, Wei J, Sui C (2016) Molecular cloning and expression of squalene epoxidase from a medicinal plant, Bupleurum chinense. Chin Herb Med 8:67–74 10.1016/S1674-6384(16)60010-2

Girol CG, Fisch KM, Heinekamp T, Günther S, Huettel W, Piel J, Brakhage AA, Müller M (2012) Regio- and stereoselective oxidative phenol coupling in Aspergillus niger. Angew Chem Int Ed 51:9788–9791 10.1002/anie.201203603

Gliszczynska A, Brodelius PE (2012). Sesquiterpene coumarins. Phytochem Rev 11:77–96 10.1007/s11101-011-9220-6

Go YS, Lee SB, Kim HJ, Kim J, Park H-Y, Kim J-K, Shibata K, Yokota T, Ohyama K, Muranaka T, Arseniyadis S, Suh MC (2012) Identification of marneral synthase, which is critical for growth and development in Arabidopsis. Plant J 72:791–804 10.1111/j.1365-313X.2012.05120.x

Goren N, Ulubelen A, Oksüz S (1988) A sesquiterpene-coumarin ether and an acetylenic compound from Tanacetum heterotumum. Phytochemistry 27:1527–1529 10.1016/0031-9422(88)80231-0

Graf E, Alexa M (1985) Uber 5 neue Umbelliferonether aus Galbanumharz. Planta Med 51:428–431 10.1055/s-2007-969539

Greger H, Hofer O (1985) Sesquiterpene-coumarin ethers and polyacetylenes from Brocchia cinerea. Phytochemistry 24:85–88 10.1016/S0031-9422(00)80812-2

Greger H, Hofer O, Robien W (1983a) Types of sesquiterpene-coumarin ethers from Achillea ochroleuca and Artemisia tripartita. Phytochemistry 22:1997–2003 10.1016/0031-9422(83)80032-6

Greger H, Hofer O, Robien W (1983b) New sesquiterpene coumarin ethers from Achillea ochroleuca. ^13^C NMR of isofraxidin-derived open-chain and bicyclic sesquiterpene ethers. J Nat Prod 46:510-516 10.1021/np50028a015

Grob CA, Baumann W (1955). Die 1,4-Eliminierung unter Fragmentierung. Helv Chim Acta 38:594–610 10.1002/hlca.19550380306

Guengerich FP, Yoshimoto FK (2018) Formation and cleavage of C–C bonds by enzymatic oxidation–reduction reactions. Chem Rev 118:6573–6655 10.1021/acs.chemrev.8b00031

Guhling O, Hobl B,Yeats T, Jetter R (2006) Cloning and characterization of a lupeol synthase involved in the synthesis of epicuticular wax crystals on stem and hypocotyl surfaces of Ricinus communis. Arch Biochem Biophys 448:60–72 10.1016/j.abb.2005.12.013

Guo T, Dang W Zhou Y, Zhou D, Meng Q, Xu L, Chen G, Lin B, Qing D, Sun Y, Hou Y, Li N (2022) Sesquiterpene coumarins isolated from Ferula bungeana and their anti-neuroinflammatory activities. Bioorg Chem 128:106102 10.1016/j.bioorg.2022.106102

Gutensohn M, Orlova I, Nguyen TTH, Davidovich-Rikanati R, Ferruzzi MG, Sitrit Y, Lewinsohn E, Pichersky E, Dudareva N (2013). Cytosolic monoterpene biosynthesis is supported by plastid-generated geranyl diphosphate substrate in transgenic tomato. Plant J 75:351–363 10.1111/tpj.12212

Güvenalp Z, Özbek H, Yerdelen KÖ, Yilmaz G, Kazaz C, Demirezer LÖ (2017) Cholinesterase inhibition and molecular docking studies of sesquiterpene coumarin ethers from Heptaptera cilicica. Rec Nat Prod 11:462–467 10.25135/rnp.58.17.03.051

Hadi, N, Omideygi R, Amini A (2011). In vitro conservation of Ferula gummosa germplasm and its callus induction Iran J Rangelands Forests Plant Breed Gen Res 19:28–38 10.22092/ijrfpbgr.2011.7435

Hajeyah AA, Griffiths WJ, Wang Y, Finch AJ, O’Donnell VB (2020) The biosynthesis of enzymatically oxidized lipids. Front Endocrinol 11:591819 10.3389/fendo.2020.591819

Hammer SC, Dominicus JM, Syrén P-O, Nestl BM, Hauer B (2012) Stereoselective Friedel–Crafts alkylation catalyzed by squalene hopene cyclases. Tetrahedron 68:7624–7629 https://doi-org.proxy.lnu.se/10.1016/j.tet.2012.06.041

Hasegawa Y, Gong X, Kuroda C (2011) Chemical diversity of iridal-type triterpenes in Iris delavayi collected in Yunnan province of China. Nat Prod Comm 6:789–792 10.1177/1934578X1100600611

Hashimoto T, Hayashi A, Amano Y, Kohno J, Iwanari H, Usuda S, Yamada Y (1991) Hyoscyamine 6β-hydroxylase, an enzyme involved in tropane alkaloid biosynthesis, is localized at the pericycle of the root. J Biol Chem 266:4648–4653 10.1016/S0021-9258(20)64371-X

He F, Zhu Y, He M, Zhang Y (2008) Molecular cloning and characterization of the gene encoding squalene epoxidase in Panax notoginseng. DNA Seq 19:270–273 10.1080/10425170701575026

Henche S, Nestl BM, Hauer B (2021) Enzymatic Friedel-Crafts alkylation using squalene-hopene cyclases. ChemCatChem 13:3405–3409 https://doi-org.proxy.lnu.se/10.1002/cctc.202100452

Herr CQ, Hausinger RP (2018) Amazing diversity in biochemical roles of Fe(II)/2-oxoglutarate oxygenases. Trends BiochemSci 43:517–532 10.1016/j.tibs.2018.04.002

Hofberger JA, Nsibo DL, Govers F, Bouwmeester K, Schranz ME (2015) A complex interplay of tandem- and whole-genome duplication drives expansion of the L-type lectin receptor kinase gene family in the brassicaceae. Genome Biol Evol 7:720–34. 10.1093/gbe/evv020

Hofer O, Greger H (1985) New sesquiterpene-coumarin ethers from Anthemis cretica. Liebigs Ann Chem 1985:1136–1144 10.1002/jlac.198519850604

Hofer O, Widhalm M, Greger H (1984) Circular dichroism of squiterpene-umbelliferone ethers and structure elucidation of a new derivative isolated from the gum resin “Asa Foetida”. Monatsh Chem 115:1207–1218 10.1007/BF00809352

Hoshino T, Kumai Y, Kudo I, Nakano S-I, Ohashi S (2004) Enzymatic cyclization reactions of geraniol, farnesol and geranylgeraniol, and those of truncated squalene analogs having C20 and C25 by recombinant squalene cyclase. Org Biomol Chem 2:2650–2657 10.1039/B407001A

Hou Y, Chen M, Sun, Z, Ma, G, Chen D, Wu H, Yang J, Li Y, Xu X (2022) The biosynthesis related enzyme, structure diversity and bioactivity abundance of indole-diterpenes: A review. Molecules 27:6870 10.3390/molecules27206870

Inoue K (2007) The chloroplast outer envelope membrane: The edge of light and excitement. J Interg Plant Biol 49:1100–111110.1111/j.1672-9072.2007.00543.x

Inoue T, Toyonaga T, Nagumo S, Nagai M (1989) Biosynthesis of 4-hydroxy-5-methylcoumarin in a Gerbera jamesonii hybrid. Phytochemistry 28:2329–2330 10.1016/S0031-9422(00)97977-9

Islam MdS, Leissing TM, Chowdhury R, Hopkinson RJ, Schofield CJ (2018) 2-Oxoglutarate-dependent-oxygenases. Annu Rev Biochem 87:585–620 10.1146/annurev-biochem-061516-044724

Jandl B, Hofer O, Kalchhauser H, Greger H (1997) Open chain sesquiterpene coumarin ethers and coniferylalcohol-4-O-farnesyl ether from Achillea ochroleuca. Nat Prod Lett 11:17–24 10.1080/10575639708043752

Jin S, Makris TM, Bryson TA, Sligar SG,. Dawson JH (2003) Epoxidation of olefins by hydroperoxo-ferric cytochrome P450. J Am Chem Soc 125:3406–3407 10.1021/ja029272n

Kang X, Wu L, Zhao C, Zhang C, Wang Q (2024) A new sesquiterpene coumarin from Ferula bungeana Kitagawa. Nat Prod Res 2024:1–5 10.1080/14786419.2024.2332490

Karppinen K, Hokkanen J, Mattila S, Neubauer P, Hohtola A (2008) Octaketide-producing type III polyketide synthase from Hypericum perforatum is expressed in dark glands accumulating hypericins. FEBS J 275:4329–4342 10.1111/j.1742-4658.2008.06576.x

Karunanithi PS, Zerbe P (2019) Terpene synthases as metabolic gatekeepers in the evolution of plant terpenoid chemical diversity. Front Plant Sci 10:1166 10.3389/fpls.2019.01166

Khayat MT, Alharbi M, Ghazawi KF, Mohamed GA, Ibrahim SRM (2023) Ferula sinkiangensis (Chou-AWei, Chinese Ferula): Traditional uses, phytoconstituents, biosynthesis, and pharmacological activities. Plants 12:902 10.3390/plants12040902

Khazaei A, Bahramnejad B, Mozafari AA, Dastan D, Mohammadi S (2019) Hairy root induction and farnesiferol B production of endemic medicinal plant Ferula pseudalliacea. 3 Biotech 9:407 10.1007/s13205-019-1935-x

Kir’yalov N.P. (1961) The structure of cocanicine and umbelliprenin, components of the neutral part of oil from Ferula cocanica. Tr Bot Inst Akad Nauk SSSR Ser 5 8:7

Kisiel W, Stojakowska A (1997) A sesquiterpene coumarin ether from transformed roots of Tanacetum parthenium. Phytochemistry 46:515–516 10.1016/S0031-9422(97)87091-4

Kranz-Finger S, Mahmoud O, Ricklefs E, Ditz N, Bakkes PJ, Urlacher VB (2018) Insights into the functional properties of the marneral oxidase CYP71A16 from Arabidopsis thaliana. BBA-Proteins Proteom 1866:2–10 10.1016/j.bbapap.2017.07.008

Kuliev ZA, Khasanov TKh (1978) Structures of ferocaulin, ferocaulinin, ferocaulidin, and ferocaulicin. Chem Nat Compd 14:267–271 https://doi-org.proxy.lnu.se/10.1007/BF00713313

Kühnel LC, Nestl BM, Hauer B (2017) Enzymatic addition of alcohols to terpenes by squalene hopene cyclase variants. ChemBioChem 18:2222–2225 10.1002/cbic.201700449

Lee Ch-L, Chiang L-Ch, Cheng L-H, Liaw Ch-Ch, Abd El-razek MH, Chang F-R, Wu Y-Ch (2009) Influenza A (H1N1) antiviral and cytotoxic agents from Ferula assa-foetida. J Nat Prod 72:1568–1572 10.1021/np900158f

Li G, Wang J, Li X, Cao L, Gao L, Lv N, Si J (2016) An unusual sesquiterpene coumarin from the seeds of Ferula sinkiangensis, J Asian Nat Prod Res 18:891–896 10.1080/10286020.2016.1168813

Li G, Wang J, Li X, Cao L, Lv N, Chen G, Zhu J, Si J (2015a) Two new sesquiterpene coumarins from the seeds of Ferula sinkiangensis. Phytochem Lett 13:123–126 10.1016/j.phytol.2015.06.002

Li H, Sun Y, Zhang Q, Zhu Y, Li S-M, Li A, Zhang C (2015b) Elucidating the cyclization cascades in xiamycin biosynthesis by substrate synthesis and enzyme characterizations. Org Lett 17:306–309 10.1021/ol503399b

Li J, Liao H-J, Tang Y, Huang J-L, Cha L, Lin T-S, Lee JL, Kurnikov IV, Kurnikova MG, Chang W-C, Chan N-L, Guo Y (2020c) Epoxidation catalyzed by the nonheme iron(II)- and 2-oxoglutarate-dependent oxygenase, AsqJ: Mechanistic elucidation of oxygen atom transfer by a ferryl intermediate. J Am Chem Soc 142:6268–6284 10.1021/jacs.0c00484

Li N, Guo T-T, Zhou D (2018) Chapter 8 - Bioactive sesquiterpene coumarins from plants. In: Atta-ur-Rahman (ed) Studies in natural products chemistry, Vol 59 Elsevier, pp. 251–282 10.1016/B978-0-444-64179-3.00008-6

Li W., Shao Y.-T., Yin T.-P., Yan H., Shen B.-C., Li Y.-Y., Xie H.-D., Sun Z.-W., Ma Y.-L. (2020b) Penisarins A and B, sesquiterpene coumarins isolated from an endophytic Penicillium sp. J Nat Prod 83:3471–3475 10.1021/acs.jnatprod.0c00393

Li Y, Leveau A, Zhao Q, Feng Q, Lu H, Miao J, Xue Z, Martin AC, Wegel E, Wang J, Orme A, Rey M-D, Karafiátová M, Vrána J, Steuernagel B, Joynsom R, Owen C, Reed J, Louveau T, Stephensom MJ, Zhang L, Huang X, Huang T, Fan D, Zhou C, Tian Q, Li W, Lu Y, Chen J, Zhao Y, Lu Y, Zhu C, Liu Z, Polturak G, Casson R, Hill L, Moore G, Melton R, Hall N, Wulff BBH, Dolezel J, Langdon T, Han B, Osbourn A (2021) Subtelomeric assembly of a multi-gene pathway for antimicrobial defense compounds in cereals. Nat Commun 12:2563 10.1038/s41467-021-22920-8

Li Z, Jiang Y, Guengerich FP, Ma L, Li S, Zhang W (2020a) Engineering cytochrome P450 enzyme systems for biomedical and biotechnological applications J Biol Chem 295:833–849 10.1016/S0021-9258(17)49939-X

Lin CI, McCarty RM, Liu HW (2017) The enzymology of organic transformations: a survey of name reactions in biological systems. Angew Chem Int Ed 56:3446–3489 10.1002/anie.201603291

Liu B, Raeth T, Beuerle T, Beerhues L (2010) A novel 4-hydroxycoumarin biosynthetic pathway. Plant Mol Biol 72:17–25 10.1007/s11103-009-9548-0

Liu Z, Cheema J, Vigouroux M, Hill L, Reed J, Paajanen P, Yant L, Osbourn A (2020) Formation and diversification of a paradigm biosynthetic gene cluster in plants. Nat Commun 11:5354 10.1038/s41467-020-19153-6

Ma C-H, Jiang N-H, Deng M-H, Chen J-W, Yang S-C, Long G-Q, Zhang G-H (2016) Cloning and characterization of three squalene epoxidase genes in Panax vietnamensis var. fuscidicus, a rare medicinal plant with high content of ocotillol-type ginsenosides. Pak J Bot 48:2453–2465

Magallón S, Gómez-Acevedo S, Sánchez-Reyes LL, Tania Hernández-Hernández T (2015) A metacalibrated time-tree documents the early rise of flowering plant phylogenetic diversity. New Phytologist 207:437–453 10.1111/nph.13264

Mandel JR, Dikow RB, Siniscalchi CM, Thapa R, Watson LE, Funk VA (2019) A fully resolved backbone phylogeny reveals numerous dispersals and explosive diversifications throughout the history of Asteraceae. PNAS 116:14083–14088 www.pnas.org/cgi/doi/10.1073/pnas.1903871116

Marco JA, Sanz JF, Yuste A, Rustaiyan A (1991) New umbelliferone sesquiterpene ethers from roots of Ligularia persica. Liebigs Ann Chem 1991:929–931 10.1002/jlac.1991199101158

Marner FJ, Kasel T (1995). Biomimetic synthesis of the iridal skeleton. J Nat Prod 58:319–323 10.1021/np50116a030

Marner FJ, Littek A, Spitzfaden C, Bell A, Jaenicke L (1989) Studies on the biosynthesis of unusual triterpenoids from sword-lilies. Planta Med 55:676 10.1055/s-2006-962281

Martinez S, Hausinger XRP (2015) Catalytic mechanisms of Fe(II)-and 2-oxoglutarate-dependent oxygenases. J Biol Chem 290:20702–20711 10.1074/jbc.R115.648691

Matos MJ, Santana L, Uriarte E, Abreu OA, Molina E. Yordi EG (2015). Coumarins - An important class of phytochemicals. In: Rao AV, Rao LG (eds) Phytochemicals - isolation, characterisation and role in human health, IntechOpen, pp 113–140 10.5772/59982

Matsuda Y, Abe I (2016) Biosynthesis of fungal meroterpenoids. Nat Prod Rep 33:26–53 10.1039/C5NP00090D

Meguro A, Motoyoshi Y, Teramoto K, Ueda S, Totsuka Y, Ando Y, Tomita T, Kim S-Y, Kimura T, Igarashi M, Sawa R, Shinada T, Nishiyama M, Kuzuyama T (2015) An unusual terpene cyclization mechanism involving a carbon-carbon bond rearrangement. Angew Chem Int Ed 54:4353–4356 10.1002/ange.201411923

Miski M, Tosun F, Aytar EC, Duran A (2015) Novel sesquiterpene coumarin ethers from the dichloromethane extract of the roots of Heptaptera cilicica. Conference Paper OP-22, 11th Interntional Symposium on the Chemistry of Natural Compounds, 1-6 October, 2015, Antalya, Turkey

Mitchell AJ, Weng J-K (2019) Unleashing the synthetic power of plant oxygenases: from mechanism to application. Plant Physiol 179:813–829 10.1104/pp.18.01223

Munakata R, Olry A, Takemura T, Tatsumi K, Ichino T, Villard C, Kageyama J, Kurata T, Nakayasu M, Jacob F, Koeduka T, Yamamoto H, Moriyoshi E, Matsukawa T, Grosjean J, Krieger C, Sugiyama A, Mizutani M, Bourgaud M, Hehn A, Yazaki K (2021) Parallel evolution of UbiA superfamily proteins into aromatic O-prenyltransferases in plants. Proc Nat Acad Sci 118:e2022294118 10.1073/pnas.2022294118

Munakata R, Takemura T, Tatsumi K, Moriyoshi Yanagihara K, Sugiyama A, Suzuki H, Seki H, Muranaka T, Kawano N, Yoshimatsu K, Kawahara N, Yamaura T, Grosjean J, Bourgaud F, Hehn A, Yazaki K (2019) Isolation of Artemisia capillaris membrane-bound di-prenyltransferase for phenylpropanoids and redesign of artepillin C in yeast. Comm Biol 2:384 10.1038/s42003-019-0630-0

Murch SJ, Rupasinghe HPV, Goodenowe D, Saxena PK (2004) A metabolomic analysis of medicinal diversity in Huang-qin (Scutellaria baicalensis Georgi) genotypes: discovery of novel compounds. Plant Cell Rep 23:419–425 10.1007/s00299-004-0862-3

Najafabadi AS, Naghavi MR, Farahmand H, Abbasi A (2017) Transcriptome and metabolome analysis of Ferula gummosa Boiss. to reveal major biosynthetic pathways of galbanum compounds. Funct Integr Genomics 17:725–737 10.1007/s10142-017-0567-7

Nes WD (2011) Biosynthesis of cholesterol and other sterols. Chem Rev 111:6423–6451 10.1021/cr200021m

Nikolaiczyk V, Kirschning A, Diaz E (2022) Lipoxygenasecatalysed co-oxidation for sustained production of oxyfunctionalized terpenoids. Flavour Fragr J. 37:234–242 10.1002/ffj.3700

Oliaro-Bosso S, Viola F, Taramino S, Tagliapietra S, Barge A, Cravotto G, Balliano G (2007) Inhibitory effect of umbelliferone aminoalkyl derivatives on oxidosqualene cyclases from S. cerevisiae, T. cruzi, P. carinii, H. sapiens, and A. thaliana: a structure–activity study. ChemMedChem 2:226–233 10.1002/cmdc.200600234

Padyana AK, Gross S, Jin L, Cianchetta C, Narayanaswamy R, Wang F, Wang R, Fang C, Lv X, Biller SA, Dang L, Mahoney CE, Nagaraja N, Pirman D, Sui Z, Popovici-Muller J, Smolen GA (2019) Structure and inhibition mechanism of the catalytic domain of human squalene epoxidase. Nat Commun 10:97 10.1038/s41467-018-07928-x.

Panchy N, Lehti-Shiu M, Shiu S-H (2016) Evolution of gene duplication in plants. Plant Physiol 171:2294–2316 10.1104/pp.16.00523

Pascal S, Taton M, Rahier A (1993) Plant sterol biosynthesis. Identification and characterization of two distinct microsomal oxidative enzymatic systems involved in sterol C4-demethylation. J Biol Chem 268:11639–11654 10.1016/S0021-9258(19)50249-6

Phillips DR, Rasbery JM, Bartel B, Matsuda SPT (2006) Biosynthetic diversity in plant triterpene cyclization. Curr Opin Plant Biol 9:305–314 https://doi-org.proxy.lnu.se/10.1016/j.pbi.2006.03.004

Piechulla B, Bartelt R, Brosemann A, Effmert U, Bouwmeester H, Hippauf F, Brandt W (2016) The α-terpineol to 1,8-cineole cyclization reaction of tobacco terpene synthases. Plant Physiol 172:2120–2131 10.1104/pp.16.01378

Pietiäinen M, Kontturi J, Paasela T, Deng X, Ainasoja M, Nyberg P, Hotti H, Teeri T (2016) Two polyketide synthases are necessary for 4-hydroxy-5-methylcoumarin biosynthesis in Gerbera hybrida. Plant J 87:548–558 10.1111/tpj.13216

Porter NA (2013) A perspective on free radical autoxidation: the physical organic chemistry of polyunsaturated fatty acid and sterol peroxidation. J Org Chem 78:3511–3524 10.1021/jo4001433

Qiao J, Liu J, Liao J, Luo Z, Ma X, Ma G (2018) Identification of key amino acid residues determining product specificity of 2,3-oxidosqualene cyclase in Siraitia grosvenorii. Catalysts 8:577 10.3390/catal8120577

Quílez del Moral JF, Pérez AI. Alejandro F. Barrero AF (2020) Chemical synthesis of terpenoids with participation of cyclizations plus rearrangements of carbocations: a current overview. Phytochem Rev 19:559–576 10.1007/s11101-019-09646-8

Rahier A (2011) Dissecting the sterol C-4 demethylation process in higher plants. From structures and genes to catalytic mechanism. Steroids 76:340–352 https://doi.org.proxy.lnu.se/10.1016/j.steroids.2010.11.011

Rasbery JM, Shan H, LeClair RJ, Norman M, Matsuda SPT, Bartel B (2007) Arabidopsis thaliana squalene epoxidase 1 is essential for root and seed development. J Biol Chem 282:17002–17013 10.1074/jbc.M611831200

Razdan T, Qadri B, Qurishi M, Khuroo M, Kachroo P (1989). Sesquiterpene esters and sesquiterpene-coumarin ethers from Ferula jaeskeana. Phytochemistry 28:3389–3393 10.1016/0031-9422(89)80353-X

Rising KA, Starks CM, Noel JP, Chappell J (2000) Demonstration of germacrene A as an intermediate in 5-epi-aristolochene synthase catalysis. J Am Chem Soc 122:1861–1866 10.1021/ja993584h

Rudolf JD, Dong L-B, Zhang X, Renata H, Shen B (2018) Cytochrome P450-catalyzed hydroxylation initiating ether formation in platensimycin biosynthesis. J Am Chem Soc 140:12349–12353 10.1021/jacs.8b08012

Saeki H, Hara R, Takahashi H, Iijima M, Munakata R, Kenmoku H, Fuku K, Sekihara A, Yasuno Y, Shinada T, Ueda D, Nishi T, Sato T, Asakawa Y, Kurosaki F, Yazaki K, Taura F (2018) An aromatic farnesyltransferase functions in biosynthesis of the anti-HIV meroterpenoid daurichromenic acid. Plant Physiol 178:535–551 10.1104/pp.18.00655

Saikia S, Nicholson MJ, Young C, Parker EJ, Scott B (2008) The genetic basis for indole-diterpene chemical diversity in filamentous fungi. Mycolog Res 112:184–199 10.1016/j.mycres.2007.06.015

Sarker SD, Nahar L (2017). Progress in the chemistry of naturally occurring coumarins. In: Kinghorn A, Falk H, Gibbons S, Kobayashi J (eds) Progress in the chemistry of organic natural products vol 106, pp 241–304, Springer, Cham. 10.1007/978-3-319-59542-9_3

Scotti L, Genovese S, Bucciarelli T, Martini F, Epifano F, Fiorito S, Preziuso F, Taddeo VA (2018) Analysis of biologically active oxyprenylated phenylpropanoids in tea tree oil using selective solid-phase extraction with UHPLC-PDA detection. J Pharm Biomed Anal 154:174–179 https://doi-org.proxy.lnu.se/10.1016/j.jpba.2018.03.004

Seitz M, Klebensberger J, Siebenhaller S, Breuer M, Siedenburg G, Jendrossek D, Hauer B (2012) Substrate specificity of a novel squalene-hopene cyclase from Zymomonas mobilis, J Mol Catal B: Enz 84:72–77 10.1016/j.molcatb.2012.02.007

Seitz M, Syren PO, Steiner L, Klebensberger J, Nestl BM, Hauer B (2013a) Synthesis of heterocyclic terpenoids by promiscuous squalene hopene cyclases. ChemBioChem 14:436–439 10.1002/cbic.201300018

Shahzadi I, Ali Z, Baek SH, Mirza, B Ahn KS (2020) Assessment of the antitumor potential of umbelliprenin, a naturally occurring sesquiterpene coumarin. Biomedicines 8:126 10.3390/biomedicines8050126

Sharifi-Rad J, Cruz-Martins N, López-Jornet P, Pons-Fuster Lopez E, Harun N, Yeskaliyeva B, Beyatli A, Sytar O, Shaheen S, Sharopov F, Taheri Y, Docea AO, Calina D, Cho WC (2021) Natural coumarins: exploring the pharmacological complexity and underlying molecular mechanisms. In: Oxidative Medicine and Cellular Longevity, Hindawi, 2021, Article ID 6492346, 19 pages 10.1155/2021/6492346

Shen Q, Zhang L, Liao Z, Wang S, Yan T, Shi P, Liu M, Fu X, Pan Q, Wang Y, Lv Z, Lu X, Zhang F, Jiang W, Ma Y, Chen M, Hao X, Li L, Tang Y, Lv G, Zhou Y, Sun X, Brodelius PE, Rose JKC, Tang K (2018) The genome of Artemisia annua provides insight into the evolution of Asteraceae family and artemisinin biosynthesis. Mol Plant 11:776–788 10.1016/j.molp.2018.03.015

Shibuya M, Sagara A, Saitoh A, Kushiro T, Ebizuka Y (2008) Biosynthesis of baccharis oxide, a triterpene with a 3,10-oxide bridge in the A-ring. Org Lett 10:5071–5074 https://doi-org.proxy.lnu.se/10.1021/ol802072y

Shibuya M, Xiang T, Katsube Y, Otsuka M, Zhang H, Ebizuka Y (2007) Origin of structural diversity in natural triterpenes: direct synthesis of seco-triterpene skeletons by oxidosqualene cyclase. J Am Chem Soc 129:1450–1455 10.1021/ja066873w

Shimizu B-I (2014) 2-Oxoglutarate-dependent dioxygenases in the biosynthesis of simple coumarins. Front Plant Sci 5:549 10.3389/fpls.2014.00549

Shomirzoeva O, Xu M-Y, Sun Z-J, Li C, Nasriddinov A, Muhidinov Z, Zhang K, Gu Q, Xu J (2021) Chemical constituents of Ferula seravschanica. Fitoterapia 149:104829 10.1016/j.fitote.2021.104829

Sidana J, Saini V, Dahiya S, Nain P, Bala S (2013) A review on Citrus – “The boon of nature”. Int J Pharm Sci Rev Res 18:20–27

Smit SJ, Lichman BR (2022) Plant biosynthetic gene clusters in the context of metabolic evolution. Nat Prod Rep 39:1465–1482 10.1039/d2np00005a

Song W, Yan S, Li Y, Feng S, Zhang J, Li J (2019) Functional characterization of squalene epoxidase and NADPH-cytochrome P450 reductase in Dioscorea zingiberensis. Biochem Biophys Res Comm 509:822–827 10.1016/j.bbrc.2019.01.010

Steele CL, Crock J, Bohlmann J, Croteau R (1998) Sesquiterpene Synthases from Grand Fir (Abies grandis): Comparison of constitutive and wound-induced activities, and cDNA isolation, characterization, and bacterial expression of δ-selinene synthase and γ-humulene synthase. J Biol Chem 273:2078–2089 10.1074/jbc.273.4.2078

Sukumaran A, McDowell T, Chen L, Renaud J, Dhaubhadel S (2018) Isoflavonoid-specific prenyltransferase gene family in soybean: GmPT01, a pterocarpan 2-dimethylallyltransferase involved in glyceollin biosynthesis. Plant J 96:966–981 10.1111/tpj.14083

Suran, D., Bolor, T., & Bayarmaa, G.-A. (2016) In vitro seed germination and callus induction of Ferula ferulaeoides (Steud.) Korov. (Apiaceae). Mongol J Biol Sci 14:53–58 https://www.biotaxa.org/mjbs/article/view/27920

Sutthivaiyakit S, Mongkolvisut W, Prabpai S, Kongsaeree P (2009) Diterpenes, sesquiterpenes, and a sesquiterpene coumarin conjugate from Jatropha integerrima. J Nat Prod 72:2024–2027 10.1021/np900342b

Takahashi S, Yeo YS, Zhao Y, O’Maille PE, Greenhagen BT, Noel JP, Coates RM, Chappell J (2007) Functional characterization of premnaspirodiene oxygenase, a cytochrome P450 catalyzing regio- and stereo-specific hydroxylations of diverse sesquiterpene substrates. J Biol Chem 282:31744–31754 10.1074/jbc.M703378200

Tanaka H, Nogushi H, Abe I (2005) Enzymatic formation of indole-containing unnatural cyclic polyprenoids by bacterial squalene:hopene cyclase. Org Lett 7:5873–5876 10.21/o1052507q

Tanaka H, Noma H, Noguchi H, Abe I (2006) Enzymatic formation of pyrrole-containing novel cyclic polyprenoids by bacterial squalene: hopene cyclase. Tetrahedron Lett 47:3085–3089 10.1016/j.tetlet.2006.02.151

Tang MC, Zou Y, Watanabe K, Walsh CT, Tang Y (2017) Oxidative cyclization in natural product biosynthesis. Chem Rev 117:5226–5333 10.1021/acs.chemrev.6b00478

Tansakul P, Shibuya M, Kushiro T, Ebizuka Y (2006) Dammarenediol-II synthase, the first dedicated enzyme for ginsenoside biosynthesis in Panax ginseng. FEBS Lett 580:5143–5149 10.1016/j.febslet.2006.08.044

Tashkhodzhaev B, Turgunov KK, Izotova LY, Kamoldinov KS (2015) Stereochemistry of samarcandin-type sesquiterpenoid coumarins. Crystal structures of feshurin and nevskin. Chem Nat Compd 51:242–246 10.1007/s10600-015-1253-4

Taton M, Rahier A (1996) Plant sterol biosynthesis: identification and characterization of higher plant Δ7-sterol C5(6)-desaturase. Arch Biochem Biophys 325:279–288 10.1006/abbi.1996.0035

Taura F, Iijima M, Yamanaka E, Takahashi H, Kenmoku H, Saeki H, Morimoto S, Asakawa Y, Kurosaki F and Morita H (2016) A novel class of plant type III polyketide synthase involved in orsellinic acid biosynthesis from Rhododendron dauricum. Front Plant Sci 7:1452 10.3389/fpls.2016.01452

Teng L, Ma GZ, Li L, Ma LY, Xu XQ (2013) Karatavicinol A, a new anti-ulcer sesquiterpene coumarin from Ferula sinkiangensis. Chem Nat Compd 49:606–609 https://doi-org.proxy.lnu.se/10.1007/s10600-013-0690-1

Thimmappa R, Geisler K, Louveau T, O’Maille P, Osbourn A (2014) Triterpene biosynthesis in plants. Annu Rev Plant Biol 65:225–257 10.1146/annurev-arplant-050312-120229

Thoma R, Schulz-Gasch T, D’Arcy B, Benz J Aebi J, Dehmlow H, Hennig M Stihle M, Ruf A (2004) Insight into steroid scaffold formation from the structure of human oxidosqualene cyclase. Nature 432:118–122 10.1038/nature02993

Tian X-H, Hong L-L, Jiao W-H, Lin H-W (2023) Natural sesquiterpene quinone/quinols: chemistry, biological activity, and synthesis. Nat Prod Rep 40:718–749 10.1039/D2NP00045H

Tosun F, Aytar EC, Beutler JA, Wilson JA, Miski M (2021) Cytotoxic sesquiterpene coumarins from the roots of Heptaptera cilicica. Rec Nat Prod 15:529–536 10.25135/rnp.242.21.02.1990

Tosun F, Beutler JA, Ransom TT, Miski M (2019) Anatolicin, a highly potent and selective cytotoxic sesquiterpene coumarin from the root extract of Heptaptera anatolica. Molecules 24:1153 10.3390/molecules24061153

Vandyshev VV, Sklyar YE, Perel’son ME, Moroz MD, Pimenov MG (1972a) Conferol, a new coumarin from the roots of Ferula conocaula and F. moschata. Khim Prir Soedin 670 10.1007/BF00564346

Vandyshev VV, Sklyar YE, Perel’son ME, Moroz MD, Pimenov MG (1972b) Conferone, a new terpenoid coumarin from the fruit of Ferula conocaula. Khim Prir Soedin 669 10.1007/BF00564345

Vanholme R, Sundin L, Seetso KC, Kim H, Liu X, Li J, De Meester B, Hoengenaert L, Goeminne G, Morreel K, Haustraet J, Tsai H-H, Schmidt W, Vanholme B, Ralph J, Boerjan W (2019) COSY Catalyzes trans-cis isomerization and lactonization in the biosynthesis of coumarins. Nature Plants 5:1066–1075 10.1038/s41477-019-0510-0

Vestena AS, Meirelles GdC, Zuanazzi JA, von Poser GL (2022) Taxonomic significance of coumarins in species from the subfamily Mutisioideae, Asteraceae. Phytochem Rev 22:85–112 https://doi-org.proxy.lnu.se/10.1007/s11101-022-09828-x

Villa-Ruano N, Pacheco-Hernández Y, Lozoya-Gloria E, Castro-Juárez CJ, Mosso-Gonzalez C, Ramirez-Garcia SA (2015) Cytochrome P450 from plants: platforms for valuable phytopharmaceuticals. Trop J Pharm Res 14:731–742 10.4314/tjpr.v14i4.24

Wang J, Huo X, Wang H, Dong A, Zheng Q, Si J (2023) Undescribed sesquiterpene coumarins from the aerial parts of Ferula sinkiangensis and their anti-inflammatory activities in lipopolysaccharide-stimulated RAW 264.7 macrophages. Phytochemistry 210:113664 10.1016/j.phytochem.2023.113664

Wang J, Wang H, Zhang M, Li X, Zhao Y, Chen G, Si J, Jiang L (2020) Sesquiterpene coumarins from Ferula sinkiangensis K.M.Shen and their cytotoxic activities. Phytochemistry 180:112531 10.1016/j.phytochem.2020.112531

Wang P, Wei G, Feng L (2022a) Research advances in oxidosqualene cyclase in plants. Forests 13:1382 10.3390/f13091383

Wang M, Zhou X, Wang Z, Chen Y (2022b), Enzyme-catalyzed allylic oxidation reactions: A mini-review. Front Chem 10:950149 10.3389/fchem.2022.950149

Wen J, Xie D-F, Price M, Ren T, Deng Y-Q, Gui L-J, Guo X-L, He X-J (2021) Backbone phylogeny and evolution of Apioideae (Apiaceae): New insights from phylogenomic analyses of plastome data. Mol Phylogenet Evol 161:107183 10.1016/j.ympev.2021.107183

Williams DC, Carroll BJ, Jin Q, Rithner GD, Lenger SR, Floss HGM, Coates RM, Williams RM, Croteau R (2000) Intramolecular proton transfer in the cyclization of geranylgeranyl diphosphate to the taxadiene precursor of taxol catalyzed by recombinant taxadiene synthase. Chem Biol 7:969–977 10.1016/S1074-5521(00)00046-6

Winkelblech J, Fan A, Li SM (2015) Prenyltransferases as key enzymes in primary and secondary metabolism. Appl Microbiol Biotechnol 99:7379-7397 10.1007/s00253-015-6811-y

Xing Y, Li N, Zhou D, Chen G, Jiao K, Wang W, Si Y, Hou Y (2017) Sesquiterpene coumarins from Ferula sinkiangensis act as neuroinflammation inhibitors. Planta Med 83:135–142 10.1055/s-0042-109271

Xiong Q, Wilson WK, Matuda SPT (2006) An Arabidopsis oxidosqualene cyclase catalyzes iridal skeleton formation via grob fragmentation. Angew Chem 118:1307–1310 10.1002/ange.200503420

Xiong Q, Zhu X, Wilson WK, Ganesan A, Matsuda SPT (2003) Enzymatic synthesis of an indole diterpene by an oxidosqualene cyclase: Mechanistic, biosynthetic, and phylogenetic implications. J Am Chem Soc 125:9002–9003 10.1021/ja036322v

Xue Z, Duan L, Liu D, Guo J, Ge S, Dicks J, ÓMàille P, Osbourn A, Qi X (2012) Divergent evolution of oxidosqualene cyclases in plants. New Phytologist 193:1022–1038 10.1111/j.1469-8137.2011.03997.x

Yamashiro T, Shiraishi A, Satake H, Nakayama K (2019) Draft genome of Tanacetum cinerariifolium, the natural source of mosquito coil. Sci Rep 9:18249 10.1038/s41598-019-54815-6

Yang TH, Fang L, Sanders S, Jayanthi S, Rajan G, Podicheti R, Thallapuranam SK, Mockaitis K, Medina-Bolivar F (2018) Stilbenoid prenyltransferases define key steps in the diversification of peanut phytoalexins. J Biol Chem 293:128–146 10.1074/jbc.RA117.000564

Yang, X., Gao, S., Guo, L, Wang B, Jia Y, Zhou J, Che Y, Jia P, Lin J, Xu T, Sun J, Ye K (2021) Three chromosome-scale Papaver genomes reveal punctuated patchwork evolution of the morphinan and noscapine biosynthesis pathway. Nat Commun 12:6030 10.1038/s41467-021-26330-8

Yazaki K, Sasaki K, Tsurumaru Y (2009) Prenylation of aromatic compounds, a key diversification of plant secondary metabolites. Phytochemistry 70:1739–1745 10.1016/j.phytochem.2009.08.023

Yonemura Y, Ohyama T, Hoshino T (2012) Chemo-enzymatic syntheses of drimane-type sesquiterpenes and the fundamental core of hongoquercin meroterpenoid by recombinant squalene-hopene cyclase. Org Biomol Chem 10:440–446 10.1039/C1OB06419C

Yoshikuni Y, Martin VJJ, Ferrin TE, Keasling JD (2006), Engineering cotton (+)-δ-cadinene synthase to an altered function: germacrene D-4-ol synthase. Chem Biol 13:91–98 10.1016/j.chembiol.2005.10.016

Yu Y, Chang P, Yu H, Ren H, Hong D, Li Z, Ying Wang Y, Song H, Huo Y, Li C (2018) Productive amyrin synthases for efficient α-amyrin synthesis in Engineered Saccharomyces cerevisiae ACS Synth Biol 7:2391–2402 https://doi-org.proxy.lnu.se/10.1021/acssynbio.8b00176

Yuan Y, Cheng S, Bian G, Yan P, Ma Z, Dai W, Chen R, Fu S, Huang H, Chi H, Cai Y, Deng Z, Liu T (2022) Efficient exploration of terpenoid biosynthetic gene clusters in filamentous fungi. Nat Catal 5:277–287 10.1038/s41929-022-00762-x

Zare R, Solouki M, Omidi M, Irvani N, Mahdi Nezad N, Rezazadeh Sh (2010) Callus induction and plant regeneration in Ferula assa foetida L. (asafetida), an endangered medicinal plant. Trakia J Sci 8:11–18

Zhai D-D, Zhong J-J (2010) Simultaneous analysis of three bioactive compounds in Artemisia annua hairy root cultures by reversed-phase high performance liquid chromatography–diode array detector. Phytochem Anal 21:524–530 10.1002/pca.1226

Zhang L-L, Chen Y, Li Z-J, Fan G, Li X (2023a) Production, function, and applications of the sesquiterpenes valencene and nootkatone: a comprehensive review. J Agric Food Chem 71:121–142 10.1021/acs.jafc.2c07543

Zhang M, Wang S, Yin J, Li C, Zhan Y, Xiao J, Liang T, Li X (2016) Molecular cloning and promoter analysis of squalene synthase and squalene epoxidase genes from Betula platyphylla. Protoplasma 253:1347–1363 https://doi-org.proxy.lnu.se/10.1007/s00709-015-0893-3

Zhang X, Li S (2017) Expansion of chemical space for natural products by uncommon P450 reactions. Nat Prod Rep 34:1061–1089 10.1039/c7np00028f

Zhang Z, Wu QY, Ge Y, Huang Z-Y, Hong R, Li A, Xu J-H, Yu H-L (2023b) Hydroxylases involved in terpenoid biosynthesis: a review. Bioresour Bioprocess 10:39 10.1186/s40643-023-00656-1

Zhou D, Li N, Zhang Y, Yan C, Jiao K, Sun Y, Ni H, Lin B, Hou Y (2016a) Biotransformation of neuro-inflammation inhibitor kellerin by Angelica sinensis (Oliv.) Diels callus. RSC Adv 6:97302–97312 10.1039/C6RA22502K

Zhou Y, Ma Y, Zeng J, Duan L, Xue X, Wang H, Lin T, Liu Z, Zeng K, Zhong Y, Zhang S, Hu Q, Liu M, Zhang H, Reed J, Moses T, Liu X, Huang P, Qing Z, Liu X, Tu P, Kuang H, Zhang Z, Osbourn A, Ro DK, Shang Y, Huang S (2016b) Convergence and divergence of bitterness biosynthesis and regulation in Cucurbitaceae. Nat Plants 2:16183 10.1038/nplants.2016.183

