## supplementary materials for "Prediction of the reaction mechanisms of sesquiterpene coumarin synthases supports a direct evolutionary link with triterpene biosynthesis"

**Table of contents**

**Plants producing sesquiterpene coumarins**
Table S1 List of plants producing sesquiterpene coumarins from *O*-farnesylated coumarins
**Chemical structures**
Figure S1 Key to the labeling of all structures of sesquiterpene coumarins
Figure S2 Structures of linear sesquiterpene coumarins with carbon skeleton ***O*-A**
Figure S3 Structures of sesquiterpene coumarins with carbon skeletons ***O*-B** to ***O*-D**
Figure S4 Structures of sesquiterpene coumarins with carbon skeletons ***O*-E** to ***O*-G**
Figure S5 Structures of sesquiterpene coumarins with carbon skeleton ***O*-H**
Figure S6 Structures of sesquiterpene with carbon skeleton ***O*-I**
Figure S7 Structures of sesquiterpene coumarins with carbon skeleton ***O*-I**, ***O*-Jb**, ***O*-Jc**, ***O*-K**, and ***O*-L**
Figure S8 Structures of sesquiterpene with carbon skeleton ***O*-Ja**
Figure S9 Structures of sesquiterpene coumarins with carbon skeletons ***O*-M** to ***O*-Q**
Figure S10 Structures of sesquiterpene coumarins with carbon skeletons ***O*-R** and ***O*-S**
Figure S11 Structures of sesquiterpene coumarins with carbon skeletons to ***O*-T** to ***O*-α**
**References**

**Table S1.** List of plants producing sesquiterpene coumarins from *O*-farnesylated coumarins. The coumarin moiety of the different sesquiterpene coumarins are color-coded: umbelliferone; scopoletin or isofraxidin; other coumarins moieties. The structures can be found in Figures S2 to S11. Sesquiterpene coumarins in bold (***O*-U-A1**) are linear and do not require any sesquiterpene coumarin synthase to be produced. The sesquiterpene moiety of sesquiterpene coumarins in blue boxes are produced before coupling to the coumarin. See Figure S1 for sesquiterpene coumarin codes. This is not a complete summary of publications dealing with the isolation and characterization of sesquiterpene coumarins from plants.

| **Species** | **Family** | **Sesquiterpene coumarins** | **Reference** |
|---|---|---|---|
| *Achillea depressa* | Asteraceae | ***O*-I-A1**; *O*-I-Da1; *O*-I-Da2; *O*-I-Ha1; *O*-I-Ha2; *O*-I-Ha3; *O*-I-Ia1; *O*-I- Ia2; *O*-I-Ia7; *O*-I- Ia8; O-I-Rc1 | [66] |
| *Achillea ochroleuca* | Asteraceae | ***O*-I-A1**; *O*-I-Ha3; *O*-I-Ha4<br>***O*-I-A1;** *O*-I-Ha2; *O*-I-Hc1; *O*-I-M1; *O*-I-Ra1; *O*-I-Rc1<br>***O*-I-A1**; ***O*-I-A2; *O*-I-A3;** *O*-I-Ha1; *O*-I-Ha2; *O*-I-Ha5; *O*-I-Ha6; *O*-I-Ia1<br>***O*-I-A1; *O*-I-A2; *O*-I-A3; *O*-I-A4** | [57]<br>[59]<br>[60]<br>[77] |
| *Achillea pseudopectinata* | Asteraceae | ***O*-I-A1**; *O*-I-Ha1; *O*-I-Ia1; *O*-I-Ia2 | [57] |
| *Aegle marmelos* | Rutaceae | *O*-8H-Y1; *O*-8H-Y2 | [8] |
| *Amaranthus retroflexus* | Amaranthaceae | ***O*-U-A1** | [46] |
| *Ammi majus* | Apiaceae | ***O*-U-A1** | [5] |
| *Angelica archangelica* | Apiaceae | ***O*-U-A1**<br>***O*-U-A1** | [136]<br>[139] |
| *Angelica sinensis** | Apiaceae | *O*-U-Ja18; *O*-U-Ja32; *O*-U-Ia33; *O*-U-Ja33; *O*-U-Ja35; *O*-U-Jc1; O-U-Jc2 | [164] |
| *Angelica sylvestris* | Apiaceae | ***O*-U-A1** | [133] |
| *Anethum graveolens* | Apiaceae | ***O*-U-A1** | [142] |
| *Anthemis cretica* | Asteraceae | *O*-I-Cb1<br>***O*-I-A2**; *O*-I-Cb1; *O*-I-Hc1; *O*-I-Ia3; *O*-I-Ia4; *O*-I-Ia11; *O*-I-O1; *O*-I-O2N | [32]<br>[68] |
| *Apium graveolens* | Apiaceae | ***O*-U-A1** | [136] |
| *Artemisia abrotanum* | Asteraceae | ***O*-I-A1**; *O*-I-Ha1 | [57] |
| *Artemisia absinthium* | Asteraceae | ***O*-U-A1** | [135] |
| *Artemisia alba* | Asteraceae | *O*-I-Ia8 | [58] |
| *Artemisia annua* | Asteraceae | *O*-S-Ia2<br>*O*-I-Ha1 | [104]<br>[162] |
| *Artemisia gmelinii* | Asteraceae | ***O*-I-A1**; *O*-I-Ha1 | [57] |
| *Artemisia persica* | Asteraceae | ***O*-I-A1**; *O*-I-Ha1<br>***O*-S-A1**; *O*-S-Ha1 | [57]<br>[65] |
| *Artemisia pontica* | Asteraceae | ***O*-I-A1**; *O*-I-Ha1 | [57] |
| *Artemisia tripartita* | Asteraceae | ***O*-I-A1; *O*-I-A2**; *O*-I-M1 | [59] |
| *Artemisia vestita* | Asteraceae | ***O*-I-A1**; *O*-I-Ha1 | [57] |
| *Artemisia vulgaris* | Asteraceae | ***O*-U-A1** | [49] |
| *Brocchia cinerea* | Asteraceae | ***O*-I-A1; *O*-S-A1**; *O*-I-Ha1; *O*-I-Ha3; *O*-I-Ha4; *O*-I-Ia1; *O*-I-Ia5, *O*-I-Ia6 | [61] |
| *Cicuta virosa* | Apiaceae | ***O*-U-A1** | [146] |
| *Citrus limon* | Rutaceae | ***O*-U-A1** | [138] |
| *Conyza obscura* | Asteraceae | ***O*-S-A1** | [33] |
| *Coriander sativum* | Apiaceae | ***O*-U-A1** | [136] |
| *Euphorbia portlandica* | Euphorbiaceae | ***O*-S-Ia1** | [105] |
| *Ferula arrigoni* | Apiaceae | ***O*-U-A1; *O*-U-A10**; *O*-I-Ia3; *O*-I-Ia7; *O*-I-Ia8; *O*-U-Ja15 | [19] |
| *Ferula assa-foetida* | Apiaceae | *O*-4H-Ia1 | [34] |
| *Ferula assa-foetida* | Apiaceae | *O*-U-Ia1; *O*-U-Ia19; *O*-U-Ia22; *O*-U-Ia24; *O*-U-Ia33; *O*-U-Ja3; *O*-U-Ja8; *O*-U-Rb1 | [1] |

| | | | |
|---|---|---|---|
| | | ***O*-U-A5; *O*-U-A7; *O*-U-A8**; *O*-U-A18; *O*-U-Rb1 | [18] |
| | | *O*-U-Rb1 | [24] |
| | | *O*-U-Ia18 | [25] |
| | | *O*-U-Ea1 | [26] |
| | | *O*-U-Ca1; *O*-U-Ha13 | [27] |
| | | *O*-U-Cb1; *O*-I-Da1 | [35] |
| | | *O*-U-Ia2; *O*-U-Ja6; *O*-U-Ja34 | [52] |
| | | *O*-U-Cb1; *O*-U-Ia25 | [53] |
| | | *O*-U-Ha27; *O*-U-Ia1; *O*-U-Ia2; *O*-U-Ia7; *O*-U-Ia8; *O*-U-Ia20; *O*-U-Ia25; *O*-U-Ia33; *O*-U-Ja4; O-U-O7 | [67] |
| | | ***O*-U-A5; *O*-U-A10; *O*-U-A16; *O*-U-A17; *O*-U-A18**; *O*-U-Ca1; *O*-U-Ca2; *O*-U-Cb1; *O*-U-Da1; *O*-U-Ha1; *O*-U-Ha7; *O*-U-Ha14; *O*-U-Ha21; *O*-U-Ia20; *O*-U-Ia25; *O*-U-O2; *O*-U-O3; *O*-U-O7; *O*-U-Rb1; *O*-U-Rb2 | [100] |
| *Ferula badrakema* | Apiaceae | *O*-U-Ha1 | [126] |
| | | *O*-U-Ha1; *O*-U-Ha2; *O*-U-Ha3; *O*-U-Ha7 *O*-U-Ha10; *O*-U-Ia2; | [72] |
| *Ferula bungeana* | Apiaceae | ***O*-U-A1**; *O*-U-Ga3; *O*-U-Ga4; *O*-U-Ga5; *O*-U-Ha2; *O*-U-Ha3; *O*-U-Ha26; *O*-U-Ia2; *O*-U-Ia3; *O*-U-Ia34; *O*-U-Ia35; *O*-U-Ia36; *O*-U-Ja4; *O*-U-Jb3 | [63] |
| | | ***O*-U-A1**; *O*-U-A25; *O*-U-Ha1; *O*-U-Ha3; *O*-U-Ia20 | [83] |
| *Ferula cocanica* | Apiaceae | ***O*-U-A1**; ***O*-U-B1** | [89] |
| *Ferula conocaula* | Apiaceae | *O*-U-Ha10; *O*-U-Ha11; *O*-U-Ha12; *O*-U-Ha13 | [95] |
| | | *O*-U-Ia16; *O*-U-Ic2 | [96] |
| | | *O*-U-Ja9 | [97] |
| | | *O*-U-Ha18; *O*-U-Ia27 | [98] |
| | | ***O*-U-A1**; *O*-U-Ha7; *O*-U-Ia13; *O*-U-Ia30 | [99] |
| | | *O*-U-Ha1 | [151] |
| | | *O*-U-Ha3 | [152] |
| | | *O*-U-Ha4 | [153] |
| *Ferula drudeana* | Apiaceae | *O*-U-Ha7; *O*-U-Ja3; *O*-U-Ja4 | [14] |
| *Ferula duranii* | Apiaceae | *O*-U-Ha7; *O*-U-Ja3 | [4] |
| *Ferula flabelliloba* | Apiaceae | ***O*-U-A1**; *O*-U-Ca2; *O*-U-Ca3; *O*-U-Da1; *O*-U-Da2; *O*-U-Da4; *O*-U-Ha3; *O*-U-Ha4; *O*-U-Ha7; *O*-U-Ha14 | [73] |
| *Ferula foetida* | Apiaceae | ***O*-U-A9; *O*-U-A10; *O*-U-A18**; *O*-U-Ha21; *O*-U-Ia6 | [3] |
| *Ferula foetidissima* | Apiaceae | *O*-U-Ja19; *O*-U-Ja27 | [93] |
| *Ferula foliosa* | Apiaceae | *O*-U-Ea7 | [79] |
| *Ferula galbaniflua* | Apiaceae | *O*-U-Ha1; *O*-U-Ha7; *O*-U-Ha6; *O*-U-Ha7; *O*-U-Ha10; *O*-U-Ha14; | [44] |
| | | *O*-U-Ha24; *O*-U-Ha25; *O*-U-Ha26; *O*-U-Hd1 | [56] |
| *Ferula gummosa* | Apiaceae | *O*-U-Ha1; *O*-U-Ha7; *O*-U-Ha10; *O*-U-Ha14; *O*-U-Ha18; *O*-U-Ia27; *O*-U-Ia28; *O*-U-Ia29; *O*-U-Ic3; *O*-U-Jb1 | [74] |
| | | *O*-U-Ha1; *O*-U-Ha3; *O*-U-Ha7 | [75] |
| | | *O*-U-Da1; *O*-U-Ia25; *O*-U-Ja32; *O*-U-Ja34 | [7] |
| *Ferula heuffelii* | Apiaceae | *O*-U-Ja4 | [121] |
| *Ferula huber-morathii* | Apiaceae | *O*-U-Ha7; *O*-U-Ja3 | [4] |
| | | *O*-U-Ia25; *O*-U-Ia33; *O*-U-Ja20; *O*-U-Ja30; *O*-U-Ja32; *O*-U-Ja34 | [21] |
| | | *O*-U-Ha1; *O*-U-Ha3; *O*-U-Ha7; *O*-U-Ia3; *O*-U-Ia23; *O*-U-Ia25; *O*-U-Ia26; *O*-U-Ia33; *O*-U-Ja3; *O*-U-Ja4; *O*-U-Ja5; *O*-U-Ja18; *O*-U-Ja20; *O*-U-Ja32; *O*-U-Ja34 | [42] |
| *Ferula karatavica* | Apiaceae | *O*-U-W1 | [22] |
| | | *O*-U-A12 | [118] |

| | | | |
|---|---|---|---|
| | | *O*-U-α1 | [23] |
| *Ferula kokanica* | Apiaceae | *O*-U-Ia20; *O*-U-Ia26; *O*-U-Ia33; *O*-U-Ja1; *O*-U-Ja2; *O*-U-Ja32; *O*-U-Ja34 | [114] |
| *Ferula kopetdaghensis* | Apiaceae | *O*-U-Da1; *O*-U-Ea3; *O*-U-Ea5<br>*O*-U-Da1; *O*-U-F1<br>*O*-U-F2; *O*-U-F3<br>*O*-U-Ja18; *O*-U-Ja20<br>*O*-U-Ea4; *O*-U-Ja7; *O*-U-Ja13 | [80]<br>[81]<br>[112]<br>[113]<br>[115] |
| *Ferula korshinskyi* | Apiaceae | ***O*-U-A22; *O*-U-A26** | [78] |
| *Ferula krylovii* | Apiaceae | *O*-U-Ea2<br>*O*-Uc1; *O*-Uc2 | [154]<br>[155] |
| *Ferula kuhistanica* | Apiaceae | *O*-U-Ga6; *O*-U-Ha7; *O*-U-Ha8; *O*-U-Ia25; *O*-U-Ia41; *O*-U-Ic2; *O*-U-Ja13; *O*-U-Ja28; *O*-U-Ja34; *O*-U-Ja37; *O*-U-Ja38; *O*-U-Ja40; *O*-U-Ja42; *O*-U-O11; *O*-U-Rb5 | [20] |
| *Ferula latialata* | Apiaceae | ***O*-U-A1**; *O*-U-Ha1; *O*-U-Ia1; *O*-U-Ia2; *O*-U-Ia7 | [150] |
| *Ferula lehmannii* | Apiaceae | *O*-U-Ca2; *O*-U-Ia1; *O*-U-Ic3 | [128] |
| *Ferula lithophila (=Peucedanum mogoltavicum)* | Apiaceae | *O*-U-A8; *O*-U-Da1; *O*-U-Ha20 | [88] |
| *Ferula loscosii* | Apiaceae | ***O*-U-A1**; *O*-U-Ia7; *O*-U-Ia8; *O*-U-Ia9 | [125] |
| *Ferula marmarica* | Apiaceae | *O*-4H-Ia2; *O*-4H-Ia3 | [2] |
| *Ferula microcarpa* | Apiaceae | *O*-U-Ea8; *O*-U-O4; *O*-U-O10 | [54] |
| *Ferula microloba* | Apiaceae | *O*-U-Ja14; *O*-U-Ja15; *O*-U-O3; *O*-U-Rb1; *O*-U-Rb2<br>*O*-U-M3 | [116]<br>[117] |
| *Ferula moschata* | Apiaceae | *O*-U-Ha1 | [151] |
| *Ferula mogoltavica* | Apiaceae | *O*-U-Ia33; *O*-U-Ja30; *O*-U-Ja31 | [86] |
| *Ferula narthex* | Apiaceae | *O*-U-Ha1; *O*-U-Ha3; *O*-U-Ic1; *O*-U-N1<br>*O*-U-Ic1; *O*-U-N2<br>*O*-U-Ha25; *O*-U-Ha27; *O*-U-Ic1; *O*-U-N1<br>***O*-U-A9; *O*-U-A10; *O*-U-A16; *O*-U-A18**; *O*-U-Ha7; *O*-U-Ha14 | [29]<br>[11]<br>[12 ]<br>[15] |
| *Ferula nevskyi* | Apiaceae | *O*-U-Ja1; *O*-U-Ja7; *O*-U-Ja29 | [132] |
| *Ferula pallida* | Apiaceae | *O*-U-Ha1; *O*-U-Ha7 | [140] |
| *Ferula penninervis* | Apiaceae | *O*-U-O4; *O*-U-O6 | [120] |
| *Ferula persica* | Apiaceae | *O*-U-Da1; *O*-U-Ia3; *O*-U-Ia25; *O*-U-Ia26; *O*-U-Ia33<br>***O*-U-A23; *O*-U-A24**; *O*-U-Ia21; *O*-U-Ja26 | [69]<br>[71] |
| *Ferula polyantha* | Apiaceae | *O*-U-Ia19; *O*-U-Ia20<br>*O*-U-Ea8; *O*-U-Ea9; *O*-U-Ea10; *O*-U-Ha17<br>*O*-U-Ea8; *O*-U-Ea9; *O*-U-Ea10; *O*-U-Ha17 | [85]<br>[87]<br>[131] |
| *Ferula pseudalliaceae* | Apiaceae | *O*-U-Da1; *O*-U-O8; *O*-U-Rb2; *O*-U-Rb3; *O*-U-Rb4; *O*-U-Rc2<br>*O*-U-Cb1; *O*-U-Da1; *O*-U-Da4; *O*-U-O7; *O*-U-O9; *O*-U-O11 | [39]<br>[40] |
| *Ferula pseudooreoselium* | Apiaceae | *O*-U-Ja4; *O*-U-Ja15 | [91] |
| *Ferula samarcandica* | Apiaceae | *O*-U-Ja3<br>*O*-U-Ga1; *O*-U-Ha1; *O*-U-Ia1; *O*-U-Ia7; *O*-U-Ia28; *O*-U-Ia33; *O*-U-Ja1; *O*-U-Ja3; *O*-U-Ja4; *O*-U-Ja7; *O*-U-Ja9; *O*-U-Ja10; *O*-U-Ja11; *O*-U-Ja12; *O*-U-Ja13; *O*-U-Ja14; *O*-U-Ja18; *O*-U-Ja21; *O*-U-Ja22; *O*-U-Ja23; *O*-U-Ja24; *O*-U-Ja25<br>*O*-U-Ia25; *O*-U-Ia33; *O*-U-Ja3; *O*-U-Ja5<br>*O*-U-Da1; *O*-U-Ga2; *O*-U-Ga5; *O*-U-Ga7; *O*-U-Ha3; *O*-U-Ia3; *O*-U-Ia37; *O*-U-Ja2; *O*-U-Ja4 | [130]<br>[82]<br>[90]<br>[163] |

| | | | |
|---|---|---|---|
| *Ferula schtschurovskyana* | Apiaceae | *O*-U-Ja1; *O*-U-Ja7 | [132] |
| *Ferula seravschanica* | Apiaceae | **_O_-U-A1; _O_-U-A4; _O_-U-A6; _O_-U-A9; _O_-U-A10; _O_-U-A18**; *O*-U-Gb1; *O*-U-Ha1; *O*-U-Ha2; *O*-U-Ha3; *O*-U-Ha5; *O*-U-Ha7; *O*-U-Ha12; *O*-U-Ha15; *O*-U-Hb1; *O*-U-Ia7; *O*-U-Ia13; *O*-U-Ic2; *O*-U-Ja36; *O*-U-Jb1; *O*-U-Jb2 | [137] |
| *Ferula sinaica* | Apiaceae | **_O_-U-A1; _O_-U-A10; _O_-U-A15;** *O*-U-Ja3; *O*-U-Ja5; *O*-U-Ja13<br>*O*-U-Ia39; *O*-U-Ib1<br>*O*-U-Ja6; *O*-U-V | [9]<br>[13]<br>[41] |
| *Ferula sinkiangensis* | Apiaceae | **_O_-U-A1; _O_-U-A3**; *O*-U-Cb1; *O*-U-Da2; *O*-U-Eb1; *O*-U-Ia8; *O*-U-Ia20; *O*-U-Ia26; *O*-U-163; *O*-U-O2; *O*-U-O5; *O*-U-Rc1; *O*-U-Rc2; *O*-U-Rc3; *O*-U-Rc4; *O*-U-S1; *O*-U-S2; *O*-U-S3; *O*-U-S4; *O*-U-X1<br>**_O_-U-A1**; *O*-U-Cb1; *O*-U-F3; *O*-U-Ha7; *O*-U-Ia7; *O*-U-Ja14; *O*-U-L1; *O*-U-O1; *O*-U-O2; *O*-U-Rc1; *O*-U-U1<br>*O*-U-K1; *O*-U-K2<br>*O*-U-Q1<br>**_O_-U-A1; _O_-U-A10; _O_-U-A11;** *O*-U-Cb1; *O*-U-Da1; *O*-U-Rb1<br>**_O_-U-A1; _O_-U-A10; _O_-U-A12**; *O*-U-Cb1; *O*-U-Da1; *O*-U-Ea2; *O*-U-Ec1; *O*-U-Ha10; *O*-U-Ha22; *O*-U-Ha23; *O*-U-Ia20; *O*-U-Ia22; *O*-U-Ia24; *O*-U-Ia25; O-U-Ia26; *O*-U-Ia31; *O*-U-Ia33; *O*-U-Ja18; *O*-U-Ja19; *O*-U-Ja28; *O*-U-Ja29; *O*-U-Ja32; *O*-U-Ja34; *O*-U-O1; *O*-U-O6; *O*-U-Rb1; *O*-U-Rc1; *O*-U-Rc2<br>*O*-U-Cb1; *O*-U-Da2; *O*-U-Ga5; *O*-U-Ga7; *O*-U-Ha7; *O*-U-Ha14; *O*-U-Ia3; *O*-U-Ia7; *O*-U-Ia13;*O*-U-Ia25; *O*-U-Ic3; *O*-U-Ja5; *O*-U-Ja13; *O*-U-Ja14; *O*-U-Ja17; *O*-U-Ja18; *O*-U-Jb1; *O*-U-Jb4; *O*-U-L1; *O*-U-O1; *O*-U-O6; *O*-U-Rc1; *O*-U-Rc5; *O*-U-Rc6; *O*-U-Rc7; *O*-U-Rc8; *O*-U-Rc9; *O*-U-X1<br>**_O_-U-A1; _O_-U-A10**; *O*-U-Cb1; *O*-U-Da1; *O*-U-Ea6; *O*-U-Ia20; *O*-U-Ia25; *O*-U-Ia26; *O*-U-Ia33; *O*-*U*-Ja18; *O*-U-Ja19; *O*-U-Ja32; *O*-U-Ja34; *O*-U-M1; *O*-U-Rb1; *O*-U-Rb2<br>*O*-U-L1; *O*-U-O1; *O*-U-O2<br>**_O_-U-A27**; *O*-U-Ga7; *O*-U-Ga9; *O*-U-Ia23; *O*-U-Ia39; *O*-U-Ja39; *O*-U-L2; *O*-U-L3; *O*-U-Rc10 | [62]<br>[101]<br>[102]<br>[103]<br>[145]<br>[157]<br>[158]<br>[159]<br>[160]<br>[37] |
| *Ferula sumbul* | Apiaceae | *O*-U-Ha1; *O*-U-Ha3; *O*-U-Ha7; *O*-U-Ia32 | [84] |
| *Ferula szowitsiana* | Apiaceae | **_O_-U-A1**; *O*-U-Cb1; *O*-U-Da1; *O*-U-M2; *O*-U-O11; *O*-U-Rb1; *O*-U-Rb2<br>**_O_-U-A1**; *O*-U-M2; *O*-U-Rb1; *O*-U-Rb2 | [70]<br>[150] |
| *Ferula tadshikorum* | Apiaceae | *O*-U-Ha6<br>**_O_-U-A8; _O_-U-A24** | [92]<br>[123] |
| *Ferula tenuissima* | Apiaceae | *O*-U-Ha7; *O*-U-Ja3 | [4] |
| *Ferula teterrima* | Apiaceae | *O*-U-Ia13<br>*O*-U-Ia15 | [124]<br>[160] |
| *Ferula tingitana* | Apiaceae | *O*-U-Ha7; *O*-U-Ja15 | [108] |
| *Ferula tunetana* | Apiaceae | **_O_-U-A1**; *O*-U-Ha9; *O*-U-Ia7; *O*-U-Ia8; *O*-U-Ja13; *O*-U-Ja16 | [76] |
| *Ferula turcica* | Apiaceae | **_O_-U-A1**; *O*-U-Ia33; *O*-U-Ja34<br>*O*-U-A21; *O*-U-Ga6; *O*-U-Ga7; *O*-U-Ga8; *O*-U-Ga9; *O*-U-Ra1 | [150]<br>[43] |
| *Ferula vesceritensis* | Apiaceae | *O*-U-Ha9; *O*-U-Ia7; *O*-U-Ia8; *O*-U-Ia10; *O*-U-Ja5; *O*-U-T1<br>*O*-U-Ha7; *O*-U-Ia7<br>*O*-U-Ha7; *O*-U-Ia25 | [10]<br>[110]<br>[119] |
| *Ferulago campestris* | Apiaceae | **_O_-U-A1** | [50] |
| *Haplophyllum patavinum* | Rutaceae | **_O_-U-A1** | [45] |
| *Heptaptera anatolica* | Apiaceae | **_O_-5DH-A2** | [16] |

| | | | |
|---|---|---|---|
| *Heptaptera anatolica* | Apiaceae | ***O*-U-A14**; *O*-U-Ha1; *O*-U-Ha3; *O*-U-Ha7 *O*-U-Ja3; *O*-U-Ja5<br>***O*-U-A1; *O*-U-A10**; *O*-U-Ia1; *O*-U-Ia3; *O*-U-Ia5; *O*-U-Ia7; *O*-U-Ia11; *O*-U-Ia12 | [16]<br>[147] |
| *Heptaptera anisoptera* | Apiaceae | ***O*-5DH-A1; *O*-5DH-A2; *O*-5DH-A3** | [16] |
| *Heptaptera anisoptera* | Apiaceae | ***O*-U-A14**; *O*-U-Ha1; *O*-U-Ha3; *O*-U-Ha7; O-U-Ja3; *O*-U-Ja5<br>***O*-U-A1**; *O*-U-Ia3; *O*-U-Ia5; *O*-U-Ia7; *O*-U-Ia11; *O*-U-Ia12; *O*-U-Ia11 | [16]<br>[17] |
| *Heptaptera cilicica* | Apiaceae | ***O*-U-A1; *O*-U-A10**; *O*-U-Ia1; *O*-U-Ia3; *O*-U-Ia4; *O*-U-Ia5; *O*-U-Ia6; *O*-U-Ia7; *O*-U-Ia11; *O*-U-Ia12; *O*-U-Ia16<br>***O*-U-A1; *O*-U-A2**; *O*-U-Ha1; *O*-U-Ha3; *O*-U-Ha7<br>***O*-U-A1; *O*-U-A10**; *O*-U-Ia1; *O*-U-Ia3; *O*-U-Ia5; *O*-U-Ia6; *O*-U-Ia7; *O*-U-Ia12 | [148]<br>[64]<br>[109] |
| *Heptaptera (Colladonia) triquetra* | Apiaceae | *O*-U-Ia7; *O*-U-Ia8<br>*O*-U-Ia7<br>***O*-U-A1; *O*-U-A10**; *O*-U-Ia2; *O*-U-Ia7<br>***O*-U-A1; *O*-U-A10**; *O*-U-Ia4: *O*-U-Ia7: *O*-U-Ia8; | [28]<br>[129]<br>[36]<br>[149] |
| *Heraclum yunngningense* | Apiaceae | ***O*-U-A1** | [143] |
| *Jatropha integerrima* | Euphorbiaceae | *O*-F-Z1 | [141] |
| *Ligularia persica* | Asteraceae | *O*-U-Ec2; *O*-U-Ha10; *O*-U-Ha14; *O-U*-Ia1; *O*-U-Ia3; *O*-U-Ia7: *O*-U-Ia8; *O*-U-Ia19; *O*-U-Ia19; *O*-U-Ia26; *O*-U-Ia33; *O*-U-Ja3; *O*-U-Ja13 | [106] |
| *Apium graveolens* | Solanaceae | ***O*-U-A1** | [48] |
| *Magydaris pastinacea* | Apiaceae | ***O*-U-A1** | [51] |
| *Magydaris tomentosa* | Apiaceae | ***O*-U-A1** | [127] |
| *Melaleuca alternifolia* | Myrtaceae | ***O*-U-A1** | [134] |
| *Nassauvia argentea* | Asteraceae | *O*-HM-Cb1; *O*-HM-Ia2 | [30] |
| *Nassauvia revoluta* | Asteraceae | *O*-HM-D1; *O*-HM-D2; *O*-HM-Ia1 | [31] |
| *Peucedanum palustre* | Apiaceae | ***O*-U-A1** | [161] |
| *Peucedanum zenkeri* | Apiaceae | ***O*-U-A1** | [107] |
| *Pimpinella anisum* | Apiaceae | ***O*-U-A1** | [142] |
| *Prangos latiloba* | Apiaceae | *O*-U-Da3 | [6] |
| *Punica granatum* | Lythraceae | ***O*-U-A1** | [47] |
| *Scabiosa comosa* | Caprefoliaceae | ***O*-U-A1** | [38] |
| *Scutellaria baicalensis* | Lamiaceae | ***O*-U-A1; *O*-U-A10** | [111] |
| *Seseli annuum* | Apiaceae | ***O*-U-A1** | [156] |
| *Spinacia oleracea* | Amaranthaceae | ***O*-U-A1** | [48] |
| *Tanacetum heterotumum* | Asteraceae | *O*-I-Ha2; *O*-I-Ha7 | [55] |
| *Tanacetum parthenium* | Asteraceae | *O*-I-Ia9 | [94] |

* Kellerin was fed to a callus culture of *Angelica sinensis*. Biotransformation of kellerin to a number sesquiterpene coumarins was confirmed. Seven sesquiterpene coumarins shown in red were isolated from the callus culture.

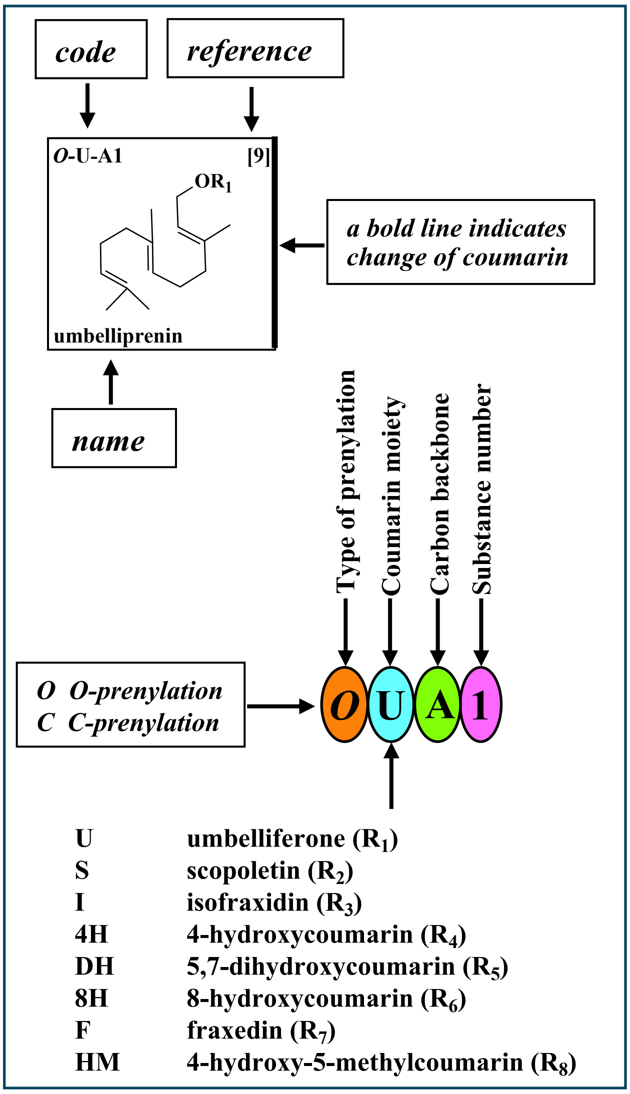


**Figure S1.** Key to the labeling of all structures of sesquiterpene coumarins in the Supplementary Figures. This label is found in the upper left corner of each box. In a few cases we suggest alternative names (in blue text) of some sesquiterpene coumarins. The names given in the publications are in red text.

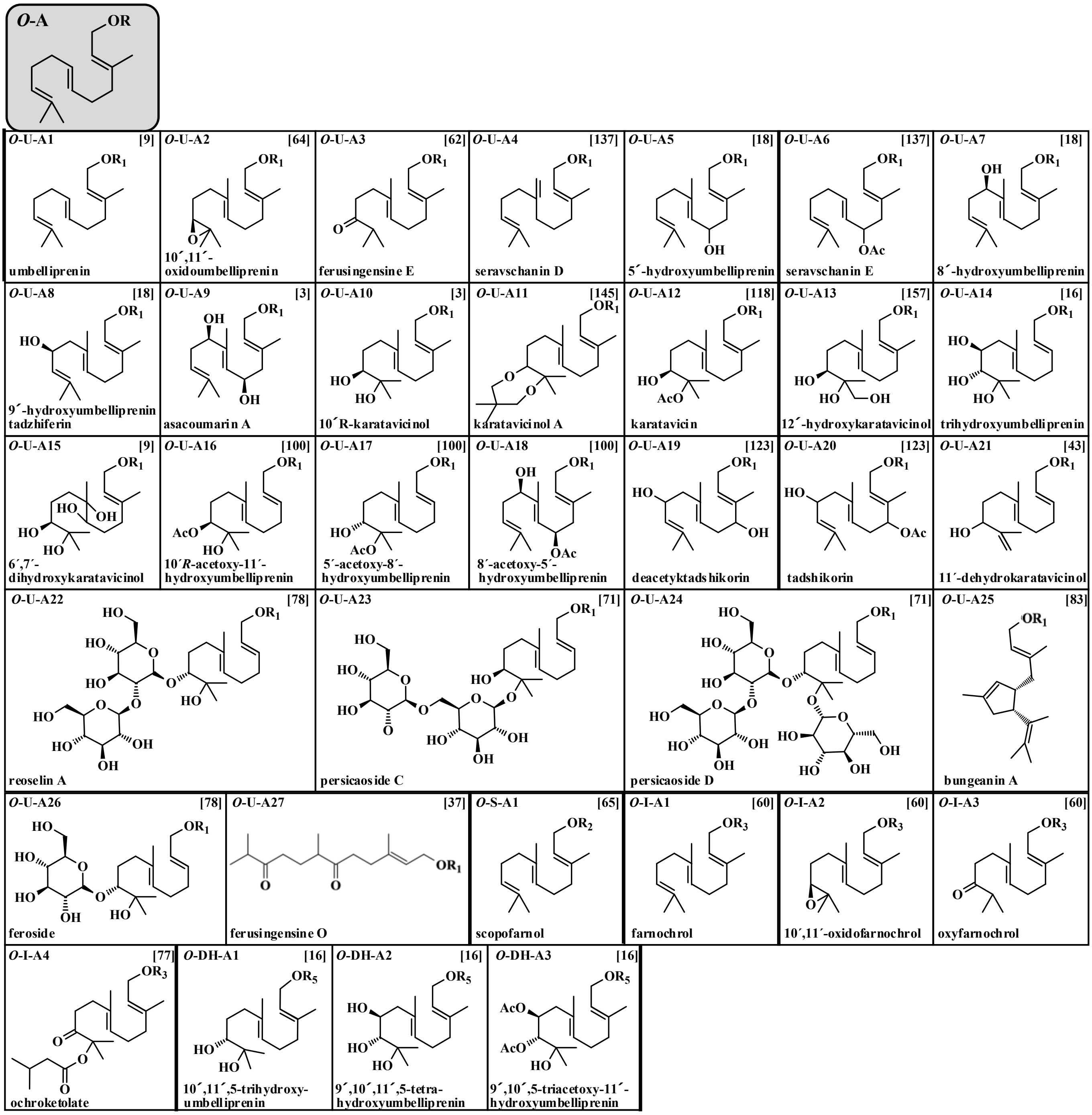


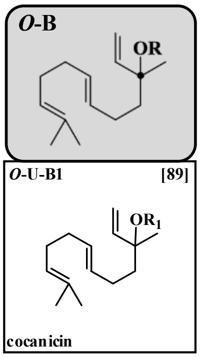


**Figure S2.** Structures of linear sesquiterpene coumarins with carbon skeleton ***O*-A** and ***O*-B.** $R_1$=umbelliferone (U) (**9**); $R_2$=scopoletin (S) (**10**); $R_3$=isofraxidin (I) (**12**); $R_5$=5,7-dihydroxycoumarin (DH) (**17**).

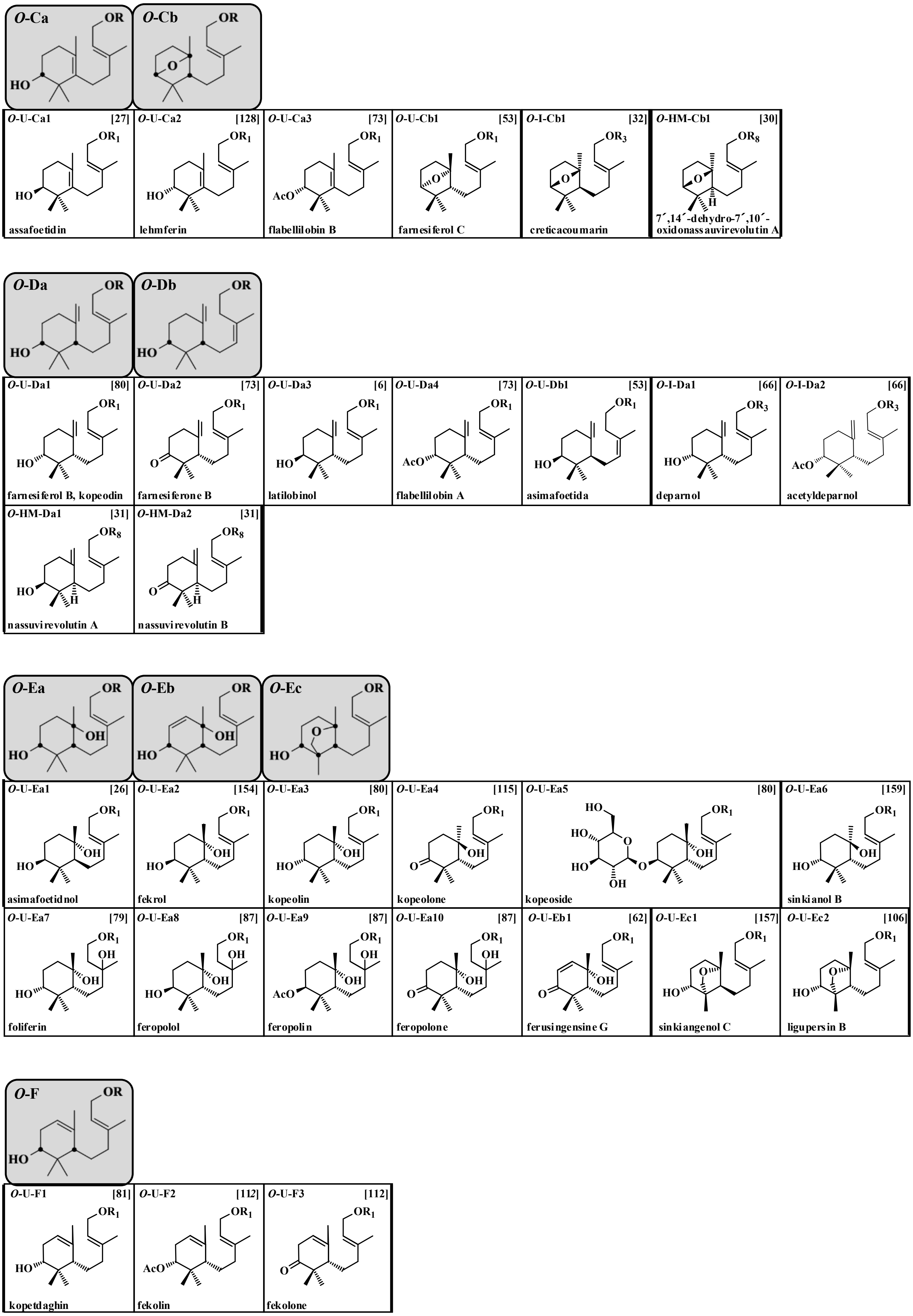


**Figure S3.** Structures of sesquiterpene coumarins with carbon skeletons ***O*-C** to ***O*-F.** $R_1$=umbelliferone (U) (**9**); $R_3$=isofraxidin (I) (**12**); $R_5$=4-hydroxy-5-methylcoumarin (DH) (**17**).

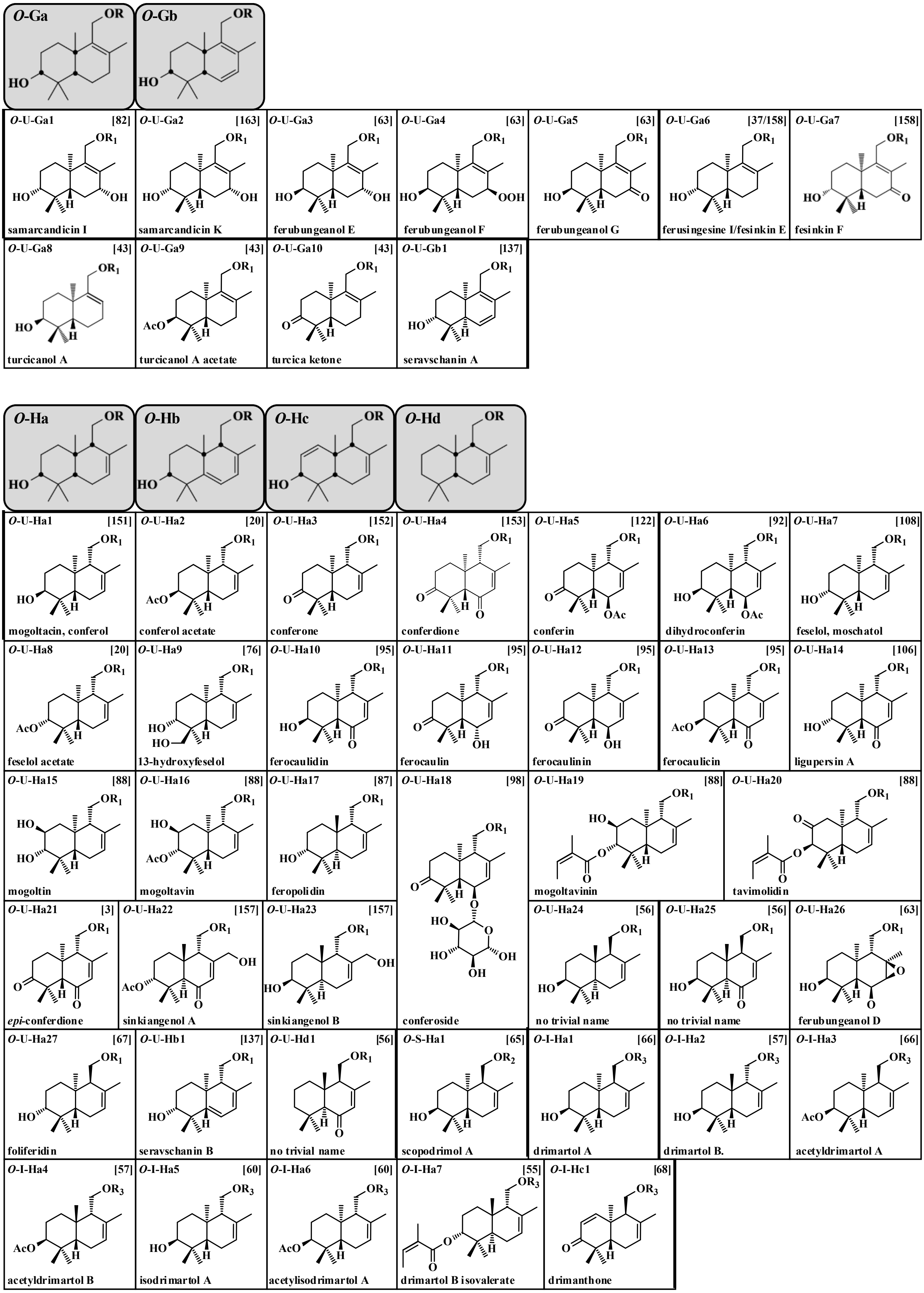


**Figure S4.** Structures of sesquiterpene coumarins with carbon skeleton ***O*-G** and ***O*-H.** $R_1$=umbelliferone (U) (**9**); $R_2$=scopoletin (S) (**10**); $R_3$=isofraxidin (I) (**12**).

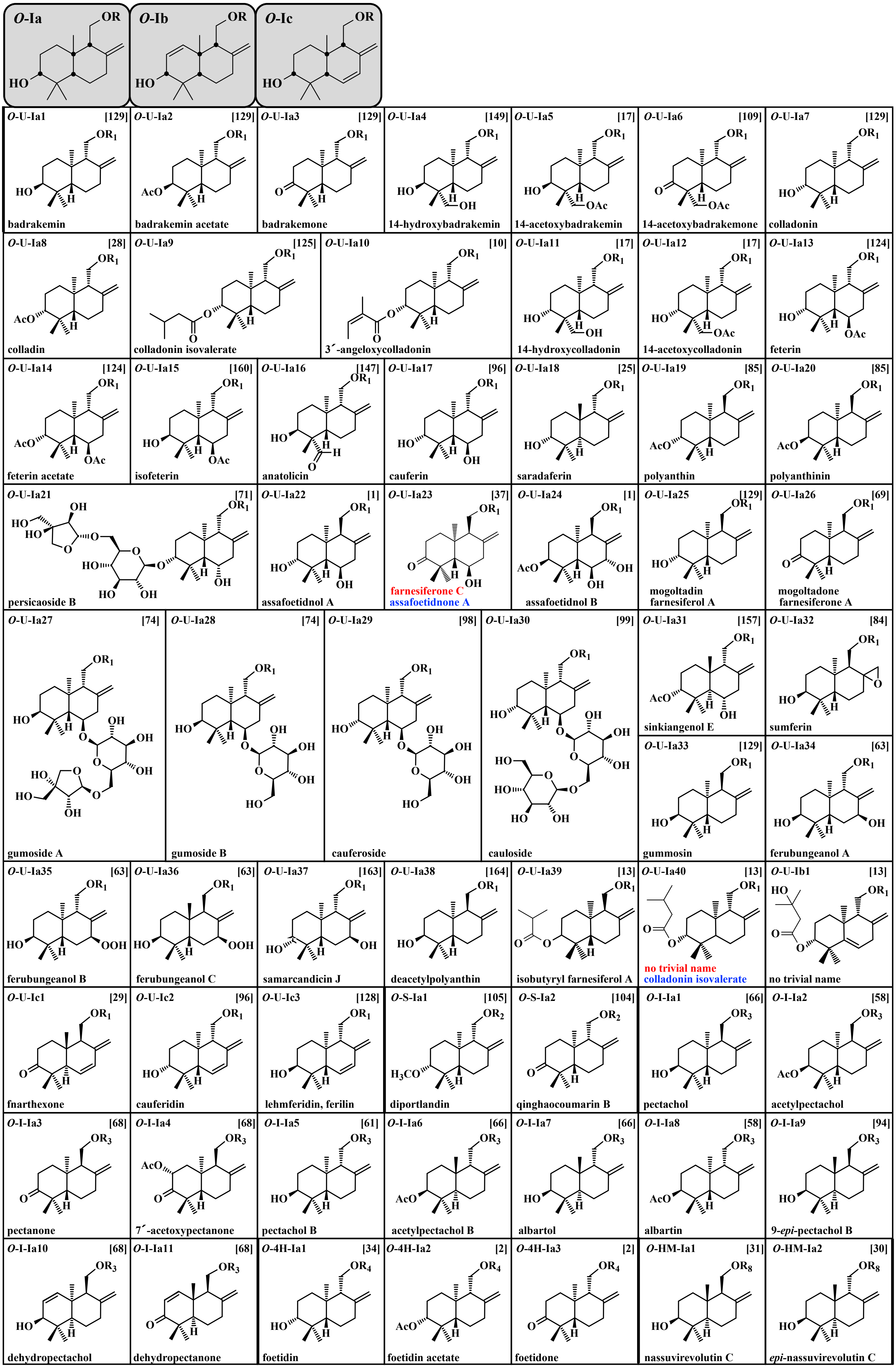




**Figure S5**. Structures of sesquiterpene with carbon skeleton ***O*-I.** $R_1$=umbelliferone (U) (**9**), $R_2$=scopoletin (S) (**10**); $R_3$=isofraxidin (I) (**12**); $R_4$=4-hydroxycoumarin (4H) (**14**); $R_8$=4-hydroxy-5-methylcoumarin (HM) (**18**).

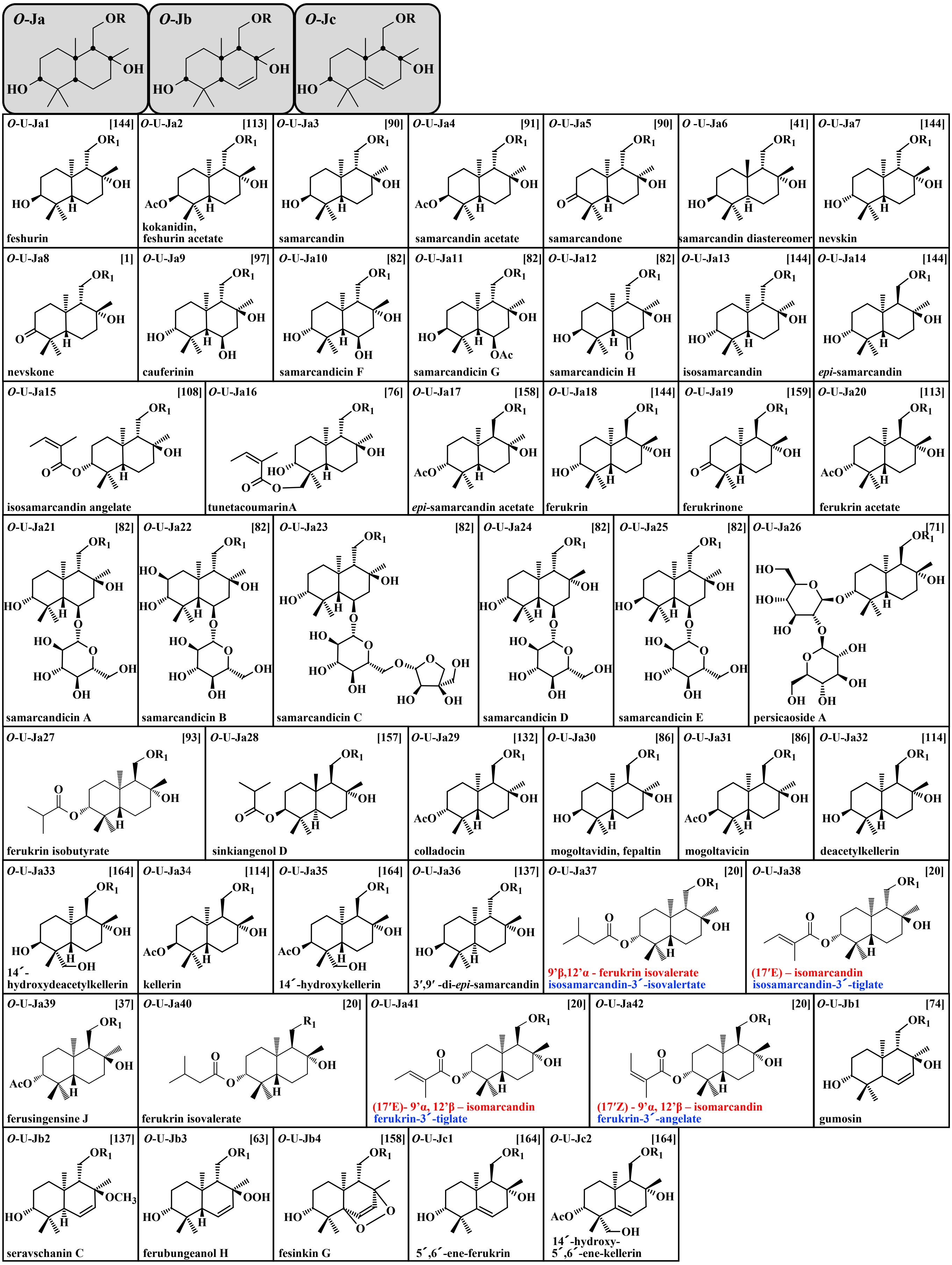


**Figure S6**. Structures of sesquiterpene with carbon skeleton ***O*-J.** $R_1$=umbelliferone (U) (**9**).

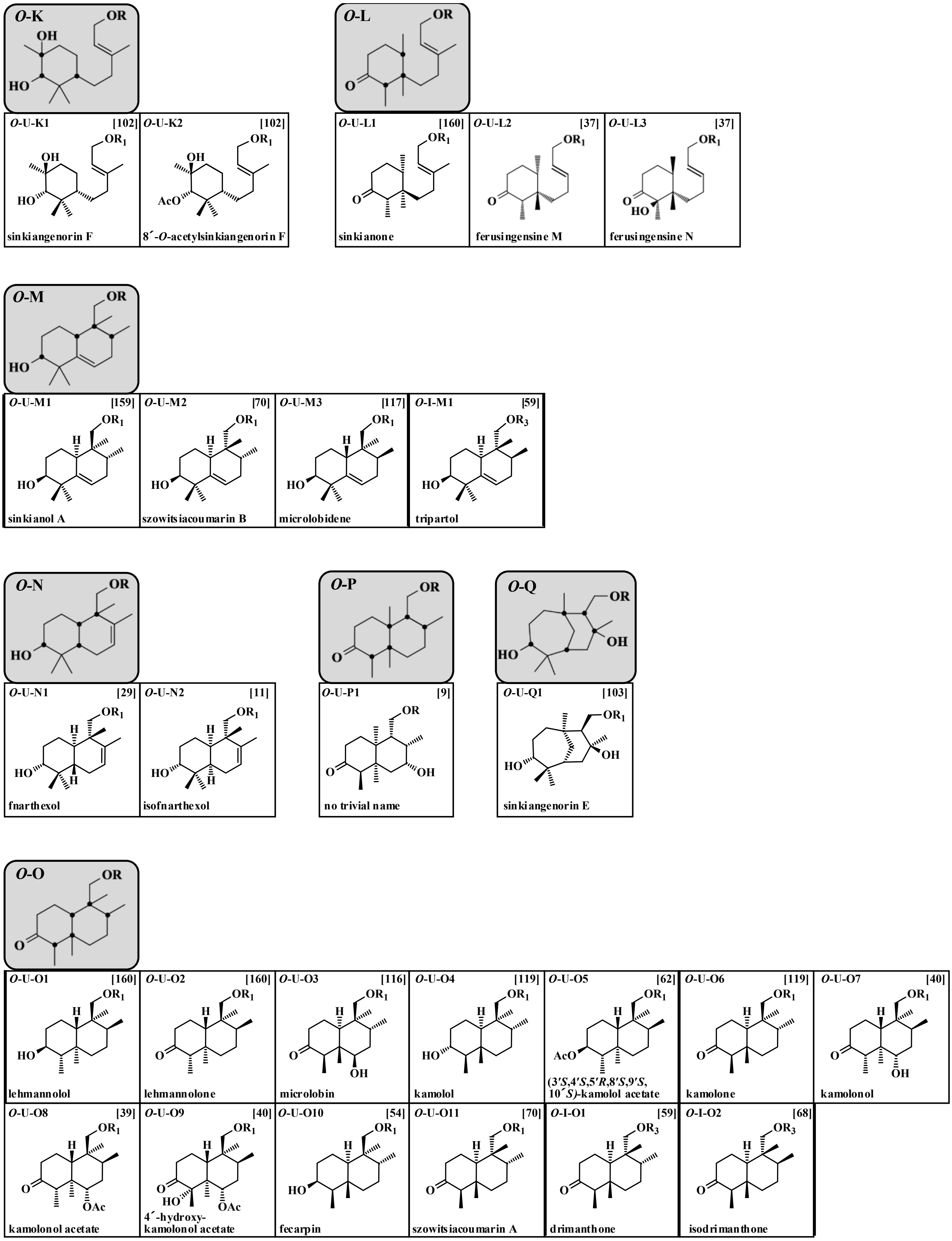


**Figure S7.** Structures of sesquiterpene coumarins with carbon skeleton ***O*-K** to ***O*-O.** $R_1$=umbelliferone (U) (**9**); $R_3$=isofraxidin (I) (**12**).

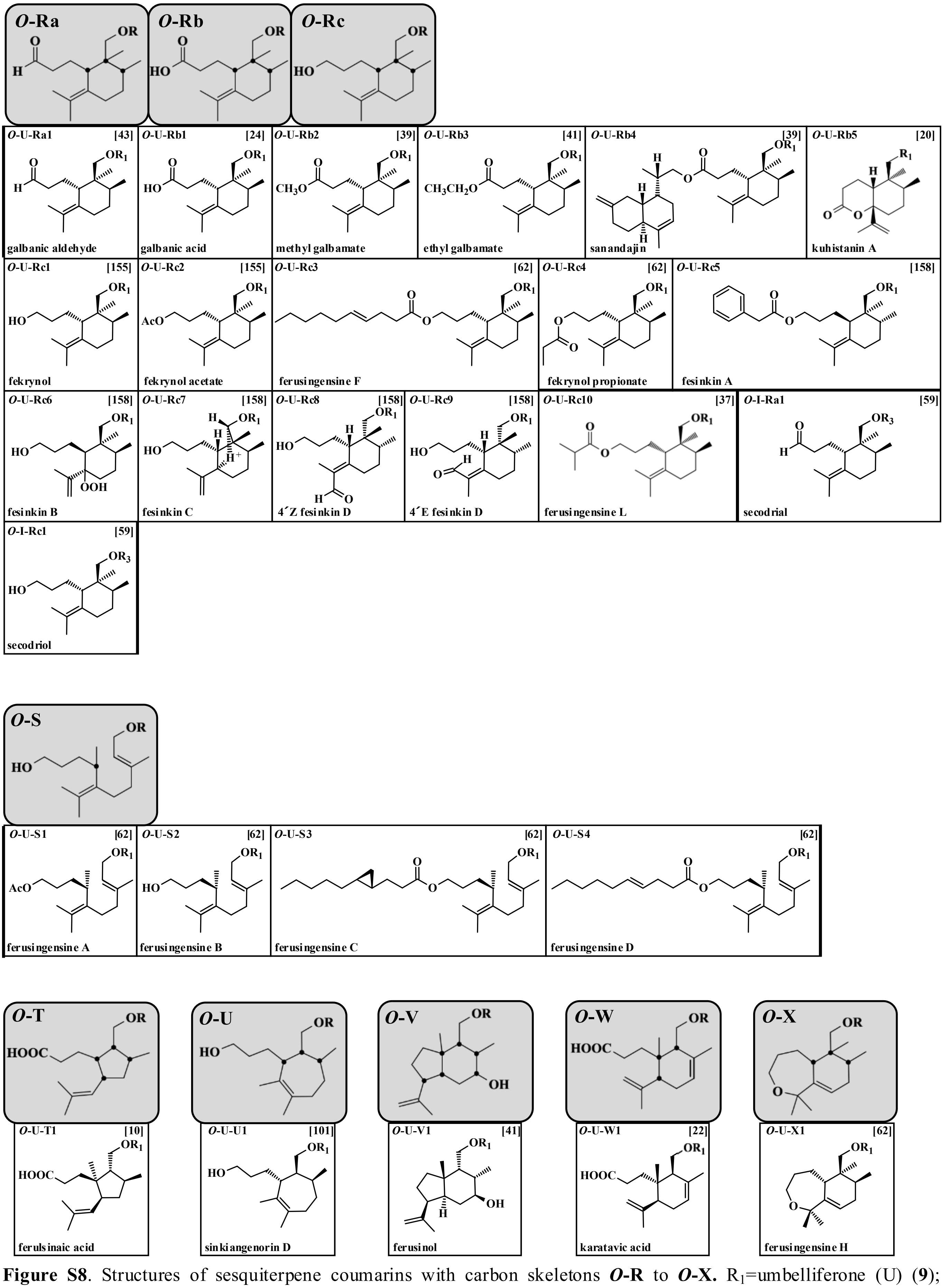


**Figure S8**. Structures of sesquiterpene coumarins with carbon skeletons ***O*-R** to ***O*-X.** $R_1$=umbelliferone (U) (**9**); $R_3$=isofraxidin (I) (**12**).

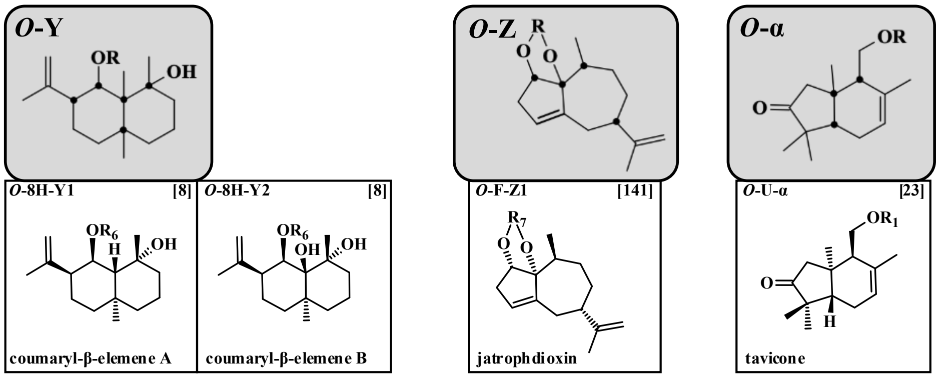


**Figure S9**. Structures of sesquiterpene coumarins with carbon skeletons ***O*-Y** to ***O*-α.** $R_1$=umbelliferone (U) (**9**); $R_6$= 8-hydroxycoumarin (8H) (**13**); $R_7$=fraxidin (F) (**11**).

**References supplementary material**


1. Abd El-Razek MH, Ohta S, Ahmed AA, Hirata T (2001) Sesquiterpene coumarins from the roots of *Ferula assa-foetida.* Phytochemistry 58:1289-1295 https://doi.org/10.1016/s0031-9422(01)00324-7
2. Abd El-Razek MH, Chang F-R, Liaw C-C, Nassar MI, Huang H-C, Chen Y-H, Yang Y-L, Wu Y-C (2004) Two sesquiterpene-coumarins from the roots of *Ferula marmarica.* Heterocycles 63:2101-2109 https://doi.org/10.3987/COM-04-10157
3. Abd El-Razek MH, Wu Y-Ch, Chang F-R (2007) Sesquiterpene coumarins from *Ferula foetida*. J Chin Chem Soc 54:235-238 https://doi.org/10.1002/jccs.200700035
4. Abdel-Kader MS, Alqarni MH, Baykan S, Oztürk B, Salkini MAA Yusufoglu HS, Alam P, Foudah AI (2022) Ecofriendly validated RP-HPTLC method for simultaneous determination of the bioactive sesquiterpene coumarins feselol and samarcandin in five *Ferula* species using green olvents. Separations 9:206 https://doi.org/10.3390/separations9080206
5. Abu-Mustafa EA, El-Bay FKA, Fayez MBE (1971) Natural coumarins XII: Umbelliprenin, a constituent of *Ammi majus* L fruits. J Pharm Sci 60:788-789 https://doi.org/10.1002/jps.2600600528
6. Abyshev AZ (1979) Latilobinol - a new terpenoid coumarin from *Prangos latiloba.* Chem Nat Compd 15:75 https://doi-org.proxy.lnu.se/10.1007/BF00570861
7. Adhami H-R, Fitz V, Lubich A, Kaehlig H, Zehl M, Krenn L (2014) Acetylcholinesterase inhibitors from galbanum, the oleo gum-resin of *Ferula gummosa* Boiss. Phytochem Lett 10:lxxxii- lxxxvii https://doi.org/10.1016/j.phytol.2014.08.023
8. Ahmad MZ, Ali M, Showkat R, Mir SR (2012) New sesqui- and diterpenic coumarin ethers from the roots of *Aegle marmelos* (L.) Corr. Nat Prod J 2:252-258 https://doi.org/10.2174/2210315511202040252
9. Ahmed AA (1999) Sesquiterpene coumarins and sesquiterpenes from *Ferula sinaica*. Phytochemistry 50:109-112 https://doi.org/10.1016/S0031-9422(98)00489-0
10. Ahmed AA, Hegazy M-EF, Zellagui A, Rhouati S, Mohamed TA, Sayed AA, Abdella MA, Ohta S, Hirata T (2007) Ferulsinaic acid, a sesquiterpene coumarin with a rare carbon skeleton from *Ferula* species. Phytochemistry 68:680-686 https://doi.org/10.1016/j.phytochem.2006.12.011
11. Alam M, Khan A, Wadood A, Khan A, Bashir S, Aman A, Jan AK, Rauf A, Ahmad B, Khan AR, Farooq U (2016) Bioassay-guided isolation of sesquiterpene coumarins from *Ferula narthex* Bioss: A new anticancer agent. Front Pharmacol 7:26 https://doi.org/10.3389/fphar.2016.00026
12. Alam M, Khan A, Wadood A, Bashir S, Aman A, Farooq U, Khan FA, Mabood F, Hussain J, Samiullah, Al-Harrasi A (2019) Bioassay-guided isolation of urease inhibitors from *Ferula narthex* Bioss. South Afr J Bot 120:247-252 https://doi.org/10.1016/j.sajb.2018.07.011
13. Al-Hazimi HMG (1986) Terpenoids and a coumarin from *Ferula sinaica.* Phytochemistry 25:2417-2419 https://doi.org/10.1016/S0031-9422(00)81710-0
14. Alqarni MH, Soliman GA, Salkini MAA, Alam P, Yusufoglu HS, Baykan S, Oztürk B, Abdel-Kader MS (2020) The potential aphrodisiac effect of *Ferula drudeana* Korovin extracts and isolated sesquiterpene coumarins in male Rats. Pharmacog Mag 16:404-409 https://doi.org/10.4103/pm.pm_551_19
15. Amin A, Tuenter E, Cos P, Maes L, Exarchou V, Apers S, Pieters L (2016) Antiprotozoal and antiglycation activities of sesquiterpene coumarins from *Ferula narthex* exudate. Molecules 21:1287 https://doi.org/10.3390/molecules21101287
16. Appendino G, Ozen HC, Nano GM, Cisero M (1992a) Sesquiterpene coumarin ethers from the genus *Heptaptera.* Phytochemistry 31:4223–4226 https://doi.org/10.1016/0031-9422(92)80447-m
17. Appendino G, Ozen HC, Tagliapietra S, Cisero M (1992b) Coumarins from *Heptaptera anisoptera*. Phytochemistry 31:3211-3213 https://doi.org/10.1016/0031-9422(92)83477-G
18. Appendino G, Tagliapietra S, Nano GM, Jakupovic J (1994) Sesquiterpene coumarin ethers from Asafetida. Phytochemistry 35:183-186 https://doi.org/10.1016/S0031-9422(00)90530-2
19. Appendino G, Jakupovic J, Alloatti S, Ballero M (1997) Daucane esters from *Ferula arrigonii*, Phytochemistry 45:1639-1643 https://doi.org/10.1016/S0031-9422(97)00250-1
20. Ashurov K, Numonov S, Guoruoluo Y, Aisa HA, Turak A (2024) Unveiling the anti-vitiligo, anti-inflammatory, and antitumor activities of sesquiterpene coumarins isolated from *Ferula kuhistanica*. Fitoterapia 176:106035 https://doi.org/10.1016/j.fitote.2024.106035
21. Aydogan F, Baykan S, Soliman GA, Yusufoglu H, Bedir E (2020) Evaluation of the potential aphrodisiac activity of sesquiterpenoids from roots of *Ferula huber-morathii* Peşmen in male rats. J Ethnopharmcol 257:112868 https://doi.org/10.1016/j.jep.2020.112868
22. Bagirov VYu, Sheichenko VI (1975) The structure of karatavic acid. Chem Nat Compd 11:734-736 https://doi.org/10.1007/bf00568456
23. Bagirov VYu, Sheichenko VI (1976) The structure and stereochemistry of tavicone. Chem Nat Compd 12:399–401 https://doi.org/10.1007/BF00564795
24. Bagirov VYu, Sheichenko VI, Veselovskaya NV, Sklyar YE, Savina AA, Kir'yanova IA (1980) Structure and stereochemistry of galbanic acid. Chem Nat Compd 16:439-441 https://doi.org/10.1007/BF00571032

25. Bandyopadhyay D, Basak B, Chatterjee A, Lai TK, Banerji A, Banerji J, Neuman A, Prangé T (2006) Saradaferin, a new sesquiterpenoid coumarin from *Ferula assafoetida.* Nat Prod Res 20:961-965 https://doi.org/10.1080/14786410600823433
26. Bandyopadhyay D, Banerje M, Laskar S, Basak B (2011) Asimafoetidnol: a new sesquiterpenoid coumarin from the gum resin of *Ferula assa-foetida.* Nat Prod Comm 6:209-212 https://doi.org/10.1177/1934578X1100600213
27. Banerji A, Mallick B, Chatterjee A, Budzikiewicz H, Breuer M (1988) Assafoetidin and ferocolicin, two sesquiterpenoid coumarins from *Ferula assafoetida* Regel. Tetrahedron Lett 29:1557-1560 https://doi.org/10.1016/S0040-4039(00)80351-2
28. Ban'kovskii AI, Ermatov NE, Perel'son ME, Bubeva-Ivanova L, Pavlova NS (1970) Structure of the coumarins colladin and colladonin II. Nat Prod Res 6:170-176 https://doi.org/10.1007/bf00941672
29. Bashir S, Alam M, Adhikari A, (Swagat) Shrestha RL, Yousuf S, Ahmad B, Parveen S, Aman A, Choudhary MI (2014) New antileishmanial sesquiterpene coumarins from *Ferula narthex* Boiss. Phytochem Lett 9:46-50. https://doi.org/10.1016/j.phytol.2014.04.009
30. Bittner M, Jakupovic J, Bohlmann F, Silva M (1988a) Coumarins and guaianolides from further Chilean representatives of the subtribe Nassauviinae. Phytochemistry 27:2867-2868 https://doi.org/10.1016/S0031-9422(00)98112-3
31. Bittner M, Jakupovic J, Bohlmann F, Silva M (1988b) 5-Methylcoumarins from *Nassauvia* species. Phytochemistry 27:3845-3847 https://doi.org/10.1016/0031-9422(88)83029-2
32. Bohlmann F, Zdero C (1975) Natürlich vorkommende Terpen-Derivate, XLVIII. Über neue Inhaltsstoffe der Gattung *Anthemis.* Chem Ber 108:1902-1910 https://doi.org/10.1002/cber.19751080613
33. Bohlmann F, Jakupovic J (1979) 8-Oxo-α-selinen und neue Scopoletin-Derivate aus *Conyza-Arten.* Phytochemistry 18:1367 https://doi.org/10.1016/0031-9422(79)83024-1
34. Buddrus J, Bauer H, Abu-Mustafa E, Khattab A, Mishaal S, EI-Khrisy EAM, Linscheid M (1985) Foetidin, a sesquiterpenoid coumarin from *Ferula assafoetida.* Phytochemistry 24:869-870 https://doi.org/10.1016/S0031-9422(00)84915-8
35. Cagliotti L, Naef H, Arigoni D, Jeger O (1959) Zur Kenntnis der Sesquiterpene und Azulene 127. Über die Inhaltsstoffe der *Asa foetida* II. Farnesiferol B und C. He1v Chim Acta 42:2557-2570
36. Çiçek Kaya A, Özbek H, Yuca H, Yilmaz G, Bingöl Z, Kazaz C, Gülçin I, Güvenalp Z (2022) Phytochemical analysis and screening of acetylcholinesterase and carbonic anhydrase I and II isoenzymes inhibitory effect of *Heptaptera triquetra* (Vent.) Tutin root. FABAD J Pharm Sci 47:381-392 https://doi.org/10.55262/fabadeczacilik.1147174
37. Dang W, Guo T, Zhou D, Meng Q, Fang M, Chen G, Lin B, Hou Y, Li N (2024) Structure-guided isolation of anti-neuroinflammatory sesquiterpene coumarins from *Ferula sinkiangensis*. Chin J Nat Med 21:1-12 https://doi.org/10.1016/S1875-5364(23)60522-9
38. Dastan D, Salehi P, Gohari AR, Ebrahimi SN, Aliahmadi A, Hamburger M (2014) Bioactive sesquiterpene coumarins from *Ferula pseudalliacea.* Planta Med 80:1118–1123 http://dx.doi.org/10.1055/s-0034-1382996
39. Dastan D, Salehi P, Gohari AR, Zimmermann S, Kaiser M, Hamburger M, Khavasi HR, Ebrahimi SN (2012) Disesquiterpene and sesquiterpene coumarins from *Ferula pseudalliacea*, and determination of their absolute configurations. Phytochemistry 78:170–178. http://dx.doi.org/10.1016/j.phytochem.2012.02.016
40. El-Bassuony AA, Gohar AA, Kabbash AM (2007) Two new sesquiterpene coumarins, ferusinol and samarcandin diastereomer from *Ferula sinaica.* Iran J Pharm Res 6:217-221 https://doi.org/10.22037/ijpr.2010.724
41. Eruçar FM, Kuran FK, Altıparmak Ülbegi G, Özbey S, Karavus¸SN, Arcan GG, Yazıcı Tütünis S, Tan N, Aksoy Sagırlı, P, Miski M (2023) Sesquiterpene coumarin ethers with selective cytotoxic activities from the roots of *Ferula huber-morathii* Pesmen (Apiaceae) and unequivocal determination of the absolute stereochemistry of samarcandin. Pharmaceuticals 16:792 https://doi.org/10.3390/ph16060792
42. Eruçar FM, Senadeera SPD, Wilson JA, Goncharova E, Beutler JA, Miski M (2023) Novel cytotoxic sesquiterpene coumarin ethers and sulfur-containing compounds from the roots of *Ferula turcica*. Molecules 28:5733. https://doi.org/10.3390/molecules28155733
43. Farhadi F, Iranshahi M, Mohtashami L, Shakeri Asil S, Iranshahy M (2021) Metabolic differences of two *Ferula* species as potential sources of galbanum: An NMR-based metabolomics study. Phytochem Anal 2021:1–9. https://doi.org/10.1002/pca.3027
44. Filippini R, Piovan A, Innocenti G, Caniato R, Cappelletti EM (1998) Production of coumarin compounds by *Haplophyllum patavinum in vivo* and *in vitro*. Phytochemistry 49:2337-2340 https://doi.org/10.1016/S0031-9422(98)00356-2
45. Fiorito S, Epifano F, Palmisano R, Genovese S, Taddeo VA (2017) A re-investigation of the phytochemical composition of the edible herb *Amaranthus retroflexus* L. J Pharm Biomed Anal 143:183–187 https://doi-org.proxy.lnu.se/10.1016/j.jpba.2017.05.051
46. Fiorito S, Ianni F, Preziuso F, Epifano F, Scotti L, Bucciarelli T, Genovese S (2019a) UHPLC-UV/Vis

quantitative analysis of hydroxylated and *O*-prenylated coumarins in Pomegranate seed extracts. *Molecules* 24:1963 https://doi.org/10.3390/molecules24101963
47. Fiorito S, Preziuso F, Epifano F, Scotti L, Bucciarelli T, Taddeo VA, Genovese S (2019b) Novel biologically active principles from spinach, goji and quinoa. Food Chem 276:262–265 https://doi-org.proxy.lnu.se/10.1016/j.foodchem.2018.10.018
48. Fiorito S, Genovese S, Palumbo L, Scotti L, Ciulla M, di Profio P, Epifano F. (2020) Umbelliprenin as a novel component of the phytochemical pool from *Artemisia* spp. J Pharm Biomed Anal 184:113205 http://doi.org/10.1016/j.jpba.2020.113205
49. Fiorito, S., Palumbo, L., Epifano, F. Fraternale D, Collevecchio C, Genovese S (2022) Modulation of the biosynthesis of oxyprenylated coumarins in calli from *Ferulago campestris* elicited by ferulic acid. Biomass Conversion and Biorefinery. https://doi.org/10.1007/s13399-022-03309-z
50. Fois B, Distinto S, Meleddu R, Deplano S, Maccioni E, Floris C, Rosa A, Nieddu M, Caboni P, Sissi C, Angeli A, Supuran CT, Cottiglia F (2020) Coumarins from *Magydaris pastinacea* as inhibitors of the tumour-associated carbonic anhydrases IX and XII: isolation, biological studies and in silico evaluation. J Enzyme Inhib Med Chem 35:539-548 https://doi.org/10.1080/14756366.2020.1713114
51. Genovese S, Taddeo VA, Epifano F, Fiorito S, Bize C, Rives A, de Medina P (2017) Characterization of the degradation profile of umbelliprenin, a bioactive prenylated coumarin of a *Ferulago* species. J Nat Prod 80:2424-2431 https://doi.org/10.1021/acs.jnatprod.7b00175
52. Ghannadi A, Fattahian K, Shokoohinia Y, Behbahani M, Shahnoush A (2014) Anti-Viral Evaluation of Sesquiterpene Coumarins from *Ferula assa-foetida* against HSV-1. Iran J Pharma Res 13:523-530
53. Ghosh A, Banerji A, Mandal S, Banerji J (2009) A new sesquiterpenoid coumarin from *Ferula assafoetida.* Nat Prod Commun 4:1023-1024 https://doi.org/10.1177/1934578X0900400801
54. Golovina LA, Khasanov TK, Saidkhodzhaev AI, Malikov VM, Rakhmankulov U (1978) Coumarins and esters of *Ferula microcarpa.* Chem Nat Compd 14:487-490 https://doi.org/10.1007/BF00567136
55. Goren N, Ulubelen A, Oksüz S (1988) A sesquiterpene-coumarin ether and an acetylenic compound from *Tanacetum heterotumum.* Phytochemistry 27:1527-1529 https://doi.org/10.1016/0031-9422(88)80231-0
56. Graf E, Alexa M (1985) Uber 5 neue Umbelliferonether aus Galbanumharz. Planta Med 51:428-431 https://doi.org/10.1055/s-2007-969539
57. Greger H, Hofer O, Nikiforov A (1982a) New sesquiterpene-coumarin ethers from *Achillea* and *Artemisia* species. J Nat Prod 45:455-461 https://doi.org/10.1021/np50022a017
58. Greger H, Haslinger E, Hofer O (1982b) Albartin – a new sesquiterpene-coumarin ether from *Artemisia alba.* Monatsh Chem 113:375-379 https://doi.org/10.1007/BF00799565
59. Greger H, Hofer O, Robien W (1983a) Types of sesquiterpene-coumarin ethers from *Achillea ochroleuca* and *Artemisia tripartita*. Phytochemistry 22:1997-2003 https://doi.org/10.1016/0031-9422(83)80032-6
60. Greger H, Hofer O, Robien W (1983b) New sesquiterpene coumarin ethers from *Achillea ochroleuca.* $^{13}$C NMR of isofraxidin-derived open-chain and bicyclic sesquiterpene ethers. J Nat Prod 46:510-516 https://doi.org/10.1021/np50028a015
61. Greger H, Hofer O (1985) Sesquiterpene-coumarin ethers and polyacetylenes from *Brocchia cinerea*. Phytochemistry 24:85-88 https://doi.org/10.1016/S0031-9422(00)80812-2
62. Guo T, Zhou D, Yang Y, Zhang X, Chen G, Lin B, Sun Y, Ni H, Liu J, Hou Y, Li N (2020) Bioactive sesquiterpene coumarins from the resin of *Ferula sinkiangensis* targeted on over-activation of microglia. Bioorg Chem 104:104338 https://doi.org/10.1016/j.bioorg.2020.104338
63. Guo T, Dang W, Zhou Y, Zhou D, Meng Q, Xu L, Chen G, Lin B, Qing D, Sun Y, Hou Y, Li N (2022) Sesquiterpene coumarins isolated from *Ferula bungeana* and their anti-neuroinflammatory activities. Bioorg Chem 128:106102 https://doi.org/10.1016/j.bioorg.2022.106102
64. Güvenalp Z, Özbek H, Yerdelen KÖ, Yilmaz G, Kazaz C, Demirezer LÖ (2017) Cholinesterase inhibition and molecular docking studies of sesquiterpene coumarin ethers from *Heptaptera cilicica*. Rec Nat Prod 11:462-467 http://doi.org/10.25135/rnp.58.17.03.051
65. Hofer O, Greger H (1984a) Scopoletin sesquiterpene ethers from *Artemisia persica*. Phytochemistry 23:181-182 https://doi.org/10.1016/0031-9422(84)83105-2
66. Hofer O, Greger H (1984b) Naturally occurring sesquiterpene-coumarin ethers, VI. New sesquiterpene-isofraxidin ethers from *Achillea depressa*. Monatsh Chem 115:477-483 https://doi.org/10.1007/BF00810009
67. Hofer O, Widhalm M, Greger H (1984c) Circular dichroism of sesquiterpene-umbelliferone ethers and structure elucidation of a new derivative isolated from the gum resin "Asa Foetida". Monatsh Chem 115:1207-1218 https://doi.org/10.1007/BF00809352
68. Hofer O, Greger H (1985) New sesquiterpene-coumarin ethers from *Anthemis cretica.* Liebigs Ann Chem 1985:1136-1144 https://doi.org/10.1002/jlac.198519850604
69. Iranshahi M, Amin G, Shafiee A (2004) A new coumarin from *Ferula persica.* Pharm Biol 42:440-442 https://doi.org/10.1080/13880200490886102

70. Iranshahi M, Arfa P, Ramezani M, Jaafari MR, Sadeghian H, Bassarello C, Piacente S, Pizza C (2007) Sesquiterpene coumarins from *Ferula szowitsiana* and *in vitro* antileishmanial activity of 7-prenyloxycoumarins against promastigotes. Phytochemistry 68:554-561 https://doi.org/10.1016/j.phytochem.2006.11.002
71. Iranshahi M, Mojaraba M, Sadeghian H, Hanafi-Bojda MY, Schneider B (2008) Polar secondary metabolites of *Ferula persica* roots. Phytochemistry 69:473–478 https://doi.org/10.1016/j.phytochem.2007.08.001
72. Iranshahi M, Kalategi F, Sahebkar A, Sardashti A, Schneider B (2010a) New sesquiterpene coumarins from the roots of *Ferula flabelliloba.* Pharm Biol 48:217–220 https://doi.org/10.3109/13880200903019226
73. Iranshahi M, Masullo M, Asili A, Hamedzadeh A, Jahanbin B, Festa M, Capasso A, Piacente S (2010b) Sesquiterpene coumarins from *Ferula gummosa.* J Nat Prod 73:1958-1962 https://doi.org/10.1021/np100487j
74. Iranshahi M, Barthomeuf C, Bayet-Robert M, Chollet P, Davoodi D, Piacente S, Rezaee R, Sahebkar A (2014) Drimane-type sesquiterpene coumarins from *Ferula gummosa* fruits enhance doxorubicin uptake in doxorubicin-resistant human breast cancer cell line. J Tradit Comp Med 4:118-1256 https://doi.org/10.4103/2225-4110.126181
75. Jabrane A, Jannet HB, Mighri Z, Mirjolet J-F, Duchamp O, Harzallah-Skhiri F, Lacaille-Dubois M-A (2010) Two new sesquiterpene derivatives from the Tunisian endemic *Ferula tunetana* POM. Chem Biodivers 7:392-399 https://doi.org/10.1002/cbdv.200900025
76. Jandl B, Hofer O, Kalchhauser H, Greger H (1997) Open chain sesquiterpene coumarin ethers and coniferylalcohol-4-O-farnesyl ether from *Achillea ochroleuca*. Nat Prod Lett 11:17-24 http://dx.doi.org/10.1080/10575639708043752
77. Kadyrov AS, Saidkhodzhaev AI, Nikonov VM (1975) The structures of feroside and of reoselin A - new glycosides from the roots of *Ferula korshinskyi*. Chem Nat Compd 11:604-608 https://doi.org/10.1007/BF00567694
78. Kadyrov AS, Saidkhodzhaev AI, Malikov VM (1978) The structure of foliferin. Chem Nat Compd 14:442-443 https://doi-org.proxy.lnu.se/10.1007/BF00565259
79. Kamilov KM, Nikonov GK (1973) Coumarins of *Ferula kopetdaghensis* and the structure of kopeolin and kopeoside. Chem Nat Compd 9:294-298 https://doi-org.proxy.lnu.se/10.1007/BF00565684
80. Kamilov KM, Nikonov GK (1974) Coumarins of *Ferula kopetdaghensis* - kopetdaghin and kopeodin (farnesiferol B). Chem Nat Compd 10:450-453 https://doi-org.proxy.lnu.se/10.1007/BF00563804
81. Kamoldinov K, Li J, Eshbakova K, Sagdullaev S, Xu G, Zhou Y, Li J, Aisa HA (2021) Sesquiterpene coumarins from *Ferula samarkandica* Korovin and their bioactivity. Phytochemistry 187:112705 https://doi.org/10.1016/j.phytochem.2021.112705
82. Kang X, Wu L, Zhao C, Zhang C, Wang Q (2024) A new sesquiterpene coumarin from *Ferula bungeana* Kitagawa. Nat Prod Res 2024:1-5 https://doi.org/10.1080/14786419.2024.2332490
83. Khaliova EK, Saidkhodzhaev AI (1998) Terpenoid coumarins of *Ferula sumbul.* Chem Nat Compd 34:506-507 https://doi-org.proxy.lnu.se/10.1007/BF02329608
84. Khasanov TKh, Saidkhodzhaev AI, Nikonov GK (1974a) Structure and configuration of polyanthin and polyanthinin - new coumarins from the roots of *Ferula polyantha.* Chem Nat Compd 10:523-524 https://doi-org.proxy.lnu.se/10.1007/BF00563828
85. Khasanov TKh, Saidkhodzhaev AI, Nikonov GK (1974b) Structure and configuration of new coumarins of the roots of *Ferula mogoltavica.* Chem Nat Compd 10:8-11 https://doi-org.proxy.lnu.se/10.1007/BF00568209
86. Khasanov TKh, Saidkhodzhaev AI, Nikonov GK (1976) Structure of feropolin, feropolol, feropolone and feropolidin. Chem Nat Compd 12:79-80 https://doi-org.proxy.lnu.se/10.1007/BF00570195
87. Khasanov TKh, Malikov VM, Melibaev S (1979) Structure and configuration of tavimolidin. Chem Nat Compd 15:417-419 https://doi-org.proxy.lnu.se/10.1007/BF00565036
88. Kir'yalov NP (1961) The structure of cocanicine and umbelliprenin, components of the neutral part of oil from *Ferula cocanica.* Tr Bot Inst Akad Nauk SSSR Ser 5:8
89. Kir'yalov NP, Movchan SD (1968) The structure of samarcandin and samarcandone, coumarin compounds from *Ferula samarcandica.* Chem Nat Compd 4:63-65 https://doi-org.proxy.lnu.se/10.1007/BF00568012
90. Kir'yalov NP, Bukreeva TV (1972) Samarcandin acetate from the roots of *Ferula pseudooreoselinum.* Chem Nat Compd 8:777-778 https://doi-org.proxy.lnu.se/10.1007/BF00564609
91. Kir´yalov NP (1980) Dihydroconferin from *Ferula tadshikorum*. In Kir´yalov NP, Sklar YE (eds) Chemistry of Natural Sciences 1:122-123. (in Russian).
92. Kir'yanova IA, Sklyar YuE (1984) Ferucrin isobutyrate and ferucrinone from *Ferula foetidissima.* Chem Nat Compd 20:617-618 https://doi-org.proxy.lnu.se/10.1007/BF00580081
93. Kisiel W, Stojakowska A (1997) A sesquiterpene coumarin ether from transformed roots of *Tanacetum parthenium.* Phytochemistry 46:515-516 https://doi.org/10.1016/S0031-9422(97)87091-4
94. Kuliev ZA, Khasanov TKh (1978a) Structures of ferocaulin, ferocaulinin, ferocaulidin, and ferocaulicin.

Chem Nat Compd 14:267-271 https://doi-org.proxy.lnu.se/10.1007/BF00713313
95. Kuliev ZA, Khasanov TKh (1978b) Structures of cauferin and cauferidin. Chem Nat Compd 14:271-274 https://doi-org.proxy.lnu.se/10.1007/BF00713314
96. Kuliev ZA, Khasanov TKh, Malikov VM (1979a) Structure and configuration of cauferinin. Chem Nat Compd 15:127-129 https://doi-org.proxy.lnu.se/10.1007/BF00570778
97. Kuliev ZA, Khasanov TKh, Malikov VM (1979b) Terpenoid coumarin glycosides of *Ferula conocaula*. Chem Nat Compd 15:414-416 https://doi-org.proxy.lnu.se/10.1007/BF00565035
98. Kuliev ZA, Khasanov TKh, Malikov VM (1982) Coumarins of *Ferula conocaula*. Chem Nat Compd 18:114-115 https://doi-org.proxy.lnu.se/10.1007/BF00581611
99. Lee Ch-L, Chiang L-Ch, Cheng L-H, Liaw Ch-Ch, Abd El-razek MH, Chang F-R, Wu Y-Ch (2009) Influenza A ($H_1N_1$) antiviral and cytotoxic agents from *Ferula assa-foetida.* J Nat Prod 72:1568-1572 https://doi.org/10.1021/np900158f
100. Li G, Li X, Cao L, Zhang L, Shen L, Zhu J, Wang J, Si J (2015a) Sesquiterpene coumarins from seeds of *Ferula sinkiangensis.* Fitoterapia 103:222–226. http://dx.doi.org/10.1016/j.fitote.2015.03.022
101. Li G, Wang J, Li X, Cao L, Lv N, Chen G, Zhu J, Si J (2015b) Two new sesquiterpene coumarins from the seeds of *Ferula sinkiangensis.* Phytochem Lett 13:123–126 https://doi.org/10.1016/j.phytol.2015.06.002
102. Li G, Wang J, Li X, Cao L, Gao L, Lv N, Si J (2016) An unusual sesquiterpene coumarin from the seeds of *Ferula sinkiangensis*. J Asian Nat Prod Res 18:891-896 https://doi.org/10.1080/10286020.2016.1168813
103. Li K-M, Dong X, Ma Y-N, Wu Z-H, Yan Y-M, Cheng Y-X (2019) Antifungal coumarins and lignans from *Artemisia annua*. Fitoterapia 134:323-328 https://doi.org/10.1016/j.fitote.2019.02.022
104. Madureira AM, Molnar A, Abreu PM, Molnar J, Ferreira MU (2004) A new sesquiterpene coumarin ether and a new abietane diterpene and their effects as inhibitors of P-glycoprotein. Planta Med 70:828-833 https://doi.org/10.1055/s-2004-827231
105. Marco JA, Sanz JF, Yuste A, Rustaiyan A (1991) New umbelliferone sesquiterpene ethers from roots of *Ligularia persica*. Liebigs Ann Chem 1991:929-931 https://doi.org/10.1002/jlac.1991199101158
106. Mbah JA, Gatsing D, Efange SMN (2010) Antibacterial agents from the seeds of *Peucedanum zenkeri* L. (Umbelliferae). Pak J Med Sci 26:314-318
107. Miski M, Ulubelen A, Lee E, Mabry TJ (1985) Sesquiterpene-coumarin ethers of *Ferula tingitana*. J Nat Prod 48:326–327 https://doi.org/10.1021/np50038a024
108. Miski M, Tosun F, Aytar EC, Duran A (2015) Novel sesquiterpene coumarin ethers from the dichloromethane extract of the roots of *Heptaptera cilicica.* Conference paper OP-22, 11th Interntional Symposium on the Chemistry of Natural Compounds, 1-6 October, 2015, Antalya, Turkey
109. Mohamed TA, Elshamy AI, Ibrahim MAA, Zellagui A, Moustafa MF, Abdelrahman AHM, Ohta S, Pare PW, Hegazy M-EF (2020) Carotane sesquiterpenes from *Ferula vesceritensis*: in silico analysis as SARS-CoV-2 binding inhibitors. RSC Adv 10:34541–34548 https://doi.org/10.1039/d0ra06901a
110. Murch SJ, Rupasinghe HPV, Goodenowe D, Saxena PK (2004) A metabolomic analysis of medicinal diversity in Huang-qin (*Scutellaria baicalensis* Georgi) genotypes: discovery of novel compounds. Plant Cell Rep 23:419–425 https://doi.org/10.1007/s00299-004-0862-3
111. Nabiev AA, Khasanov TKh, Malikov VM (1978) New terpenoid coumarins of *Ferula kopetdaghensis*. Chem Nat Compd 14:440 https://doi-org.proxy.lnu.se/10.1007/BF00565257
112. Nabiev AA, Khasanov TKh, Malikov VM (1979) A chemical study of the roots of *Ferula kopetdagensis*. Chem Nat Compd 15:14-16 https://doi-org.proxy.lnu.se/10.1007/BF00570841
113. Nabiev AA, Khasanov TKh, Malikov VM (1982a) Terpenoid coumarins of *Ferula kokanica.* Chem Nat Compd 18:547-549 https://doi-org.proxy.lnu.se/10.1007/BF00575034
114. Nabiev AA, Khasanov TKh, Malikov VM (1982b) A new terpenoid coumarin from *Ferula kopetdaghensis*. Chem Nat Compd 18:44-46 https://doi-org.proxy.lnu.se/10.1007/BF00581594
115. Nabiev AA, Malikov VM (1983a) Microlobin a new coumarin from *Ferula microloba.* Chem Nat Compd 19:664–667 https://doi-org.proxy.lnu.se/10.1007/BF00575163
116. Nabiev AA, Malikov VM (1983b) Microlobidene, a terpenoid coumarin from *Ferula microloba* with a new type of carbon skeleton. Chem Nat Compd 19:743–744 https://doi-org.proxy.lnu.se/10.1007/BF00575187
117. Nabiev AA, Malikov VM Khasanov TKh (1983c) Karatavicin - a new coumarin from *Ferula karatavica.* Chem Nat Compd 19:498 https://doi-org.proxy.lnu.se/10.1007/BF00575721
118. Oughlissi-Dehak K, Lawton P, Michalet S, Bayet C, Darbour N, Hadj-Mahammed M, Badjah-Hadj-Ahmed YA, Dijoux-Franca M-G, Guilet D (2008) Sesquiterpenes from aerial parts of *Ferula vesceritensis*. Phytochemistry 69:1933-1938 https://doi.org/10.1016/j.phytochem.2008.03.010
119. Paknikar SK, Kirtany JK (1974) The terpenoid parts of kamolone and kamolol of *Ferula penninervis*. Experientia 30:224-225 https://doi.org/10.1007/BF01934792
120. Pavlovic I, Krunic A, Nikolic D, Radenkovic M, Brankovic S, Niketic M, Petrovic S (2014) Chloroform extract of underground parts of *Ferula heuffelii*: Secondary metabolites and spasmolytic activity. Chem Biodiver 11:1417-1427 https://doi.org/10.1002/cbdv.201400094

121. Perel'son ME, Vandyshev VV, Sklyar YE (1975) The structure and stereochemistry of conferin. Chem Nat Compd **11**, 252–253 https://doi.org/10.1007/BF00570683
122. Perel'son ME, Bandyshev VV, Sklyar YE, Vezhkhovska-Renke K, Veselovskaya NV, Pimenov MG (1976) New terpenoid coumarins from *Ferula tadshikorum.* Chem Nat Compd 12:533-537 https://doi-org.proxy.lnu.se/10.1007/BF00565176
Beselovskaya NV, Sklyar YE (1984) Deacetyltadzhikorin from *Ferula tadshikorum*. Chem Nat Compd 20:363 https://doi.org/10.1007/BF00575772
123. Perel'son ME, Sokolova AI, Sklyar YE (1978) The structure of feterin - a new terpenoid coumarin from *Ferula teterrima*. Chem Nat Compd 14:263-267 https://doi-org.proxy.lnu.se/10.1007/BF00713312
124. Pinar M, Rodriguez B (1977) A new coumarin from *Ferula loscosii* and the correct structure of colladonin. Phytochemistry 16:1987-1989 https://doi.org/10.1016/0031-9422(77)80109-X
125. Rassouli FB, Matin MM, Iranshahi M, Bahrami AR, Neshati V, Mollazadeh S, Neshati Z (2009) Mogoltacin enhances vincristine cytotoxicity in human transitional cell carcinoma (TCC) cell line**.** Phytomedicine 16**:**181-187 https://doi.org/10.1016/j.phymed.2008.06.011
126. Rosselli S, Maggio A, Bellone G, Formisano C, Basile A, Cicala C, Alfieri A, Mascolo N, Bruno M (2007) Antibacterial and anticoagulant activities of coumarins isolated from the flowers of *Magydaris tomentosa.* Planta Med 73:116-120 https://doi.org/10.1055/s-2006-951772
127. Sagitdinova GV, Saidkhodzhaev AI, Malikov VM (1983) Structure and stereochemistry of the coumarins of *Ferula lehmannii*. Chem Nat Compd 19:672-675 https://doi-org.proxy.lnu.se/10.1007/BF00575165
128. Saidkhodzhaev AI, Nikonov GK (1973) The configuration of badrakemin and gummosin, and the identity of isobadrakemin, colladonin and farnesiferol A. Chem Nat Compd 9:462-464 https://doi-org.proxy.lnu.se/10.1007/BF00568629
129. Saidkhodzhaev AI, Malikov VM (1978a) The stereochemistry of terpenoid coumarins. Chem Nat Compd 14:601-605 https://doi-org.proxy.lnu.se/10.1007/BF00937607
130. Saidkhodzhaev AI, Malikov VM (1978b) The stereochemistry of feropolol, feropolin, feropolone, and feropolidin. Chem Nat Compd 14:681–682 https://doi.org/10.1007/BF00937630
131. Saidkhodzhaev AI, Kadyrov ASh, Malikov VM (1979) Stereochemistry of feshurin, nevskin, and colladocin. Chem Nat Compd 15:266-268 https://doi-org.proxy.lnu.se/10.1007/BF00566071
132. Sarker SD, Nahar L, Rahman MM, Siakalima M, Middleton M, Byres M, Kumarasamy Y, Murphy E (2005) Bioactivity of umbelliprenin, the major component found in the seeda of *Angelica sylvestris*. Ars Pharm 46:35-41
133. Scotti L, Genovese S, Bucciarelli T, Martini F, Epifano F, Fiorito S, Preziuso F, Taddeo VA (2018) Analysis of biologically active oxyprenylated phenylpropanoids in tea tree oil using selective solid-phase extraction with UHPLC-PDA detection. J Pharm Biomed Anal 154:174–179 https://doi-org.proxy.lnu.se/10.1016/j.jpba.2018.03.004
134. Shahzadi I, Ali Z, Baek SH, Mirza, B Ahn KS (2020) Assessment of the antitumor potential of umbelliprenin, a naturally occurring sesquiterpene coumarin. Biomedicines 8:126 https://doi.org/10.3390/biomedicines8050126
135. Shakeri A, Iranshahy M, Iranshahi M (2014) Biological properties and molecular targets of umbelliprenin—A mini-review. J Asian Nat Prod Res 16:884–889 https://doi.org/10.1080/10286020.2014.917630
136. Shomirzoeva O, Xu M-Y, Sun Z-J, Li C, Nasriddinov A, Muhidinov Z, Zhang K, Gu Q, Xu J (2021) Chemical constituents of *Ferula seravschanica.* Fitoterapia 149:104829 https://doi.org/10.1016/j.fitote.2021.104829
137. Sidana J, Saini V, Dahiya S, Nain P, Bala S (2013) A review on Citrus – "The boon of nature". Int J Pharm Sci Rev Res 18:20-27
138. Steck W, Bailey BK (1969) Leaf coumarins of *Angelica archangelical.* Can J Chem 47:2425-2430 https://doi.org/10.1139/v69-396
139. Su B-N, Takaishi Y, Honda G, Itoh M, Takeda Y, Kodzhimatov OK, Ashurmetov O (2000) Sesquiterpene Coumarins and Related Derivatives from *Ferula pallida.* J Nat Prod 63:436-440 https://doi.org/10.1021/np990262i
140. Sutthivaiyakit S, Mongkolvisut W, Prabpai S, Kongsaeree P (2009) Diterpenes, sesquiterpenes, and a sesquiterpene-coumarin conjugate from *Jatropha integerrima.* J Nat Prod 72:2024–2027 https://doi.org/10.1021/np900342b
141. Taddeo VA, Epifano F, Preziuso F, Fiorito S, Caron N, Rives A, de Medina P, Poirot M, Sylvente-Poirot S, Genovese S (2019) HPLC analysis and skin whitening effects of umbelliprenin-containing extracts of *Anethum graveolens*, *Pimpinella anisum*, and *Ferulago campestris*. Molecules 24:501 https://doi.org/10.3390/molecules24030501
142. Taniguchi M, Yokota O, Shibano M, Wang N-H, Baba K (2005) Four coumarins from *Heracleum yunngningense*. Chem Pharm Bull 53:701-704 https://doi.org/10.1248/cpb.53.701
143. Tashkhodzhaev B, Turgunov KK, Izotova LY, Kamoldinov KS (2015) Stereochemistry of samarcandin-

type sesquiterpenoid coumarins. Crystal structures of feshurin and nevskin. Chem Nat Compd 51:242-246 http://dx.doi.org/10.1007/s10600-015-1253-4
144. Teng L, Ma GZ, Li L, Ma LY, Xu XQ (2013) Karatavicinol A, a new anti-ulcer sesquiterpene coumarin from *Ferula sinkiangensis.* Chem Nat Compd 49:606-609 https://doi-org.proxy.lnu.se/10.1007/s10600-013-0690-1
145. Tian Y-Q, Zhang Z-X, Xu H-H (2013) Laboratory and field evaluations on insecticidal activity of *Cicuta virosa* L. var. *latisecta* Celak. Ind Crops Prod 41:90-93 https://doi.org/10.1016/j.indcrop.2012.04.015
146. Tosun F, Beutler JA, Ransom TT, Miski M (2019) Anatolicin, a highly potent and selective cytotoxic sesquiterpene coumarin from the root extract of *Heptaptera anatolica.* Molecules 24:1153 http://dx.doi.org/10.3390/molecules24061153
147. Tosun F, Aytar EC, Beutler JA, Wilson JA, Miski M (2021) Cytotoxic sesquiterpene coumarins from the roots of *Heptaptera cilicica.* Rec Nat Prod 15:529-536 http://doi.org/10.25135/rnp.242.21.02.1990
148. Tosun F, Beutler JA, Miski M (2023) Coumarins from the dichloromethane root extract of *Haptaptera triquetra* and their cytotoxic activities. Rec Nat Prod 17:998-1005 http://doi.org/10.25135/rnp.415.2307.2846
149. Tuncay HO, Akalın E, Doğru-Koca A, Eruçar FM, Miski M (2023) Two new *Ferula* (Apiaceae) species from central Anatolia: *Ferula turcica* and *Ferula latialata*. Horticulturae 9:144 https://doi.org/10.3390/horticulturae9020144
150. Vandyshev VV, Sklyar YE, Perel'son ME, Moroz MD, Pimenov MG (1972a) Conferol, a New Coumarin from the Roots of *Ferula conocaula* and *F. moschata.* Khim Prir Soedin 670 https://doi.org/10.1007/BF00564346
151. Vandyshev VV, Sklyar YE, Perel'son ME, Moroz MD, Pimenov MG (1972b) Conferone, a New Terpenoid Coumarin from the Fruit of *Ferula conocaula*. Khim Prir Soedin 669 https://doi.org/10.1007/BF00564345
152. Vandyshev VV, Sklyar YE, Perel'son ME, Moroz MD (1974) Conferdione a new coumarin from *Ferula conocaula.* Chem Nat Compd 10:670-671 https://doi.org/10.1007/BF00567873
153. Veselovskaya NV, Sklyar YE, Fesenko DA, Pimenov MG (1979) Fekrol, a new terpenoid coumarin from *Ferula krylovii.* Chem Nat Comp 15:755-756 https://doi.org/10.1007/BF00565585
154. Veselovskaya NV, Sklyar YE, Savina AA (1981) Fekrynol and its acetate from *Ferula krylovii.* Chem Nat Compd 17:589-590 https://doi-org.proxy.lnu.se/10.1007/BF00574389
155. Vuckovic I, Trajkovic V, Macura S, Tesevic V, Janackovic P, Milosevljevic S (2007) A novel cytotoxic lignan from *Seseli annuum* L. Phytother Res 21:790–792 https://doi.org/10.1002/ptr.2152
156. Wang H, Liu Y, Wang Y, Xu T, Xia G, Huang X (2023) Umbelliprenin induces autophagy and apoptosis while inhibits cancer cell stemness in pancreatic cancer cells. Cancer Med 12:15277–15288 https://doi.org/10.1002/cam4.6170
157. Wang J, Wang H, Zhang M, Li X, Zhao Y, Chen G, Si J, Jiang L (2020) Sesquiterpene coumarins from *Ferula sinkiangensis* KM Shen and their cytotoxic activities. Phytochemistry 180:112531 https://doi.org/10.1016/j.phytochem.2020.112531
158. Wang J, Huo X, Wang H, Dong A, Zheng Q, Si J (2023) Undescribed sesquiterpene coumarins from the aerial parts of *Ferula sinkiangensis* and their anti-inflammatory activities in lipopolysaccharide-stimulated RAW 264.7 macrophages. Phytochemistry 210:113664 https://doi.org/10.1016/j.phytochem.2023.113664
159. Xing Y, Li N, Zhou D, Chen G, Jiao K, Wang W, Si Y, Hou Y (2017) Sesquiterpene coumarins from *Ferula sinkiangensis* act as neuroinflammation inhibitors. Planta Med 83:135-142 http://dx.doi.org/10.1055/s-0042-109271
160. Yang J-R, An Z, Li Z-H, Jing S, Qin H-L (2006) Sesquiterpene coumarins from the roots of *Ferula sinkiangensis* and *Ferula teterrima*. Chem Pharm Bull 54:1595-1598 https://doi.org/10.1248/cpb.54.1595
161. Yrjönen T, Eeva M, Kauppila TJ, Martiskainen O, Summanen J, Vuorela P, Vuorela H (2016) Profiling of coumarins in *Peucedanum palustre* (L.) Moench populations growing in Finland. Chem Biodiv 13:700–709 https://doi.org/10.1002/cbdv.201500198
162. Zhai D-D, Zhong J-J (2010) Simultaneous analysis of three bioactive compounds in *Artemisia annua* hairy root cultures by reversed-phase high performance liquid chromatography–diode array detector. Phytochem Anal 21:524–530 http://dx.doi.org/10.1002/pca.1226
163. Zhang M-M,, Kamoldinov K, Nueraihemaiti M, Turdu G, Zou G-A, Aisa HA (2024) Sesquiterpene coumarins with anti-vitiligo and cytotoxic activities from *Ferula samarkandica*, Phytochem Lett 61:21-28 https://doi-org.proxy.lnu.se/10.1016/j.phytol.2024.03.004
164. Zhou D, Li N, Zhang Y, Yan C, Jiao K, Sun Y, Ni H, Lin B, Hou Y (2016) Biotransformation of neuro-inflammation inhibitor kellerin by *Angelica sinensis* (Oliv.) Diels callus. RSC Adv 6:97302-97312 http://dx.doi.org/10.1039/C6RA22502K
